# A Portrait of a Young Fish: Redescription of *Pteronisculus gunnari* (Nielsen, 1942) from a juvenile specimen from the Early Triassic of East Greenland, with implications for ontogenetic development in early actinopterygians

**DOI:** 10.1101/2024.07.06.598961

**Authors:** Iacopo Cavicchini, Thodoris Argyriou, Vincent Fernandez, Katheen Dollman, Sam Giles

## Abstract

The Early–Middle Triassic actinopterygian genus *Pteronisculus* (White, 1933) is part of the Triassic Early Fish Fauna (TEFF), a cosmopolitan group of taxa that thrived in the aftermath of the end-Permian mass extinction. *Pteronisculus* is considered an important non-neopterygian outgroup taxon in many works dealing with the interrelationships of early crown actinopterygians, but the phylogenetic relationships of many of TEFF genera are debated, with the topology of the lineages giving rise to crown actinopterygians consequently unclear. This is despite exceptional, three-dimensionally preservation of an abundance of fossils associated with TEFF fishes. *P. gunnari*, from the Induan (Early Triassic) Kap Stosch Formation, East Greenland, is known in less detail than other species of the genus. Here, we use X-ray micro-computed tomography to comprehensively redescribe the three-dimensionally preserved holotype of *P. gunnari*, including a detailed description of the internal anatomy. The specimen shows morphological features previously undescribed for the genus, including paired premaxillae, medially-directed teeth on the palate, canals for the buccohypophysial canal and internal carotids piercing the parasphenoid, and numerous parotic toothplates. Scale covering is complete, but the braincase and palatoquadrate are ossified as multiple elements, indicating that the specimen was not fully mature and allowing new insights into ossification patterns and ontogenetic development of non-neopterygian actinopterygians. These new anatomical data enrich our understanding of both the morphological complexity and the interrelationships of actinopterygians from the Triassic Early Fish Fauna.

## Introduction

During the Early Triassic (252-247 Ma) a group of ray-finned fish taxa attained a wide distribution and left an abundant—and often exceptionally preserved—fossil record (Brinkmann et al., 2010). The coherence of this fauna first became apparent in the 1970s (Schaeffer & Mangus, 1976), prompting the naming of the Triassic Early Fish Fauna (TEFF) (Romano, 2021; Tintori et al., 2014). The TEFF includes taxa with debated phylogenetic placements, but generally considered non-neopterygians (e.g., *Saurichthys*, *Birgeria*, *Australosomus, Boreosomus, Pteronisculus*), possible stem neopterygians (*Bobasatrania*) and crown group neopterygians (e.g., *Watsonulus*) (T. Argyriou et al., 2018; Brinkmann et al., 2010; Friedman, 2015; Brian G Gardiner et al., 2005; López-Arbarello & Sferco, 2018; Olsen, 1984; Romano, 2021; Tintori et al., 2014; Xu, Gao, et al., 2014). Surprisingly, given the phylogenetic uncertainties, many of the boreal TEFF taxa are among the best known fossil actinopterygians, due to detailed monographic descriptions (Nielsen, 1949; Stensio, 1925; Stensiö, 1921, 1932). This cosmopolitan and diverse fauna appears to thrive in the immediate aftermath of the Permo-Triassic mass extinction (PTME), a catastrophe that wiped out up to 96% of all species (Chen & Benton, 2012; Erwin et al., 2002), but how this event has affected actinopterygians is still unclear.

The abrupt rise and diversification of the TEFF could represent a genuine recovery fauna, or an artefact related to the very poor late Permian actinopterygian fossil record (Friedman & Sallan, 2012; and Henderson et al., 2023; but see Henderson et al., 2022; Smithwick & Stubbs, 2018; Vázquez & Clapham, 2017). Some TEFF genera, like *Bobasatrania*, *Saurichthys* and possibly *Pteronisculus*, are indeed known from the late Permian (Brian George Gardiner & Jubb, 1975; Tintori et al., 2014), but broader macroevolutionary patterns are the subject of ongoing debate (Friedman, 2015; Romano, 2021; Romano et al., 2016; Smithwick & Stubbs, 2018).

The genus *Pteronisculus* was originally erected by White (1933), who described two new species of actinopterygians from the Early Triassic of north-eastern Madagascar (*P. macropterus* and *P. cicatrosus*, the latter being the type species). A few years later, *Glaucolepis* (Stensiö, 1921), initially erected to contain incomplete fossil fish material from the Early Triassic of Svalbard, was synonymized with *Pteronisculus* (White & Moy-Thomas, 1940). The temporal and geographical range of this genus grew with successive descriptions of new species from the Early Triassic of east Greenland (Nielsen, 1942), north-western Madagascar (Lehman, 1952), Nevada, USA (Romano et al., 2019) and the Middle Triassic of Luoping, China (Ren & Xu, 2021; Xu, Shen, et al., 2014). Fragmentary material has also been referred to the genus from the late Permian of South Africa (Brian George Gardiner & Jubb, 1975) and the Early Triassic of Alberta and British Columbia, Canada (Lambe, 1916; Schaeffer & Mangus, 1976). The anatomy of *Pteronisculus* is known in great detail, especially from five species from the Early Triassic of East Greenland (Nielsen 1942) and four from north-western Madagascar (Lehman 1952). This taxon was originally considered a member of the ‘Palaeonisciformes’, an assemblage of taxa of generalized appearance now known to be paraphyletic (Gardiner & Schaeffer, 1989); in this work we use the term “palaeopterygian” to avoid historical associations of monophyly (see Friedman & Giles, 2016 and references therein). *Pteronisculus* is routinely included in phylogenetic analyses of crown neopterygians, either as a formal outgroup or as a representative of non-neopterygian morphology more generally (Brian G Gardiner et al., 2005; Hurley et al., 2007; López-Arbarello & Sferco, 2018; Olsen, 1984; Xu, Gao, et al., 2014). In more targeted investigations of early actinopterygian relationships, *Pteronisculus* has fluctuated between the actinopterygian and neopterygian stem (e.g., Lund, Poplin & McCarty, 1995; Cloutier & Arratia, 2004 and references therein; Hurley et al., 2007; Xu, Gao & Finarelli, 2014, etc.). More recently, it has been consistently recovered as a stem actinopterygian (but see Figueroa et al., 2019), although with limited agreement as to its immediate sister taxa or position relative to the actinopterygian crown node: as sister taxon to the Carboniferous *Cosmoptychius* and Bear Gulch actinopterygians (Giles et al., 2017); in a clade with *Cyranorhis*, which is resolved as the sister taxon to a clade comprising the Triassic *Boreosomus*, *Cosmoptychius*, *Mesopoma* and a subset of Bear Gulch taxa (Ren & Xu, 2021), or as sister to *Beagiascus* and a heterogeneous group of Carboniferous (*Cosmoptychius*, *Kansasiella*, *Coccocephalichthys*, *Lawrenciella*), Permian (*Luederia*), and Early Triassic genera (*Boreosomus*) (Thodoris Argyriou et al., 2022).

Although 13 species of *Pteronisculus* are described in the literature (plus two to three more dubious attributions; see e.g., Brian George Gardiner & Jubb, 1975; Romano et al., 2019; Schaeffer & Mangus, 1976), only Ren & Xu (2021) and Caron et al. (2023) have included more than one taxon in a phylogenetic analyses, and the latter only consider neurocranial characters. In large part, this can be attributed to the uneven amount of information available for each taxon. *P. stensioi* and *P. magnus* from East Greenland have been described in great detail from 3D material (Nielsen, 1942), and as are most commonly used in phylogenetic analyses. The dermal skeleton of the Malagasy species is well described, but due to preservational biases the braincase is only described in *P. cicatrosus* (Beltan, 1996; Lehman, 1952). The recently described Middle Triassic Chinese species *P. nielseni* and *P. changae*, and the Early Triassic *P. nevadanus* from Nevada, have been described based upon several complete but flattened specimens, with the result that the dermal skeleton is almost completely known but internal anatomy is lacking (Ren & Xu, 2021; Romano et al., 2019; Xu, Shen, et al., 2014). *P. gyrolepidoides* is known from very fragmentary and incomplete remains (Stensiö, 1921). *P. arcticus*, *P. aldingeri*, and *P. gunnari* are also known from three-dimensional material and are referred to throughout Nielsen’s (1942) monograph, but are rarely described in their own right, mostly relying on comparisons with *P. magnus* or *P. stensioi*.

Here, we use X-ray computed tomography (XCT) to reinvestigate the holotype of *P. gunnari*, previously only partially described by Nielsen (1942). Its 3D preservation allows us to describe a wealth of morphological features previously unknown in the genus, including a small coronoid process formed by the dentary, medially-directed teeth on the palate, an intercalar, and a propterygium fused to the first lepidotrichium. We present an expanded phylogenetic analysis incorporating multiple species of *Pteronisculus*, confirming its monophyly. Its position nested deep within a clade with Carboniferous origins suggests it may be unsuitable for representing the morphology of generalized early actinopterygians.

## Materials and methods

### Fossil material and locality

NHMD–73588 (*ex*-MGUH VP.791), curated in the Natural History Museum of Denmark (Copenhagen), is a complete, articulated specimen including the skull, paired fins, median fins, body and caudal fin, lacking only the extreme tip of the dorsal lobe of the tail (Fig. 1). The cranium in particular is preserved in 3D and almost perfect articulation. The specimen was likely preserved within a concretion and has been extensively prepared to expose the skull in the round and the right side of the body and fins. It was found between 1932 and 1937 in the Kap Stosch area, West locality of the Spath Plateau, East Greenland (Nielsen 1942), in Griesbachian (Induan) strata of the Kap Stosch Formation (Surlyk et al., 2017).

**Figure 1:**
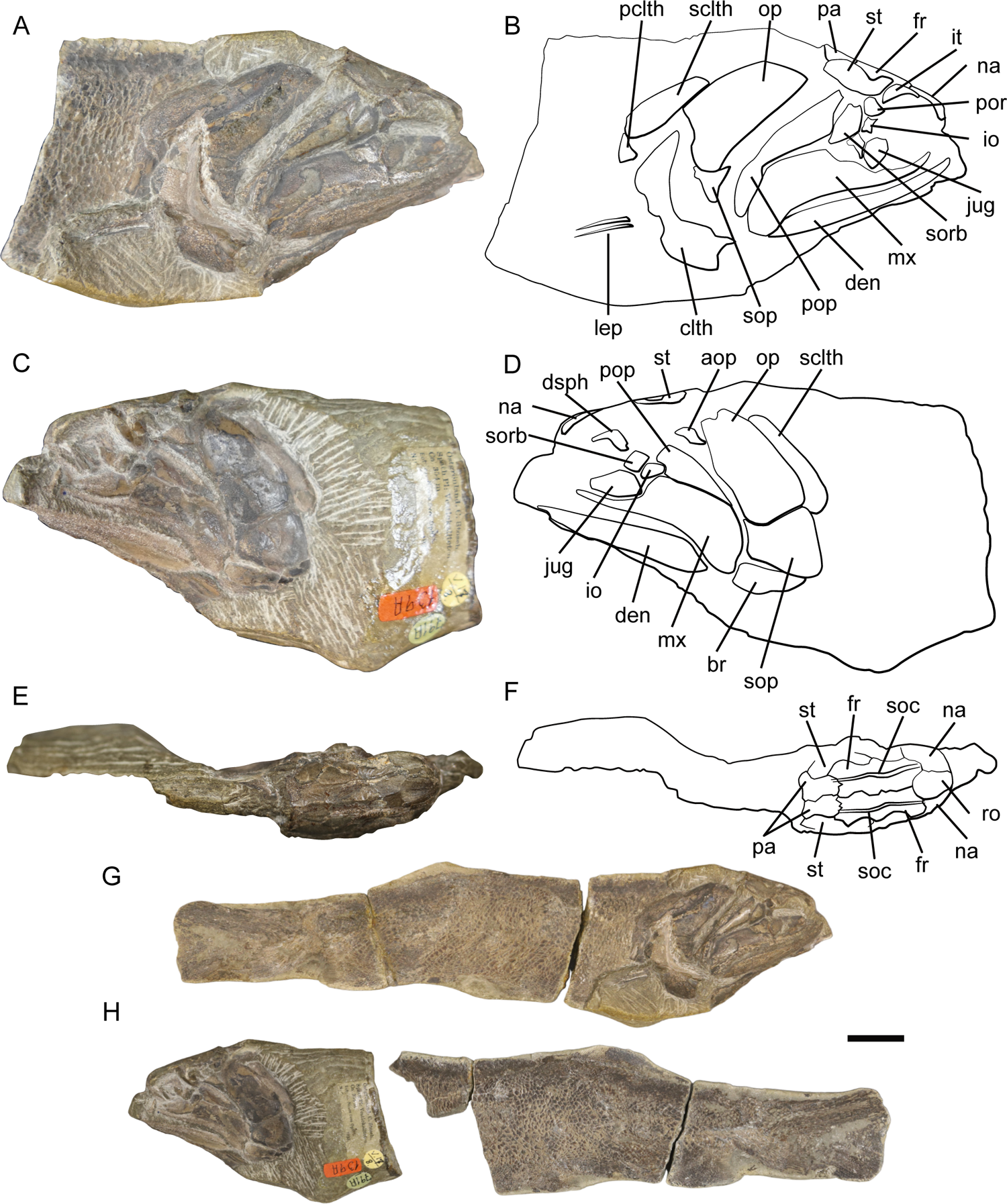
External photos and line drawings of Pteronisculus gunnari NHMD 73588, anterior half. Right lateral view (A) and interpretive line drawing (B). Left lateral view (C) and interpretive line drawing (D). Dorsal view (E) and interpretive line drawing (F). Entire specimen in right lateral (G) and left lateral view (H). Scale bar: 10 mm for A-F, 5 mm for G-H. Abbreviations: aop, accessory operculum; br, branchiostegal ray; clth, cleithrum; dsph, dermosphenotic; den, dentary; fr, frontal; io, infraorbital; it, intertemporal; jug, jugal; lep, lepidotrichia; mx, maxilla; na, nasal; op, operculum; pa, parietal; pclth, postcleithrum; pop, preoperculum; por, postorbital process; ro, rostral; sorb, suborbital; sclth, supracleithrum; soc, supraorbital canal; sop, suboperculum; st, supratemporal.

The fossil comes from Fish Zone III, one of five highly fossiliferous strata that were discovered and sampled intensively in the late 1920s and the 1930s (Nielsen, 1942; Surlyk et al., 2017). The Triassic ichthyofauna is rich and includes actinopterygians (e.g., *Birgeria*, *Saurichthys*, *Australosomus*, *Boreosomus*, *Bobasatrania*), sarcopterygians (e.g., *Laugia*, *Sassenia*, *Whiteia*) and chondrichthyans (e.g., *Polyacrodus*, *Nemacanthus*) (Brinkmann et al., 2010; Stensiö, 1932). NHMD–73588 was described and chosen as the holotype of *Pteronisculus gunnari* by Nielsen (1942), who originally assigned it the specimen field number 18 and carried out mechanical preparation. The specimen itself does not figure in any plates in Nielsen’s monograph, but the dermal bones of the skull roof, plus the rostral and nasals, are figured in a line drawing (Nielsen 1942: fig. 26A, p. 133).

### X-ray Computed Tomography

The anterior portion of NHMD–73588, containing the three-dimensionally preserved cranium, was imaged using X-ray computed tomography at the XTM Facility, Palaeobiology Research Group, University of Bristol using a Nikon XT H 225 ST system. The parameters used are: 224 kV, 191 μA, 1 mm copper filter, and a cone beam magnification resulting in an isotropic voxel size of 22.5 μm. The resulting images were processed using Mimics Materialise software (v.20-23, biomedical.materialise.com/mimics; Materialise, Leuven, Belgium) to create three-dimensional anatomical models. Most features are clearly visible in tomograms, although the smallest teeth (e.g. those on the oral surface of the palate, the vomers and the parasphenoid) appear to be on the limit of scan resolution. After segmentation, 3D models were exported in .ply format and processed in Blender, v. 3.4, for imaging and for digital reconstructions (https://www.blender.org/).

### Synchrotron Tomography

The central region of the skull of NHMD–73588 was also imaged at the European Synchrotron Radiation Facility (ESRF) in Grenoble, France using propagation phase contrast synchrotron X-ray micro-Computed tomography at the BM05 beamline of the ESRF. The beam setup consisted of white beam from the 0.86 T 2-pole wiggler with fixed gap, filtered with 3.93 mm of Molybdenum, resulting in a total integrated detected energy of 133 keV. The acquisition setup consisted of an indirect detector, positioned 3.5 m downstream of the sample for propagation phase contrast, comprising a 2 mm thick LuAG (Lu 3 Al 5 O 12: Ce) scintillator with a visible light reflective layer; a 0.615x magnification from a Hasselblad macro 120 mm f/4 (Victor Hasselblad AB, Gothenburg, Sweden); and a sCMOS PCO.edge 4.2 (PCO, Kelheim, Germany).

Measurements on recorded radiograph returned an isotropic pixel size of 10.14 µm. With this setup, the recorded field of view was 2048 x 304 pixels (horizontal x vertical), i.e., 20.77 x 3.08 mm (limited vertically by the size of the beam). To image the full specimen, 11 tomographic acquisition were performed, moving the specimen by 2 mm along the vertical axis between each acquisition. To compensate for the limited horizontal field of view, the centre of rotation was shifted from the centre of the image by 900 pixels (9.13 mm), resulting in reconstructed tomograms of 3848 x 3848 pixels (39.02^2^ mm^2^). Each acquisition consisted of 5100 projections, each projection resulting from the accumulation of 5 frames of 10 msec exposure each (total integrated exposure of 50 msect), over a rotation of 360°. For each acquisition, 41 flatfield images (image without the specimen) and 40 dark current images (image without beam) were recorded. Tomographic reconstruction was done with PyHST2 (Mirone et al., 2014) using the single distance phase retrieval approach (Paganin et al., 2002) with a δ/β = 1000 and an unsharp mask (sigma = 1.2, coefficient = 4). The resulting 32-bits tomogram were converted to 16-bits using the 0.001% exclusion values from the 3D histogram generated by PyHST2. The post-processing also included ring correction (Lyckegaard et al., 2011; Matlab code for ring correction available at https://github.com/HiPCTProject/Tomo_Recon), normalisation of dense pyrite inclusions (Cau et al., 2017) and 3D cropping of the data.

### Phylogenetic analysis

Our phylogenetic dataset is modified from that of Argyriou et al. (2022), in which *Pteronisculus stensioi* is present. We added NHMD–73588 and five additional species from the literature: *P. gunnari*, *P. magnus* (Nielsen, 1942), *P. changae* (Ren & Xu, 2021), *P. nielseni* (Xu et al., 2014) and *P. cicatrosus* (Lehman, 1952), for a total of 123 taxa. NHMD–73588 was coded as a separate terminal from *P. gunnari* in order to investigate the impact of additional character information derived from XCT analysis.

We also added three characters from the character matrix of Ren & Xu (2021): lacrimal contributing to oral margin (absent/present); teeth on lacrimal (absent/present); position of dermosphenotic (dermosphenotic extending well below dermopterotic/dermosphenotic located at same horizontal level of dermopterotic). This brought the total to 303 characters. Several character states for *Pteronisculus stensioi* were modified from those in Argyriou et al. (2022) to reflect the new information: char. 37: 1 to 0; char. 38: - to 0; char. 39: 0 to 1; char. 40: 1 to 0; char. 41: 0 to 1; char. 66: 0 to 1; char. 68: 0 to 1; char. 111: 1 to 0; char. 127: 1 to 0; char. 166: 0 to ?; char. 224:to 0; char. 225:to 1; char. 231: 0 to 1; char. 249: 0 to 1; char. 281: 1 to 0.

The equally weighted and implied weighted parsimony phylogenetic analyses were conducted in TNT (v. 1.6) (Goloboff et al., 2008; Goloboff & Morales, 2023). The outgroup was constrained as follows for both analyses: (*Dicksonosteus arcticus* (*Entelognathus primordialis* ((*Acanthodes bronni* (*Cladodoides wildungensis Ozarcus mapesae*))(“*Ligulalepis*” (*Dialipina salguerioensis*))). For the equally weighted analysis, the Consistency Index and Retention Index were calculated using the apposite TNT script (stats.run) on a random subset of 100 most parsimonious trees; Bremer support values were calculated using the built-in function of the software, performing a Tbr analysis on the 10752 most parsimonious trees, retaining trees that were suboptimal by up to 10 steps.

For the implied weights analysis, after some preliminary trials we conducted the analysis with a very gentle concavity (K = 100) (Goloboff et al., 2018), recovering 135 best-fit trees and calculating a strict consensus tree.

### Systematic palaeontology

Superclass **Osteichthyes** Huxley, 1880 Class **Actinopterygii** sensu Goodrich, 1930 Genus ***Pteronisculus*** White, 1933 Species *Pteronisculus gunnari* Nielsen, 1942

### Revised diagnosis

Small actinopterygian characterized by the following combination of characters: toothed lacrimal contributing to the orbit and the oral margin; separate intercalar ossification of the braincase; parasphenoid pierced by paired foramina for the internal carotids; tooth cover on anterior corpus of parasphenoid reduced to a median strip in anterior third; at least three pairs of parotic toothplates; absence of dermal basipterygoid processes; canals or cavities in premaxillae, maxillae, dermohyal and dermosphenotic that are not connected to the main lateral line network; anterior and posterior accessory vomers; medially oriented teeth on ectopterygoid and dermopalatines; paired premaxillae; additional prearticular plate on the medial side of the lower jaw; low process on the dorsal margin of the dentary; unperforated propterygium fused to the first lepidotrichium of the pectoral fin.

Differentiated from other species of *Pteronisculus* by: presence of a bony pedicel carrying the buccohypophyseal canal into the parasphenoid; unperforated ventral surface of the posterior myodome; presence of a single ceratohyal; uncinate processes on epibranchials I, II and III; wide entopterygoid process directed dorsolaterally; weak anterodorsal process on the suboperculum; course of the lateral line canal in the supracleithrum almost anteroposterior.

### Description External

The specimen has historically been mechanically prepared, removing the matrix to expose the skull roof, cheek, opercular series and shoulder girdle. As a result, most of the dermal bones in these areas remain as no more than impressions on the surface of the specimen (Fig. 1) and are not visible in tomograms. Parts of the skeleton and the matrix have been substituted by pyrite, particularly the anterior portion of the lower jaws, pectoral fins and scales, and sensory canals; these pyritized areas could not be segmented. The portion of the specimen that was imaged using XCT preserves most of the skull, the jaws, both pectoral girdles and pectoral fins and a portion of the axial skeleton (Fig. 1A-F). It has been subject to a small amount of lateral compression, and some degree of shear in an anterolateral direction. Despite this, most of the bones of the skull and the pectoral girdle are undeformed and preserved in 3D, and range from articulated to loosely associated (Fig. 2A-D). The jaws and external dermal bones in particular are largely preserved in articulation.

**Figure 2:**
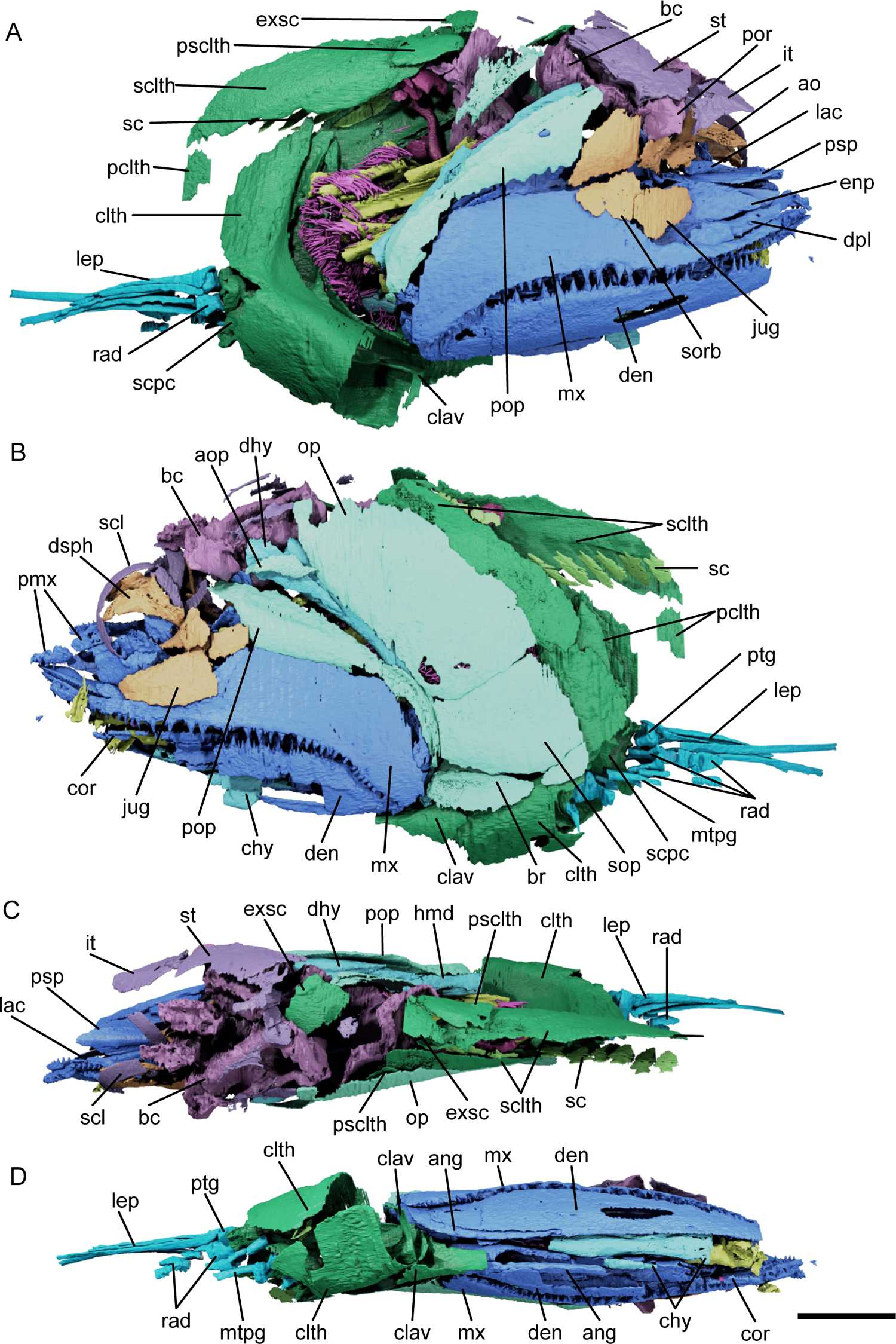
Renders of external and internal anatomy of *Pteronisculus gunnari* NHMD– 75388. Right lateral view (A), left lateral view (B), dorsal view (C) and ventral view (D). Some lepidotrichia and scales omitted. Scale bar: 10 mm. Colour coding of skeleton: orange, circumorbital; dark blue, jaws and palate; dark purple, skull roof and sclerotic ring; light purple, braincase; light blue, cheek and opercular series; turquoise, hyoid arch; yellow, branchial arches; magenta, gill rakers and toothplates; green, shoulder girdle; light green, scales; cyan, pectoral fin. Abbreviations: ang, angular; ao, antorbital; aop, accessory operculum; bc, braincase; br, branchiostegal ray; chy, ceratohyal; clth, cleithrum; clav, clavicle; cor, coronoid; dpl, dermopalatine; dsph, dermosphenotic; dhy, dermohyal; den, dentary; enp, entopterygoid; exsc, extrascapular; hmd, hyomandibula; it, intertemporal; jug, jugal; lac, lacrimal; lep, lepidotrichia; mtpg, metapterygium; mx, maxilla; op, operculum; pclth, postcleithrum; pmx, premaxilla; pop, preoperculum; por, postorbital process; psclth, presupracleithrum; psp, parasphenoid; ptg, propterygium; rad, radials; sc, scale; scl, sclerotic ring; sclth, supracleithrum; scpc, scapulocoracoid; sorb, suborbital; sop, suboperculum; st, supratemporal.

### Snout

The snout comprises the antorbital, premaxilla, rostral and nasal. The rostral and nasal are visible only as impressions on the surface of the specimen, in relative articulation (Fig. 1E - F). The right antorbital and premaxillae have been dislodged from life position.

Only the right antorbital is preserved (Fig. 2A; Fig. 3A-C, ao). It is roughly trapezoidal in lateral view. The dorsal margin of the bone bears an articulation surface for the nasal, and is pierced by a dorsally directed portion of the supraorbital canal. The medial surface is divided into a larger, ventrally directed articular surface and a smaller, dorsally directed one above and below the infraorbital canal. The external surface of the antorbital is ornamented with large tubercules, and is pierced by foramina that connect to the infraorbital canal and the ethmoidal commissure (Fig. 3A, C) The two have almost identical, vertically curved rami of canal (Fig. 3C, ioc, lcm) traverse the bone and converge in a triple junction with the shorter supraorbital canal (Fig. 3C, soc). The ethmoidal commissure exits the antorbital medially, while the infraorbital canal enters through a smaller opening in the posterior margin from the lacrimal.

**Figure 3:**
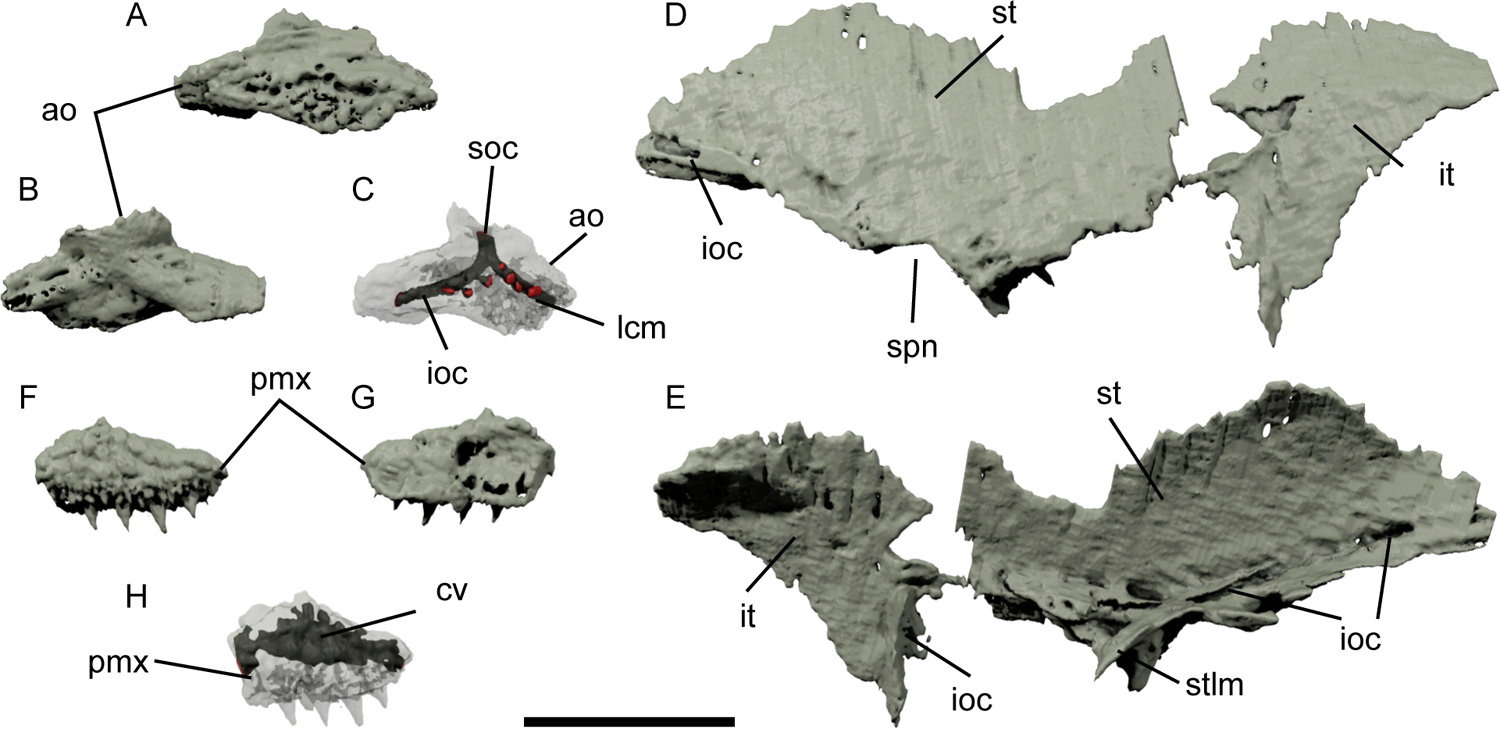
Renders of the snout and skull roof bones of Pteronisculus gunnari NHMD 73588. Right antorbital in anterior view (A), medial view (B) and partially transparent to show the internal ramifications of the infraorbital and supraorbital canals (C). Right supratemporal and intertemporal in lateral view (D) and medial view (E). Left premaxilla in external view (F), medial view (G) and partially transparent to show the infraorbital canal (H). Scale bar: 5 mm. Abbreviations: ao, antorbital; cv, cavity; ioc, infraorbital canal; it, intertemporal; lcm, lateral commissure; pmx, premaxilla; soc, supraorbital canal; spn, spiracular notch; st, supratemporal; stlm, supratemporal lamina.

The premaxilla is small and triangular, with a tuberculate external surface (Fig. 1B; Fig. 3F-H, pmx). It has a straight, tooth-bearing ventral margin, with 4-5 principal teeth and numerous smaller ones; the tooth-bearing margin creates a small horizontal shelf lingually. The bone is deeper anteriorly, where it contacts its antimere. A cavity runs the length of the bone, entering from the lateral margin and without a clear exit on the medial side (Fig. 3H, cv).

The rostral is visible only as an impression on the surface of the specimen (Fig. 1F, ro). Its curved dorsal margin contacts both frontals, while the lateral margins contact the nasals. Nares are not visible. Similarly, the nasal is only visible as an impression (Fig. 1B, D, F, na). The groove for the supraorbital canal can be traced from the nasal into the frontal (Fig. 1F, soc).

### Skull roof

The skull roof comprises frontals, parietals, intertemporals and supratemporals. They are present mostly as impressions on the dorsal surface of the specimen (Fig. 1A-F), and as such it is only possible to partially see their extent and the sutures between them. Some unidentifiable fragments were segmented below the surface of the specimen (Fig. 2A-C).

Only impressions and small patches of tuberculate ornamentation can be identified of the frontal (Fig. 1B, F, fr). It is also possible to follow the supraorbital canal, which has partially been infilled with pyrite (Fig. 1F, soc). The sutures between the frontals and between the frontals and the parietals are irregular, so much so that the left frontal is somewhat wider than the right. The parietal is preserved in a similar manner to the frontal (Fig. 1A, F, pa). Its suture with the frontals is irregular, while that with the supratemporal is more linear.

Portions of the right intertemporal are partially preserved, but it is mostly reconstructed from impressions on the surface of the specimen. It is triangular in lateral view, and broadens posteriorly to meet the supratemporal (Fig. 1B; Fig. 2A, C; Fig. 3D-E; Fig. 4A, it); the suture between the two bones is incomplete, but they share the same mediolateral convexity. The infraorbital canal is continuous between the two ossifications and exits the intertemporal ventrally to enter the dermosphenotic, although the canal is incomplete (Fig. 3D-E; Fig. 4A, ioc). The anteroventral margin of the intertemporal is curved to accommodate the dermosphenotic. The medial margin is incomplete.

**Figure 4:**
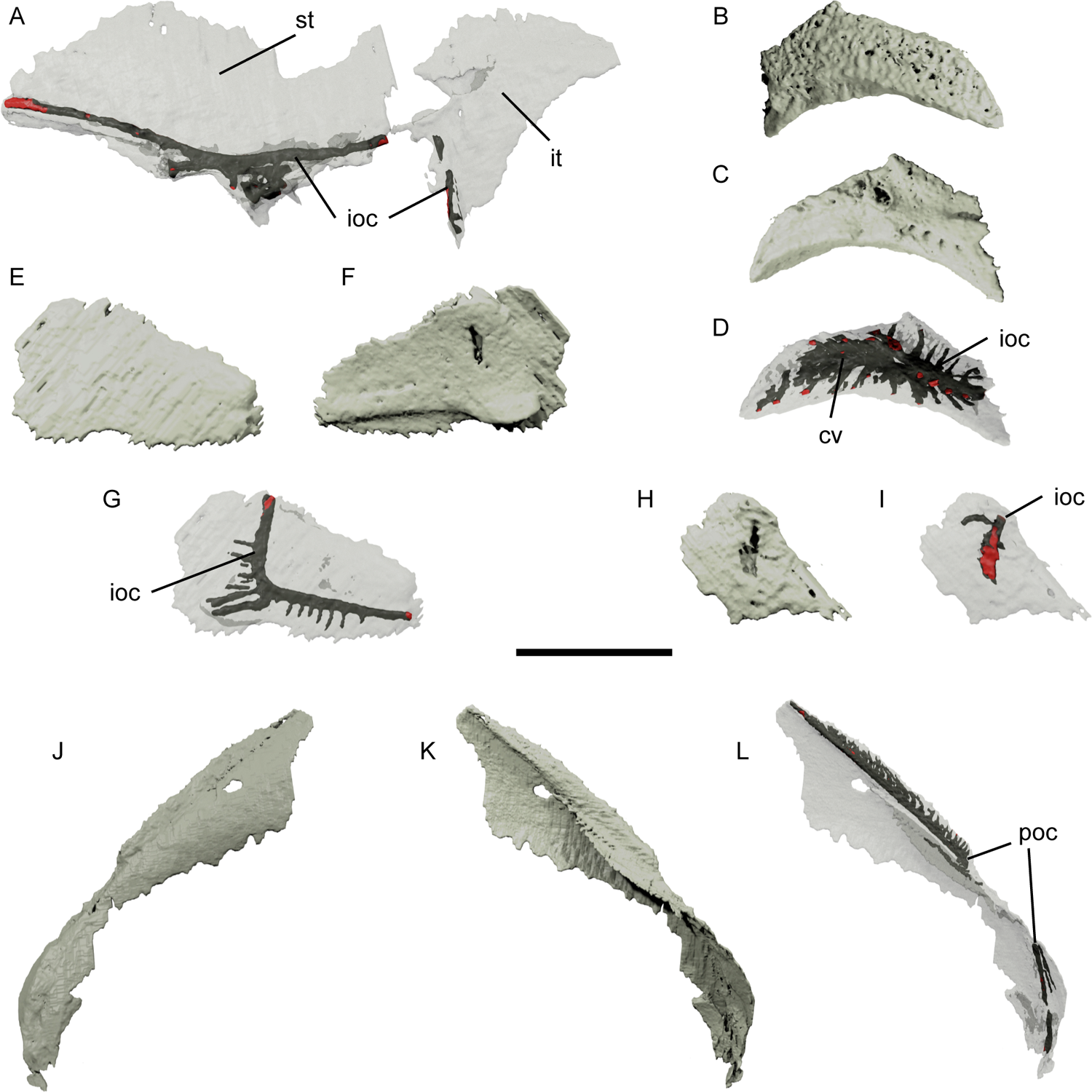
Renders of the skull roof, cheek and infraorbital bones of *Pteronisculus gunnari* NHMD–75388. Right supratemporal and intertemporal in lateral view, partially transparent to show the infraorbital canal (A). Right dermosphenotic in lateral view (B), medial view (C) and partially transparent to show the infraorbital canal and additional cavities (D). Right jugal in lateral view (E), medial view (F) and partially transparent to show the infraorbital canal (G). Left infraorbital in lateral view (H) and partially transparent to show the infraorbital canal (I). Right preoperculum in lateral view (J), medial view (K) and partially transparent to show the preopercular canal (L). Scale bar: 5 mm for A-I; 8 mm for J-L. Abbreviations: cv, cavity; ioc, infraorbital canal; it, intertemporal; poc, preopercular canal; st, supratemporal.

The supratemporal is mostly preserved as impression, although some portions of bone remain, particularly around the sensory canals and close to the braincase (Fig. 1B, D, F; Fig. 2A, C; Fig. 3D, E; Fig. 4A, st). As with other skull roof bones, the dermal ornament is tuberculate. Although the posterior, dorsal and anterior margins are incompletely preserved, the supratemporal is mediolaterally convex. Its lateral margin is produced as two lobes, separated by a distinct spiracular notch (Fig. 3D, spn). The infraorbital canal traverses the entire anteroposterior length, following the lateral margin of the supratemporal. A complex series of smaller canals and cavities depart from and join the infraorbital canal anterior to the supratemporal notch, opening on the ventral and medial surface of the bone (Fig. 3D-E; Fig. 4A, ioc). However, the canal does not appear to exit through the lateral margin of the supratemporal towards the preoperculum. Multiple thin laminae descend from the sensory canal anteroventral and posteroventral to the spiracular notch (Fig. 3E, stlm); these likely correspond to the ventral lamina described by Nielsen (1942, pp. 119-120).

### Cheek and infraorbitals

The cheek and infraorbital series consist of a dermosphenotic, jugal, third infraorbital, two suborbitals, accessory operculum, and preoperculum. Most of these bones are exposed on the surface of the specimen and as a result have been mechanically prepared to a greater or lesser extent (Fig. 1A-D); the suborbitals are completely reconstructed from surface impressions.

The dermosphenotic is roughly crescent-shaped in lateral view (Fig. 1D; 2B; Fig. 4B-D, dsph), with a curved ventral margin contributing to the orbit. It can be divided into smaller, ventrolateral and larger dorsolateral triangular surfaces, both bearing tubercular ornamentation. Its medial surface is pierced by multiple foramina (Fig. 4C). There appear to be two main cavities within the dermosphenotic, both connected to multiple branching rami, with little to no communication between the cavities. The smaller cavity hosts the infraorbital canal, traversing the bone dorsoventrally, entering from the medial surface and exiting from the ventral margin (Fig. 4F, ioc). Smaller vessels ramify from the canal in two rows, dorsally and ventro-medially. The second cavity is complex in shape, with many vessels branching posteriorly, posterodorsally and antero-dorsally, towards both the external and medial surface (Fig. 4F, cv).

The jugal is also partially preserved as an impression, and its margins cannot be fully reconstructed. It is wide and short, almost triangular in lateral view, and carries the infraorbital canal (Fig. 2A-B; Fig. 4E-G, jug). The dorsal margin contributes to the ventral margin of the orbit for the anterior half of its length, while the posterior half articulates dorsally with the third infraorbital. The infraorbital canal enters the bone from its dorsal margin and after a short distance bends anteriorly, running the length of the bone before exiting it into the lacrimal (Fig. 4G, ifc). Smaller branches emanate posteriorly and ventrally, but never towards the orbit. A low ridge on the medial surface traces the anterior arm of the canal, developing posteriorly into a small, ventromedially directed protrusion close to the posterior margin of the bone (Fig. 4E).

An additional, trapezoidal infraorbital, positioned dorsal to the jugal, is partially preserved (Fig. 1B, D; Fig. 4H-I, io). The infraorbital canal travels the bone dorsoventrally (Fig. 4I, ioc).

A total of three suborbitals are preserved as superficial impressions, two on the right side and one on the left (Fig. 1B, D; Fig. 2A, sorb). They are laminar, with convex external and concave internal faces, and are trapezoidal in lateral view. The dorsal suborbital is considerably bigger than the ventral.

As with other cheek bones, the preoperculum has been prepared, and its dorsal limb is partially reconstructed from its impression (Fig. 1B, D; Fig. 2A-C; Fig. 4J-L, pop). The dorsal limb is elongate and roughly triangular in lateral view, with an anterior margin that is recessed to accommodate the suborbitals. The ventral limb of the bone is shorter and crescent shaped, although its anterior margin is probably not fully preserved. The preopercular canal travels the length of the preoperculum, close to its dorsal margin, with its path marked by a ridge on the medial surface of the bone (Fig. 4L, poc). The ridge becomes a ventromedially extending lamina for the posterior half of the dorsal limb (Fig. 4K), and is only faintly present on the ventral half. Within the dorsal triangular portion of the preoperculum, the canal has smaller, anteriorly directed branches that become curved more strongly dorsally further back along the limb (Fig. 4L, poc). The canal in the ventral limb of the preoperculum is directed ventrally and has a few long, ventrolaterally directed branches. The canal exits the ventralmost margin of the bone through a small, hooked process (Fig. 4L).

An accessory operculum is also present on the left side, although it can only be described from surface impressions (Fig. 1D; Fig. 2B, aop). It is small and triangular, tapering to a point anterodorsally and posteroventrally. It bears ridged ornament.

### Hyoid Arch

The hyoid arch comprises the dermohyal, hyomandibula, symplectic and a single ceratohyal. Part of the left ceratohyal and both left and right hypohyals are obscured by pyrite.

The dermohyal is slender and triangular, tapering to a point posteroventrally (Fig. 2B, D; Fig. 5A-C, H, dhy). It is not fused to the hyomandibula, but lies along its dorsal margin, projecting some way dorsal to its anterior head. A raised triangular area on the external face is ornamented with small tubercles; the rest of the dermohyal is smooth, and may have been overlapped by other bones. A distinct lamina is developed along much of the length of the dermohyal on its medial surface, somewhat posterior and ventral to the ridge of ornament. This lamina is developed as a hook-like process at the dorsal margin of the dermohyal and gradually tapers posteriorly. Unusually, the dermohyal carries a canal, which lies underneath the triangular ornamented area (Fig. 5C, cn). Small foramina open onto the dorsal margin between the two ridges.

**Figure 5:**
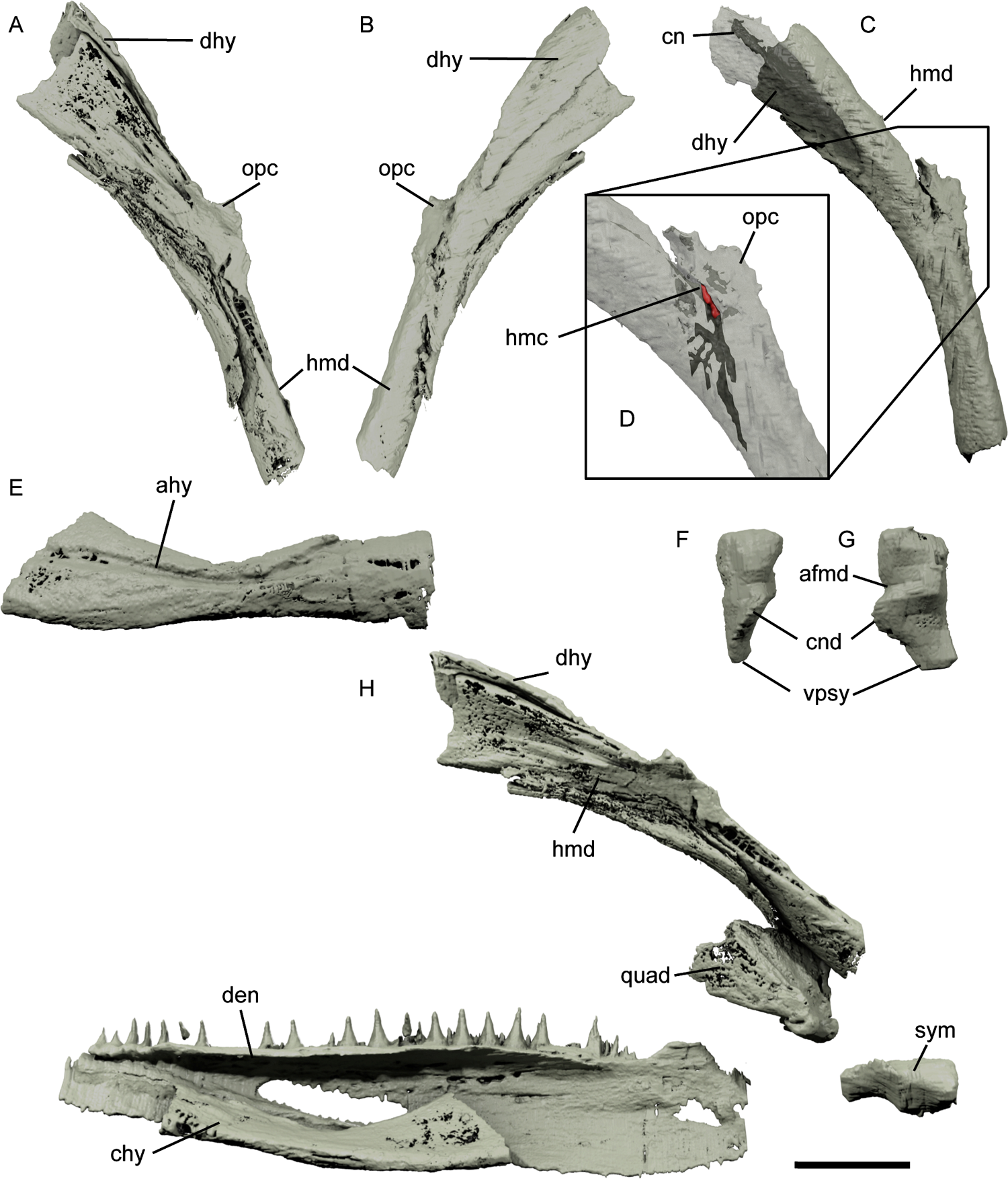
Renders of the hyoid arch and jaw of *Pteronisculus gunnari* NHMD–75388. Right hyomandibula and dermohyal in medial view (A), lateral view (B) and with the dermohyal partially transparent to show the canal (C). Close up of the middle portion of the hyomandibula, rendered partially transparent (D). Right ceratohyal in lateral view (E). Right symplectic in lateral view (F) and medial view (G). Right hyoid arch and selected jaw ossifications in medial view (H). Scale bar: 4 mm for A-C; 8 mm for D; 5.5 mm for E-G; 0.5 cm for H. Abbreviations: afmd, groove for afferent mandibular artery; chy, ceratohyal; ahy, groove for afferent hyoid artery; cn, canal; cnd, anterior condyle of symplectic; den, dentary; dhy, dermohyal; hmc, hyomandibular canal; hmd, hyomandibula; opc, opercular process; quad, quadrate; sym, symplectic; vpsy, ventral process of symplectic.

The hyomandibula is elongate and almost straight, with a dorsal arm that is flattened in cross section and a ventral arm that is oval (Fig. 2C; Fig. 5A-C, H, hmd). A small opercular process is present halfway along the dorsal margin of the bone, marking its maximum curvature point (Fig. 5A-B, D, opc). The process is bilobed: the larger portion lies parallel to the hyomandibula, with a smaller, stumpy portion directed dorsomedially.

The foramen for the hyomandibular nerve pierces the lateral face of the hyomandibula near the base of the process (Fig. 5D, hmc). In addition, two grooves are visible on the surface of the bone, more evident on the right hyomandibula: the first groove runs along the anteroventral margin of the dorsal arm, while the second runs on the lateral face of the ventral arm up to the opercular process (Fig. 5A-B). Each end of the hyomandibula is incompletely ossified. A small, conical symplectic lies anteroventral to the hyomandibula (Fig. 5F-H, sy). As previously described by Argyriou et al. (2022), the symplectic has a rounded dorsal face, tapers ventrally (Fig. 5F-G, vpsy), and bears a condyle and groove for the afferent mandibular artery on its anterolateral face (Fig. 5F-G, afmd, cnd). A single ceratohyal completes the preserved ossified components of the hyoid arch.

It is laterally flattened and approximately half the length of the lower jaw, with a strongly curved dorsal margin and a gently curved ventral margin (Fig. 2B, D; Fig. 5E, H, chy). Its medial surface is slightly concave and external surface slightly convex, creating a narrow ellipsoidal cross section. A deep, sinuous groove for the afferent hyoid artery marks the lateral face (Fig. 5E, chyg).

### Upper jaw and palate

The upper jaw and palate comprise external dermal bones (lacrimal, maxilla, quadratojugal), dermal palatal bones (dermometapterygoid, entopterygoid, ectopterygoid, anterior and posterior accessory vomer, dermopalatines) and endochondral palatoquadrate ossifications (autopalatine, metapterygoid, quadrate). Most components are well preserved, in relative life positions or closely associated, and essentially complete.

The tooth-bearing external dermal bones comprise the lacrimal and maxilla. The lacrimal is narrow, with a gently curved, posterior portion, and a triangular, tooth-bearing anterior portion (Fig. 2A, C; Fig. 6F-H, lac). The dorsal margin contributes to the ventral margin of the orbit for about ¾ of its length, indicated by a slim ridge. The ventral margin bears a similarly fragile ridge and a medial shelf. Both margins are slightly dorsoventrally curved. The anterior portion of the bone is wedge-shaped and has a ventral, toothed surface bearing 6-7 main teeth and numerous smaller lateral cusps. The infraorbital canal enters the bone from the posterior margin and runs in an antero-posterior and slightly dorsal direction, extending above the teeth, exiting as the dorsal margin of the bone leaves the orbital margin (Fig. 6H, ioc).

**Figure 6:**
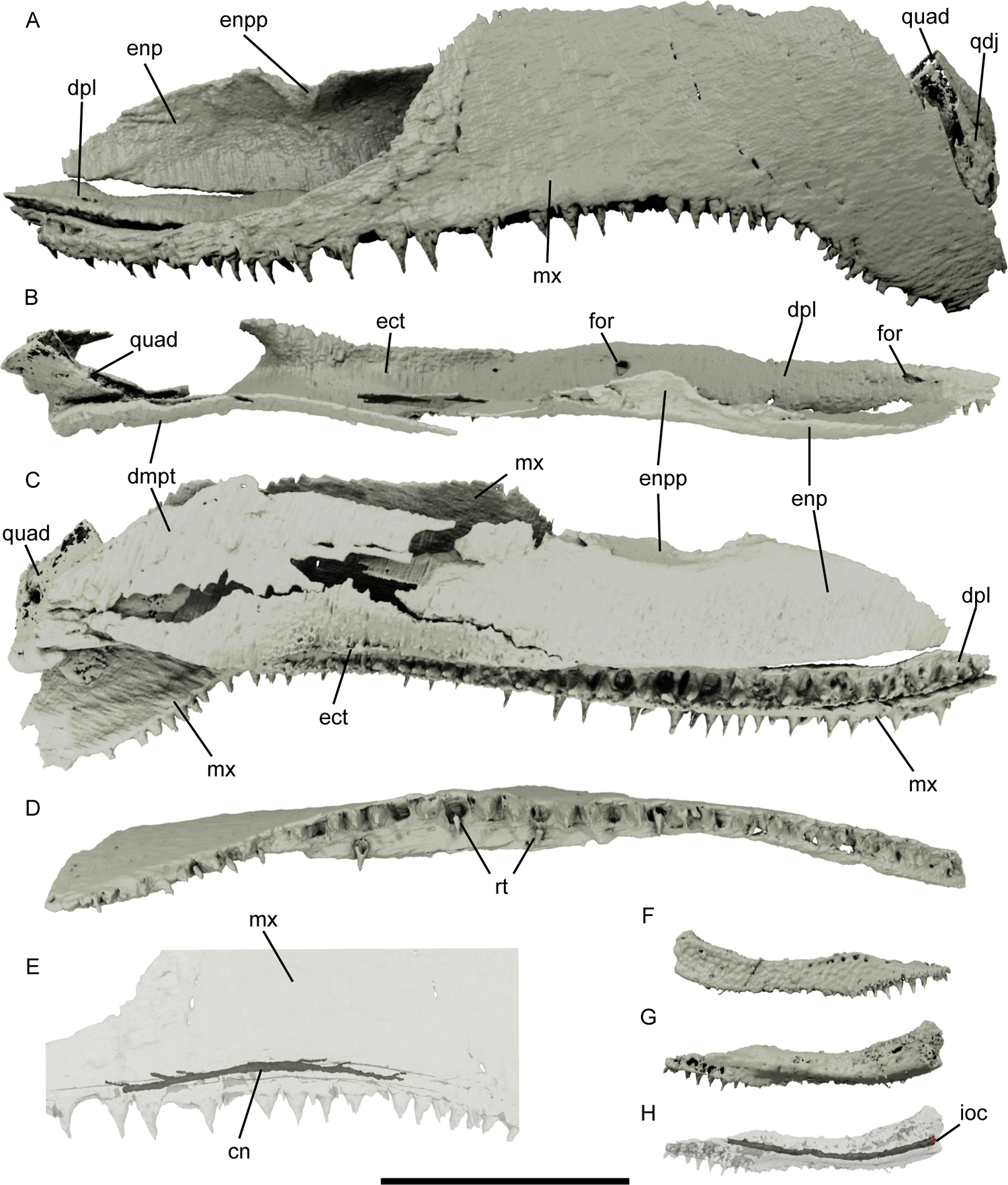
Renders of the upper jaw and palate of *Pteronisculus gunnari* NHMD–75388. Left upper jaw and dermal palate in lateral view (A); dorsal view with maxilla removed (B); and medial view (C). Right maxilla in ventral view (D). Close up of middle portion of maxilla in lateral view rendered partially transparent to show canal (E). Right lacrimal in lateral view (F), medial view (G) and partially transparent to show the infraorbital canal (H). Scale bar: 10 mm. Abbreviations: cn, canal; dpl, dermopalatine; dmpt, dermetapterygoid; ect, ectopterygoid; enp, entopterygoid; enpp, entopterygoid process; for, foramen; ioc, infraorbital canal; mx, maxilla; quad, quadrate; qdj, quadratojugal; rt, replacement tooth.

The maxilla can be completely described, although the anteriormost portion of the suborbital ramus is embedded in pyrite and the majority of the postorbital plate is moldic (Fig. 1B, D; Fig. 2A-B; Fig. 6A, C-E, mx). Overall, it has a slightly convex external surface. The suborbital ramus is narrow, forms about two fifths of the length of the maxilla, and curves slightly dorsally and medially. It consists of a narrow, medial horizontal lamina and a ventral tooth-bearing portion. The lamina extends some way anterior to the tooth-bearing portion of the maxilla, forming an overlap surface upon which the posterior ramus of the lacrimal rests (Fig. 6A, C-D).

The postorbital plate of the maxilla is considerably higher: the anterodorsal margin rises in a curve following the posterior margin of the jugal, the dorsal margin is flat underlying the preoperculum, and the posterior margin falls sharply, descending some way past the quadratojugal (Fig. 6A). As a result, the posteroventral portion of the maxilla covers part of the mandible. A narrow strip along the anterodorsal margin of the maxilla is slightly depressed, forming an overlap surface for suborbital and infraorbitals. The horizontal lamina of the maxilla is thicker and wider below the postorbital plate. A canal runs anteroposteriorly along the maxilla lateral to the thickened portion of the lamina, just dorsal to the tooth row (Fig. 6E, cn); the canal also has small branches, directed posteriorly in the postorbital portion and anteriorly in the suborbital portion. As the maxilla deepens ventrally, the lamina gradually tapers out.

Teeth are borne along the entire ventral margin of the maxilla, which is sinusoidal, with the result that the small, posteriormost teeth point anteroventrally (Fig. 6A, C-D). Two kinds of teeth are present: at least 45 larger, pointed cusps, and numerous irregularly spaced smaller teeth lateral to the main tooth row (Fig 6-7, mx). In addition, around 11 empty sockets are present in each main tooth row, some with what appear to be small replacement teeth partially emplaced (Fig. 6D, rt). Each tooth is capped with acrodin: being a denser material than the rest of the teeth, this appears as a small, bright cone in tomograms. The quadratojugal is very thin and triangular in lateral view, and appears to be incompletely preserved (Fig. 6A, qdj). It lies between the external surface of the quadrate and the medial surface of the preoperculum.

**Figure 7:**
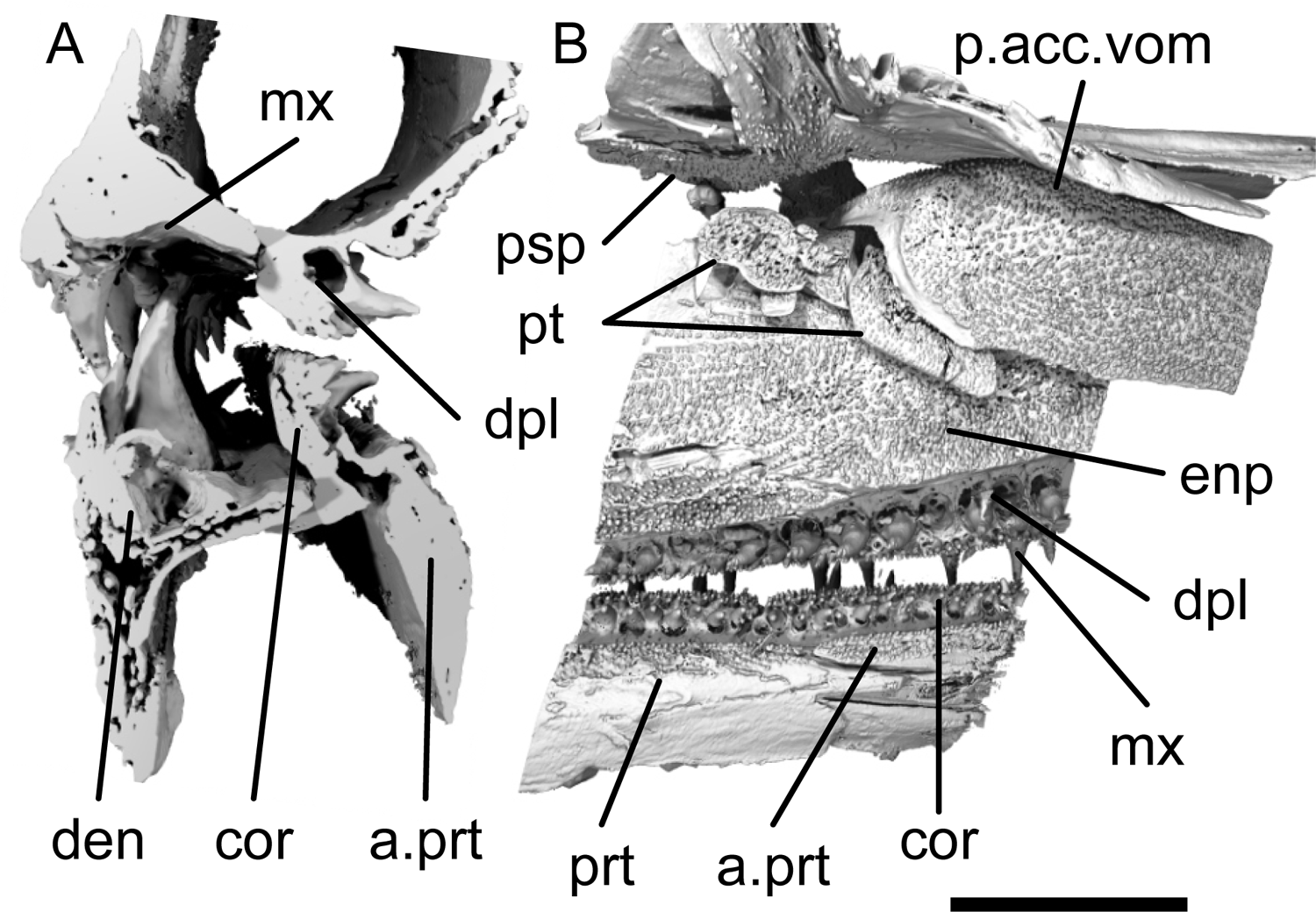
Synchrotron renders of the parasphenoid, left jaws and parotic toothplates of *Pteronisculus gunnari* NHMD–73588. Transverse section through right upper and lower jaws (A) and medial view of the left upper jaw, palate, and lower jaw (B). Scale bar: 5 mm for A; 2.5 mm for B. Abbreviations: a.prt, additional prearticular ossification; cor, coronoid; den, dentary; dpl, dermopalatine; enp, entopterygoid; mx, maxilla; p.acc.vom, posterior accessory vomer; prt, prearticular; psp, parasphenoid; pt, parotic toothplate.

An ectopterygoid, dermopalatine series, entopterygoid and dermometapterygoid sheath the internal surface of the palate. A single, large trapezoidal dermometapterygoid, which thins and fragments anteriorly, forms the posterodorsal margin of the dermal palate (Fig. 6B-C, dmp). It lies medial to both the quadrate and the metapterygoid, with its anteroventral margin contacting the ectopterygoid and the entopterygoid. A ridge on the lateral surface follows the posterodorsal and dorsal margin. The medial surface bears very diminutive teeth (Fig. 7A).

The ectopterygoid has a complex morphology, framing the adductor fossa posteriorly and tapering to a point anteriorly (Fig. 6B-C, ect; Fig. 8A-B, ect). It is continuous with the dermopalatines ventrally and anteriorly (Fig. 6A-C, dpl; Fig. 8 A-B, dpl). The medial face comprises a triangular, vertical lamina, bearing small teeth across almost its entire surface; these teeth are largest towards the ventral margin of the ectopterygoid. The dorsal margin of the lamina contacts the dermometapterygoid in its posterior half and the entopterygoid more anteriorly, before tapering out (Fig. 6C). The ectopterygoid contacts the maxilla via a laterally directed lamina, which is wider posteriorly (Fig. 6B). This lamina and the dorsolateral surface of the medial face are continuous and form a dorsally directed surface that is separated from the tooth row via a thickened ridge and, more ventrally, deep trough (Fig. 7B). Both the ridge and trough are continued by the entopterygoid and dermopalatine series anteriorly. The ectopterygoid tooth row is borne on a separate, ventral lamina. Its teeth are recurved and project medially, at right angles to those of the dentary.

**Figure 8:**
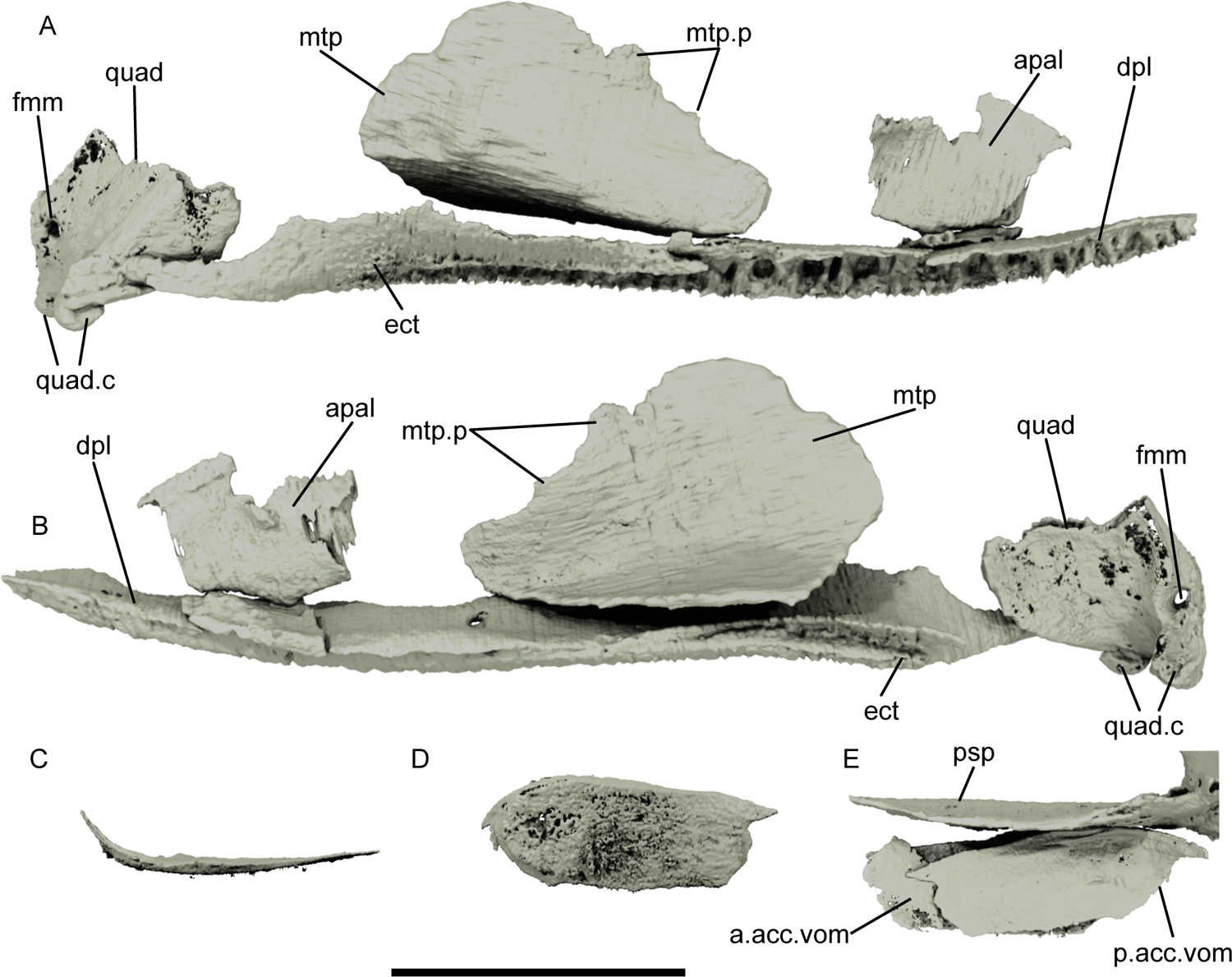
Renders of the palate, parasphenoid and accessory vomers of *Pteronisculus gunnari* NHMD–75388. Left palatal ossifications in medial view (A) and lateral view (B). Left posterior accessory vomer in medial view (C) and ventral view (D). Accessory vomers and parasphenoid in left lateral view (E). Scale bar: 10 mm. Abbreviations: a.acc.vom, anterior accessory vomer; apal, autopalatine; dpal, dermopalatine; ect, ectopterygoid; fmm, foramen for mandibular nerve; mpt, metapterygoid; mpt.p, metapterygoid processes; os, minor palatoquadrate ossification; p.acc.vom, posterior accessory vomer; psp, parasphenoid; quad, quadrate; quad.c, quadrate condyles.

The entopterygoid is elongate and ellipsoidal (Fig. 6A-C, enp). Its medial surface is almost entirely covered in minuscule teeth, except for a small strip parallel to the dorsal margin (Fig. 7A). About halfway along the dorsal margin, just anterior to the metapterygoid, a stout triangular process projects dorsomedially. This may have articulated with the parasphenoid (Fig. 6A-C, enpp). The ventral margin of the entopterygoid is thickened into a ridge, continuous with the ectopterygoid posteriorly and dermopalatines ventrally, from which it is separated by a deep trough. The posterior margin contacts the dermometapterygoid. It is not possible to identify the boundary between the ectopterygoid and dermopalatine tooth, or the number of dermopalatines (der, Fig. 8A-B, dpl), but the teeth extend level to the anterior margin of the maxilla. The tooth-bearing lamina is thickened into a ridge, from which extends a thin lateral lamina, and there is a considerable, gap between the maxillary tooth row and the palatal tooth row (Fig. 7B). Numerous teeth are present on the dermopalatines: a main row dorsally, with a number of empty sockets, and numerous irregularly spaced smaller teeth ventrally. As with the ectopterygoid, the main teeth of the dermopalatine series recurved and project medially––and in some cases almost dorsally—at right angles to the maxillary teeth (Fig. 7A-B, dpl).

Dorsal to the tooth row, the dermopalatines extend as a thin flange towards the entopterygoid. This surface is pierced by two large foramina: a larger, more posterior opening, positioned approximately halfway along the length of the palate, which pierces the bone dorsoventrally and has a minor branch towards the medial side of the bone; and a second, more anterior foramen, which is smaller and opens on the dorsal and ventral margins (Fig. 6B, for). The ventrolateral margin of the tooth-bearing lamina is pierced by numerous small foramina.

A large, posterior accessory vomer is present on each side of the skull. It is rectangular and gently concave in dorsal view, with a medial surface covered in minuscule teeth (Fig. 8E-G, p.vo). Both the anterodorsal and posterodorsal corners are developed as triangular projections; the posterodorsal projection is continuous with a thickened ridge on the lateral face of the posterior accessory vomer, which together taper to a narrow, dorsolaterally-directed point (Fig. 8E-G).

A flat, thin ossification bearing small teeth that lies between the posterior accessory vomers is interpreted as the anterior accessory vomer (Fig. 8E, a.vo). Rather than being present as a single ossification, the palatoquadrate is ossified as a number of separate elements (Fig. 8A-B). The autopalatines are the most anterior, and are made of a triangular portion applied on the dorsal surface of the dermopalatines, and a vertical polygonal portion (Fig. 8A-B, apal). In the right autopalatine the two portions are continuous, while in the left autopalatine are discontinuous, probably as a result of post-mortem damage (Fig. 8A-B, apal).

The metapterygoid is the largest palatoquadrate ossification. It lies midway along the upper jaw and is strongly curved, with almost vertical and ventrolateral laminae (Fig. 8A-B, mtp). The anterodorsal margin of the vertical face bears two low, vertical processes, more evident on the left metapterygoid (Fig. 8A-B, mtp.n).

The quadrate is robustly ossified (Fig. 6A-C; 8A-B, quad). The posterior and medial surfaces are triangular and are divided by a low, anterodorsally directed ridge that originates from the internal condyle. The anterior face is concave, forming the medial and posterior margins of the adductor fossa, and the lateral surface is small and crescent-shaped; its ventral portion contacts the quadratojugal. The condyles are borne on the ventral surface near the posterior margin (Fig. 8A-B, quad.c). The lateral condyle is larger and somewhat medially-directed; the medial condyle is smaller and more rounded. The posterior margin of the quadrate forms a curved ridge, pierced by a large foramen for the internal mandibular branch of the facial nerve (Fig. 8A-B, fmm).

### Lower jaw

The lower jaw is composed of external dermal bones (dentary, angular, surangular), endochondral ossifications (Meckel’s cartilage) and internal dermal bones (prearticular, coronoids, accessory toothplates). The tooth-bearing portions of the right prearticular and coronoids are slightly flattened against the dentary and the prearticular and quadrate are displaced laterally, distorting the outline of the adductor fossa, but in general the mandible is largely articulated (Fig. 9C-D). The left mandible is preserved in life position, although the anterior half of the dentary is pyritized and cannot be segmented (Fig. 9A-B). Both symphyses are concealed within pyrite and thus the anteriormost morphology of the dentary and coronoids is unclear.

**Figure 9:**
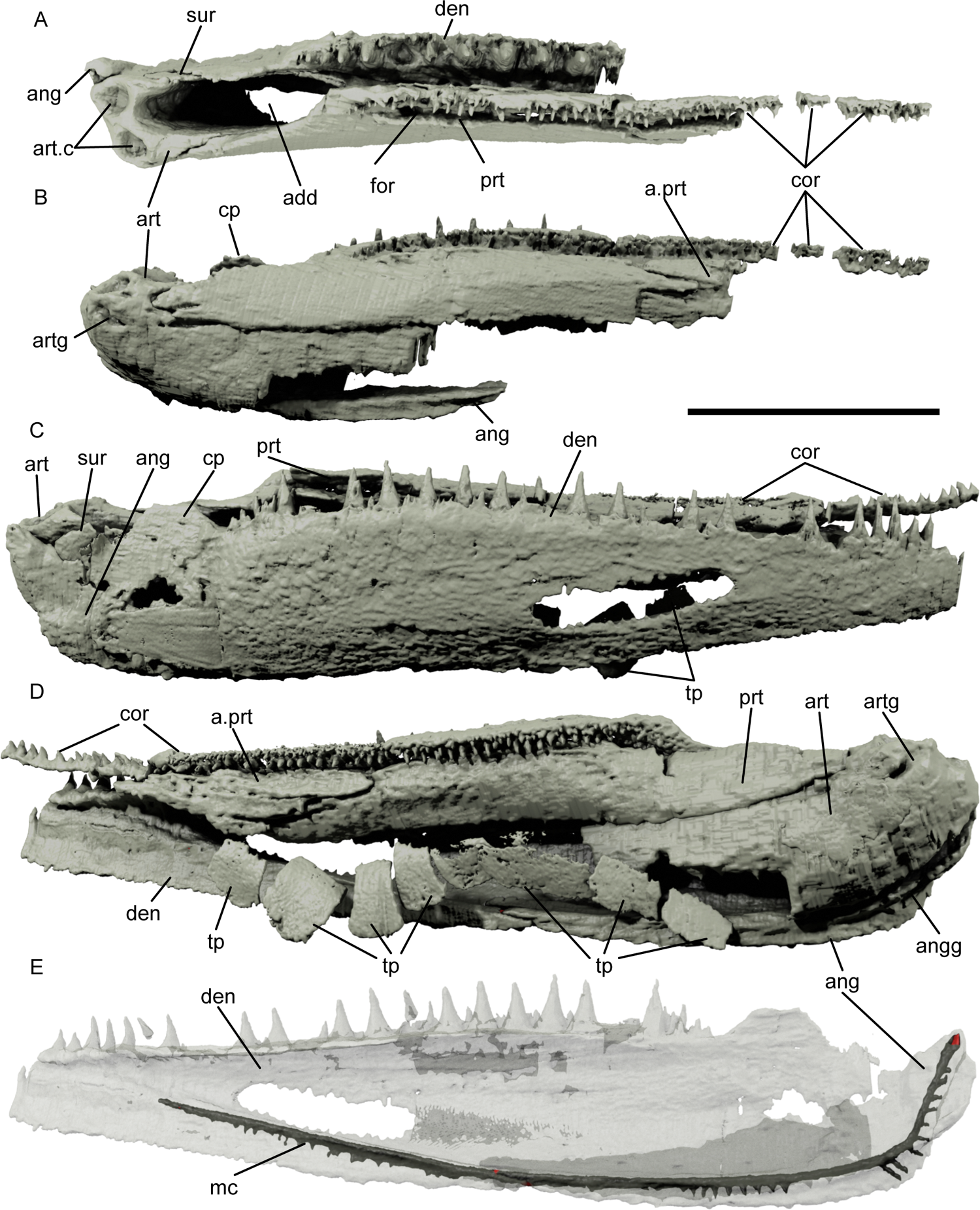
Renders of the lower jaw of Pteronisculus gunnari NHMD 73588. Left lower jaw in dorsal view (A) and medial view (B). Right lower jaw in lateral view (C) and medial view (D). Right dentary and angular in medial view, partially transparent to show the mandibular canal. Scale bar: 10 mm. Abbreviations: a.prt, additional prearticular ossification; add, adductor fossa; ang, angular; angg, groove on the angular; art, articular; art.c, articular cotyles; artg, groove on the articular; cor, coronoid; cp, coronoid process; den, dentary; for, foramen; mc, mandibular canal; prt, prearticular; sur, surangular; tp, toothplate.

The dentary is stout, tapering only slightly anteriorly (Fig. 1B, D; 2A-B, D; 5H; 9A-E, den). Its lateral surface is convex and ornamented with a combination of short ridges ventrally and tubercules dorsally. The dorsal margin bears small lateral teeth, although as they are exposed on the surface and have been partially prepared, many are lost. A main row of larger, conical teeth is present medially (Fig. 9). The right dentary preserves 23 cusps in this tooth row, with the tips covered by a cap of acrodin, and 5 empty sockets.

The tooth row begins anterior to a rounded concavity on the external surface, which in life was overlapped by the ventralmost portion of the maxilla. Behind this depression, the dentary is the sole contributor to a low and thin “coronoid” process (Fig. 9B-C, cp).

A broad horizontal lamina of the dentary extends medially towards a corresponding lateral lamina of the prearticular and coronoids, such that a wide gap separates the dentary tooth row from that of the inner dentition (Fig. 7B, 9A). The dentary lamina is separated from the rim of the adductor fossa by a flange of the prearticular, but otherwise runs for the whole length of the tooth row.

The ventral margin of the mandible is formed by the angular and dentary. The mandibular canal closely follows this ventral margin while it is hosted by the angular, but as soon as the canal passes into the dentary it angles dorsally, probably to meet the tooth row at the anterior tip of the bone (Fig. 9E, mc).

The angular is roughly triangular, forming the posterolateral margin of the mandible and tapering anteriorly to almost the midpoint of the mandible, reaching roughly the end of the horizontal lamina of the prearticular (Fig. 9A, C-E, ang). It contacts the articular and surangular medially, contributes to the lateral side of the adductor fossa, and is largely covered by the dentary laterally. A dorsoventral groove runs down its posterior margin and part of the ventral margin (Fig. 9D, angg). A ridge runs along the entire length of the angular, hosting the mandibular canal (Fig. 9E, mc). The surangular lies dorsal and partially anterior to the angular, and forms the area of the mandible overlapped by the maxilla (Fig. 9A, C, sur). It contacts the horizontal lamina of the prearticular anteriorly, and contributes to the rim of the adductor fossa.

The articular is robust, with two rounded posterodorsal depressions corresponding to the condyles of the quadrate on its dorsal margin (Fig. 9A-D, art; Fig. 9A, qf). A faint ridge runs down the posterior and ventral margin of the articular, originating from the rim of the lateral condyle. Ventral to the condyles, the medial surface is covered in thin dermal bone. A groove runs along the medial surface of the articular, between it and the dermal sheathing bones (Fig. 9B, D, artg). The anterior surface of the articular frames the posterior margin of the adductor fossa, and a narrow lateral projection contacts the surangular. The Meckel’s cartilage is ossified anterior to the articular for approximately a third of the length of the mandible. It becomes increasingly poorly ossified anteriorly, with the perichondral bone hardly developed near its anterior margin. As pyrite obscures the anterior portion of the lower jaws, it is not possible to say whether a mentomeckelian ossification was present.

The prearticular forms much of the medial surface of the lower jaw and the anteromedial margin of the adductor fossa (Fig. 9A-D, prt). Dorsally, it contributes to the internal tooth row for roughly one quarter of its length; Teeth of a variety of sizes are present but organisation into distinct rows is poor. The more ventral teeth are generally larger, with a number of smaller teeth positioned more dorsally; all are directed medially, at 90 degrees to those of the dentary, and have their roots in a dorsolaterally oriented lamina (Fig. 7B, 9A). A deep groove and ventral ridge separate the tooth row from the more ventral portion of the prearticular. These both fade posteriorly, and an elliptical foramen is present near the posterior termination (Fig. 9A, for). Minuscule teeth are present on the medial surface of the ridge, and this tooth covering extends down for about half of the width of the medial surface of the prearticular (Fig. 7A, prt). Ventrolaterally from the tooth row, the prearticular forms a lateral, horizontal lamina that contacts the dentary and frames the anterolateral rim of the adductor fossa (Fig. 9A, add). Anteriorly, the horizontal lamina of the prearticular tapers, and the groove between the dentary and the coronoids is floored by the lamina from the dentary. The tooth-bearing portion passes anteriorly into the coronoids. Ventral to the first coronoid, the prearticular interdigitates with a separate ossification with a similar, entirely dermal composition (Fig. 7, 9B, D, a.prt). Pyrite obscures the anterior extent of this additional bone. A similar bone was described by Lehman (1952) and Nielsen (1942, pp.163-164, figure 38), but wrongly assuming that it had an endochondral component.

At least 3 coronoids are present, sutured to, but distinct from, the prearticular (Fig. 9A-D, cor). The coronoids of the right jaw have rotated somewhat medially. The dorsal portion of each coronoid is directed dorsomedially and surmounted by a low dorsolateral wall, covered in small, irregularly spaced teeth (Fig. 7B). Pyrite invasion means that details cannot be determined anteriorly.

A number of small laminar bones, sub-rectangular in lateral view, with a concave lateral surfaces and convex medial surfaces, lie medial to the lower jaws (Fig. 9D, tp). The dorsal margin is thicker than the ventral, and each plate bears minuscule teeth. Together, these form an elongate row. Some toothplates may be lost or displaced, as eight are associated with the right mandible and six with the left.

### Braincase and parasphenoid

The braincase is incompletely ossified, lacking the ethmoidal and most of the orbitotemporal regions. It is ossified in six different pieces: median basisphenoid, median basiocciput, left and right intercalars, and left and right otic-orbitotemporal regions. Each is slightly displaced from their relative life positions. The areas between each of these ossifications, as well as the ethmoid region and the remainder of the orbitotemporal region, would have been filled with cartilage in life. Some areas of the braincase, particularly in the orbitotemporal region and the lateral face of the otic region, are incompletely ossified; this will be highlighted in the description.

The basioccipital portion of the braincase is almost completely ossified, and is composed of posterior, posterolateral, anterodorsal and ventral surfaces (Fig. 10A, C-D, F). The posterior surface has three openings: a ventral, ellipsoidal aortic canal; a central, rounded notochordal canal, and a dorsal, triangular foramen magnum (Fig. 10A, D, F, aop, nc, fm). The first two face posteriorly while the third faces posteroventrally. The floor and roof of the notochordal canal are incompletely ossified on the midline posteriorly, thus partially connecting the three openings. The dorsal margin of the occipital region is probably incompletely ossified in the specimen; the margin forms the “v” shaped posterior edge of the posterior dorsal fontanelle in the middle, which is the highest point of the occipital portion, then extends laterally to meet the posterolateral surfaces and the lateral margin (Fig. 10A, C, pdf).

**Figure 10:**
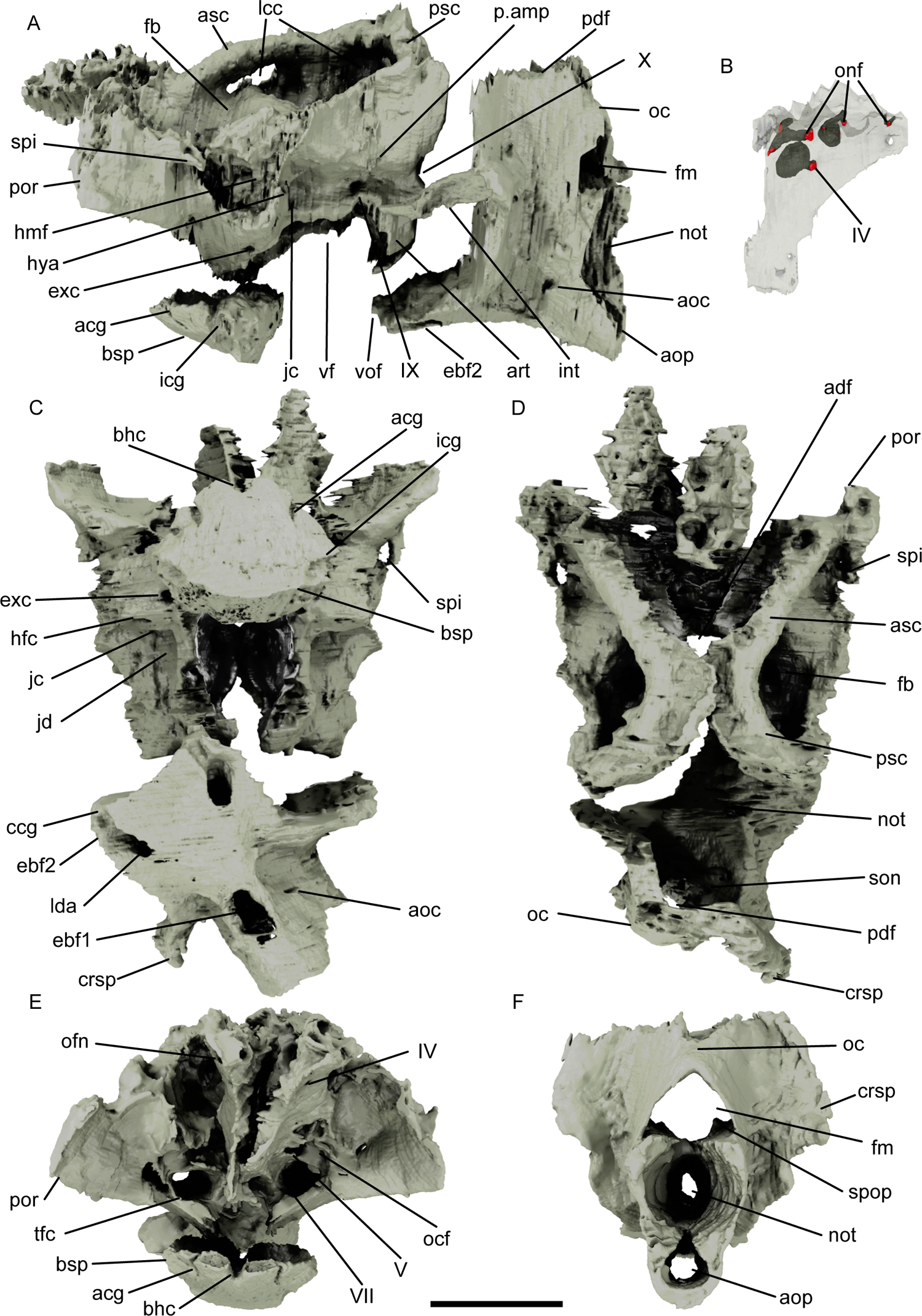
Renders of the braincase of *Pteronisculus gunnari* NHMD–75388. Braincase in left lateral view (A). Right pterosphenoid in lateral view rendered partially transparent (B). Braincase in ventral view (C), dorsal view (D) and anterior view (D). Occiput in posterior view (F). Scale bar: 5 mm. Abbreviations: acg, anterior carotid groove; adf, anterior dorsal fontanelle; aoc, occipital artery canal; aop, posterior opening for the aorta; art, suprapharyngobranchial articulation; asc, anterior semicircular canal; bhc, buccohypophyseal canal; bsp, basisphenoid; ccg, common carotid groove; not, notochordal canal; co, occipital crest; crsp, craniospinal process; ebf1-2, efferent branchial artery foramen; exc, external carotid foramen; fb, fossa bridgei; fm, foramen magnum; hfc, foramen for the hyomandibular trunk of the facial nerve; hmf, hyomandibular facet; hya, foramen for the hyoid artery; icg, internal carotid groove; int, intercalar; jc, jugular canal; jd, jugular depression; lcc, lateral cranial canal; lda, lateral dorsal aorta; my, myodome; oc, occipital crest; ocf, otic canal foramen; onf, ophthalmic nerve foramen; p.amp, parampullary process; pdf, posterior dorsal fontanelle; por, postorbital process; psc, posterior semicircular canal; son, spinoccipital nerve; spi, spiracular canal; spop, spinoccipital process; vf, vestibular fontanelle; vof, ventral otic fissure; IV, trochlear nerve; V, trigeminal nerve; VII, facial nerve; IX, glossopharyngeal nerve; X, vagus nerve.

The aortic canal is enclosed for approximately two thirds of the length of the basiocciput, and its floor is pierced by a median rectangular foramen for the first epibranchial artery (Fig. 10C, ebf1). Dorsal to this, two smaller, paired canals for the occipital arteries pierce the walls of the aortic canal and exit the posterolateral surface of the occipital region (Fig. 10A, C, aoc). The aortic canal bifurcates anterior to the epibranchial artery foramen, and the lateral dorsal aortae diverge from this point and run in an anterolateral direction (Fig. 10C, 11G, lda). These emerge into grooves approximately three quarters of the way along the ventral surface of the basiocciput, bifurcating again into anterior (for the common carotids) and dorsolateral (for the branchial efferent artery) grooves immediately before the ventral otic fissure (Fig. 10C, 11G, ccg, ebf2).

**Figure 11:**
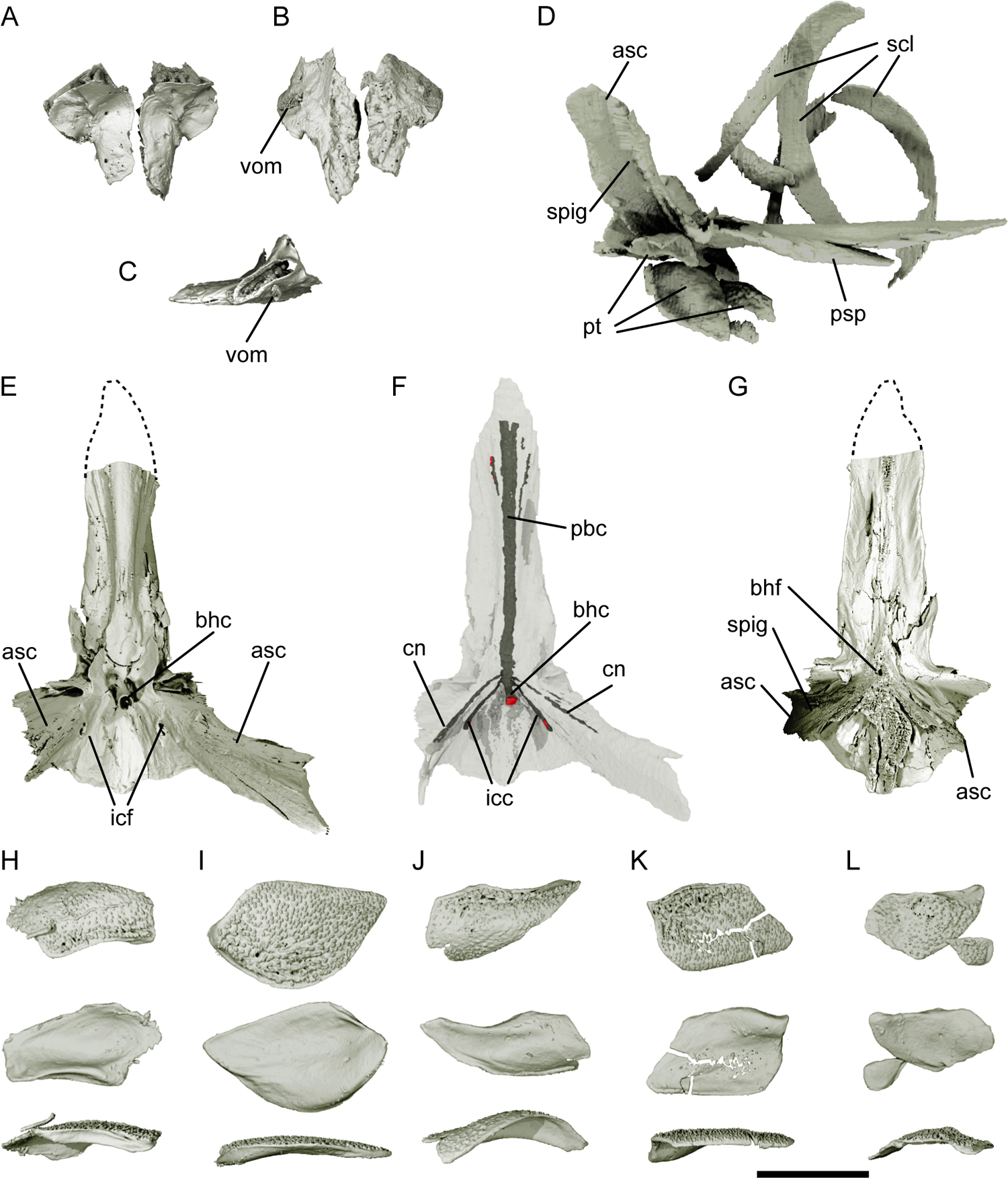
Renders of the ethmoids, parasphenoid, sclerotic rings, and parotic toothplates of *Pteronisculus gunnari* NHMD–75388. Ethmoids in dorsal view (A), ventral view (B) and right ethmoid in lateral view (C). Parasphenoid, sclerotic rings fragments and parotic toothplates in right lateral view (D). Parasphenoid in dorsal view (E), rendered partially transparent (F), and in ventral view (G). Parotic toothplates in ventral view (anterior to right, top row), dorsal view (anterior to left, middle row) and lateral view (anterior to right, bottom row) (H-L). A-C, E, G, H-L rendered from synchrotron data; D, F rendered from microCT data. Dashed lines in E and G indicate the anterior extent of the parasphenoid, as it was not part of the synchrotron data. Scale bar: 5 mm for A-G; 3.5 mm for H-L. Abbreviations: asc, ascending process; bhc, buccohypophyseal canal; bhf, buccohypophyseal foramen; cn, canal; icc, internal carotid canal; icf, internal carotid foramen; pbc, parabasal canal; psp, parasphenoid; pt, parotic toothplate; scl, sclerotic ossicle; spig, spiracular groove; vom, possible vomer.

The notochordal canal is the largest opening on the posterior face of the braincase, but rapidly narrows anteriorly (Fig. 10A, D, F, 11A, not); the walls are not lined with perichondral bone. The notochordal canal opens into the anterior face of the occipital ossification (Fig. 10D, not). The foramen magnum expands almost immediately into the cranial cavity (Fig. 10A, F, 11F, fm). Its short ventral margin is pierced by two pairs of small foramina: the posterior pair host the spinoccipital nerves, while the anterior pair host branches of the occipital artery (Fig. 10D, 11F-G, son, oa). The occipital artery canals have secondary branches. Just posterior to the spinoccipital nerve foramina are two small processes, originating from the foramen magnum floor in dorsomedial direction (Fig. 10F, spop).

The posterolateral surfaces of the occipital region flare laterally around the three medial posterior openings and meet dorsally above the foramen magnum to form a blunt occipital crest (Fig. 10A, D, F, oc). The anterolateral margin of these posterolateral surfaces forms the posterior margin of the otoccipital fissure; the widest point of the occipital region is marked, on the lateral margins, by the posteriorly directed craniospinal process, more evident on the right side in this specimen (Fig. 10C-D, F, crsp). Ventral to the process, a notch in the margin marks the contact with the intercalar, while the likely exit for the vagus (X) nerve is anterior to the process (Fig. 10A, int, X). The otoccipital fissure becomes almost horizontal ventrally, marking the base of the vestibular fontanelle (Fig. 10A, vf). The ventral otic fissure (Fig. 10A, vof) constitutes the anterior margin of the occipital region, forming a concavity that matches a convexity on the posterior margin of the basisphenoid.

A small, elongate ossification corresponding to the intercalar of Pradel et al. (2016) is preserved in articulation with the braincase (Fig. 10A, int). In life, the intercalar would have been in contact with the occipital and otic regions, articulating just ventral to the craniospinal process and the vagus (X) nerve exit and extending partially into the jugular depression. It is pointed anteriorly, with a blunt posterior extremity. its dorsal and ventral margins are sigmoidal in lateral view and the bone has a relatively deep concavity on its lateral surface (Fig. 10A, 16D, int). The bone is pierced by small canals on the ventral and lateral surfaces.

The otic and orbitotemporal regions are co-ossified, although the left and right components are not preserved in continuity across the midline (Fig. 10D). Dorsally, the otic region is completely ossified only around the semicircular canals, and as such it is impossible to establish the extent of the anterior and posterior dorsal fontanelles. Lateral to the semicircular canals, the roof of the otic region is mostly complete, and the lateral faces are also completely mineralized to the level of the vestibular fontanelles, with the exception of the articulation surface for the hyomandibula (Fig. 10A, C-D). Ossification becomes poorer anteriorly, passing into the orbitotemporal region, except for the lateral portions of the postorbital processes (Fig. 10E).

The posterior margins of the otic region form the anterior margin of the otoccipital fissure laterally, and the margin of the posterior dorsal fontanelle dorsally. The vagus (X) nerve passed through a visible indentation in the posterolateral margin, giving it a sinuousoidal outline (Fig. 10A, X). The almost straight dorsal margin marks the lateral border of the fossa bridgei, although here too are signs of incomplete ossification towards the spiracular canal (Fig. 10A, C-D, spi). The fossae bridgei are formed of two deep depressions, connected by a triangular “saddle” above the ampulla of the horizontal semicircular canal (Fig. 10A, D, fb). The posterior depression is rhombic in dorsal view and occupies most of the volume between the posterior and horizontal semicircular canals, reaching its lowest point below the plane of the horizontal semicircular canal; the anterior depression is shallower, positioned lateral to the ampulla of the anterior semicircular canal, and accommodated openings for the spiracular canal and the otic canal (Fig. 10A, C, fb). The lateral cranial canal exits the cranial cavity through a large, rounded canal in the loop on the posterior semicircular canal (Fig. 10A, lcc). It also reconnects with the cranial cavity through the loop of the anterior semicircular canal, lateral to the fossa bridgei (Fig. 10A, lcc), although this feature may be due to incomplete ossification relating to the juvenile nature of the specimen.

The ventral margin face of the otic region is developed into an ovoidal parampullary process (Fig. 10A, p.amp) somewhat posterior to the large vestibular fontanelle. The jugular depression runs just dorsal to the vestibular fontanelle, in a straight line from the posterior margin to the jugular canal (Fig. 10A, C, jd); the glossopharyngeal (IX) nerve exits the cerebral cavity about halfway through this depression via a small foramen (Fig. 10A, IX). The area around the articulation surface for the hyomandibula is incompletely ossified on both sides, extending to include both the spiracular canal and a portion of the postorbital process, and thus appears to cover a much larger area than it would in life (Fig. 10A, hmf). The area around the spiracular canal is incompletely ossified on both sides (Fig. 10A, C-D, spi). It is continuous ventrally with a short spiracular groove that in life would have passed onto the lateral face of the ascending process of the parasphenoid.

Three foramina pierce the postorbital process posterior to the hyomandibula facet: a large, anterior opening for the jugular vein, a ventrolateral opening for the hyomandibular trunk of the facial nerve, and a lateral opening for the hyoid artery (Fig. 10A, C, jc, hfc, hya). The smaller two almost immediately join the main jugular canal, which runs in anterior and slightly ventral direction, to open into the trigeminofacialis chamber on the posterior wall of the orbit (Fig. 10E, tfc). Two additional small canals branch from the jugular canal before it enters the trigeminofacialis chamber: a ventral one for the external carotid, which pierces the lateral face of the braincase below the articulation with the hyomandibula (Fig. 10A,C, exc), and a small lateral one which does not appear to exit the braincase wall. The facial (VII) nerve canal enters the jugular canal just before it opens into the trigeminofacialis chamber, while the trigeminal (V) nerve exit foramen is just above the opening of the jugular canal, in a depression that also hosts the ventral foramen of the otic canal (Fig. 10E, V, VII, ocf).

The postorbital process marks the widest point of the braincase (Fig. 10A, C-E, por); it juts anterolaterally, almost in a plane with the anterior semicircular canal. Its dorsal margin, as well as the portion that connects it to the interorbital wall, are poorly ossified. The dorsal half of the posterior myodome and the, the area surrounding the internal carotids and the optic (II) nerve are completely unossified (Fig. 10E).

The orbitotemporal region is only partially ossified, preserving portions of the posterodorsal interorbital wall and areas surrounding the trochlear (IV) nerve and ophthalmic nerves (Fig. 10A, C-E). The right posterodorsal interorbital wall is separated from the rest of the right otic region. Ossification extends ventrally as far as the oculomotor (III) nerve foramen, although it is too poor for this region to be segmented. The dorsalmost portion of the orbitotemporal region has various foramina preserved: five foramina for the ophthalmic nerves are present in a line, arching very close to the dorsal margin of the preserved interorbital wall (Fig. 10B, E, ofn), while the trochlear (IV) nerve is situated more ventrally (Fig. 10B, E, IV).

A separate basisphenoid cup is preserved ventral to the rest of the braincase, slightly disarticulated but still in association (Fig. 10A, C, E, bsp). It is roughly cup-shaped, with a deeply concave, elliptical dorsal surface, a wide, convex anteroventral surface and an almost vertical, convex posterior surface. The anterior and posterior surfaces meet ventrally at an acute angle, forming a transversal ridge; the posterior surface marks the anterior margin of the ventral otic fissure. Both the anterior and posterodorsal margins are notched on the midline (Fig. 10E). The anterior notch abutted the buccohypophyseal canal pedicle of the parasphenoid (Fig. 10C, E, bhc). A pair of grooves that mark the anterior portion of the basisphenoid housed the anterior carotids, and a more lateral pair housed the internal carotids (Fig. 10A, C, E, acg, icg). Both pairs of grooves are deep dorsally and fade ventrally.

The ethmoids are mineralized separately from the remainder of the braincase, and have been displaced to lie lateral and somewhat dorsal to the parasphenoid. They are roughly trapezoidal, with a straight medial margin and curved anterior margins (Fig. 11A-B). Laterally, the anterior portion is wider than the posterior portion, with these two regions separated by a distinct notch for the autopalatine articulation facet. A large, ovoid articulation facet for articulation with the anterior portion of the palatoquadrate (autopalatine) occupies the entire anterolateral margin. Both the dorsal and ventral surfaces are pierced by small canals, and the ventral surface also bears a curved groove near its medial margin. A small toothplate applied to the anterolateral extremity of the ventral surface, may represent a vomer (Fig. 11B-C, vom).

Multiple fragments from both sclerotic rings are commingled within the orbit (Fig. 11C, scl). Each ring appears to be made up of at least three or four plates, each of which is very thin and has a medial margin slightly thicker than the lateral margin. The plates differ in dimensions, and it is difficult to tell how complete they are.

The parasphenoid is well preserved, extending to the ethmoid region and terminating anterior to the ventral otic fissure. The anterior corpus is long, with a longitudinal medial crest that gives this portion of the bone a triangular cross section. The right margin is slightly damaged in its posterior half (Fig. 11D-G, psp). Small teeth are borne on a narrow medial strip on the anterior third of the ventral surface of the anterior corpus, either side of which are large overlap areas that would have been covered by the accessory vomers (Fig. 11G). The anterior corpus gradually widens posteriorly towards the ascending processes. Dermal basipterygoid processes are absent. The ascending process is broad, bipartite, and in life would have covered most of the posterior face of the sphenotic (Fig. 11D-E, G, asc), with two faces separated by the spiracular groove (Fig. 11D, G, spig): the anterior part of the process is short, rounded, and directed laterally, whereas the more posterior is tall and directed dorsally. Tiny teeth cover the base of the posterior face (Fig. 11G). Roughly half of the left ascending process is fragmented, although it is complete on the right side, reaching the level of the spiracular canal foramen. A deep spiracular groove incises the ascending processes, narrowing to a ventral point and almost meeting its antimere on the midline either side of the small buccohypophyseal foramen (Fig. 11G, bhf, spig).

The buccohypophyseal canal enters the dorsal margin of the parasphenoid via a substantially raised pedicel between the ascending processes (Fig. 11E-F, bhc), with only a small corresponding foramen on the ventral face of the bone (Fig. 11G, bhf). An enclosed midline parabasal canal runs for almost the full length of the parasphenoid, diverging and exiting close to its anterolateral tip (Fig. 11F, pbc). Two additional small canals are emitted from near the origin of the parabasal canal. The more posterior of these leaves the dorsal margin of the parasphenoid near the base of the ascending process, and likely accommodated the internal carotid (Fig. 11E-F, icf, icc), The more anterior runs through the anteriormost portion of the ascending processes and does not appear to exit the parasphenoid (Fig. 11F, cn).

Posterior to the buccohypophyseal foramen, the dorsal and ventral surfaces of the parasphenoid diverge. The dorsal surface is continuous laterally with the ascending processes and cups the basisphenoid ossification. In contrast, the ventral surface is reduced to a narrow, multipartite pedicel covered in tiny teeth almost beyond the limit of the scan resolution. Deep concave embayments flank this pedicel (Fig. 11D, E). These embayments accommodate parotic toothplates, six of which are preserved in a disarticulated position ventral to the posterior corpus of the parasphenoid (Fig. 7A, 11H-L, pt). They are thin and vary from roughly rectangular to teardrop-shaped, with a deeply concave dorsal surface and a convex ventral surface bearing minuscule teeth. One margin of each plate is thickened into a ridge. In life, they appear to have formed pairs and were likely arrayed in two rows, posterior to the buccohypophyseal foramen.

### Endocast

Limited ossification of the internal walls of the cranial cavity, combined with the presence of multiple separate and displaced braincase ossifications, make it difficult to reconstruct much of the endocast. The lateral faces of the mid- and hindbrain regions of the endocast and the labyrinth are best preserved. Of the forebrain region of the endocast, only the posterolateral limits of the optic lobes can be identified. The walls of the midbrain region are also mostly unossified: the anteriormost features that can be described are the anterior ampullae, the canal for the trigeminal (V) nerve and the cerebellar auricles (Fig. 12C-D).

**Figure 12:**
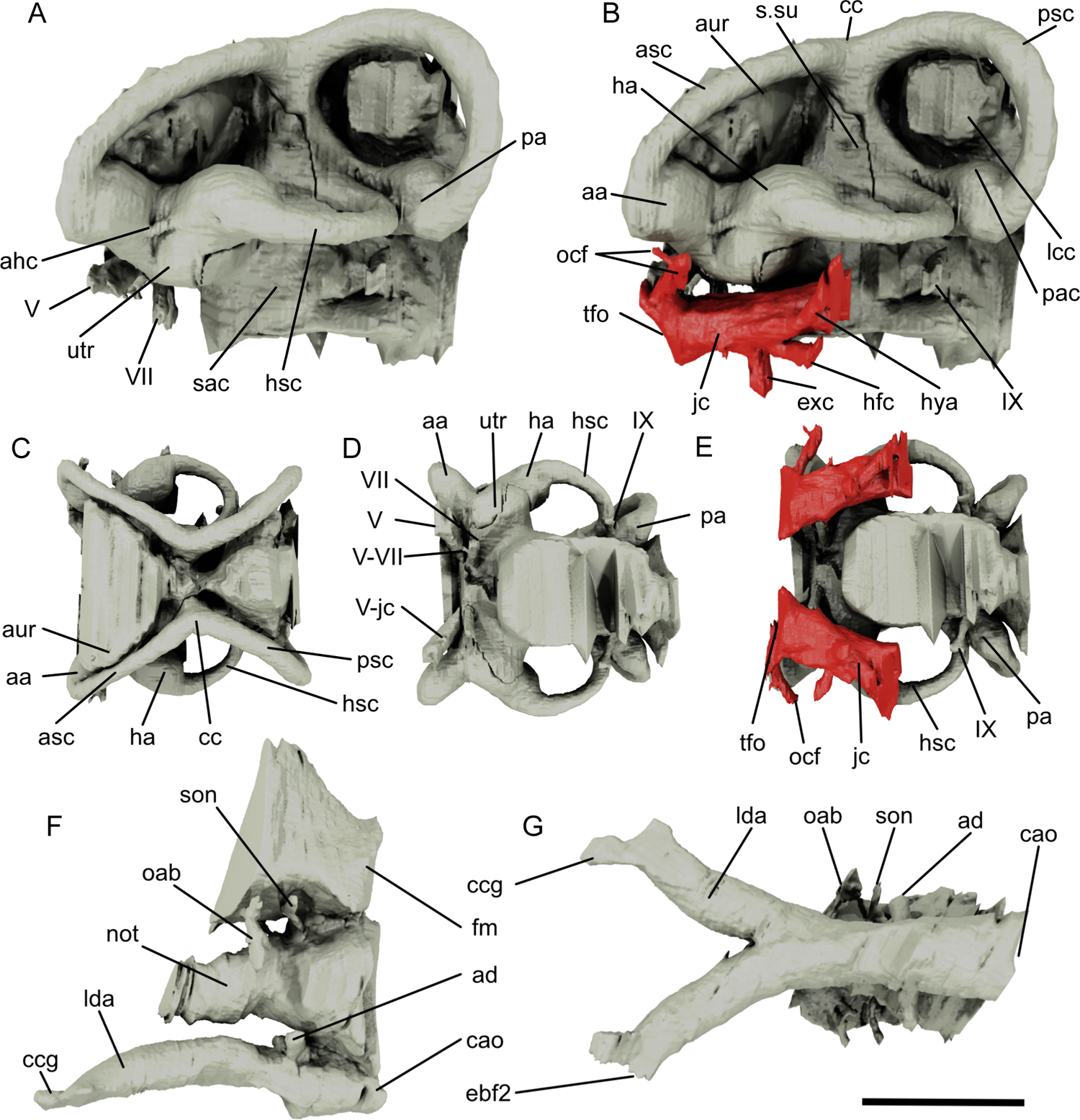
Renders of the endocast of *Pteronisculus gunnari* NHMD–75388. Endocast in left lateral view (A) and with the jugular canal in red (B). Endocast in dorsal view (C), ventral view (D) and with the jugular canal in red (E). Posterior portion of the endocast and basicranial circulation in left lateral view (F) and ventral view (G). Scale bar: 5 mm for A, B; 4 mm for C-E; 6 mm for F, G. Abbreviations: aa, anterior ampulla; ad, artery canal; ahc, connection between anterior and horizontal ampullae; asc, anterior semicircular canal; aur, cerebellar auricle; cao, canal for dorsal aorta; cc, crus commune; ccg, common carotid; ebf2, efferent branchial artery; exc, external carotid foramen; fm, foramen magnum; ha, horizontal ampulla; hfc, hyomandibular trunk of the facial nerve; hsc, horizontal semicircular canal; hya, hyoid artery canal; jc, jugular canal; lcc, lateral cranial canal; lda, lateral dorsal aorta; not, notocord; oab, occipital artery branch; ocf, otic canal; pa, posterior ampulla; pac, preampullary canal; psc, posterior semicircular canal; s.su, sinus superior; sac, saccular recess; son, spinoccipital nerve; tfo, trigeminofacialis opening; utr, utriculus; V, trigeminal nerve; V-jc, trigeminal-jugular canal connection; V-VII, trigeminal-facial nerve connection; VII, facial nerve; IX, glossopharyngeal nerve.

The hindbrain region can be almost completely reconstructed (Fig. 12A-E). It is hourglass-shaped in dorsal view, but the relative displacement of the two halves of the orbitotemporal region must be taken into account when considering the endocast proportions in Figure 12(C-E). Its highest point is medial to the posterior semicircular canal. The cranial cavity, lateral cranial canal and fossa bridgei are continuous; the medial walls of the cranial cavity are incompletely ossified such that the lateral cranial canal has both an anterior and posterior connection through the loops of the corresponding semicircular canals (Fig. 9A; 12A-B, lcc). The cerebellar auricles are just visible as slight bulges medial to the anterior semicircular canals (Fig. 12B-C, aur). The cranial walls surrounding the midbrain and forebrain regions of the endocast are not ossified, and thus these portions cannot be described.

Unlike much of the rest of the endocast, the perichondral bone surrounding the bony labyrinth is almost completely preserved, permitting a full description. The sinus superior is roughly triangular in lateral view (Fig. 12B, ss) and is partially separated from the rest of the endocast by flaps of bone. The posterior semicircular canal is thicker than the others and describes an almost perfect circle, but the corresponding ampulla is the smallest (Fig. 12A-E, psc, pa). A short length of preampullary canal separates the posterior ampulla from the cranial cavity (Fig. 12A-B, pac). The anterior semicircular canal has a very slight curvature, is dorsoventrally compressed (Fig. 12A-E, asc) and enters its ampulla through its anterodorsal margin (Fig. 12A-E, aa).

The posterior and anterior canals meet at the crus commune at a roughly 120° angle in dorsal view, while in lateral view the dorsal margins of the two canals form almost a straight line (Fig. 12A-B, cc). The horizontal semicircular canal is the smallest, but its ampulla is the largest and communicates with the anterior ampulla by means of a narrow canal (Fig. 12A, ahc). Both the posterior end of the horizontal semicircular canal and the preampullary portion of the posterior semicircular canal run parallel for a distance before entering the cranial cavity at the ventral limit of the sinus superior. The utriculus is about twice the size of any of the ampullae and communicates with the ventromedial side of the horizontal ampulla and the posteromedial side of the anterior ampulla (Fig. 12A-B, D, utr). The lateral margin of the saccular chamber is vertical, but its exact shape cannot be determined (Fig. 12A, sac). A short canal carries the glossopharyngeal (IX) nerve laterally into the jugular depression from a point anterior and ventral to the posterior ampulla (Fig. 9A, IX; Fig. 12A-B, D-E, IX). Very narrow canals connect the facial (VII) and the trigeminal (V) nerves and the trigeminal (V) nerve with the jugular canal, while the otic canal has a small branch in anterodorsal direction before exiting into the fossa bridgei posterodorsally (Fig. 12A-E, ocf, V, V-jc, V-VII, VII).

### Operculogular Series

The operculogular series is partially preserved in the specimen and includes the opercular, subopercular and first branchiostegal ray of the left series. As these are mostly preserved on the surface of the specimen, ornamentation is almost completely lost. Other elements of the operculogular series are either prepared away or embedded in pyrite clusters.

The operculum is rectangular, with the shorter side about half the length of the long sides (Fig. 1B, D; Fig. 2B-C, op). The posteroventral margin is straight, while the dorsal and anteroventral margins are slightly curved, and anterodorsally the bone tapers slightly. Much of the suboperculum can only be reconstructed from its impression on the surface, although a fragment of the right is preserved (Fig. 1B, D; Fig. 2B, sop). It is roughly trapezoidal, a little under half the height of the operculum, with an anterior margin shorter than the posterior margin. A small, thin anterodorsal process is present, although this, and the dorsal margin of the suboperculum, were overlapped by the operculum.

The sole undisputed branchiostegal is roughly triangular and somewhat wider than the suboperculum (Fig. 1D; Fig. 2B, br). It bears an anteroposteriorly directed low ridge parallel with its dorsal margin and traces of tubercular ornamentation on its lateral surface. Another thin and long ossification, tapering posteriorly to an asymmetrical tip and with a rounded anterior margin, could be another branchiostegal displaced between the cleithra.

### Gill skeleton

The branchial skeleton is preserved almost in its entirety and largely in articulation. It comprises three incompletely ossified basibranchials, four hypobranchials, five ceratobranchials, four epibranchials, three infrapharyngobranchials and one suprapharyngobranchial. Hundreds of gill filaments and toothplates are also preserved all around the branchial skeleton.

Three basibranchials are present. They have limited internal endochondral ossification and their extremities are unfinished, typically appearing as open, concave surfaces (Fig. 13A-B, bbI-III). The dorsal surface of each is slightly convex, smooth and rectangular. The first basibranchial is triangular in cross section, while the other two have a trapezoidal cross section, which becomes triangular ventrally. Basibranchial I has a convex anterior face, which likely articulated with the hypohyals. Basibranchial II is the most completely ossified element. Two pairs of rectangular depressions are present on its ventral side, one pair near the anterior margin and one pair either side of the ventral process, probably for articulation with the second and third pairs of hypobranchials (Fig. 13A, bbII). Basibranchial III, despite being poorly ossified, has a similar ventral process flanked by depressions (Fig. 13A, bbIII), likely for the fourth hypobranchials.

**Figure 13:**
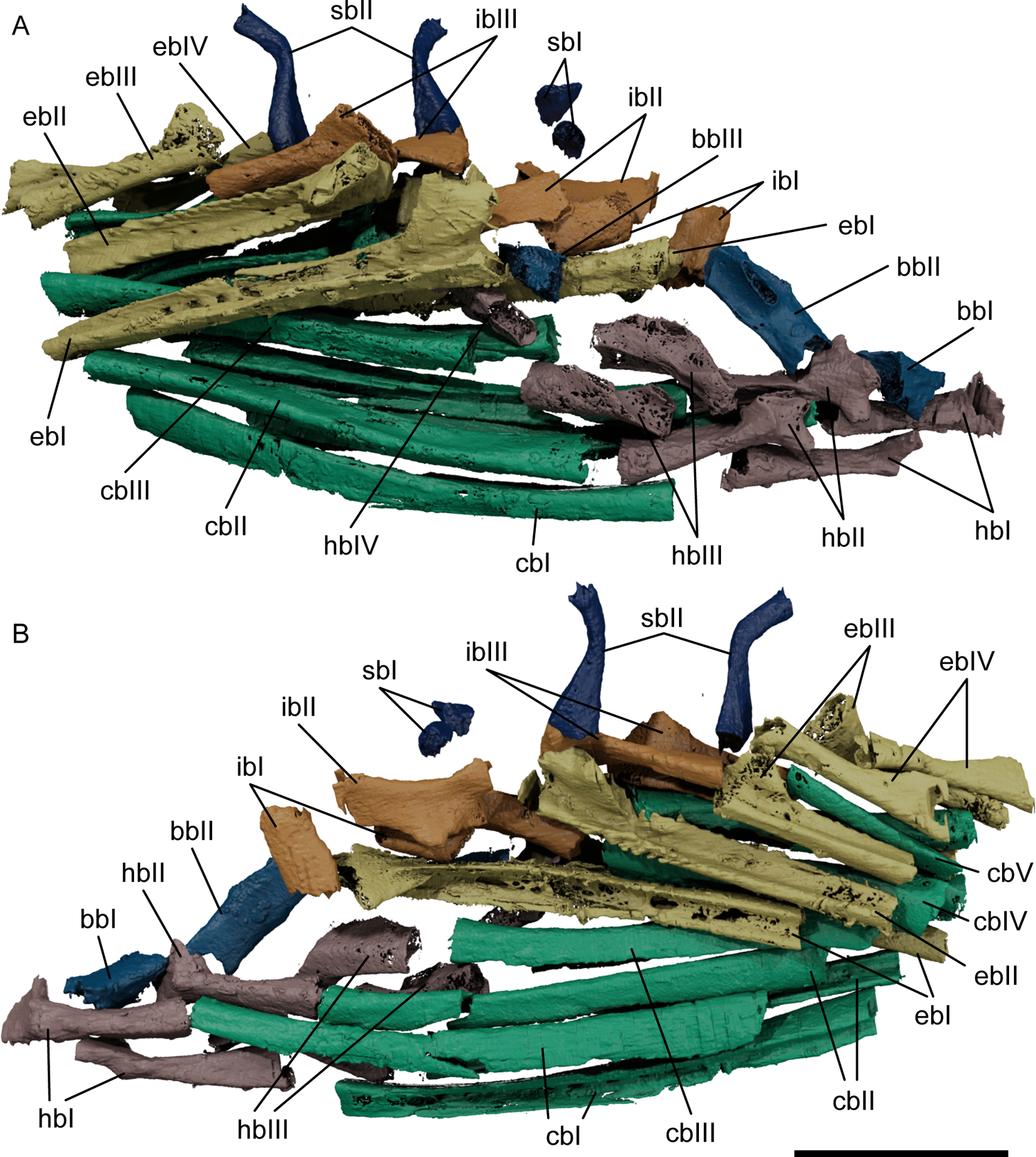
Renders of the gill skeleton of *Pteronisculus gunnari* NHMD–75388. Gill skeleton in right lateral view (A) and left lateral view (B). Scale bar: 10 mm. Colour coding: blue, basibranchials; light brown, hypobranchials; purple, ceratobranchials; green, epibranchials; magenta, infrapharyngobranchials; dark green, suprapharyngobranchials. Abbreviations: bbI-III, basibranchials; cbI-V, ceratobranchials; ebI-IV, epibranchials; hbI-IV, hypobranchials; ibI-III, infrapharyngobranchials; sbI-II, suprapharyngobranchials.

Hypobranchials are preserved in association with the first four gill arches. Hypobranchial I is straight and slender with a rectangular cross section, a triangular, medially directed head, and a slightly expended posterior end (Fig. 13A-B, hbI). Hypobranchial II is similar but slightly shorter, with a bigger, triradiate head (Fig. 13A-B, hbII). The articulation facets for the basibranchials would have been almost vertical in life position. The third hypobranchial is very different, being shorter and twisted. The anterior half of its ventral surface is deeply grooved, while the medial surface bears an anterior hook-like process that articulates with basibranchial II (Fig. 13A-B, hbIII). Hypobranchial IV is much smaller, with a grooved ventral surface giving it an n-shaped cross section, and a slightly flared posterior margin (Fig. 13A-B, hbIV).

Ceratobranchials I – IV are elongate, slightly curved ossifications that decrease in length from the first to the fourth and bear a ventral groove (Fig. 13A-B, cbI-IV). Ceratobranchials I and II have a trapezoidal cross section anteriorly but develop a longitudinal groove on the ventral surface that runs the length of the bone, resulting in an n-shaped cross section posteriorly (Fig. 13A-B, cbI-II). The third ceratobranchial differs in bearing a ventral groove along its entire length, with a corresponding n-shaped cross section, and is slightly sinusoidal in dorsal view (Fig. 13A-B, cbIII). Ceratobranchial IV is also grooved along its entire ventral surface and sinusoidal in dorsal view. Additionally, it has an expanded anterior head with a ventromedially-directed process; the groove runs dorsal to this process (Fig. 13A-B, cbIV). The fifth ceratobranchial is short, thin, and rod-like, lacking a ventral groove (Fig. 13A-B, cbV).

Epibranchials I-III are rod-like, U-shaped in cross section, with a dorsally-directed uncinate process directed on their anterior heads, and a longitudinal groove on their dorsal surface (Fig. 13A-B, ebI-III). Both the total length and the size of the uncinate process decrease from epibranchial I to epibranchial III, but otherwise the bones are very similar. Epibranchial IV is markedly different, being more cylindrical, with anterior and posterior expanses at angles to each other (Fig. 13A-B, ebIV). It lacks both an uncinate process and ventral groove.

A single suprapharyngobranchial I is associated with the first gill arch. It is small and stubby, articulating between the uncinate process of epibranchial I and the parampullary process on the braincase (Fig. 13A-B, sbI). A much longer suprapharyngobranchial II is associated with the second gill arch. It is sinusoid, comprising two arms that meet at an obtuse angle: the anteroventral arm is longer and flares at its base, giving it a conical appearance, while the dorsal arm is about half as long and does not flare at its extremity (Fig. 13A-B, sb II). Infrapharyngobranchial I is short, roughly rectangular in lateral view, and ellipsoidal in cross section (Fig. 13A-B, ibI). The second and third infrapharyngobranchials are larger than infrapharyngobranchial I and dorsoventrally flattened. Infrapharyngobranchial II has a concave dorsal margin, a small lateral process and a flared, bilobate posterior extremity where it contacts the anterior margin of the corresponding epibranchial (Fig. 13A-B, ibII). The third pair is the largest of the three, flares at both extremities and has a longitudinal twist between the anterior and posterior extremities of about 15° (Fig. 13A-B, ibIII).

Hundreds of thin, curved, smooth gill rays are present in the specimen, in loose association with the endoskeletal gill arches but somewhat removed from life position (Fig. 14M). Each ray is very long and needle-like, with a tapered distal extremity and expanded proximal extremity. They are distributed in the areas around the ceratobranchials and the epibranchials, and in clusters medial to the opercula; the length, curvature and number of the rays all increase from the anterior portion of the ceratobranchials towards the epibranchials. Thickness, on the other hand, is maximum around the posterior half of the ceratobranchials. Where they are closer to life position, they are aligned perpendicular to the long axis of the gill ossifications, with the expanded head oriented close to the longitudinal groove of the host ceratobranchials or epibranchials. Small openings arranged longitudinal rows either side of the epibranchial/ceratobranchial groove may represent attachment sites for the gill rays. Medial to each operculum, three to five gill rays are preserved a short line, with greatly expanded bulbous heads (Fig. 14L).

**Figure 14:**
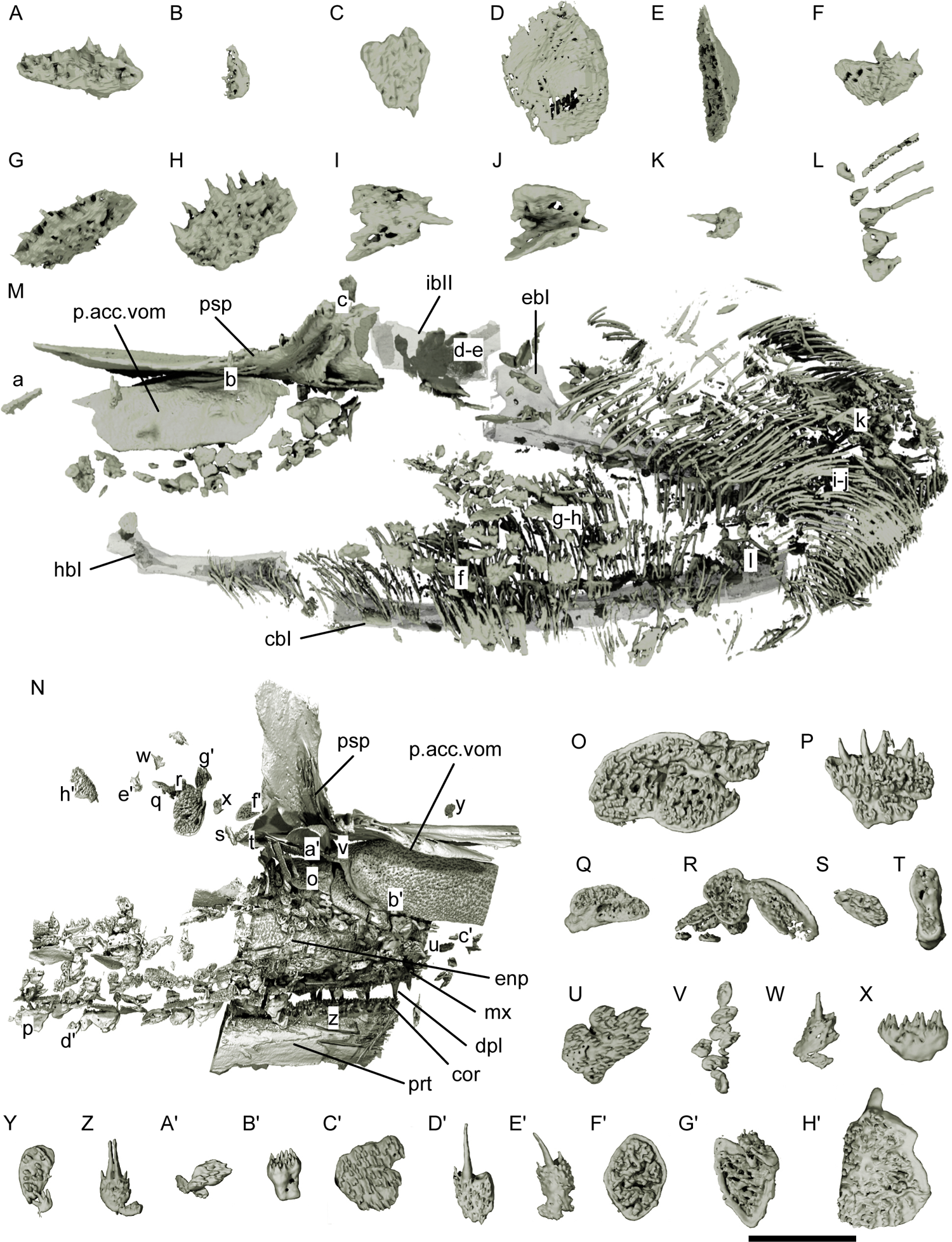
Distribution of toothplates and gill rays in *Pteronisculus gunnari* NHMD-73588. Morphotype 1 toothplate in ventral view (A). Morphotype 2 toothplate in lateral view (B). Morphotype 3 toothplate in ventral view (C). Morphotype 3 toothplate in dorsal (D) and lateral view (E). Morphotype 4 toothplate in lateral view (F). Morphotype 4 toothplate in dorsolateral (G) and lateral view (H). Morphotype 5 toothplate in lateral (I) and medial view (J). Morphotype 6 toothplate in lateral view (K). Gill rays with bulbous heads in lateral view (L). Toothplates and gill rays in life position in left lateral view, with branchial arch rendered partially transparent; position of individual toothplates (A-L) indicated by corresponding lower case letters in left lateral view (M). Toothplates in life position in right medial view (N); position of individual toothplates (O-H’) indicated by corresponding lower case letters. A-M rendered from microCT data; N-H’ rendered from synchrotron data.. Scale bar: 2 mm for A-L, O-H’; 5 mm for M-N. Abbreviations: cbI, ceratobranchial I; cor, coronoid; dpl, dermopalatine; ebI, epibranchial I; enp, entopterygoid; hbI, hypobranchial I; ibII, infrapharyngobranchial II; mx, maxilla; p.acc.vom, posterior accessory vomer; prt, prearticular; psp, parasphenoid.

### Toothplates and gill rakers

Hundreds of small toothplates are concentrated around the branchial skeleton, between the jaws and below the braincase and parasphenoid (Fig. 14M-N). The teeth range in size from barely visible at the XCT scan resolution, to a maximum length of about half a millimetre, which is comparable with the smaller teeth on the jaws. The morphological diversity is wide, but they can be divided into 6 main morphotypes:

1. three to four toothplates with similar morphologies are found in the snout area, close to the premaxillae (Fig. 14A). Thin and relatively long (about 2 mm), they bear many small teeth of different sizes and with no particular arrangement;
2. a couple of very small (1 mm in length) toothplates are closely associated with the ethmoids (Fig. 14B, Y). They have a straight toothed margin with tiny, disorganised teeth, while the opposite margin is deeper on one side and tapers down on the other;
3. flattened plates, either very thin and polygonal (Fig. 14C, Q, AH) or with a flat side covered in minuscule teeth and a slightly convex opposite side (Fig. 14D-E, U, AF-AG). These are found more commonly in the area below the parasphenoid and the braincase, reaching lengths of up to 2-3 mm;
4. plates with a polygonal, flattened base plate and between one and 10-11 conical teeth on the medial margin of the plate (Fig. 14F-H, P, X). These plates sometimes have one or more missing teeth—presumably lost during replacement—as shown in Figure 14G. They are concentrated between the lower jaws and can be found aligned on the lateral surfaces of the ceratobranchials, forming a “wall” between the coronoids and the gill basket. Some of them, positioned anterior to the gill bars, may represent gill rakers. They bear the biggest teeth of all toothplates and are normally 1-2 mm long;
5. plates with a concave-convex base plate, rather than flat (Fig. 14I-J, W). These can bear 1-3 teeth of different sizes and are mostly found posteriorly to the ceratobranchials and around the epibranchials, and thus also likely represent gill rakers. The bigger ones can be about 1 mm wide and 1.5 mm long;
6. plates with a rounded bulb-like root from which 1-4 teeth can sprout are found in the same areas of morphotype 5, about 1-2 mm long including the teeth (Fig. 14K, Z, AD - AE). When multiple teeth are present, one or two are considerably bigger and can have an acrodin cap. These plates appear to represent a third gill raker morphotype.

### Pectoral girdle and fin

The bones of the dermal pectoral girdle comprise the extrascapulars, presupracleithrum, supracleithrum, cleithrum, postcleithrum and clavicle; the endoskeletal girdle is formed from the scapulocoracoid; the pectoral fin endoskeleton comprises proximal and distal radials, lepidotrichia and fringing fulcra. Most of the bones of both pectoral girdles are preserved in the specimen, although the scapulocoracoid shows signs of incomplete ossification and many radials are partially engulfed by pyrite, especially in the right pectoral fin.

A single pair of lateral extrascapulars are preserved in the specimen (Fig. 2A,C; 15A-B exsc). They are laminar, with a convex external side and a concave medial side, roughly triangular in dorsal view. The right extrascapular is partially embedded in pyrite, while the left is more complete and preserves the three-way junction of the supratemporal commissure medially, cephalic lateral line posteriorly, and supraorbital canal anteriorly.

Small fragments situated close to one of the lateral extrascapulars likely represent median extrascapular elements or additional ossifications anterior the extrascapulars as described by Nielsen (1942: fig. 26c; p.116).

Both left and right presupracleithra are preserved, loosely associated with the supracleithra although somewhat displaced (Fig. 2A, C; 15A-B, psclth). Each presupracleithrum is roughly triangular and laminar, with a convex lateral side and concave medial side. The external surface is unornamented, except for a small area near the posteroventral edge of the bone which bears some tubercules, and sensory canals are absent.

The supracleithrum is large, thin, and roughly rectangular (Fig. 1B, D; Fig. 2A-C; Fig. 15A-B, sclth). It is convex, but has a distinct concavity on the anterodorsal portion of the lateral surface. The entire lateral surface is ornamented with ridges. Two canals enter the anterodorsal margin of the supracleithrum. The more dorsal, for the lateral line, is sinusoidal with a slight dorsoventral bend and exits the bone from its medial surface, roughly halfway along the posterodorsal margin (Fig. 15B, llc). A more ventral cavity, comprising two parallel channels, roughly follows the anteroventral margin, before exiting along a narrow ridge on the medial surface (Fig. 15B, cv). A narrow ridge, roughly centred on the medial surface of the bone, parallel to the long margins, may have abutted the dorsal margin of the cleithrum.

**Figure 15:**
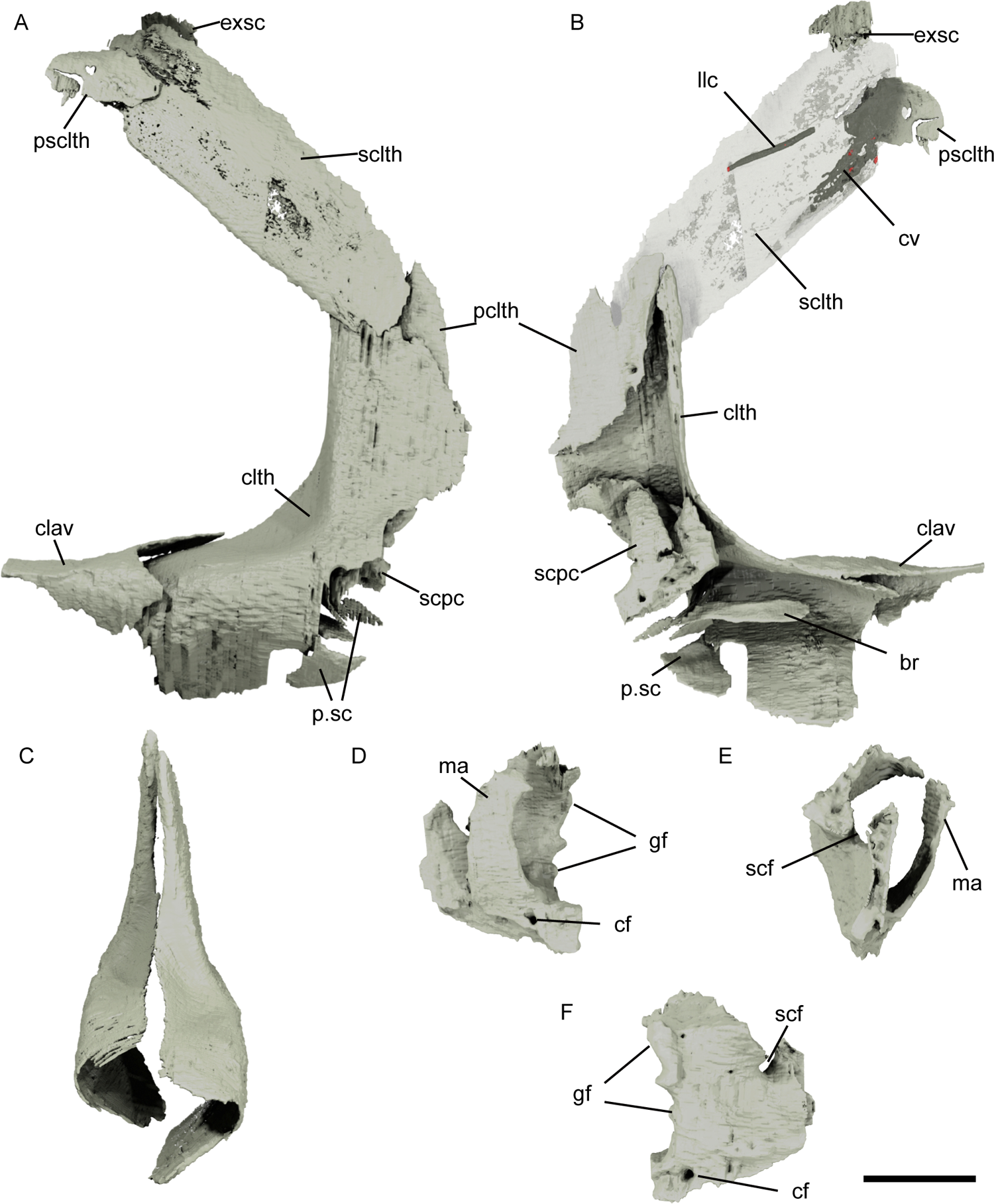
Renders of the shoulder girdle and pectoral fin endoskeleton of *Pteronisculus gunnari* NHMD–75388. Left shoulder girdle and scapulocoracoid in lateral view (A) and medial view, with supracleithrum rendered partially transparent (B). Cleithra in anterior view (C). Right scapulocoracoid in medial view (D), anterior view (E) and lateral view (F). Scale bar: 7 mm for A-C; 5 mm for D-F. Abbreviations: br, branchiostegal ray; cf, coracoid foramen; clth, cleithrum; cv, cavity; clav, clavicle; exsc, extrascapular; gf, glenoid fossa; llc, lateral line canal; ma, mesocoracoid arch; p.sc, pectoral fin scales; pclth, postcleithrum; psclth, presupracleithrum; scf, supracoracoid foramen; sclth, supracleithrum; scpc, scapulocoracoid.

The cleithrum has a tall, pointed dorsal arm with a slightly recurved medial margin, as well as curved mediolateral, anterior and lateral faces that create a continuous surface (Fig. 1B; Fig. 2A-D; Fig. 15A-B, clth). The posterior and posteroventral margins, particularly around the pectoral notch, are poorly preserved; the right is exposed on the surface and the left is obscured by pyrite. The dorsal arm passes into an elongate, thin triangular postbranchial lamina, which is widest at the base of the dorsal arm, close to the pectoral notch. The postbranchial lamina forms an approximately 90° angle with the triangular, curved external lamina, and with the anterior arm through a deep, anteromedially facing concavity. An unornamented strip along the anterior margin of the cleithrum would have been overlapped by the clavicle. The lateral and dorsal surfaces are otherwise ornamented with ridges. A small, thin, tear-shaped postcleithrum sits medial to the posterodorsal margin (Fig. 2A-B; Fig. 15A-B, pclth). It is relatively long, with a bifurcated anterodorsal margin and a posteroventral margin that tapers to a point. Its external surface is ornamented in delicate tubercles. The anterior margin is thickened, with a ridge to accommodate the cleithrum. The clavicle is strongly curved and overlies part of the anterior margin of the cleithrum (Fig. 1A-B, D; 15A-B, clav), but it is unclear how much it overlapped the ventrolateral face. It is partially engulfed by pyrite anteriorly and ventrally, obscuring some aspects of its anatomy. Convex dorsal and ventrolateral triangular faces join at an acute angle that mimics the angle between the same faces of the cleithrum. The more complete left clavicle has a concave posterior margin on the dorsal face, and an elongate posteromedial process that extends along the horizontal face of the cleithrum.

Both scapulocoracoids are preserved in the specimen, although most of the coracoid plate, and parts of the anterior and mesocoracoid processes are incompletely mineralized (Fig. 2A; Fig. 15A-E; Fig. 16A-B, scpc). The unossified areas are identical in both scapulocoracoids.

As the anterior process is unmineralized, the anterior portion of the bone is present only as a small, dorsally directed stump. The supracoracoid foramen is round and opens posterodorsally, but its dorsal margin is also incompletely mineralized (Fig. 15C, E, scf).

The mesocoracoid arch is thin and has a gentle anteroposterior curvature, more evident on the posterior margin; it is oriented such that it does not obscure the supracoracoid foramen in medial view (Fig. 15D-E, ma). Its lateral surface encloses the dorsal muscle canal medially, while dorsally the arch is incomplete, and as a result the canal is wide and roughly triangular in section. The coracoid foramen lies just ventral to the posteroventral margin of the root of the mesocoracoid arch and pierces the bone in posterolateral direction (Fig. 15C-D, cf).

The dorsal and coracoid (ventral) portions of the bone are almost entirely unmineralized, so it is not possible to determine the shape of the ventral muscle canal or how the scapulocoracoid contacted the cleithrum. Similarly, there is no mineralized dorsal coracoid process. In contrast, the glenoid fossa is fully mineralized on the posterior margin of the bone (Fig. 15C-D, gf): in life position, it would have formed a roughly 30-45° angle with the vertical posterior margin of the cleithrum. Its concave margin has 3-4 protrusions for articulation with the endoskeleton radials, directed posterolaterally to ventrolaterally.

All radials appear to be preserved, but most are partially or wholly engulfed in pyrite and thus cannot be fully described. It is possible to identify the propterygium and metapterygium of the right fin, and an additional four to five proximal radials and at least three distal radials (Fig. 16A-E, ptg, mptg, pr.rad1-4, ds.rad). The right propterygium is well preserved and in situ, articulating with the lateral margin of the glenoid fossa (Fig. 16A-B, D-E, ptg). It is complex in shape, with dorsal and ventral portions almost divided in two by a deep grove on the lateral surface. The propterygium is imperforate, and this groove may have transmitted the properygial canal; alternatively, this may have passed medial to the propterygium, between it and the next radial. The propterygium is also fused to the first, fused lepidotrichia (Fig. 16A-B, D-E, ptg, lep). The first radial is bulbous and blocky close to the glenoid fossa and flares out medially (Fig. 16A-B, D-E, pr.rad1); similarly, the second radial is smaller towards its articulation with the scapulocoracoid and flares out towards the distal radials, although is much flatter than the first radial (Fig. 16A-B, pr.rad2). The remaining proximal radials are rod-like with slightly flared proximal heads, but they could not be fully segmented (Fig.16A-C, pr.rad3-4). The metapterygium is roughly twice as long as the first radial and rod-like in section (Fig. 16A-C, mptg). Several distal radials are also present. Three have a flattened, roughly triangular or trapezoidal shape (Fig. 16A-C, ds.rad).

**Figure 16:**
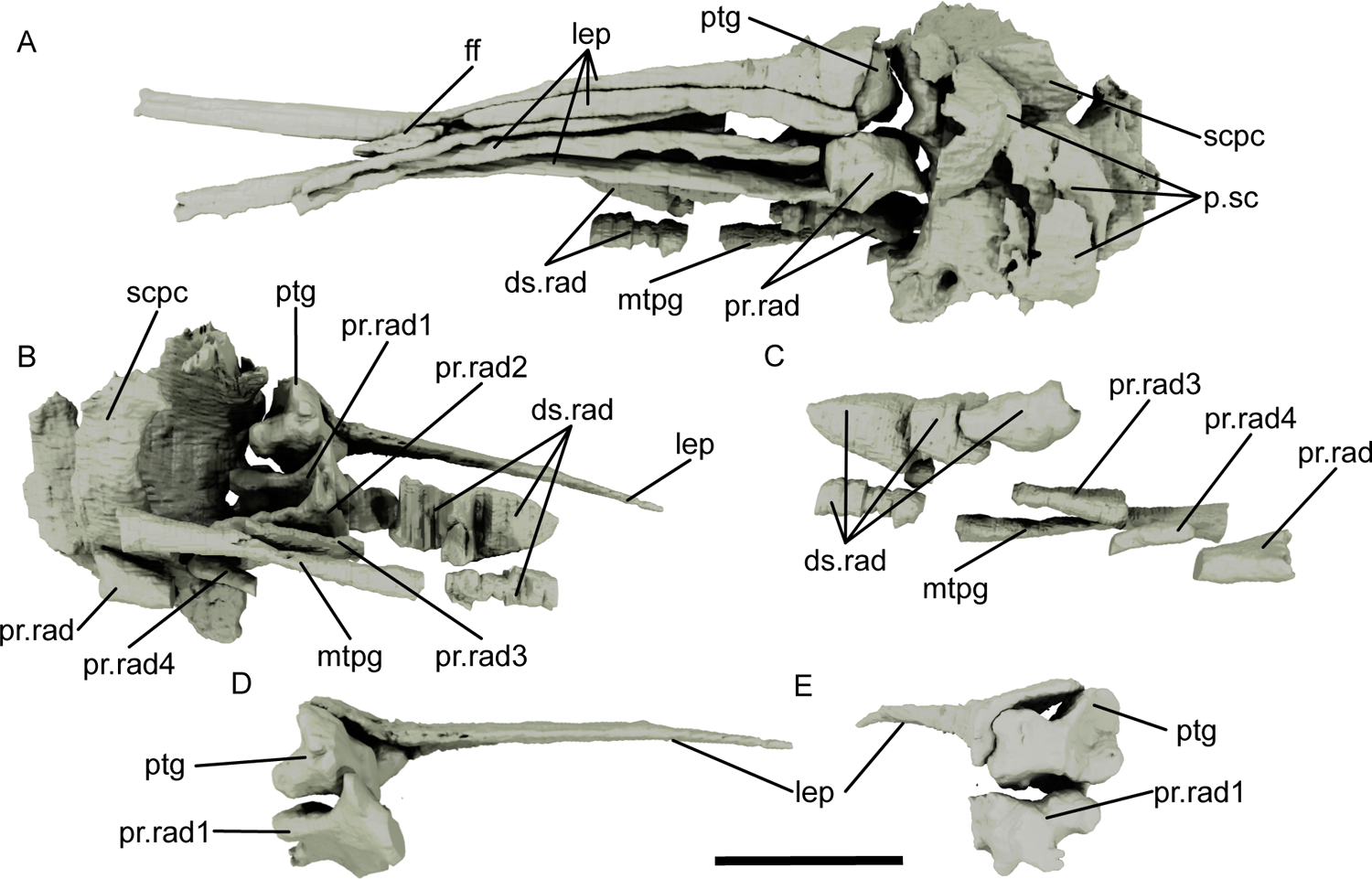
Renders of the pectoral fin of *Pteronisculus gunnari* NHMD–75388. Right pectoral fin in lateral view (A) and medial view (only first lepidotrich figured) (B). Distal radials, last three proximal radials and metapterygium in lateral view (C). Propterygium and first lepidotrich in posteromedial view (D) and anterolateral view (E). Scale bar: 5 mm. Abbreviations: ds.rad, distal radial; ff, fringing fulcra; lep, lepidotrichia; ptg, propterygium; mtpg, metapterygium; pr.rad1-4, proximal radial; p.sc, pectoral fin scale; scpc, scapulocoracoid.

The lepidotrichia are long, slender and slightly recurved; a few representative elements were segmented and are shown in Fig. 1B, 2A-B and D (lep). A minimum of 30 are present for each fin. The first lepidotrichium is the shortest, formed of several conjoined lepidotrichia, and is fused to the propterygium (Fig. 16A-B, D-E, lep, ptg). Each lepidotrichium becomes segmented after a certain length, but the two halves do not diverge but rather remain joined together. Multiple, very thin ossifications of variable dimensions and curvature localised around both scapulocoracoids and the radials are interpreted as fin scales (Fig. 15A-B; Fig. 16A, p.sc). Fringing fulcra are also present (Fig. 16A, ff).

### Axial skeleton

The axial skeleton comprises basidorsals, basiventrals and scales. Elements become increasingly difficult to segment more posteriorly due to artefacts from pyrite growth. At least 14-15 pairs of basidorsals are preserved, but only one complete basiventral, plus an incomplete basiventral associated with the second pairs of basidorsals.

The basidorsals are paired, and become more posteriorly in the vertebral column. The neural arch is oval in cross section and oriented posterodorsally (Fig. 17A-C, na). In smaller basidorsals, the neural arch is detached. Close to its base, a small process juts out anterodorsally (Fig. 17A-B, pr); the same process creates a small ridge on the medial surface of the basidorsal (Fig. 17A-B, r). These features become progressively smaller as the basidorsals become smaller, until they are barely visible. The epineural process is broad and has an ovoidal cross section (Fig. 17A-D, ep). It is arched ventrally in the first pair of basidorsals, then becomes smaller and loses its curvature in smaller basidorsals.

**Figure 17:**
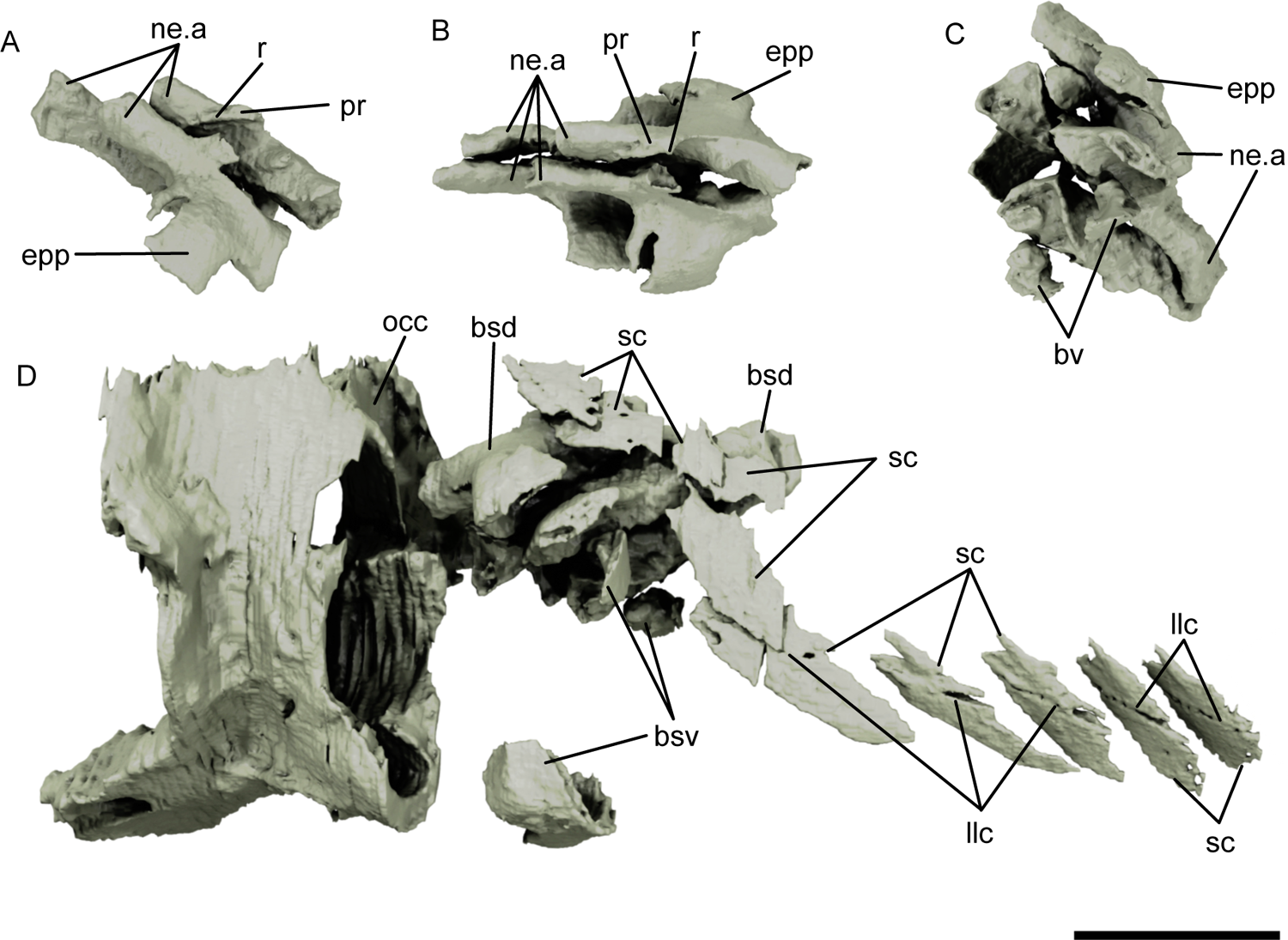
Renders of the axial skeleton of *Pteronisculus gunnari* NHMD–75388. Basidorsals in right lateral view (A), dorsal view (B) and posteroventral view (C). Occipital region of the braincase, portion of the axial skeleton and scales in left posterolateral view (D). Scale bar: 5 mm. Abbreviations: bsd, basidorsal; bsv, basiventral; epp, epineural process; int, intercalar; llc, lateral line canal; ne.a, neural arch; occ, occiput; pr, process; r, ridge; sc, scale.

Basiventrals are only associated with the first two pairs of basidorsals (Fig. 17C-D, bv). The only complete basiventral is “U”-shaped and has width comparable with the lower third of the notochordal canal. The anterior margin is notched. More basiventral fragments can be seen around the first two pairs of basidorsals (Fig. 17D, bv).

Numerous scales are preserved in the specimen, albeit disarticulated and in loose association (Fig. 2A-C, sc). The scale covering is essentially complete, as scales can be found all around the body. Lateral line scales show pores, corresponding to the lateral line canal. All scales are rhombic in shape and present peg-and-socket articulation on the dorsal margin (Fig. 17D, llc). The anterior scales are narrower and more elongated than posterior ones, while the dorsal and ventral scales are considerably smaller and with a stronger curvature.

### Phylogenetic results

Our equally weighted parsimony analysis retrieved 10752 most parsimonious trees of 1693 steps (Consistency Index: 0.198; Retention Index: 0.674); the strict consensus tree is shown in Figure 18. Sarcopterygians are recovered as monophyletic (Bremer Decay Index, BDI: 3), although most included taxa branch from a basal polytomy. Sarcopterygians are resolved in more detail in the implied weights analysis (See Supplementary Material, Figure S1).

**Figure 18:**
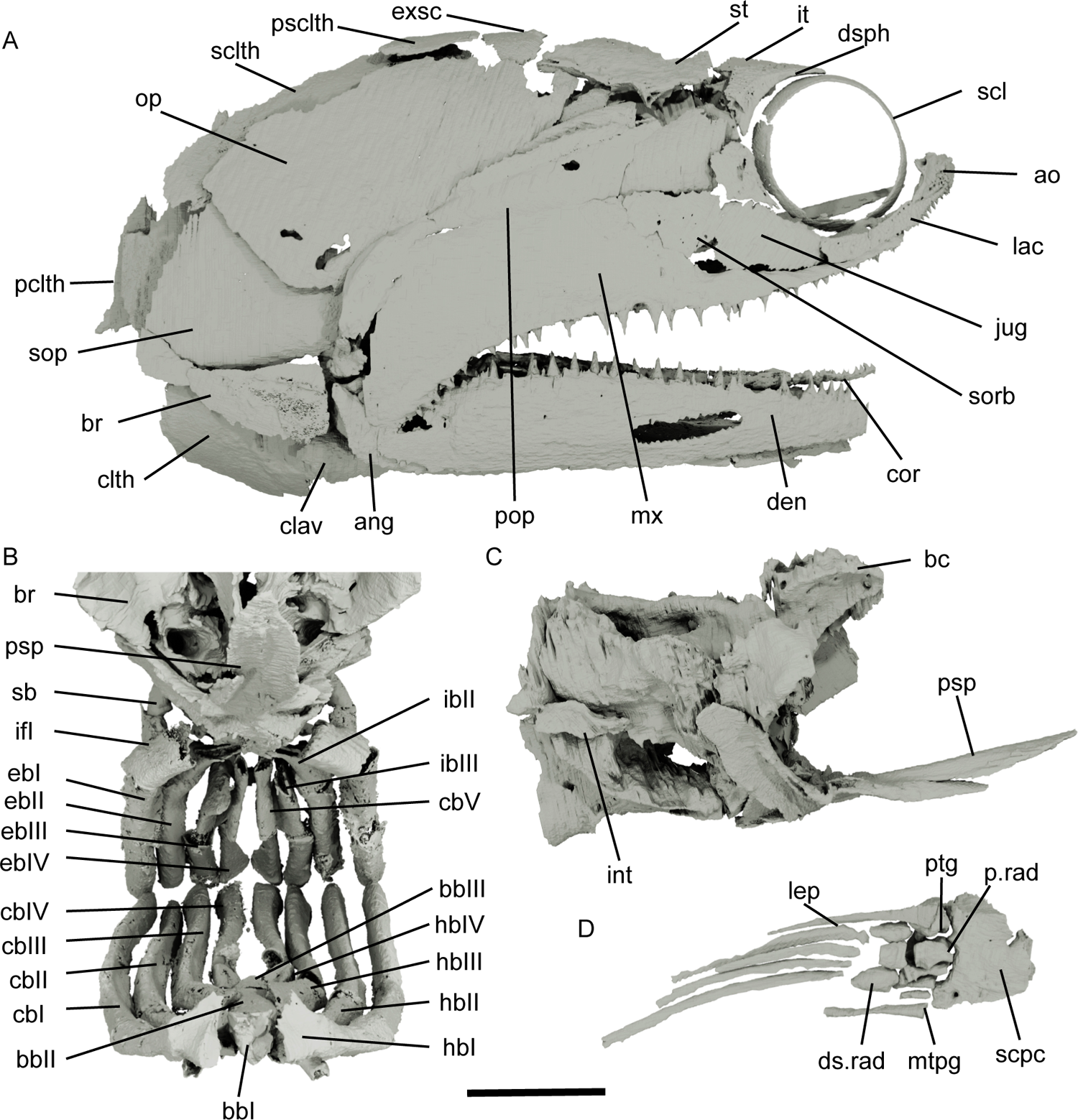
Strict consensus tree of 10752 most parsimonious trees, equal weights analysis with outgroup topology constrained. Tree length: 1693 steps. Consistency Index: 0.198; Retention Index: 0.674. Bremer indexes higher than 2 are indicated above nodes. Bootstrap percentages higher than 50% are indicated below nodes. Crown group nodes (black circle) and total groups (white circle) are labelled. Shading: purple, Chondrichthyes (total group); red, Sarcopterygii (total group); yellow, *Pteronisculus*; orange, Scanilepiformes+Cladistia; light blue, Chondrostei (crown-group); green, Teleostei (total group).

Actinopterygii is also recovered as monophyletic but receives very weak support (BDI: 1). *Meemannia* is resolved as the earliest actinopterygian, sister to a monophyletic *Cheirolepis* and all other actinopterygians. *Tegeolepis* and *Osorioichthys* fall as successive sister taxa to remaining actinopterygians (BDI: 3). Most other Palaeozoic taxa are recovered as singletons or small clades, but the topology is very weakly supported (BDI: 1-2). Several Carboniferous (*Trawdenia*, *Kansasiella*, *Coccocephalichthys*, *Lawrenciella*), Permian (*Luederia*) and Triassic (*Boreosomus*) taxa form a basal polytomy, sister to a broad clade that includes *Pteronisculus* and remaining actinopterygians. This clade, mostly made of Carboniferous taxa, is supported by an antorbital bone (ch. 50), two suborbitals (ch. 55), an operculum twice as high as the suboperculum (ch. 111) and jointed pectoral radials (ch. 247). We recover, at successive nodes: *Cosmoptychius* as the earliest-diverging member; a clade containing the rhadinichthyids *Cyranorhis* and *Wendyichthys*; and *Beagiascus* + *Aesopichthys* + *Kalops* forming the sister group to a monophyletic *Pteronisculus*. The topology of the actinopterygian stem, as well as this monophyletic clade, are recovered in our implied weights analysis in the same way (See Supplementary Material, Figure S1).

The monophyly of *Pteronisculus* is supported by the presence of more than two infraorbitals (ch. 52), mandibular canal arched dorsally in the anterior half of the mandible (ch. 77), dorsalmost branchiostegal ray deeper than the adjacent one (ch. 120), lacrimal contributing to oral margin (ch. 301) and presence of teeth on the lacrimal (ch. 302) (BDI: 2). Within the genus, the Chinese and Malagasy species cluster together with weak support, united by the lack of premaxillae (ch. 2). The Greenland species are the sister group of the other species (BDI: 2), supported by the absence of contact between extrascapulae at the midline (ch. 47), presence of a sensory canal/pit line of the maxilla (ch. 71) and presence of an accessory operculum (ch. 109). *P. gunnari* and NHMD–73588 are recovered as sister taxa (Fig. 18).

We recover *Australosomus* as the sister taxon to crown actinopterygians. Scanilepiformes and Cladistia, are recovered as a clade (BDI: 3), which in turn is the sister group to Actinopteri (Fig. 18). We recover a very weakly supported stem-chondrostean assemblage comprising *Brachydegma*, the macrodont *Brazilichthys*, birgeriids (BDI: 5) and saurichthyids (BDI: 5). The Carboniferous *Platysomus* and Eurynotiformes (*Fouldenia*, *Styracopterus, Amphicentrum*), as well as other deep bodied Carboniferous, Permian and Triassic forms (*Discoserra*, *Ebenaqua*, *Bobasatrania*) are resolved as stem neopterygians, but the topology of their relationships is weakly supported (BDI: 1-2) (Fig. 19) and in fact the order these taxa are recovered in our implied weights analysis is slightly different, while the relationships between Scanilepiformes, Cladistia and chondrosteans are largely confirmed (See Supplementary Material, Figure S1). Other relationships are largely as reported by Argyriou et al. (2022).

**Figure 19:**
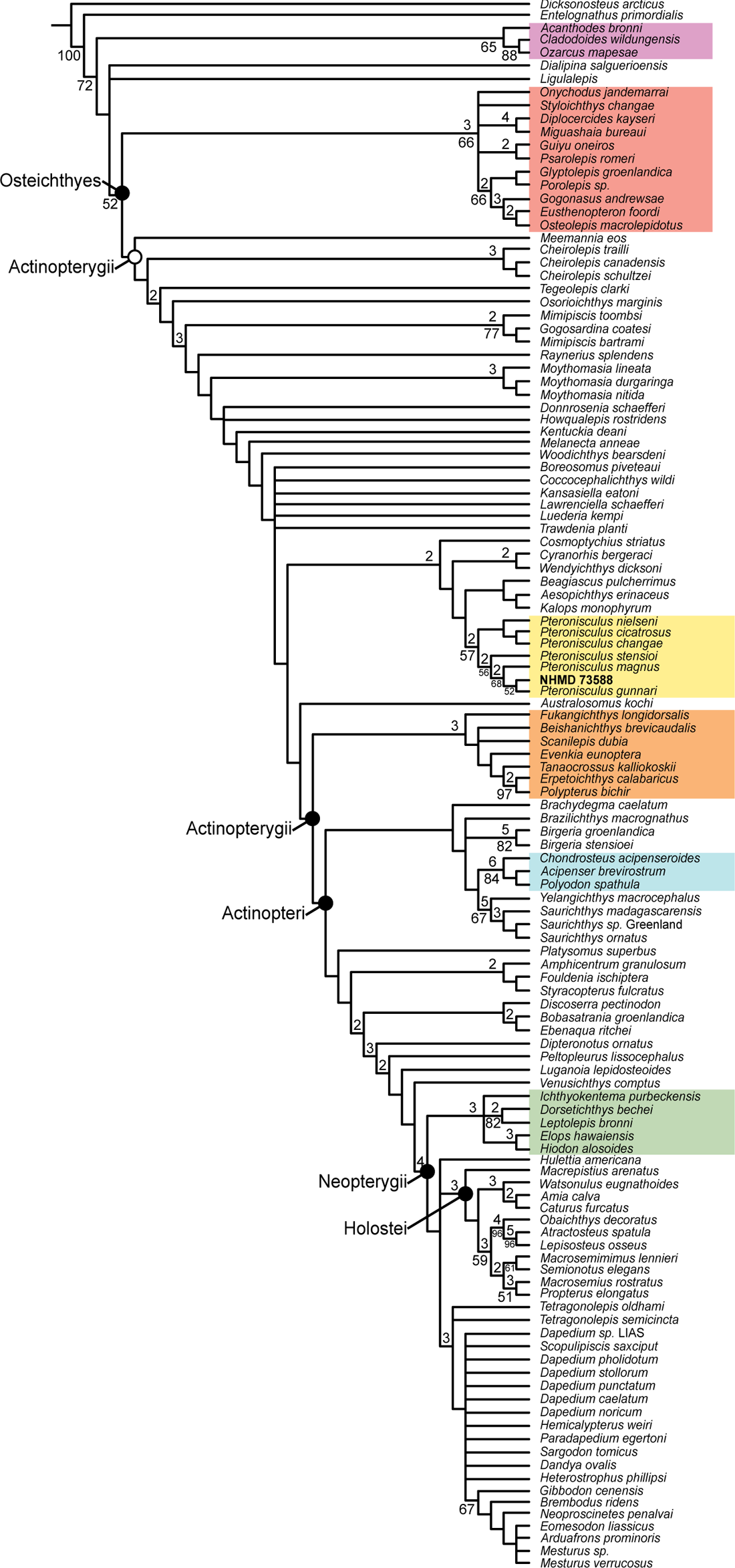
Composite reconstructions of the skeleton of *Pteronisculus gunnari* NHMD–73588. Skull and dermal pectoral girdle in right lateral view (A). Gill skeleton, parasphenoid and braincase in anterior view (B). Braincase, parasphenoid and intercalar in right lateral view (C). Right pectoral fin endoskeleton and selected lepidotrichia in ventrolateral view (D). Scale bar: 10 mm for A, D; 6 mm for B-C. Abbreviations: ang, angular; ao, antorbital; bbI-III, basibranchials; bc, braincase; br, branchiostegal ray; cbI-V, ceratobranchials; clth, cleithrum; clav, clavicle; cor, coronoid; ds.rad, distal radial; dsph, dermosphenotic; den, dentary; ebI-IV, epibranchials; exsc, extrascapular; hbI-IV, hypobranchials; ibI-III, infrapharyngobranchials; int, intercalar; it, intertemporal; jug, jugal; lac, lacrimal; lep, lepidotrichia; mtpg, metapterygium; mx, maxilla; op, operculum; p.rad, proximal radial; pclth, postcleithrum; pop, preoperculum; psclth, presupracleithrum; psp, parasphenoid; ptg, propterygium; sb, suprapharyngobranchials; scl, sclerotic ring; sclth, supracleithrum; scpc, scapulocoracoid; sorb, suborbital; sop, suboperculum; st, supratemporal.

## Discussion

### Updates to previous descriptions

Our redescription of the holotype of *Pteronisculus gunnari* adds meaningful details to our knowledge of this species, particularly regarding aspects of internal anatomy not observed by previous authors (Nielsen, 1942). The preservation is so detailed that it allowed us to reconstruct portions of the anatomy (Fig. 19).

### Snout

The presence of paired premaxillae is confirmed in *P. gunnari* for the first time. Premaxillae are tentatively described in *P. stensioi* and *P. magnus*, and the condition is unclear in other Greenlandic species owing to poor preservation of the snout region (Nielsen, 1942); for the same reason, the condition is reported as unknown by White (1933) for *P. cicatrosus.* Premaxillae are hypothetically reconstructed in the remaining Malagasy species, although not explicitly described by Lehman (1952). Although the premaxillae of *P. gunnari* host a cavity through the length of the bone (Fig. 3), these do not align with the antorbital infraorbital canal and are thus unlikely to have carried the ethmoidal commissure. The nature of the cavity is uncertain: since it only opens onto the medial surface of the bone, it is hard to associate it with sensory organs like electroreceptors (King et al., 2018).

### Parasphenoid

Previous accounts of the parasphenoid of *P. gunnari* are short (Nielsen, 1942). We provide a detailed account, recognising intrageneric diversity in *Pteronisculus*: reduced tooth covering of the anterior corpus, the presence of a dorsal bony pedicel for the buccohypophyseal canal and a ventral buccohypophyseal opening, foramina for the internal carotids, the presence of an enclosed median parabasal canal, and the absence of any kind of dermal basipterygoid processes distinguish *P. gunnari* from other species. In *P. magna* and *P. stensioi*, the dermal basipterygoid process is restricted to a sheath of dermal bone underlying the corresponding endoskeletal process (Lehman, 1952; Nielsen, 1942). In other palaeopterygians, such as the Carboniferous *Kentuckia* (Rayner, 1952), the process is entirely endoskeletal, and it is also absent in chondrosteans, amiids, Recent teleosts, and some neopterygian clades, where it may have been secondarily lost (Brian George Gardiner, 1984; Patterson, 1975). The lack of an endoskeletal basipterygoid process in this specimen of *P. gunnari* may be related to the submature condition of the specimen, rather than a true absence. Foramina for the internal carotids piercing the parasphenoid are a feature more commonly associated with neopterygians or teleosts (Brian George Gardiner, 1984; Patterson, 1975). The foramina are aligned with the posterior grooves on the lateral surface of the basisphenoid, confirming that the circulation partially passed through the median parabasal canal of the parasphenoid (Fig. 10, icg), like in e.g., *Saurichthys* (T. Argyriou et al., 2018), rather than in grooves on the dorsal surface of the anterior corpus, as was previously described for *P. magnus* (Nielsen, 1942). In other respects, the parasphenoid resembles that of generalised post-Devonian actinopterygians, as it does not extend past the ventral otic fissure (Brian George Gardiner, 1984; Patterson, 1975). The number of parotic toothplates associated with the parasphenoid is higher than previously reported for *P. gunnari*, with at least three pairs rather than one (Nielsen, 1942), although the peculiar morphologies are difficult to rearrange in life positions (Fig. 11).

### Braincase

Although the braincase is only partially ossified in NHMD–73588, we are able to largely confirm previous descriptions and provide new anatomical information, including of the ethmoids. Absence of a foramen on the floor of the posterior myodome, presence of an intercalar bone, and the presence of small canals between the anterior and horizontal ampullae, jugular canal and fossa bridgei, and various nerves are the most evident differences from previous reports (Nielsen, 1942). The craniospinal process of palaeopterygians was considered homologous to the intercalar of neopterygians, but lacking any dermal outgrowths (Patterson, 1975). An independent, membranous intercalar with an endochondral core was historically considered a neopterygian character, while it is entirely membranous in *Amia* and extant teleosts (Patterson, 1975; Pradel et al., 2016; Schaeffer, 1971). The discovery of an intercalar in the stem actinopterygians *Lawrenciella*, where it appears to be dermal (Pradel et al., 2016), possibly in *Palaeoneiros* (Giles et al., 2023a) and now in *Pteronisculus* with a dermal component, suggests that this ossification had a deeper origin, but may have been overlooked in previous descriptions of palaeopterygian braincases due to its small size or difficulties in preparation. The tomograms also show small canals connecting the trigeminal (V) and facial nerves (VII), the anterior and horizontal ampullae of the bony labyrinth, the trigeminal nerve (V) and the roof of the jugular canal, and the trigeminofacialis chamber and the floor of the fossa bridgei (Fig. 11). Additional microCT-based descriptions of Palaeozoic and Mesozoic actinopterygian braincases will be needed to assess whether these represent species-specific characters, variation due to incomplete ossification, or are more broadly distributed.

The unfinished nature of the endochondral ossification of the braincase allows us to infer the position of some of the primordial ossification centres, which are otherwise difficult to discern in the compact neurocrania of generalized Paleozoic–early Mesozoic actinopterygians (Nielsen 1942, 1949; Patterson 1975; but see some exceptions in Gardiner 1984; Argyriou et al. 2022). The ethmoids mineralize from independent ossification centres. The posteromedial portion of the interorbital wall is separated from the otic region by wide spaces and likely constitutes the pterosphenoid (Fig. 10); denser trabecular ossification around the canal for the trochlear nerve may indicate the original mineralization point. As the basisphenoid is present as a single element, and is most densely mineralized along the midline, it is most probably unpaired. Perichondral lining of the basisphenoid is largely complete, except for two anterodorsal facets and the ventral otic fissure, indicating it may have been the first to begin ossifying (Fig. 10, bsp). There are no clear boundaries or ossification centres that could be associated with a prootic ossification, and the area is overall poorly ossified. The postorbital process is separated from the rest of the braincase by two almost vertical bands of reduced ossification, in the prootic area and the hyomandibular facet. The very dense endochondral ossification in the dorsolateral portion of the postorbital process could represent the ossification centre for the sphenotic. There are no meaningful variations in the density of endochondral ossification in the rest of the otic region, so discriminating potential pterotic and opisthotic ossifications is difficult. In the occipital region, a horizontal band of less dense endochondral bone, dorsal to the occipital arteries, probably divides a ventral basioccipital from the exoccipitals, although there are no clear indications of where ossification likely initiated. There are no signs of the presence of a supraoccipital ossification centre. Our inferences show good agreement with Patterson (1975), who indicated where the ossification centres may lie on a braincase of *P. magnus* described by Nielsen (1942).

### Dermal cheek

Most dermal ossifications are fundamentally as described in previous accounts. However, tomograms show that several dermal bones, including the dermosphenotic (Fig. 4), dermohyal (Fig. 5) and horizontal shelf of the maxilla (Fig. 6), contain additional cavities that are separate from the sensory canals, and connect to the surface of the bone via small foramina. The nature of these cavities is unclear, but they may have supplemented the primary sensory canal network: either as remnant of the “p” canal for the dermosphenotic (Clement et al., 2018; Northcutt, 1989), or related to Lorenzinian ampullae, as was suggested for the Late Devonian *Howqualepis* and the stem osteichthyan ‘*Ligulalepis*’ (King et al., 2018; J. Long, 1988).

### Palate

Previous descriptions of *Pteronisculus* only reported one vomerine ossification (‘vomer’ of Nielsen, 1942, p.138), and the upper internal tooth rows were depicted as pointing ventrally (Lehman, 1952; Nielsen, 1942). We document the presence of a vomer (applied to the ventral surface of the ethmoids), and anterior and posterior accessory vomers, the latter corresponding with the ‘vomer’ (sensu Nielsen) of previous accounts (Fig. 9, 11). The internal tooth rows borne on the ectopterygoid and dermopalatines are directed medially in NHMD–73588, and the internal and external rows are separated by an edentulous space that accommodated the dentary teeth (Fig. 6-7). This condition, along with a tooth-bearing lacrimal, are also present in a macrodont actinopterygian from the Pennsylvanian of New Mexico (Friedman et al., in review). Discovering what this means in terms of the relationships between these two taxa will require further study.

### Lower jaw

The dentary of NHMD–73588 bears a low process, lateral to the midpoint of the adductor fossa (Fig. 9). A line drawing of *P. magna* (Nielsen 1942, p. 166) is suggestive of a bump on the surangular, but this structure is not described in the text. A similar small process formed by a single ossification of the lower jaw is reported in a small number of Palaeozoic taxa, including the prearticular of *Eurynotus* (Friedman et al., 2018), dentary of *Aesopichthys* (C. Poplin & Lund, 2000), and surangular of *Coccocephalichthys* (M. Poplin & Véran, 1996) and *Trawdenia planti* (M. I. Coates & Tietjen, 2018). Although analogous in position, the shape and composition of these processes suggests they are not homologous with the coronoid process of neopterygians (e.g. Thodoris Argyriou et al., 2022; Olsen, 1984; Xu, 2019), which is originally formed by the dentary and surangular, with modifications in various clades of teleosts (Arratia, 2015; Friedman, 2015; López-Arbarello & Sferco, 2018). However, it is likely that further investigation of palaeopterygians will uncover additional evidence for the widespread distribution this feature.

### Operculogular series

We report the presence of a modest anterodorsal process on the suboperculum. This process is sturdy in Devonian taxa like *Mimia*, *Moythomasia* or *Donnrosenia* (Brian George Gardiner, 1984; J. A. Long et al., 2008), but reduced to a weak strip in *P. gunnari*. The process is overlaid by the operculum and preoperculum, and CT-aided descriptions of additional taxa may indicate it is more widespread than previously thought.

### Gill skeleton

The gill skeleton of *P. gunnari* was not previously described, only equated to the description of *P. stensioi*, and the morphology of the basibranchial series for all species within the genus was largely unknown (Nielsen, 1942, p. 195). We describe the gill skeleton in its entirety, including the unfinished basibranchial series, and clarify the presence of uncinate processes on epibranchials II and III (contra Nielsen, 1942), the hook-like articulation of hypobranchial III with the basibranchial series, the simple polyhedral shape of hypobranchial IV and the long, bent shape of suprapharyngibranchial II (Fig. 13). The gill skeleton can be a useful diagnostic tool in extant fish, but its usefulness is limited in fossils due to a lack of detailed descriptions (Beckett et al., 2018). The development of a more extension comparative dataset may uncover new anatomical features of use in resolving relationships between early actinopterygians.

### Shoulder girdle and pectoral fin

We report a different course for the lateral line canal in the supracleithrum and provide the first description of the pectoral fin of *P. gunnari*, which was previously noted as identical to that of *P. magnus* (Nielsen, 1942). The preserved path of the lateral line canal through the supracleithrum of *P. gunnari* is more anterior than in other species, but the anterior portion of the bones has been replaced by pyrite and the complete course of the canal is not visible in the tomograms (Fig. 15). An additional cavity, separate from the sensory canal, as described in other dermal bones, is also present. We also clarified the shape of the propterygium and the first three proximal radials, and showed that the first lepidotrichium is fused to an unperforated propterygium, rather than articulated with it as previously described. In addition, we do not find that the fifth and sixth proximal radials are fused, as it was reported for a specimen of *P. gunnari* (Nielsen, 1942). Our updated description shows that the propterygium is imperforate (contra Jessen, 1972), a condition which is known, among palaeopterygians, only in *Cheirolepis* (Giles et al., 2015) and *Trawdenia*. In *Trawdenia*, the propterygial canal is suggested to have passed medial to the lateralmost propterygium, between it and the fused second radial (interpreted as a second propterygium by Coates & Tietjen, 2018). In addition, the first three radials of *P. gunnari* have an irregular morphology, reminiscent of Palaeozoic taxa like *Trawdenia*— except for the peculiar propterygial conformation (M. I. Coates & Tietjen, 2018)—or *Mimia* and *Moythomasia* (Brian George Gardiner, 1984).

### Axial skeleton

We confirm most details of the previous description, but clarify the size and shape of the epineural processes on the vertebrae (Fig. 17; “proximal plates” of Nielsen, 1942).

### Phylogenetic affinity of *Pteronisculus*

#### *Pteronisculus* and its taxonomic placement

Both our phylogenetic analyses recover a monophyletic *Pteronisculus*, albeit poorly supported, with NHMD–73588 forming the sister terminal to *P. gunnari*, supporting the specific attribution of Nielsen (1942). Two unambiguous synapomorphies support the monophyly of the genus: a toothed lacrimal, although we note that this has recently been reported in a Carboniferous macrodont (Friedman et al., in review); and a lacrimal contributing to the oral margin, which was reported in the original description of *Cyranorhis* by Lund and Poplin (1997) but considered absent in a later reassessment by Mickle (2015). This latter character is also purportedly present in the Late Triassic *Turseodus*, although this taxon is not included in our data matrix (Lund & Poplin, 1997; Schaeffer, 1952). Other characters supporting the monophyly of *Pteronisculus* in our phylogenetic analysis show variable degrees of homoplasy: the presence of more than two infraorbitals and the dorsalmost branchiostegal ray being deeper than the adjacent ray are common conditions in late Palaeozoic and early Mesozoic actinopterygians; and an anteriorly arched mandibular canal is present in *Australosomus*, some dapediids and Devonian forms like *Cheirolepis* and *Moythomasi*a.

Within the genus *Pteronisculus*, the Greenlandic species form a clade, sister to the Malagasy and Chinese species. While Early Triassic palaeogeography lends support to this (Scotese & Schettino, 2017), the latter clade is united only by the lack of premaxillae, which is fairly homoplastic among actinopterygians, (C. Poplin & Lund, 1997) and may have been misidentified in some *Pteronisculus* species as in *P. gunnari*. In addition, both clades have older species in terminal positions (*P. gunnari* and *P. cicatrosus;* (Lehman, 1952; Nielsen, 1942). New data in the future will hopefully clarify the relationships among *Pteronisculus* species.

We recover *Pteronisculus* in a broader clade that includes taxa from the Carboniferous of the United Kingdom and the United States. However, the characters supporting this are also homoplastic to a greater or lesser degree. The number of suborbitals and the relative dimensions of the operculum and suboperculum are highly variable across actinopterygians. Similarly, an independent antorbital (housing the infraorbital-ethmoidal commissure-supraorbital commissure) is common to many Carboniferous actinopterygians, as well as neopterygians (Brian George Gardiner, 1963; C. Poplin & Lund, 1995, 1997), and is optimized as evolving multiple times. In addition, this character unites many Palaeozoic taxa (including the majority of the *Pteronisculus* clade) for which the variability in the shape and relative positions of dermal snout bones is not fully captured in our analysis (e.g., vertical antorbital and tooth-bearing premaxilla in *Aesopichthys*, tooth-bearing antorbital and premaxilla in *Cosmoptychius* and *Watsonichthys*, absence of premaxilla and notched antorbital in *Cyranorhis* and *Wendichthys*, edentulous antorbital contributing to the oral margin and dentigerous premaxilla in *Kalops,* etc) (Brian George Gardiner, 1963; Lund & Poplin, 1997; C. Poplin & Lund, 2000; C. M. Poplin & Lund, 2002). Potential phylogenetic signals are obscured by the common poor preservation of the snout area in fossils and multiple, often conflicting bone naming conventions that make comparisons difficult (Kathryn E. Mickle, 2015; C. Poplin & Lund, 1995, 1997). We suggest that future phylogenetic analyses should incorporate characters that describe the features and arrangement of dermal bones of the snout in relation to morphological elements that are univocally identifiable (e.g., number of anamestic/canal bearing bones, contribution to oral/orbit margins, number of toothed ossifications, position relative to the nares, number of paired/unpaired ossifications, etc.).

The final character uniting this clade, the presence of jointed radials, is present (among actinopterygians) only in *Pteronisculus*, *Cheirolepis canadensis* and *Cosmoptychius* (Arratia & Cloutier, 1996; Brian George Gardiner, 1963), but the state is unknown in more than half of the taxa in our matrix, including most members of the *Pteronisculus* clade. While this may represent a robust synapomorphy, more data on the three-dimensional anatomy of palaeopterygian pectoral fins is needed to evaluate this.

The Palaeozoic affinities of *Pteronisculus* (e.g., rhadinichthyids, cosmoptychiids, macrodonts) are evident from our analysis, with the taxon recovered deep within a long-lived Carboniferous radiation which terminated in the Middle Triassic (Fig. 17).

Unfortunately, broader problems with early actinopterygian evolution persist, with most branches weakly supported and largely unresolved. On a small scale, this obscures relationships among different families, and on a larger scale, obscures evolutionary trends during the Palaeozoic-Mesozoic transition (López-Arbarello & Sferco, 2018; Lauren C. Sallan, 2014). A combination of descriptions of new taxa, updates to previous descriptions and improved phylogenetic matrices are needed to discern the relationships among Devonian and Carboniferous palaeopterygians, and palaeopterygians and neopterygians.

#### Differences from previous analyses

Most phylogenetic analyses do not include multiple species of *Pteronisculus*, and the genus is commonly used in isolation to represent a generalized non-neopterygian taxon (e.g. López-Arbarello & Sferco, 2018; Olsen, 1984; Xu, 2021). Where *Pteronisculus* has been included among a broader sample of palaeopterygians, its position has been variable: for example confirming its Palaeozoic affinities (Richard Cloutier & Arratia, 2004; Lund et al., 1995; Kathryn E Mickle, 2012); in a position close to the neopterygian stem (Brian G Gardiner et al., 2005; Wen et al., 2012); or in a broad polytomy (Hurley et al., 2007). The analyses that have employed iterations of the matrix used here, have recovered *Pteronisculus* either in an extensive polytomy containing Carboniferous-Permian taxa and the actinopterygian crown (Latimer & Giles, 2018); or associated with Carboniferous actinopterygians, either as sister taxon to multiple taxa (T. Argyriou et al., 2018; Ren & Xu, 2021) or as an early-diverging member of the radiation (Thodoris Argyriou et al., 2022; Giles et al., 2023a). The link between *Pteronisculus* and taxa from the Palaeozoic, in particular from the Carboniferous Bear Gulch fauna, is confirmed in our analysis. However, we recover *Pteronisculus* deeply nested within the Carboniferous radiation, rather than as an early diverging member.

Only Ren & Xu (2021) have previously included more than two species of *Pteronisculus* in a phylogenetic analysis (*P. stensioi*, *P. nielseni*, *P. changae*, *P. cicatrosus* and *P. magnus*). Their relationships and placement of *Pteronisculus* are broadly similar to our results, but the topology of the genus and the characters supporting it differ from our analysis. They recover the genus as closely related to *Cyranorhis* and in a clade comprising *Boreosomus*, *Trawdenia* (=*Mesopoma*; Coates & Tietjen, 2019), *Cosmoptychius*, *Beagiscus*, *Kalops* and *Aesopichthys*. Within the genus, *P. nielseni* is recovered as the earliest diverging, implying an eastern Tethys origin for *Pteronisculus*. In addition, (Ren & Xu, 2021) only report three characters supporting monophyly: a toothed lacrimal, presence of more than two infraorbitals and the supratemporal ending at the level of the posterior margin of the parietal. While Caron et al. (2023) include two species in their analysis (*P. stensioi* and *P. magnus*) they focus only on the neurocranium, and the notable differences between the braincase of the two species result in a paraphyletic *Pteronisculus*.

### Juvenile features and comparative anatomy

Several endoskeletal features of NHMD–73588 indicate it was a juvenile or immature specimen. Although complete growth series are known for some Palaeozoic and early Mesozoic fishes (e.g. R Cloutier, 2010a; Maxwell et al., 2018), juvenile actinopterygians are comparatively rare. Growth series of styracopterid actinopterygians from the Mississippian of Scotland reveals allometric growth of the fins and demonstrates an increase in the number of scales through ontogeny (Lauren Cole Sallan & Coates, 2013). Similarly, juvenile actinopterygians from Bear Gulch (Namurian) and Mazon Creek (Pennsylvanian) show scale covering spreading from the lateral line and the tail, and progressive ossification of the skull and pectoral girdle (Karen Anne Lowney, 1980; Schultze & Bardack, 1987). Isolated juvenile palaeopterygian specimens are relatively more common, and have been recognized by the reduced scale covering and/or skull dermal plate development (M. Coates, 1993; Hutchinson, 1973; Karen A Lowney, 1980), the degree of sensory canal enclosure (Maxwell et al., 2018) or limited endochondral ossification of the skull and pectoral girdle (Nielsen, 1942, 1949).

Here we summarise the key features of NHMD–73588 compared to fully mature individuals, along with implications for ontogenetic ossification patterns in Palaeozoic– early Mesozoic actinopterygians.

#### Endoskeleton: palate

In most palaeopterygians, the palatoquadrate ossified as a single element in the adult (Arratia & Schultze, 1991 and references therein; Brian George Gardiner, 1984), while in other coeval taxa (e.g., *Saurichthys*) the palatoquadrate ossifies as multiple elements (T. Argyriou et al., 2018). Some limited ontogenetic data with a bearing on early ossification patterns is known from Palaeozoic–early Mesozoic taxa, e.g. the Devonian *Mimia* (Brian George Gardiner, 1984), *Pteronisculus* and *Boreosomus* (Nielsen, 1942), and *Australosomus* (Nielsen, 1949), although a complete growth series is lacking. The condition in NHMD–73588, where the quadrate, metapterygoid and autopalatine are independently ossified, corroborates Nielsen’s report for some specimens of *Pteronisculus* (1942, p.143) and the previously mentioned ontogenetic data: ossification of the palatoquadrate began early in ontogeny with separate metapterygoid, quadrate and autopalatine ossification centres, which later fuse into a single element in the adult.

The palatoquadrate ossification patterns seen in extant non-teleost actinopterygians can be considered modifications of the palaeopterygian pattern. This is particularly evident in polypterids: the palatoquadrate of the Middle Triassic stem polypterid *Fukangichthys* is ossified as a single unit (Giles et al., 2017), while in *Polypterus*, the endoskeletal palate is originally a single cartilage, but it ossifies into separate autopalatine, metapterygoid and quadrate elements (Arratia & Schultze, 1991; M Jollie, 1984; Wacker et al., 2001). The condition in *Amia* is very similar, except that in the adult part of the cartilage persists between the three ossifications and the autopalatine ossifies late in ontogeny (Arratia & Schultze, 1991; Grande & Bemis, 1998). In *Lepisosteus*, the autopalatine is absent (Arratia & Schultze, 1991; Malcolm Jollie, 1984). In extant chondrosteans, the palatoquadrate is highly specialized, comprising the autopalatine plate, palatoquadrate bridge and quadrate flange, and remains cartilaginous through life (Tsessarsky, 2023; Warth et al., 2017).

A palatoquadrate ossified as a single unit in the adult, formed from three ossification centres during ontogeny, appears to be the primitive state in actinopterygians and probably for Osteichthyes (Brian George Gardiner, 1984). In all extant lineages (excluding chondrosteans), there is a similar trend of fragmentation of the palatoquadrate into three bones (Arratia & Schultze, 1991). NHMD–73588 confirms that *Pteronisculus* retains the primitive single palatoquadrate ossification, and also that the pattern of ossification involved three initial centres that are homologous to the bones in the palatoquadrate of more recent actinopterygians.

#### Endoskeleton: braincase

The braincase of early actinopterygians is well ossified in most mature specimens, with poorer endochondral and perichondral development in juvenile or immature specimens. Available data from fossil taxa suggests that the braincase is preformed in cartilage and ossification commences from multiple centres, proceeding towards the ethmoidal region during ontogeny; once maturity was reached, the braincase was normally composed of two ossifications separated by the oticoccipital fissure and ventral otic fissure (Caron et al., 2023; Brian George Gardiner, 1984; Giles et al., 2023a; Nielsen, 1942; Pradel et al., 2016). The ethmoidal region could, in some instances, ossify quite late, or not at all (M. Coates, 1998; Brian George Gardiner, 1984; Hamel & Poplin, 2008; Patterson, 1975). This pattern is not the only one known in early actinopterygians: macrodont taxa show limited or no ossification of the braincase, with independent ossifications separated by large gaps filled by cartilage in life (Thodoris Argyriou et al., 2022, Friedman et al. 2023 pending publication; Figueroa et al., 2019), and other taxa appear to show little or no endochondral ossification into adulthood (Giles et al., 2015; Giles et al., 2023b; Pearson & Westoll, 1979).

NHMD–73588 has a completely ossified occiput, some unfinished areas in the otic region, minimal ossification in the sphenoid region, a basisphenoid and a small ethmoid (Fig. 9, 11). This is in agreement with the ossification pattern seen in most early actinopterygians, but not entirely with past descriptions of the genus, in which the ethmoidal region is normally described as poorly ossified, or with ossification starting from the dorsal areas (Nielsen 1942; pp. 16-18, 58, 91-92, 97). The opening between the cerebral cavity, the anterior portion of the lateral cranial canal and the fossa bridgei (Fig. 9, lcc) is particularly noteworthy, as it indicates that in the pterotic and possibly the opisthotic and semicircular canals ossified before the endocranial walls and roof. This ephemeral connection also highlights the potential inaccuracies that can arise from incorrect interpretation of juvenile features (as discussed in Lauren Cole Sallan & Coates, 2013): in this case, an anterior connection of the lateral cranial canal and the cranial cavity is typically optimized as a character of neopterygians (Giles et al., 2018; Patterson, 1975), while in adult specimens of *Pteronisculus* and other palaeopterygians there is only a posterior connection through the loop of the posterior semicircular canal (Brian George Gardiner, 1984; Nielsen, 1942).

Braincase ossification in extant non-teleosts is markedly different from palaeopterygians: in *Acipenser* the braincase is largely cartilaginous, except in the largest individuals (Hilton et al., 2022; Hilton et al., 2011), while in *Polypterus,* amiids and gars the adult braincase is composed of discrete ossifications linked by greater or lesser expanses of cartilage (Bjerring, 1991; Brian George Gardiner, 1984; Grande & Bemis, 1998). The condition we see in NHMD–73588 confirms that the palaeopterygian braincase contained ossification centres homologous to those of later actinopterygians; the difference with more recent forms lies in the ossification process, which in palaeopterygians progressed to the point of complete fusion between all ossifications, save for the cranial fissures (Patterson, 1975). These considerations largely confirm previous hypotheses on the acquisition of the patterns seen in extant lineages (Arratia, 2015; Brian George Gardiner, 1984; López-Arbarello & Sferco, 2018; Patterson, 1975), with the addition that an independent intercalar has its roots outside of Neopterygii.

#### Endoskeleton: gill arches

As with other endoskeletal elements, ontogenetic growth of the basibranchial series begins with development of a perichondral rind and continues with endochondral ossification, although articular areas remain unfinished throughout life.

Nielsen (1942) describes *Pteronisculus* specimens in which the basibranchial series is poorly mineralized. A similar situation is reported for specimens of *Mimia* and *Moythomasia*, in which the basibranchial forms a single ossification in adults, but probably fused from three centres during ontogeny (Brian George Gardiner, 1984). In NHMD– 73588, basibranchial I probably represents the area between the articular surfaces for hypobranchials I and II, while basibranchial II is ossified between the articulation of hypobranchials II and III. The relationship between hypobranchials IV and basibranchial III is more difficult to assess given the small dimensions of the latter (Fig. 13). The incomplete ossification of the basibranchials is thus confirmed to be a good indicator of non-adult palaeopterygian specimens.

There is great variability in the development of the basibranchial series in extant non-teleosts. *Polypterus* has a single basibranchial cartilage, which ossifies as a single unit during ontogeny (Brian George Gardiner, 1984; M Jollie, 1984). Acipenseriformes have up to 5 cartilaginous basibranchials early in ontogeny: in time, three ossifications (copulae) coalesce, with the largest, most anterior one containing up to the first three cartilaginous elements (Warth et al., 2017). *Polyodon* and *Amia* have three cartilaginous basibranchials early in ontogeny, but while they remain cartilaginous in the former taxon, a portion of the first basibranchial ossifies between the articulations of the second and third gill arches in the latter (Grande & Bemis, 1991, 1998). Gars have two cartilaginous elements and only a portion of the second, bigger element ossifies during ontogeny (Grande, 2010; Malcolm Jollie, 1984).

The condition in NHMD–73588 appears to confirm a primitive actinopterygian condition comprising three cartilaginous elements with three initial ossification centres in the basibranchial series, which fuse into a single mineralized element in the earliest palaeopterygians (e.g. *Mimipiscis*, *Raynerius*) but remain as three separate ossified elements in more derived taxa, including *Pteronisculus*.

#### Endoskeleton: shoulder girdle

Detailed accounts of the endoskeletal components of palaeopterygian pectoral fins are rare (M. I. Coates & Tietjen, 2018; Giles et al., 2015; Giles et al., 2023b). The cleithrum also appears to develop very early in the ontogeny of palaeopterygians, further complicating the observation of the endoskeletal shoulder girdle in juvenile specimens (M. Coates, 1993; Karen A Lowney, 1980; Lauren Cole Sallan & Coates, 2013). Reports of incompletely ossified or unossified scapulocoracoids can be found in Nielsen (1942; 1949) for specimens of *Pteronisculus*, *Boreosomus* and *Australosomus*, but without any indication of the pattern of ossification. The condition in NHMD–73588 implies that the scapulocoracoid started to ossify around the dorsal muscle canal (Fig. 14C-E), but it remains unclear whether the ossification proceeded from one or multiple centres. Multiple scapulocoracoid plates associated with a larger ossification are also present in the small-bodied Famennian *Palaeoneiros*, which is interpreted as an adult or near-adult (Giles et al., 2023a), supporting the hypothesis that the scapulocoracoid formed from multiple ossification centres.

In extant non-teleosts, the scapulocoracoid of *Amia* persists as a single, cartilaginous element in all but the largest specimens, where small ossification centres may appear (Grande & Bemis, 1998); in Acipenseriformes, the single scapulocoracoid cartilage can ossify in adult specimens (Hilton & Bemis, 1999; Jollie, 1980). In *Lepisosteus*, the endochondral pectoral girdle is made of an ossified scapula and a cartilaginous mesocoracoid element (Malcolm Jollie, 1984). *Polypterus* has an endochondral pectoral girdle unique among actinopterygians, made of independent rod-like scapula and coracoid ossifications (M Jollie, 1984). Further information is needed on the endoskeletal girdles of palaeopterygians in order for this data to be interpreted in a wider context. However, despite the impossibility to confirm the presence of multiple ossification centres, the condition in the juvenile NHMD–73588 indicates that a fusion of possibly different centres would have happened early in ontogeny for palaeopterygians.

#### Dermal skeleton: squamation

Incomplete scale covering is a well-documented sign of young age for ray-finned fishes. In some cases, larval-stage specimens completely lack scales, and scale covering proceeds from the lobe of the caudal fin towards the head, or from around the cleithrum towards the tail; in either case, lateral line scales tend to form before other flank scales (R Cloutier, 2010b; Karen Anne Lowney, 1980; Lauren Cole Sallan & Coates, 2013; Schultze & Bardack, 1987). NHMD–73588 has an essentially complete scale covering and all dermal bones of the skull and pectoral girdle are present (Fig. 1G-H), indicating this specimen was probably closer to a sub-adult at the time of death. This is in line with the known growth series of palaeopterygians (Karen Anne Lowney, 1980; Schultze & Bardack, 1987), while also showing that, in the ontogeny of *Pteronisculus*, scale covering was completed before endochondral ossification.

In *Polypterus* and *Amia*, squamation development closely resembles that of palaeopterygians, with scales appearing first on the lateral line, then extending down to the tail and across the flanks (Bartsch et al., 1997; Grande & Bemis, 1998). A similar pattern is seen in *Lepisosteus*, with the first scales covering the lateral line in the posterior region of the body (Malcolm Jollie, 1984). *Acipenser* differs in having longitudinally-arranged plates (or scutes) rather than scales; the first row appears on the dorsal margin of the body, while on the flanks the plates start appearing anteriorly, and the rows develop towards the tail (Jollie, 1980). Apart from very specialized cases, squamation patterns seem very conservative among ray-finned fish, and more useful in identifying ontogenetic stages than taxonomic affinities.

NHMD–73588 adds a useful snapshot to our knowledge of ontogenetic changes in palaeopterygians, with implications for development in fossil and extant actinopterygians. In order to expand these considerations and map how ontogenetic development changes across clades, further descriptions of juveniles at a range of different ontogenetic stages, encompassing both dermal and endoskeletal development, are needed.

## Conclusion

Our redescription of *Pteronisculus gunnari* represents the first XCT-based reinvestigation of a member of the Triassic Early Fish Fauna. We reveal new anatomical details of the braincase, parasphenoid, palatoquadrate, gill skeleton and pectoral fin endoskeleton, improving our knowledge of *P. gunnari* and of the intrageneric diversity of *Pteronisculus*, while revealing many features reminiscent of Palaeozoic forms. The juvenile or sub-adult state of NHMD–73588 allowed us to draw useful comparisons on the endoskeletal development of palaeopterygians and extant non-teleosts. A revised phylogenetic analysis confirms the Palaeozoic affinities of *Pteronisculus*, implying a ghost lineage through the Permian and suggesting it is a poor candidate for generalized “non-neopterygian” anatomies. However, broader interrelationships of palaeopterygians remain unclear, as does the topology of neopterygian evolution. Future work should continue to focus on detailed redescriptions of Palaeozoic and early Mesozoic actinopterygians in tandem with revised and expanded phylogenetic analyses.

## Supporting information

Supplementary material phylogenetic analyses

## Acknowledgments

We thank Bent Lindow and Laura Cotton (Natural History Museum, Denmark for access to specimens and Liz Martin-Silverstone (University of Bristol) for assistance with CT scanning. We acknowledge the European Synchrotron Radiation Facility for provision of synchrotron radiation facilities under proposal number ES-1299, and we would like to thank BM05 staff for assistance in using the beamline. We thank Matt Friedman and Rodrigo Tinoco Figueroa (University of Michigan) and Soph Fasey (University of Birmingham) for assistance at the ESRF. IC was supported by a Central England NERC Training Alliance (CENTA) PhD scholarship via grant NE/S007350/1. SG was supported by a Royal Society Dorothy Hodgkin Research Fellowship (no. DH160098).

## Authors contributions

Conceptualisation: SG

Data curation: IC, VF, KD, SG

Formal analysis: IC, SG

Funding acquisition: SG

Investigation: IC, TA, VF, KD, SG

Methodology: IC, TA, SG

Project administration: SG

Resources: SG

Software: N/A

Supervision: SG

Validation: IC, SG

Visualisation: IC, SG

Writing: IC, SG

Writing – review and editing: IC, TA, VF, SG

## Data availability

The raw data (projections) for the synchrotron tomography scan of links: NHMD–73588 are deposited at: 10.15151/esrf-dc-1714814156. For review, segmentation files and surface files are available via temporary dropbox links: NHMD–73588 CT data .mcs file: https://www.dropbox.com/scl/fi/k4q9q4c38g2w518tu3yqu/NHMD_73588_Pteronisculus_gunnari_CT_.mcs?rlkey=dursljpby52uxh825tirzpodi&dl=0

NHMD–73588 CT data .ply files (zipped): https://www.dropbox.com/scl/fi/yquba4c6c38b1xbhfc3ov/NHMD_73588_Pteronisculus_gunnari_CT_ply_files.zip?rlkey=9i4e9zak63eqfxvk4v0672fhp&dl=0

NHMD–73588 synchrotron data .mcs file: https://www.dropbox.com/scl/fi/y3fis99falikghgagwire/NHMD_73588a_10.14um_normhalo_cropped_ESRF.mcs?rlkey=97gclmau4ljuq1a22f8xtilir&dl=0

NHMD–73588 synchrotron data .ply files (zipped): https://www.dropbox.com/scl/fi/xiv31cm4esz7qmsqe4696/NHMD_73588A_10.14um_normholo_synchro_plys.zip?rlkey=5sssixyi620uulxhlqjo4xf6r&dl=0

These, along with the tomogram TIFF stacks, will be deposited in a stable repository upon acceptance.

## Supplementary Material section

Additional supporting information can be accessed at: doi to be populated

## Bibliography

Argyriou, T., Giles, S., & Friedman, M. (2022). A Permian fish reveals widespread distribution of neopterygian-like jaw suspension. Elife, 11.

Argyriou, T., Giles, S., Friedman, M., Romano, C., Kogan, I., & Sanchez-Villagra, M. R. (2018). Internal cranial anatomy of Early Triassic species of daggerSaurichthys (Actinopterygii: daggerSaurichthyiformes): implications for the phylogenetic placement of daggersaurichthyiforms. BMC Evol Biol, 18(1), 161. 10.1186/s12862-018-1264-4

Arratia, G. (2015). Complexities of early Teleostei and the evolution of particular morphological structures through time. Copeia, 103(4), 999–1025.

Arratia, G., & Cloutier, R. (1996). Reassessment of the morphology of Cheirolepis canadensis (Actinopterygii). *Devonian fishes and plants of Miguasha, Quebec*, Canada, 165–197.

Arratia, G., & Schultze, H. P. (1991). Palatoquadrate and its ossifications: development and homology within osteichthyans. Journal of morphology, 208(1), 1–81.

Bartsch, P., Gemballa, S., & Piotrowski, T. (1997). The embryonic and larval development of Polypterus senegalusCuvier, 1829: its staging with reference to external and skeletal features, behaviour and locomotory habits. Acta Zoologica, 78(4), 309–328.

Beckett, H., Giles, S., & Friedman, M. (2018). Comparative anatomy of the gill skeleton of fossil Aulopiformes (Teleostei: Eurypterygii). Journal of Systematic Palaeontology, 16(14), 1221–1245.

Beltan, L. (1996). Overview of systematics, paleobiology, and paleoecology of Triassic fishes of northwestern Madagascar. Mesozoic Fishes− Systematics and Paleoecology, 479–500.

Bjerring, H. C. (1991). Two intracranial ligaments supporting the brain of the brachiopterygian fish Polypterus senegalus. Acta Zoologica, 72(1), 41–47.

Brinkmann, W., Romano, C., Bucher, D., & Ware, D. (2010). Palaeogeography and Stratigraphy of Advanced Gnathostomian Fishes (Chondrichthyes and Osteichthys) in the Early Triassic and from selected Anisian Localities (Report 1863 - 2009). Zentralblatt für Geologie und Paläontologie II, 2009(5/6), 765–812.

Caron, A., Venkataraman, V., Tietjen, K., & Coates, M. (2023). A fish for Phoebe: a new actinopterygian from the Upper Carboniferous Coal Measures of Saddleworth, Greater Manchester, UK, and a revision of Kansasiella eatoni. *Zoological Journal of the Linnean Society*, zlad011.

Cau, A., Beyrand, V., Voeten, D. F., Fernandez, V., Tafforeau, P., Stein, K., Barsbold, R., Tsogtbaatar, K., Currie, P. J., & Godefroit, P. (2017). Synchrotron scanning reveals amphibious ecomorphology in a new clade of bird-like dinosaurs. Nature, 552(7685), 395–399.

Chen, Z.-Q., & Benton, M. J. (2012). The timing and pattern of biotic recovery following the end-Permian mass extinction. Nature Geoscience, 5(6), 375–383. 10.1038/ngeo1475

Clement, A. M., King, B., Giles, S., Choo, B., Ahlberg, P. E., Young, G. C., & Long, J. A. (2018). Neurocranial anatomy of an enigmatic Early Devonian fish sheds light on early osteichthyan evolution. Elife, 7, e34349.

Cloutier, R. (2010a). The fossil record of fish ontogenies: Insights into developmental patterns and processes. (Ed.),^(Eds.). Seminars in cell & developmental biology.

Cloutier, R. (2010b). The fossil record of fish ontogenies: Insights into developmental patterns and processes. (Ed.),^(Eds.). Seminars in cell developmental biology.

Cloutier, R., & Arratia, G. (2004). Early diversification of actinopterygians. Recent advances in the origin and early radiation of vertebrates, 217–270.

Coates, M. (1993). Actinopterygian and acanthodian fishes from the Viséan of East Kirkton, West Lothian, Scotland. Earth and Environmental Science Transactions of the Royal Society of Edinburgh, 84(3-4), 317–327.

Coates, M. (1998). Actinopterygians from the Namurian of Bearsden, Scotland, with comments on early actinopterygian neurocrania. Zoological Journal of the Linnean Society, 122(1-2), 27–59.

Coates, M. I., & Tietjen, K. (2018). ‘This strange little palaeoniscid’: a new early actinopterygian genus, and commentary on pectoral fin conditions and function. Earth and Environmental Science Transactions of the Royal Society of Edinburgh, 109(1-2), 15–31.

Erwin, D. H., Bowring, S. A., & Yugan, J. (2002). End-Permian mass extinctions: a review. Special Papers-Geological Society of America, 363–384.

Figueroa, R. T., Friedman, M., & Gallo, V. (2019). Cranial anatomy of the predatory actinopterygian Brazilichthys macrognathus from the Permian (Cisuralian) Pedra de Fogo Formation, Parnaíba Basin, Brazil. Journal of Vertebrate Paleontology, 39(3), e1639722.

Friedman, M. (2015). The early evolution of ray-finned fishes. Palaeontology, 58(2), 213–228. 10.1111/pala.12150

Friedman, M., & Giles, S. (2016). Actinopterygians: The Ray-Finned Fishes—An Explosion of Diversity. In J. A. Clack, R. R. Fay, & A. N. Popper (Eds.), Evolution of the Vertebrate Ear: Evidence from the Fossil Record (pp. 17-49). Springer International Publishing. 10.1007/978-3-319-46661-3_2

Friedman, M., Pierce, S. E., Coates, M., & Giles, S. (2018). Feeding structures in the ray-finned fish Eurynotus crenatus (Actinopterygii: Eurynotiformes): implications for trophic diversification among Carboniferous actinopterygians. Earth and Environmental Science Transactions of the Royal Society of Edinburgh, 109(1-2), 33–47.

Friedman, M., & Sallan, L. C. (2012). Five hundred million years of extinction and recovery: A phanerozoic survey of large-scale diversity patterns in fishes. Palaeontology, 55(4), 707–742. 10.1111/j.1475-4983.2012.01165.x

Gardiner, B. G. (1963). Certain palaeoniscoid fishes and the evolution of the snout in actinopterygians. *Bulletin of the British Museum (Natural History)*, Geology Series, 8, 2 pls.

Gardiner, B. G. (1984). The relationships of the palaeoniscid fishes, a review based on new specimens of Mimia and Moythomasia from the Upper Devonian of Western Australia. *Bulletin of the British Museum (Natural History)*, Geology Series, 37(4), 173–428.

Gardiner, B. G., & Jubb, R. A. (1975). A new palaeoniscid from the Lower Beaufort Series of South Africa. South African Museum.

Gardiner, B. G., Schaeffer, B., & Masserie, J. A. (2005). A review of the lower actinopterygian phylogeny. Zoological Journal of the Linnean Society, 144(4), 511–525.

Giles, S., Coates, M. I., Garwood, R. J., Brazeau, M. D., Atwood, R., Johanson, Z., & Friedman, M. (2015). Endoskeletal structure in Cheirolepis (Osteichthyes, Actinopterygii), An early ray-finned fish. Palaeontology, 58(5), 849–870. 10.1111/pala.12182

Giles, S., Feilich, K., Warnock, R. C., Pierce, S. E., & Friedman, M. (2023a). A Late Devonian actinopterygian suggests high lineage survivorship across the end-Devonian mass extinction. Nature Ecology & Evolution, 7(1), 10–19.

Giles, S., Feilich, K., Warnock, R. C., Pierce, S. E., & Friedman, M. (2023b). A Late Devonian actinopterygian suggests high lineage survivorship across the end-Devonian mass extinction. Nature Ecology Evolution, 7(1), 10–19.

Giles, S., Rogers, M., & Friedman, M. (2018). Bony labyrinth morphology in early neopterygian fishes (Actinopterygii: Neopterygii). Journal of morphology, 279(4), 426–440.

Giles, S., Xu, G.-H., Near, T. J., & Friedman, M. (2017). Early members of ‘living fossil’lineage imply later origin of modern ray-finned fishes. Nature, 549(7671), 265–268.

Goloboff, P. A., Farris, J. S., & Nixon, K. C. (2008). TNT, a free program for phylogenetic analysis. Cladistics, 24(5), 774–786.

Goloboff, P. A., & Morales, M. E. (2023). TNT version 1.6, with a graphical interface for MacOS and Linux, including new routines in parallel. Cladistics, 39(2), 144–153.

Goloboff, P. A., Torres, A., & Arias, J. S. (2018). Weighted parsimony outperforms other methods of phylogenetic inference under models appropriate for morphology. Cladistics, 34(4), 407–437.

Grande, L. (2010). An empirical synthetic pattern study of gars (Lepisosteiformes) and closely related species, based mostly on skeletal anatomy. The resurrection of Holostei. Ichthyology & Herpetology, 10(2A), 1.

Grande, L., & Bemis, W. E. (1991). Osteology and phylogenetic relationships of fossil and recent paddlefishes (Polyodontidae) with comments on the interrelationships of Acipenseriformes. Journal of Vertebrate Paleontology, 11(S1), 1–121.

Grande, L., & Bemis, W. E. (1998). A comprehensive phylogenetic study of amiid fishes (Amiidae) based on comparative skeletal anatomy. An empirical search for interconnected patterns of natural history. Journal of Vertebrate Paleontology, 18(sup1), 1–696.

Hamel, M.-H., & Poplin, C. (2008). The braincase anatomy of Lawrenciella schaefferi, actinopterygian from the Upper Carboniferous of Kansas (USA). Journal of Vertebrate Paleontology, 28(4), 989–1006.

Henderson, S., Dunne, E. M., Fasey, S. A., & Giles, S. (2023). The early diversification of ray[finned fishes (Actinopterygii): hypotheses, challenges and future prospects. Biological Reviews, 98(1), 284–315.

Henderson, S., Dunne, E. M., & Giles, S. (2022). Sampling biases obscure the early diversification of the largest living vertebrate group. Proceedings of the Royal Society B, 289(1985), 20220916.

Hilton, E. J., & Bemis, W. E. (1999). Skeletal variation in shortnose sturgeon (Acipenser brevirostrum) from the Connecticut River: implications for comparative osteological studies of fossil and living fishes. Mesozoic fishes, 2, 69–94.

Hilton, E. J., Dillman, C. B., Paraschiv, M., & Suciu, R. (2022). Cranial morphology of the Stellate Sturgeon, Acipenser stellatus Pallas 1771 (Acipenseriformes, Acipenseridae), with notes on the skulls of other sturgeons. Acta Zoologica, 103(1), 57–77.

Hilton, E. J., Grande, L., & Bemis, W. E. (2011). Skeletal anatomy of the shortnose sturgeon, Acipenser brevirostrum Lesueur, 1818, and the systematics of sturgeons (Acipenseriformes, Acipenseridae). Fieldiana Life and Earth Sciences, 2011(3), 1–168.

Hurley, I. A., Mueller, R. L., Dunn, K. A., Schmidt, E. J., Friedman, M., Ho, R. K., Prince, V. E., Yang, Z., Thomas, M. G., & Coates, M. I. (2007). A new time-scale for ray-finned fish evolution. Proceedings of the Royal Society B: Biological Sciences, 274(1609), 489–498. 10.1098/rspb.2006.3749

Hutchinson, P. (1973). A REVISION OF THE REDFIELDIIFORM AND PERLEIDIFORM FISHES FROM THE TRIASSIC OF BEKKER’S KRAAL (SOUTH AFRICA) AND BROOKVALE (NEW SOUTH WALES.

Jessen, H. L. (1972). Schultergürtel und Pectoralflosse bei Actinopterygiern (Schultergürtel und Pectoralflosse bei Actinopterygiern (pp. 1-101).

Jollie, M. (1980). Development of head and pectoral girdle skeleton and scales in Acipenser. Copeia, 226–249.

Jollie, M. (1984). Development of cranial and pectoral girdle bones of Lepisosteus with a note on scales. Copeia, 476–502.

Jollie, M. (1984). Development of the head and pectoral skeleton of Polypterus with a note on scales (Pisces: Actinopterygii). Journal of Zoology, 204(4), 469–507.

King, B., Hu, Y., & Long, J. A. (2018). Electroreception in early vertebrates: survey, evidence and new information. Palaeontology, 61(3), 325–358.

Lambe, L. M. (1916). Ganoid fishes from near Banff, Alberta. Transactions of Royal Society of Canada, ser. 3, *sect. 4*, *10*, 35–44.

Latimer, A. E., & Giles, S. (2018). A giant dapediid from the Late Triassic of Switzerland and insights into neopterygian phylogeny. Royal Society open science, 5(8), 180497.

Lehman, J.-P. (1952). Etude complémentaire des poissons de l’Eotrias de Madagascar. Kungliga Svenska Vetenskaps-akademiens Hangdlingar, 2(6), 1–201.

Long, J. (1988). New palaeoniscoid fishes from the Late Devonian and Early Carboniferous of Victoria.

Long, J. A., Choo, B., & Young, G. C. (2008). A new basal actinopterygian fish from the Middle Devonian Aztec Siltstone of Antarctica. Antarctic Science, 20(4), 393–412.

López-Arbarello, A., & Sferco, E. (2018). Neopterygian phylogeny: the merger assay. Royal Society open science, 5(3), 172337.

Lowney, K. A. (1980). Certain Bear Gulch (Namurian A, Montana) Actinopterygii (Osteichthyes) and a reevaluation of the evolution of the Paleozoic actinopterygians. New York University.

Lowney, K. A. (1980). A revision of the family Haplolepidae (Actinopterygii, Paleonisciformes) from Linton, Ohio (Westphalian D, Pennsylvanian). Journal of Paleontology, 942–953.

Lund, R., & Poplin, C. (1997). The rhadinichthyids (paleoniscoid actinopterygians) from the bear gulch limestone of Montana (USA, lower carboniferous). Journal of Vertebrate Paleontology, 17(3), 466–486.

Lund, R., Poplin, C., & McCarthy, K. (1995). Preliminary analysis of theinterrelationships of some paleozoic actinopterygii. Geobios, 28, 215–220.

Lyckegaard, A., Johnson, G., & Tafforeau, P. (2011). Correction of ring artifacts in X-ray tomographic images. International Journal of Tomography and Statistics, 18(F11), 1–9.

Maxwell, E. E., Argyriou, T., Stockar, R., & Furrer, H. (2018). Re[evaluation of the ontogeny and reproductive biology of the Triassic fish Saurichthys (Actinopterygii, Saurichthyidae). Palaeontology, 61(4), 559–574.

Mickle, K. E. (2012). Unraveling the Systematics of Palaeoniscoid Fishes--Lower Actinopterygians in Need of a Complete Phylogenetic Revision, University of Kansas].

Mickle, K. E. (2015). Identification of the Bones of the Snout in Fossil Lower Actinopterygians—A New Nomenclature Scheme Based on Characters. Copeia, 103(4), 838–857. 10.1643/cg-14-110

Mirone, A., Brun, E., Gouillart, E., Tafforeau, P., & Kieffer, J. (2014). The PyHST2 hybrid distributed code for high speed tomographic reconstruction with iterative reconstruction and a priori knowledge capabilities. Nuclear Instruments and Methods in Physics Research Section B: Beam Interactions with Materials and Atoms, 324, 41–48.

Nielsen, E. (1942). Studies on Triassic Fishes from East Greenland 1. Glaucolepis and Boreosomus (Vol. 3). Palaeozoologica Groenlandica.

Nielsen, E. (1949). Studies on Triassic Fishes from East Greenland: Australosomus and Birgeria. Reitzel.

Northcutt, R. G. (1989, 1989//). The Phylogenetic Distribution and Innervation of Craniate Mechanoreceptive Lateral Lines. (Ed.),^(Eds.). The Mechanosensory Lateral Line, New York, NY.

Olsen, P. E. (1984). The skull and pectoral girdle of the parasemionotid fish Watsonulus Eugnathoides from the Early Triassic Sakamena Group of Madagascar, with comments on the relationships of the holostean fishes. Journal of Vertebrate Paleontology, 4(3), 481–499. 10.1080/02724634.1984.10012024

Paganin, D., Mayo, S. C., Gureyev, T. E., Miller, P. R., & Wilkins, S. W. (2002). Simultaneous phase and amplitude extraction from a single defocused image of a homogeneous object. Journal of microscopy, 206(1), 33–40.

Patterson, C. (1975). The braincase of pholidophorid and leptolepid fishes, with a review of the actinopterygian braincase. *Philosophical Transactions of the Royal Society of London. B*, Biological Sciences, 269(899), 275–579.

Pearson, D. M., & Westoll, T. S. (1979). The Devonian actinopterygian Cheirolepis Agassiz. Earth and Environmental Science Transactions of the Royal Society of Edinburgh, 70(13-14), 337–399.

Poplin, C., & Lund, R. (1995). Fates of the rostral; postrostral and premaxillary in the early history of actinopterygians. Geobios, 28, 225–230.

Poplin, C., & Lund, R. (1997). Evolution of the premaxillary in the primitive fossil actinopterygians. Geodiversitas, 19(3), 557–565.

Poplin, C., & Lund, R. (2000). Two new deep-bodied palaeoniscoid actinopterygians from Bear Gulch (Montana, USA, Lower Carboniferous). Journal of Vertebrate Paleontology, 20(3), 428–449. 10.1671/0272-4634(2000)020[0428:TNDBPA]2.0.CO;2

Poplin, C. M., & Lund, R. (2002). Two Carboniferous fine-eyed palaeoniscoids (Pisces, Actinopterygii) from Bear Gulch (USA). Journal of Paleontology, 76(6), 1014–1028. 10.1666/0022-3360(2002)0761014:TCFEPP2.0.CO;2

Poplin, M., & Véran, M. (1996). A revision of the actinopterygian fish Coccocephalus wildi from the Upper Carboniferous of Lancashire. Special Papers in Palaeontology, 52, 7–30.

Pradel, A., Maisey, J. G., Mapes, R. H., & Kruta, I. (2016). First evidence of an intercalar bone in the braincase of “palaeonisciform” actinopterygians, with a virtual reconstruction of a new braincase of Lawrenciella Poplin, 1984 from the Carboniferous of Oklahoma. Geodiversitas, 38(4), 489–504.

Rayner, D. H. (1952). III.—On the Cranial Structure of an Early Palæoniscid, Kentuckia, gen. nov. Earth and Environmental Science Transactions of the Royal Society of Edinburgh, 62(1), 53–83. 10.1017/S0080456800009248

Ren, Y., & Xu, G. (2021). A new species of Pteronisculus from the Middle Triassic (Anisian) of Luoping, Yunnan, China, and phylogenetic relationships of early actinopterygian fishes. Vertebrata PalAsiatica, 59(3), 169–199.

Romano, C. (2021). A Hiatus Obscures the Early Evolution of Modern Lineages of Bony Fishes [Review]. Frontiers in Earth Science, 8. 10.3389/feart.2020.618853

Romano, C., Koot, M. B., Kogan, I., Brayard, A., Minikh, A. V., Brinkmann, W., Bucher, H., & Kriwet, J. (2016). Permian-Triassic Osteichthyes (bony fishes): Diversity dynamics and body size evolution. Biological Reviews, 91(1), 106–147. 10.1111/brv.12161

Romano, C., López-Arbarello, A., Ware, D., Jenks, J. F., & Brinkmann, W. (2019). Marine Early Triassic Actinopterygii from the Candelaria Hills (Esmeralda County, Nevada, USA). Journal of Paleontology, 93(5), 971–1000. 10.1017/jpa.2019.18

Sallan, L. C. (2014). Major issues in the origins of ray-finned fish (Actinopterygii) biodiversity. Biological Reviews, 89(4), 950–971. 10.1111/brv.12086

Sallan, L. C., & Coates, M. I. (2013). Styracopterid (Actinopterygii) ontogeny and the multiple origins of post-Hangenberg deep-bodied fishes. Zoological Journal of the Linnean Society, 169(1), 156–199.

Schaeffer, B. (1952). The palaeoniscoid fish Turseodus from the Upper Triassic Newark group. American Museum novitates; no. 1581.

Schaeffer, B. (1971). The braincase of the holostean fish Macrepistius, with comments on neurocranial ossification in the Actinopterygii. American Museum novitates; no. 2459.

Schaeffer, B., & Mangus, M. (1976). An early Triassic fish assemblage from British Columbia. Bulletin of the American Museum of Natural History, 156(5), 519–563. http://digitallibrary.amnh.org/dspace/handle/2246/619

Schultze, H.-P., & Bardack, D. (1987). Diversity and size changes in palaeonisciform fishes (Actinopterygii, Pisces) from the Pennsylvanian Mazon Creek Fauna, Illinois, USA. Journal of Vertebrate Paleontology, 7(1), 1–23.

Scotese, C., & Schettino, A. (2017). Late Permian-Early Jurassic paleogeography of western Tethys and the world (Permo-Triassic salt provinces of Europe, North Africa and the Atlantic margins (pp. 57-95). Elsevier.

Smithwick, F. M., & Stubbs, T. L. (2018). Phanerozoic survivors: Actinopterygian evolution through the Permo[Triassic and Triassic[Jurassic mass extinction events. Evolution, 72(2), 348–362.

Stensio, E. A. (1925). Triassic fishes from Spitzbergen. JG, 33(7), 753–753.

Stensiö, E. A. (1921). Triassic fishes from Spitzbergen. A. Holzhausen.

Stensiö, E. A. (1932). Triassic Fishes from East Greenland: Collected by the Danish Expeditions in 1929-31:[de Østgrønlandske Expeditioner Til Kong Christian Den X’s Land Udført i Aarene 1926-27 Od 1929-30 Under Ledelse Af Lauge Koch]. Reitzel.

Surlyk, F., Bjerager, M., Piasecki, S., & Stemmerik, L. (2017). Stratigraphy of the marine Lower Triassic succession at Kap Stosch, Hold with Hope, North-East Greenland. Bulletin of the Geological Society of Denmark, 65, 87–123.

Tintori, A., Hitij, T., Jiang, D., Lombardo, C., & Sun, Z. (2014). Triassic actinopterygian fishes: The recovery after the end-Permian crisis. Integrative Zoology, 9(4), 394–411. 10.1111/1749-4877.12077

Tsessarsky, A. (2023). What is missing from the sturgeon jaw: Developmental morphology of the upper jaw in Acipenser. Journal of Anatomy.

Vázquez, P., & Clapham, M. E. (2017). Extinction selectivity among marine fishes during multistressor global change in the end-Permian and end-Triassic crises. Geology, 45(5), 395–398.

Wacker, K., Bartsch, P., & Clemen, G. (2001). The development of the tooth pattern and dentigerous bones in Polypterus senegalus (Cladistia, Actinopterygii). Annals of Anatomy-Anatomischer Anzeiger, 183(1), 37–52.

Warth, P., Hilton, E. J., Naumann, B., Olsson, L., & Konstantinidis, P. (2017). Development of the skull and pectoral girdle in S iberian sturgeon, Acipenser baerii, and R ussian sturgeon, Acipenser gueldenstaedtii (A cipenseriformes: A cipenseridae). Journal of morphology, 278(3), 418–442.

Wen, W., Zhang, Q.-Y., Hu, S.-X., Zhou, C.-Y., Xie, T., Huang, J.-Y., Chen, Z. Q., & Benton, M. J. (2012). A new basal actinopterygian fish from the Anisian (Middle Triassic) of Luoping, Yunnan Province, southwest China. Acta Palaeontologica Polonica, 57(1), 149–160.

White, E. I., & Moy-Thomas, J. A. (1940). VII.—Notes on the Nomenclature of Fossil Fishes.—Part II. Homonyms D-L. Annals and Magazine of Natural History, 6(31), 98–103. 10.1080/03745481.1940.9723659

Xu, G.-H. (2019). Osteology and phylogeny of Robustichthys luopingensis, the largest holostean fish in the Middle Triassic. PeerJ, 7, e7184.

Xu, G.-H. (2021). A new stem-neopterygian fish from the Middle Triassic (Anisian) of Yunnan, China, with a reassessment of the relationships of early neopterygian clades. Zoological Journal of the Linnean Society, 191(2), 375–394.

Xu, G.-H., Gao, K.-Q., & Finarelli, J. A. (2014). A revision of the Middle Triassic scanilepiform fish Fukangichthys longidorsalis from Xinjiang, China, with comments on the phylogeny of the Actinopteri. Journal of Vertebrate Paleontology, 34(4), 747–759.

Xu, G.-H., Shen, C.-C., & Zhao, L.-J. (2014). Pteronisculus nielseni sp. nov., a new stem-actinopteran fish from the Middle Triassic of Luoping, Yunnan Province, China. 古脊椎动物学报, 52(4), 364–380.

