## Supplementary material phylogenetic analyses for "A Portrait of a Young Fish: Redescription of *Pteronisculus gunnari* (Nielsen, 1942) from a juvenile specimen from the Early Triassic of East Greenland, with implications for ontogenetic development in early actinopterygians": Cavicchini_et_al_2024 copy.txt

#NEXUS
[written Mon Dec 18 11:24:06 GMT 2023 by Mesquite  version 3.61 (build 927) at MacBook-Pro-di-Iacopo.local/127.0.0.1]

BEGIN TAXA;
	TITLE Taxa;
	DIMENSIONS NTAX=123;
	TAXLABELS
		Acanthodes_bronni Acipenser_brevirostrum Aesopichthys_erinaceus Amia_calva Amphicentrum_granulosum_ Atractosteus_spatula Australosomus_kochi Beagiascus_pulcherrimus Beishanichthys_brevicaudalis Birgeria_groenlandica Birgeria_stensioei Bobasatrania_groenlandica Boreosomus_piveteaui Brazilichthys_macrognathus Caturus_furcatus Cheirolepis_canadensis Cheirolepis_schultzei Cheirolepis_trailli Chondrosteus_acipenseroides Cladodoides_wildungensis Coccocephalichthys_wildi Cosmoptychius_striatus Cyranorhis_bergeraci Dapedium_sp_LIAS Scopulipiscis_saxciput Dapedium_pholidotum Dapedium_stollorum Dapedium_punctatum Dapedium_caelatum Dapedium_noricum Hemicalypterus_weiri Paradapedium_egertoni Sargodon_tomicus Dandya_ovalis Heterostrophus_phillipsi Tetragonolepis_oldhami Dialipina_salguerioensis Dicksonosteus_arcticus Diplocercides_kayseri Dipteronotus_ornatus Discoserra_pectinodon Donnrosenia_schaefferi Dorsetichthys_bechei_ Ebenaqua_ritchei Elops_hawaiensis Entelognathus_primordialis Erpetoichthys_calabaricus Eusthenopteron_foordi Evenkia_eunoptera Fouldenia_ischiptera Fukangichthys_longidorsalis Glyptolepis_groenlandica Gogonasus_andrewsae Gogosardina_coatesi Guiyu_oneiros Hiodon_alosoides Howqualepis_rostridens Hulettia_americana Ichthyokentema_purbeckensis Kalops_monophyrum Kansasiella_eatoni_ Kentuckia_deani Lawrenciella_schaefferi Lepisosteus_osseus Leptolepis_bronni '''Ligulalepis''' Luederia_kempi Luganoia_lepidosteoides Macrepistius_arenatus Macrosemimimus_lennieri Macrosemius_rostratus Meemannia_eos Melanecta_anneae Trawdenia_planti Miguashaia_bureaui Mimipiscis_bartrami Mimipiscis_toombsi Moythomasia_durgaringa Moythomasia_lineata Moythomasia_nitida Obaichthys_decoratus Onychodus_jandemarrai Osorioichthys_marginis Osteolepis_macrolepidotus Ozarcus_mapesae Peltopleurus_lissocephalus Platysomus_superbus Polyodon_spathula Polypterus_bichir Porolepis_sp. Propterus_elongatus Psarolepis_romeri Pteronisculus_stensioi Raynerius_splendens Saurichthys_madagascarensis Saurichthys_Greenland Saurichthys_ornatus Scanilepis_dubia Semionotus_elegans Styloichthys_changae Styracopterus_fulcratus Tanaocrossus_kalliokoskii Tegeolepis_clarki Tetragonolepis_semicincta Venusichthys_comptus Watsonulus_eugnathoides Wendyichthys_dicksoni Woodichthys_bearsdeni Gibbodon_cenensis Yelangichthys_macrocephalus Eomesodon_liassicus Brembodus_ridens Arduafrons_prominoris Mesturus_sp Mesturus_verrucosus Neoproscinetes_penalvai Brachydegma_caelatum NHMD_73588 Pteronisculus_cicatrosus Pteronisculus_changae Pteronisculus_magnus Pteronisculus_nielseni Pteronisculus_gunnari 
	;

END;


BEGIN CHARACTERS;
	TITLE  'Matrix modified from "Argyriou et al Brachydegma Sep 2020_sg.nex"';
	DIMENSIONS  NCHAR=303;
	FORMAT DATATYPE = STANDARD RESPECTCASE GAP = - MISSING = ? SYMBOLS = "  0 1 2 3 4 5 6 7 8 9";
	CHARSTATELABELS 
		1 '[G 1] Large dermal plates' /  absent present,
		2 '[G 2] Sensory lines' /  preserved_as_open_grooves pass_through_canals,
		3 Premaxilla_as_distinct_ossification_ /  present_ absent_,
		4 '[CH 1; G3] Premaxillae, contact at midline' /  present absent,
		5 Premaxilla_fused_at_midline /  absent present,
		6 '[G 4] Premaxilla' /  Reaches_or_extends_past_anterior_margin_of_orbit Confined_to_region_anterior_to_orbit,
		7 '[G 5] Premaxilla contributes to orbital margin' /  absent present,
		8 Teeth_on_premaxillae /  present absent,
		9 Mobile_premaxilla /  absent present,
		10 Olfactory_nerve_pierces_premaxilla /  absent present,
		11 Nasal_process_of_premaxilla /  absent short 'long, reaches skull roof',
		12 Sensory_canal_on_premaxilla /  present_ absent_,
		13 '[CH 3; G 6] Postrostrals (element[s] immediately anterior to frontals but not in contact with premaxillae)' /  present absent,
		14 '[CH 4; G 7] Single median dermal bone capping snout' /  absent present,
		15 Median_rostral /  'plate-like' 'tube-like',
		16 '[G 8] Pores for rostral organ' /  absent present,
		17 '[CH 8; G 10] Nasal bone as single consolidated ossification (i.e. bone(s) carrying supraorbital canal between premaxilla and anterior margin of frontals)' /  absent present,
		18 Contact_of_nasals_on_midline /  separated_by_dermal_bones contacting_or_separated_by_gap_unfilled_by_bone,
		19 Nasal_contributes_to_orbital_margin /  absent present,
		20 '[CH 57; G 11] Mesial margin of (anterior) nasal' /  not_notched notched,
		21 BOTH_NOSTRILS_ACCOMMODATED_WITHIN_SINGLE_DERMAL_OSSIFICATION /  absent present,
		22 '[CH 6; G12] Posterior nostril in complete communication with orbital fenestra' /  absent present,
		23 '[CH 7; G 13] Posterior nostril – contribution to margin by premaxillae' /  absent present,
		24 '[G 14] Tectals (sensu Cloutier & Ahlberg 1996, not counting the posterior tectal of Jarvik)' /  absent present,
		25 '[CH 75; G 36] Dermal intracranial joint' /  absent_ present_,
		26 '[CH 9; G 15] Pineal foramen' /  present absent,
		27 '[G 16] Pineal eminence' /  absent present,
		28 '[CH 10; G17] Shape of parietals (sarcopterygian postparietals): ' /  'rectangular, with long axis parallel to midline' quadrate,
		29 '[CH 11; G 18] Relative lengths of frontals and parietals (sarcopterygian parietals and postparietals)' /  frontal_shorter_than_parietal frontal_approximately_equal_to_parietal_ frontal_longer_than_parietal,
		30 Frontals_broad_posteriorly_and_tapering_anteriorly /  absent present,
		31 '[G 19] Anterior pit line' /  absent present,
		32 '[G 20] Otic canal extends through parietals' /  absent present,
		33 POSTERIOR_Junction_between_supraorbital_and_infraorbital_canals /  absent present,
		34 Anterior_branch_of_infraorbital_sensory_canal_ /  absent present,
		35 '[G 21] Tabular' /  present absent,
		36 '[G 22] Tabular pit line' /  absent_ present_,
		37 '[CH 64; G 28]  Number of bones carrying otic portion of lateral line canal between dermosphenotic and posterior edge of skull roof. ' /  'at least two (i.e. intertemporal and supratemporal)' 'one (i.e. dermopterotic)',
		38 '[CH 12; G 23] Intertemporal – relative length' /  shorter_than_supratemporal of_similar_length_to_supratemporal longer_than_supratemporal,
		39 '[CH 13; G 24] Intertemporal – contact with supratemporal anterior to that between frontal and parietal' /  absent present,
		40 '[G 27] Intertemporal contacts nasal' /  absent present,
		41 '[CH 69; G 29] Supratemporal – narrow anterolateral flange forming ventral margin of spiracular opening' /  absent present,
		42 Parietal_fused_to_dermopterotic /  absent present,
		43 Bone_carrying_otic_portion_of_lateral_line_canal_extends_past_posterior_margin_of_parietals /  absent present,
		44 '[CH 15; G 30] Number of paired extrascapulars' /  one_pair two_pairs three_or_more_pairs,
		45 '[G 31] Extrascapular reaches lateral edge of skull roof' /  absent_ present_,
		46 '[CH 71; G 32] Single median extrascapular' /  present absent,
		47 '[G 33] Extrascapulae contact each other at midline' /  absent present,
		48 '[CH 70; G 34] Medially-directed branch of sensory canal in extrascapulae' /  present absent,
		49 '[G 35] Extratemporal ' /  absent_ present_,
		50 '[CH 59; G 38] Antorbital bone' /  absent present,
		51 'Tube-like canal bearing anterior arm of antorbital:' /  absent present,
		52 '[CH 61; G 39] Infraorbitals' /  one two more_than_two,
		53 '[CH 16; G 40] Anterior expansion of lacrimal' /  absent present,
		54 '[CH 17; G 41] Notch in anterior margin of jugal' /  absent present,
		55 '[CH 18; G 42] Suborbitals (non-canal bearing ossifications separating jugal and maxilla)' /  absent one two three_or_more,
		56 '[G 43] Multiple rami of infraorbital canal in jugal' /  absent present,
		57 '[CH 54; G 25] Dermosphenotic with distinct posterior ramus' /  absent present,
		58 '[CH 14; G 26] Dermosphenotic – contact with frontals blocked by intertemporal or dermopterotic' /  absent present,
		59 Supraorbital /  absent one_or_two_ three_or_more,
		60 'Anterior-most infraorbital anterior to orbit (i.e. does not contribute to orbital margin)' /  absent present,
		61 Three_or_more_lachrymals /  absent present,
		62 Circumorbital_ring_ /  Supraorbitals_do_not_contact_infraorbitals_at_the_anterior_rim_of_the_orbit. 'Supraorbitals contact infraorbitals, closing the orbit.',
		63 '[CH 62; G 44] Jugal canal' /  absent present,
		64 '[CH 53; G 45] Dermohyal' /  absent present,
		65 '[G 46] Head of dermohyal projects above dorsal margin of operculum' /  absent present,
		66 '[G 47] Dermohyal' /  fused_to_hyomandibular separate_from_hyomandibular,
		67 '[G 49] Complete enclosure of spiracle by bones bearing otic and infraorbital canals' /  absent_ present_,
		68 '[G 57] Maxilla' /  absent present,
		69 '[G 58] Expanded dorsal lamina of maxilla' /  absent_ present_,
		70 '[G 60] Contribution by maxilla to posterior margin of cheek' /  absent present,
		71 '[G 61] Sensory canal/pit line associated with maxilla' /  absent present,
		72 Teeth_on_maxilla /  present absent,
		73 Mobile_maxilla_in_cheek /  absent present,
		74 'Peg-like anterior process of maxilla' /  absent present,
		75 Posterior_maxillary_notch_ /  absent present,
		76 Supramaxilla /  absent one two,
		77 '[CH 21; G 63] Course of mandibular canal' /  traces_ventral_margin_of_jaw_along_entire_length_ arches_dorsally_in_anterior_half_of_jaw_,
		78 '[G 64] Mandibular canal reaches anterior margin of mandible' /  present absent,
		79 '[CH 74; G 65] Mandibular canal' /  primarily_carried_by_infradentaries primarily_carried_by_dentary,
		80 '[G 66] Relative length of dentary' /  'long (constitutes most of the length of the lower jaw)' 'short (constitutes less than half of jaw length)',
		81 Teeth_on_dentary /  present absent,
		82 '[CH 22; G 67] Dentary with conspicuously reflexed distal tip' /  absent_ present_,
		83 '[CH 24; G 68] Enlarged series of parasymphysial teeth on dentary' /  absent_ present_,
		84 '[CH 73; G69] Facet for parasymphysial tooth-whorl on anterior dentary' /  present_ absent_,
		85 '[G 70] Teeth of outer dental arcade' /  several_rows_of_disorganized_teeth_ 'two rows, with large teeth lingually and small teeth labially ' single_row_of_teeth,
		86 Jaw_margins_overlain_by_lateral_lamina /  absent_ present_,
		87 '[CH 25; G 71] Acrodin caps on teeth' /  absent_ present_,
		88 Plicidentine /  absent present,
		89 '[CH 27; G 73] Ossification of mentomeckelian region:' /  present_ absent_,
		90 '[CH 23; G 76] Number of infradentaries' /  more_than_two_ 'two (angular and surangular) ' 'one (angular only)',
		91 '[G 74] Coronoids (sensu stricto, excluding parasymphysial tooth whorl or anterior coronoid)' /  present_ absent_,
		92 '[G 75] Number of coronoids' /  five__ four_or_more three two one,
		93 '[G 76] Posterior coronoid' /  morphologically_similar_to_anterior_coronoids expanded,
		94 '[G 78] Coronoid process of lower jaw' /  absent present,
		95 Coronoid_process_contributed_to_by /  prearticular_only surangular_only dentary_plus_postdentary_bones angular_only,
		96 Leptolepid_notch /  absent present,
		97 ACCESSORY_HYOID_ELEMENT_INVOLVEMENT_IN_JAW_JOINT /  absent present,
		98 '[G 79] Retroarticular process' /  present absent,
		99 Palatal_bite /  absent_ present_,
		100 '[G 82] Palatal articulation with basipterygoid process' /  articulation_not_obvious_ via_basipterygoid_fenestra 'via metapterygoid process/notch',
		101 Palatoquadrate_ossifications /  comineralized separate_ossification_centers,
		102 Palatoquadrate_symphysis_hf30_ /  absent present,
		103 Dorsal_margin_of_palate /  high_posterior_extension flat_dorsal_margin,
		104 Metapterygoid_posterior_to_quadrate /  absent present,
		105 '[G 81] Number of dermopalatines' /  multiple single,
		106 Prearticular /  present absent,
		107 Vomers /  paired single,
		108 Vomer_sutured_to_parasphenoid /  absent present,
		109 '[CH 19; G 50] Accessory operculum' /  _absent present_,
		110 OPERCULUM /  absent present,
		111 '[CH 67; G 51] Operculum - relative size ' /  at_least_twice_as_high_as_suboperculum subequal smaller_than_suboperculum,
		112 Subopercle /  present absent,
		113 '[CH 68; G 52] Anterodorsal process of suboperculum' /  absent_ present_,
		114 Anteroventral_process_of_suboperculum /  absent present,
		115 '[G 62] Number of cheek bones bearing pre-opercular canal posterior to jugal' /  one multiple series_of_small_ossicles,
		116 Preoperculum_orientation /  prounounced_dorsal_limb_ vertical_ pronounced_ventral_limb_,
		117 Junction_between_preopercular_and_more_anterior_cheek_bones /  'Infraorbitals (including jugal) or suborbitals suture with or abut preopercular' 'Infraorbitals (including jugals) and suborbitals broadly overlap preopercular' completely_obscure,
		118 Posterior_border_of_preopeculum_notched_ventrally_ /  absent present,
		119 Interopercle /  absent present,
		120 '[CH 72; G 53] Branchiostegal rays - dorsal-most in series' /  of_similar_depth_to_adjacent_branchiostegal_ray_ deeper_than_adjacent_branchiostegal_ray__,
		121 Lateral_gulars /  present absent,
		122 '[CH 20; G 54] Lateral gulars' /  extending_most_of_the_length_of_the_lower_jaw_ 'restricted to the anterior third of the lower jaw (no longer than the width of three branchiostegals) ',
		123 '[G 55] Median gular ' /  absent_ present_,
		124 '[G 56] Relative length of median gular' /  much_shorter_than_jaw_length more_than_half_of_law_length,
		125 '[G 83] Fenestra ventrolateralis' /  absent present,
		126 '[G 84] Palatal opening surrounded by premaxilla, maxilla, dermopalatine and vomer (choana)' /  absent present,
		127 '[G 85] Internasal cavity' /  absent_ present_,
		128 '[G 86] Interorbital septum' /  broad_ narrow_,
		129 Optic_foramen /  dorsally_positioned_ 'ventrally positioned (i.e. abuts parasphenoid) ',
		130 '[G 87] Pronounced median anterior crista on dorsal surface of braincase' /  absent_ present_,
		131 '[G 88] Expanded anterior dorsal fontanelle ' /  absent present,
		132 '[G 89] Endoskeletal intracranial joint' /  absent_ present_,
		133 '[G 90] Eye stalk or unfinished area for similar structure' /  absent_ present_,
		134 '[G 91] Roof of posterior myodome perforated by palatine branch of facial nerve (VII)' /  absent_ present_,
		135 '[G 92] Foramen for abducens nerve (VI) dorsally positioned (level with optic foramen (II))' /  absent_ present_,
		136 '[G 93] Anterodorsal myodome' /  paired_ single_ absent,
		137 '[G 94] Posterior myodome' /  absent paired median,
		138 '[G 96] Basicranial fenestra' /  absent_ present_,
		139 '[G 98] Endoskeletal spiracular canal' /  open partial_closure_of_spiracular_bar complete_enclosure_in_canal_,
		140 '[G 100] Basipterygoid process' /  present_ absent_,
		141 '[G 101] Basipterygoid process with vertically oriented component' /  absent present,
		142 '[G 103] Dermal component to basipterygoid process' /  absent_ present_,
		143 Hyoid_facet /  directed_posteroventrally horizontal,
		144 '[G 104] Fossa bridgei' /  absent_ present_,
		145 '[G 105] Posttemporal fossae' /  absent_ present_,
		146 '[G 106] Vestibular fontanelle' /  absent_ present_,
		147 '[G 107] Ventral cranial fissure and vestibular fontanelle' /  separated_by_bridge_of_bone_ conflUent_,
		148 '[G 108] Accessory fenestration in otic capsule ' /  absent_ present_,
		149 '[G 109] Otoccipital fissure' /  absent_ present_,
		150 '[G 110] Median projection overhanging posterior part of posterior dorsal fontanelle' /  absent_ present_,
		151 '[G 111] Median projection overhanging anterior part of posterior dorsal fontanelle' /  absent_ present_,
		152 CRANIOSPINAL_PROCESSES /  absent present,
		153 '[G 112] Dorsal aorta' /  open_in_groove_ canal_notched_posteriorly completely_enclosed_in_canal_,
		154 '[G 113] Dorsal aorta pierced by canal/s for exit of eff.a.2' /  absent present,
		155 '[G 114] Dorsal aorta pierced by canal/s for exit of eff.a.1' /  absent present,
		156 '[G 115] Bifurcation of dorsal aorta' /  posterior_to_occiput_ anterior_to_occiput_,
		157 '[G 116] Birfurcation of dorsal aorta into lateral dorsal aortae' /  open_ enclosed_in_canal_ below_parasphenoid,
		158 Braincase_ossifications_differentiated /  absent present,
		159 Basisphenoid /  present absent_or_very_reduced,
		160 'Opisthotic-pterotic relationship' /  opisthotic_larger_than_subotic opisthotic_and_pterotic_equal_in_size,
		161 Epioccipital /  present absent,
		162 Forward_extension_of_the_exoccipital_around_the_vagus_nerve /  absent present,
		163 Sphenotic_with_small_dermal_component /  absent present,
		164 Pterotic /  present absent,
		165 Opisthotic_bone /  present absent,
		166 Intercalar /  present absent,
		167 OCCIPITAL_REGION_OSSIFICATION_PATTERN /  basioccipital_and_exoccipitals_as_separate_ossifications 'co-mineralized',
		168 Supraoccipital_bone__ /  absent present,
		169 Membranous_outgrowth_of_intercalar /  absent_ present_,
		170 'Post-temporal fossa' /  absent present,
		171 'Sub-temporal fossa' /  absent present,
		172 Dilatator_fossa /  absent present,
		173 '[G 117] Parasphenoid' /  absent present,
		174 '[G 118] Parasphenoid' /  'terminates at/anterior to ventral otic fissure ' extends_across_ventral_otic_fissure_ extends_to_basioccipital_,
		175 '[CH 28; G 120] Ascending process of the parasphenoid' /  absent_ present,
		176 '[CH 29; G 121] Parasphenoid with multifid anterior margin' /  absent_ present_,
		177 '[G 124] Buccohypophyseal canal pierces parasphenoid' /  present_ absent_,
		178 Parasphenoid_teeth /  small large absent,
		179 PARASPHENOID_PIERCED_BY_ASCENDING_COMMON_CAROTIDS /  absent present,
		180 Parasphenoid_pierced_by_internal_carotid_artery /  absent present,
		181 Parasphenoid_pierced_by_efferent_pseudobranchial_artery /  absent present,
		182 Aortic_notch_in_parasphenoid /  absent_ present_,
		183 Parabasal_canal_ /  present absent,
		184 '[G 125] Anterolaterally divergent olfactory tracts' /  absent_ present_,
		185 '[G 126] Elongate olfactory tract(s)' /  absent_ present_,
		186 '[G 127] Olfactory nerves carried in a single tract' /  present_ absent_,
		187 ARRANGEMENT_OF_OLFACTORY_NERVE_IN_ORBITAL_REGION /  enclosed_in_olfactory_canal 'traversing the orbit lateral to the interorbital septum, at times leaving a groove on the latter',
		188 '[G 128] Hypophyseal chamber' /  projects_posteroventrally_ projects_ventrally_or_anteroventrally,
		189 '[G 129] Optic lobes' /  narrower_than_cerebellum_ same_width_or_wider_than_cerebellum_,
		190 Optic_lobes /  smaller_than_telencephalon larger_than_telencephalon,
		191 '[G 130] Optic tectum divided into bilateral halves' /  absent_ present_,
		192 '[G 131] Cerebellar corpus' /  absent_ present_,
		193 Cerebellar_corpus /  divided_bilaterally undivided_,
		194 Position_of_cerebellar_corpus /  _enters_fourth_ventricle arches_above_fourth_ventricle,
		195 Cerebellar_corpus_with_median_anteriorly_projecting_portion /  absent present,
		196 '[G 132] Horizontal semicircular canal' /  joins_vestibular_region_dorsal_to_ampulla_for_the_posterior_semicircular_canal_ joins_vestibular_region_level_with_ampulla_for_the_posterior_semicircular_canal_,
		197 '[G 133] Junction between ampulla of posterior semicircular canal and cranial cavity' /  separated_by_short_length_of_canal__ confluent_,
		198 '[G 134] Crus commune of anterior and posterior semicircular canal' /  dorsal_to_endocranial_roof_ ventral_to_endocranial_roof_,
		199 '[G 135] Lateral cranial canal' /  absent present_,
		200 Lateral_cranial_canal_connects_to_cranial_cavity_anteriorly /  absent present,
		201 LATERAL_CRANIAL_CANAL_CONNECTS_TO_LATERAL_WALL_OF_BRAINCASE /  absent present,
		202 INTRAMURAL_DIVERTICULA_OPENING_IN_FOSSA_BRIDGEI /  absent present,
		203 '[CH 30 in part; G 147] Enameloid on dermal bones and scales' /  absent_ present_,
		204 '[G 148] Extensive pore-canal network' /  absent present_,
		205 '[CH 30 in part; G 149] Enamel' /  'single-layered ' 'multi-layered ',
		206 '[CH 30 in part; G 150] Enamel layers' /  applied_directly_to_one_another_ separated_by_layers_of_dentine__,
		207 Scales_on_body /  present absent,
		208 '[G 151] Scales' /  micromeric macromeric,
		209 '[CH 32 in part; G 152] Scales with ‘peg and socket articulation''' /  absent_ present_,
		210 '[CH 32 in part; G 153] Peg on rhomboid scale' /  narrow_ broad_,
		211 '[CH 33; G 154] Anterodorsal process on scale' /  absent_ present_,
		212 '[CH 35; G 155] Scales with well developed pores on surface' /  absent_ present_,
		213 Small_scales_below_dorsal_fin /  absent present,
		214 '[G 158] Lepidotrichia' /  absent_ present_,
		215 '[CH 37; G 159] Fringing fulcra' /  absent_ present_,
		216 '[G 139] Double headed hyomandibular' /  absent present,
		217 '[G 140] Perforate hyomandibula' /  absent_ present_,
		218 '[G 141] Opercular process' /  absent present,
		219 '[G 136] Ceratohyal' /  single_ossification two_ossifications_,
		220 '[G 137] Anterior ossification of ceratohyal' /  no_medial_constriction 'medial constriction (hourglass-shaped)',
		221 '[G 138] Anterior ceratohyal' /  no_groove_ groove_for_afferent_hyoidean_artery_,
		222 'OSSIFIED ACCESSORY HYOID ELEMENTS ABS/PRES' /  absent present,
		223 'IF PRESENT, NUMBER OF ACCESSORY HYOID ELEMENTSONE OR TWO ' /  one two,
		224 Symplectic_shape_c68 /  'tube/splint like' hatchet 'l-shaped',
		225 'SYMPLECTIC, POSITION' /  posterior_to_the_posterior_margin_of_quadrate medial_to_the_posterior_margin_of_quadrate,
		226 '[G 145] Hypohyal' /  absent present,
		227 '[G 143] Basihyal' /  absent present,
		228 '[G 146] Gill arches' /  largely_restricted_to_area_under_braincase extend_far_posterior_to_braincase,
		229 Number_of_ceratobranchials /  five_ four_,
		230 Number_of_hypobranchials /  three four,
		231 Uncinate_processes_on_epibranchials /  absent present,
		232 '[G 142] Endoskeletal urohyal' /  absent present,
		233 Urohyal_formed_as_a_tendon_bone_of_the_sternohyoideus_muscle /  absent present,
		234 '[CH 39; G 162] Presupracleithrum' /  absent_ present_,
		235 Presupracleithrum_ /  single multiple,
		236 '[G 160] Dorsal margin of cleithrum' /  pointed broad_and_rounded,
		237 Medial_wing_on_cleithrum /  absent present,
		238 '[G 161] Anocleithrum' /  bone_developed_as_postcleithrum bone_developed_as_anocleithrum_sensu_stricto bone_absent,
		239 Clavicle /  present_as_a_broad_plate much_reduced_or_absent_,
		240 Serrated_organ /  absent present,
		241 Interclavicle /  present absent,
		242 '[CH 40; G 167] Triradiate scapulocoracoid' /  absent_ present_,
		243 '[G 163] Perforate propterygium' /  absent_ present_,
		244 '[CH 41; G 164] Anterior rays embrace propterygium' /  absent_ present_ fused,
		245 Propterygium_fused_to_first_ray /  absent present,
		246 '[CH 43; G 168] Pectoral fin endoskeleton' /  'extends far beyond body wall (fins lobate)  ' 'barely extends beyond body wall (fins not lobate)  ',
		247 '[G 166] Pectoral fin radials' /  unjointed_ jointed_,
		248 '[G 169] Fin articulation' /  monobasal polybasal,
		249 '[CH 44; G 171] Pectoral fin-ray segmentation' /  roughly_even_segmentation_to_fin_base_ proximal_segments_elongate_with_terminal_segmentation_ no_significant_segmentation_on_pectoral_fin_ terminal_segments_elongate_with_proximal_segmentation,
		250 '[G 170] Pectoral fin' /  '[leaf-like]' '(not leaf-like)',
		251 '[G 172] Paired fin spines' /  absent present,
		252 '[CH 38; G 173] Pelvic fins' /  absent_ present_,
		253 '[CH 45; G 174] Pelvic fin insertion' /  'shorter than fin depth (short based) ' 'longer than fin depth (long based) ',
		254 '[G 177] Basal scutes on fins' /  absent present,
		255 '[CH 48; G 178] Dorsal scutes anterior to dorsal fin' /  absent_ 'few limited to region immediately anterior to fin (basal fulcra only)' 'many, extending to posterior of skull roof (complete set of dorsal ridge scales)',
		256 '[CH 49; G 179] Ventral scutes between hypochordal lobe of caudal fin and anal fin' /  absent_ present_,
		257 '[CH 50; G 180] Ventral scutes anterior to anal fin' /  absent_ present_,
		258 '[CH 52; G 181] Dorsal fin(s)' /  two_ one_,
		259 '[CH 51; G 182] Relative positions of anal and (second) dorsal fin' /  anal_shifted_anteriorly_relative_to_dorsal_ fins_opposite_one_another_ anal_shifted_posteriorly_relative_to_dorsal,
		260 'Median fins (except caudal fin)' /  rays_more_numerous_than_radials_ rays_and_radials_equal,
		261 Proximal_and_middle_radials_of_dorsal_fin /  proximal_and_middle_radials_of_similar_size proximal_radials_substantially_enlarged,
		262 Posteriormost_proximal_radial_of_dorsal_fin /  enlarged_plate smaller_than_more_anterior_radials,
		263 '[CH 46; G 175] Epichordal lobe of caudal fin' /  present_ absent_,
		264 '[CH 47; G 176] Fulcra along dorsal ridge of caudal fin' /  absent_ present_,
		265 Caudal_fin_geometry /  long_chordal_lobe short_chordal_lobe,
		266 Posterior_margin_of_caudal_fin /  forked unforked,
		267 EPINEURAL_PROCESSES /  absent present,
		268 'Diplospondyly in mid-caudal region' /  absent present,
		269 Median_neural_spines_in_caudal_region /  absent present,
		270 Uroneural /  absent present,
		271 Division_of_hypurals_into_dorsal_and_ventral_groups /  absent present,
		272 Number_of_caudal_lepidotrichs_borne_per_hypural /  multiple single,
		273 Opistocoelous_vertebrae /  _absent present_,
		274 Ossified_ribs /  present absent,
		275 Antorbital_deeper_than_anteriormost_infraorbital /  absent present,
		276 Infraorbitals_fragmented_ /  absent present,
		277 Suborbitals_fragmented /  absent present,
		278 Suborbitals_extend_ventral_to_orbit_ /  absent present,
		279 Coronoids_contribute_to_lateral_dentition_field /  absent present,
		280 Jaw_articulation_ventral_or_anterior_to_orbit /  absent present,
		281 Parasphenoid_wings_around_basiocciput /  absent present,
		282 Dorsal_extension_of_parasphenoid_between_orbits_ /  absent present,
		283 Ligament_pit_on_posterior_face_of_braincase /  absent present,
		284 '''Hem-like'' median fins' /  absent present,
		285 Anal_fin_insertion /  short long,
		286 Denticulate_ridge_scales /  absent present,
		287 Ossified_centra /  absent present,
		288 Supratemporal_lateral_line_commissure_passes_through_parietal /  abs pres,
		289 Enlarged_vomerine_teeth /  abs pre,
		290 Enlarged_prearticular_teeth /  abs pres,
		291 Prearticular_symphysis /  abs pres,
		292 Dermal_supraoccipital /  abs pres,
		293 Preoperculum_taller_than_operculum /  shorter taller,
		294 Preoperculum_wider_than_operculum /  narrower wider,
		295 Branchiostegals_contact_next_in_sequence /  abs pres,
		296 Dorsal_lateral_line /  abs pres,
		297 Parasphenoid_inflected_ventrally_below_orbit /  abs pres,
		298 Sagittal_flange_on_neural_spine /  abs pres,
		299 Sagittal_flange_on_haemal_sine /  abs pres,
		300 Pleural_ribs_alate /  absent present,
		301 Lacrimal_contributing_to_oral_margin /  absent present,
		302 Teeth_on_lacrimal /  absent present,
		303 Dermosphenotic_position /  extending_well_below_dermopterotic located_at_same_horizontal_level_than_dermopterotic ; 
	MATRIX
	Acanthodes_bronni             0????????????????-??????0?-?????01?-??????-?????-?????????????1??????????????????????-???????0-?0?021000?-?-?-----?--??0??0-????0000?????00000000???10000--0-?0??????????0000---------1????????????01?0-??00--000---?0000010?10--??1010??-??-????0000101?1110000002????00??????-?---?-?0--?00-?--------00??-???
	Acipenser_brevirostrum        111---------00?0000-01-?010000-0101-1----00110-00??1-001001--000--00-----------010---0-?0-1--0-01100111-111000-0002---011-0-0-0000000-?20021--0100--0--10--1{1 2}10?100?01?0-000121112000111010000011011010--1----010-0001000001010??1010{0 1}0000-000000101?10101110121112000110000100001?0---0000000000--0--10000-000
	Aesopichthys_erinaceus        1101-0000001110010110100010120?0001-00110-1111100101--1101000-011101000000000010000120???2???120?1??????????0110010100010?11???????0??????2?????????1????????00----------?????????????????????????????????????011010011011???????????????100?00??????0??01010021012???0100????????10?000???011?0???01010????00-
	Amia_calva                    1101-100012010001100000001012010101-1----00011100110000110000-10--010000111100100001201?110101201100101000010100100210101-11000?000001112021--1100-00--00--1211-00111000110112111000010001-110111110010--000--010---?100111001010101011000-00011100101011101000001211110110110010010--000000?010100010100000001
	Amphicentrum_granulosum_      11000101000?110?10110?0001-020?0001-1----01?1??000-1100000000-00--11100000000?1010---1-??20?-0-0??120010????0120010100000???0001???0????20?0010??0-00???21011?????0??????1?011100?0001?????????????????????????????????010???????????????0-0??0?????????????????????????0?????????-0--00-000?0?001?010100???000
	Atractosteus_spatula          11000100-12010101100000001012010111-1----01211100112--3-10111101110100100100001000011001110201200100101100010110100210011-0-000100000-022020011100-00--00--1211-01101100-10112111000010??11???????????????101?0110000110111101020101011000-0101110010001110100000?111111110010011000100100000-10000000100000000
	Australosomus_kochi           111?????-??-??0??0??????01002010001-1----01011000??0-001--0---010111100000001010000120??011--0-001020000?0000110000100000?100001001001022021--?1011010012001001----------0?110101200000011-1110????1110--0????01101001101100110??101001000-00000110?000111010000010010??1010100001?0---0-0001-1000001010000-00-
	Beagiascus_pulcherrimus       11000000000111001011010001012010001-00110-0210-?0101003111000-?10??111?00000???00001101???????-0?1??????????1110000000010110??????????????????????????????????????????????????????????????????????????????????011?10011??????????????????110?00??????1??21010011?11????100????????1010?0???000?0???010100???000
	Beishanichthys_brevicaudalis  1100?01000?0110010110100010120?0??1-1----00111?000-100210020?00101?1100000000?100001201??2???1?0????????????01100000000000??????????????????????????????????????????????????1?????????????????????????????????011010?10??????????????????0-0?0??????????11010000012???1011????????-000?0???01-?0???010100???000
	Birgeria_groenlandica         111--1-0----110010110000010?2010001-1----01001100102003010200001?1011100000000100001101?01???110?100100000--01?0000000000110000100?0010?202?--010???1--1???1210-?-0?????-0001210120001?????????????????0??????010?00011000?0?10??101?0?000-00200100??001110100000120001100001000-1?0100010000-0?-0001010000-000
	Birgeria_stensioei            111--1-0----110010110000010120?0001-1----01?????0102003000200001110111000010???000011010?10??110?1001000?0--010000000000?????????????????????????????????????1??????????????1210100001????????????????????????1------11001?0110???????0000-0?20010000001?101010001200011000??000-1?010?010?00-0??0?01??0000-000
	Bobasatrania_groenlandica     111----1----0000101101-0010120?00{0 1}1-1----0?0011000-2100110000-00--0100011000001010---0-??2???0-?0?0??0?0????01-1--11000-?-0-???10???0??????0??????????????????1?????????????121??2????????????????????????????011???0110?1010????1???????0-0001?????????0?01001011210?11000??????1-0--00???1100000?000?10??-000
	Boreosomus_piveteaui          11001011000011001111010?01002010011-1----01100-000-2003100000-11010111100000?010000110?0010300-0010200010100011000010001011000?1010001?12020010101001{0 1}{0 1}12111000----------000101010-1000010-?1111100??11010????01101001100111?10??1?1??00?0-?020011???0??1?01000101200?010010000?-1-00000-0000-000--010100??-00-
	Brazilichthys_macrognathus    110001-00000110?101101000????????11-?????????????0-2?1000??00-?????11?000000001000001010010??0-0?10??000?0???1?????????0001000??????????????????????????????????????????????1?1012000?????????????????????1????????????011???????1?1??0???????00??1?1??1??????????????????????????-0--00-0???????00???1?0??????
	Caturus_furcatus              1100010001?110101100000001012000110-1----00011100112102?01200110--0100001111???0000120??010301201102101??0010100100210?11-1100010??0?????021--1??0-00??0?????10?0001100011??12111000000???????????????11-0??????????0110111001010??1??1??1000??1?0????????01000001211111100?1000-000000000?00000100000??0000001
	Cheirolepis_canadensis        110?01-000000000000?0?010{0 1}0010?0001-000-0-10011000-1010010000-010?111110000010100001100??10?00-0????????????11000000000000???0??????????????????????????????????????????????1?00??????????????????????????????000-0?011??????????????????0-0?00????0001101011010110???0101????????-0--00???010?0?0?00010????00-
	Cheirolepis_schultzei         11000000000?000000000?01010020?00?1-1---0010111?00-1010010000-010?1111?0000010100001?00????????0?????????0??1100000000000?1??0??????????????????????????????????????????????1????0????????????????????????????000-0??11????????????????????0?00??????0??010??0????????????????????-0--?0???????????01010????000
	Cheirolepis_trailli           1100001000000000000100010{0 1}001010001-00000-10010000-1010010000-01101111?0000010100001100??10100-0?10100000000110000000000001000?0??00?????010000001001?010--1-00----------000100000000001110???????????????111000101?011000???????????????10000000001000101011011110???0100???00??1-0--00-0000000?00010100??-00-
	Chondrosteus_acipenseroides   111---------1?????????-?010020????1-1----0?2?0-?0??0?0??00?????0--00-------??01010---0-?0?1--??0??0??1??10??012000??-??0?????????????????????0??????????????????????????????1211?2?0011???????????????????----1-----?10000010????????????0-0?0?????????12101?00001????110???0000?1?0??-0???00-?????0??1?000-000
	Cladodoides_wildungensis      0?-?-??-----??-??--?-???-?-??-?????-?????--?????-?-???-???----?????-???-----????-?1??-00?????0---?0?00?-?---?-?-?---?--?-??-??0000001?0?00010-0000-01000---0-00----------0000---------1101000000---0100---????00???????0?????0---????????--?-?---????????????????????????????????------0--0-?----------0----???
	Coccocephalichthys_wildi      11????0?????11?01010010?01002010011-1----00110-00112000?0010?01?0?0111?00?00??100001101??1???1100101000000?0?1101?00000?????00?10??000?02020000?01001??12101100----------0?010101000000???????????????????????01???????0?1???10?????????????0?????????????????????????????????????00--00-0?????0?00010??0??????
	Cosmoptychius_striatus        110??1000?00?1001011010001012??0001-1----?0?1??00101002100000-010?01110000001?10000?10???2???????1020???????110000000001?110???1?????????0?001???1101??111111?0??00?????????101?1?00000???????????????1???????011010011??????????????????0-0?000????011110011010012???1100?????0??100000-??000?0???01010????000
	Cyranorhis_bergeraci          111---------1100101101-001012010001-00100-1111100101002110000-010101110000000?10000120?????????0???20???????0100000000000110??????????????????????????????????????????????????????????????????????????????????011?100110?1???????????????100??0?1????1??11010010112???0100????????1000?0???000?0?00010100????0-
	Dapedium_sp_LIAS              11001100001?1100110?0000010---0-??-------1-111100102--3-0-1000?0--0100001101??11000120??0?0?0120??1??0??00??0110100210101-1100010?0001?120?0011100-00??02001?0-----------111121112-10000010111?????110110?????011??0?110?1?????0?????????100?01?1???????21010?????????1?11??1?0??0011111111110?0????0010????---
	Scopulipiscis_saxciput        11??????????????????????010120?00???1----0-?????????????????????????????????????????????????????????????????????????????????0??10?10010?2020011100-00--?2001??0??????????111121?02-1000????111?????110100???????????????????????????????????????????????????????????????????????????????111????0???????00??????
	Dapedium_pholidotum           110011000011110011010000010---?-01-------1-21110-102--3-0-200010--0100001?011101000120???0030120????00?010010110000220101-11??????????????????1??????????????0-----------?????????????????????????????????????011??0111011?10?1??1???????10??01?1????1??110100201121??1111????????011111???110?0????0010????---
	Dapedium_stollorum            11010100001110001101000001012010011-1----00211100102--3-001000?10101000011011101000120???0030??0??1?1???????0110100210101-11?????????????????????????????????0????????????????????????????????????????????????011??0111??????????????????1{0 1}??01?1????10111010020112???1111????????011111???110??????0010????---
	Dapedium_punctatum            110?010000??100011000000010---?-??-------1-2111?-102--3-0-1101?0--?100?0110????1000120???0????????1?????????01000012101????????????????????????????????0????????????????????1?????????????????????????????????011??0111011???????????????100?01??????1??110??020?12???1111????????011111???110??????00100???---
	Dapedium_caelatum             1100010000?111001100000?01012010?11-1----01?????0102--3-??100000--010001110???01000120???0020120??0?1?????0?0100000210101-11???????00??????0011??0-?0??02??110-----------???1211?2????????????????????????????011??011?0???????????????????0?01?1????1???1011020?12???1111????????01?11111?110??0???00100???---
	Dapedium_noricum              110?01000???100011?00?0?0100201???1-1----002111?0??2--3-00100??0--?100?11?00????000120????0?????????????????0110100220101-11????????????????????????????????????????????????1????2????????????????????????????011??0?11???????????????????0-?????????{0 2}??110100???12???1111????????000111???11000???00010????---
	Hemicalypterus_weiri          11000100001100-0110?0??001002000011-1----002111001121-3-00100100--?1000111000???000120???1???1????0?1???????010010?210101-11??????????????????????????????????????????????????????????????????????????????????011??0-11??????????????????{0 1}0??01?1????1??1101002-1121?11111?????1??100101???11100???00010????000
	Paradapedium_egertoni         110?010000??1000110000000100201???1-1----002111?0??2--3-???00??0--?1?0?01???0?10000120????0???2???001?10?0??010010021010??????????????????????????????????????????????????????????????????????????????????????011??0111011???????????????0-0?01??????1??110100211121?11111???????????????????1???0?00010????--?
	Sargodon_tomicus              1?0?0100001?1?0?1100?01?010120????1-1----0001?1?0??2--3-0??001?0--?1?0?11??????1?00120???1????????1???????11011010022?10?????????????????????????????????????????????????????????????????????????????????????????????1???????????????????{0 1}0??21??????1??110100211121111111????0????00101???1110?1??00010?00?--?
	Dandya_ovalis                 110??1?00???100?11?00?0?010120101?1-1----{0 1}02111?0??2?-3-???00??0--?100?11?0?????000120????????????0?????????010010122010???????????????????0?1??????????????????????????????1?1???????????????????????????????011??0?11??????????????????0-0?2??????????11?100???12???1111?????????001?1???11?001??00010?00????
	Heterostrophus_phillipsi      110?010000?1110011?10?00010010????101----0???????102--3-01100??0--?100001100??10000120???1??????????????????0100100211101-11????????????????????????????????????????????????1?????????????????????????????????011010?11??????????????????100?01?1????1??11?100?011????1111???????0??1101???1100????000100???--?
	Tetragonolepis_oldhami        110??1000???1?00?0??????010?20????1-1----0001???0?????3-???????0--?1?0?01???0?10000120??????????????????????0110010210101-11??????????????????????????????????????????????????????????????????????????????????011????11????????????????????0??1?1????1??1101002?1121?11111???????0???1?1???111?01??00010???????
	Dialipina_salguerioensis      100???1000??01?0?0??????01012100001-1----00?????????103?00????????110??00??????00?01000??0??????????????????????????0???000-??????????????????????????????????????????????????????????????????????????????1???011010?10????????????????????0?0????????????0110?0001???0??1????????????00???000?????0??10????---
	Dicksonosteus_arcticus        100??1-000000100????????00-00000?0??1?????0010-1???10000??0??-?0???0--------?????????0????????????00000???--00????0?????????000000001?????0???0000-00??00--0?00??????????0001000000000?1010???00??????0????????????????????????????????????????????????????????????????????????????0--?0--0????0-??0????0???--?
	Diplocercides_kayseri         110?01-000??00?10??0000111000010001-1----0?010-0???100?0???00?10--00?-------00010???00?1010?113010021000?0?00100?010000-000-0?000?010???0100000010-10??00--1-00----------0001000000000?1110?0?????????????????01---0???1?????110-???????????0????????????10??0??????????1?????????????00-0000???000000?00???00?
	Dipteronotus_ornatus          110??10000??1100100100??010120?000??1----?0??????1?0-00???1--1?????10000000000100001201???1????0??0?1????????1100001000?1?????????????????????1???????????????????0?????????111012000?????????????????????????01?0?0?11????????????????????0?2????????????01002?112????110??100???00---0-0?0100?0??01010000?--?
	Discoserra_pectinodon         110000100001010010110100010120?0000-1----002110000-2003-01100000--010000?0000010000120???20??0-0010?1??0?0??01-1--120000010-????0??00??02?2?--11?0-0?????????0-----------1??121?0??0??????????????????????????0110?011101?00010??????????110??1?1???00???10100201120001111??10???1-01101???01100?0010010000-000
	Donnrosenia_schaefferi        110?001?000011001011010000-01010001-00100-10111?00-???0?10000-0?0?1111?000001110000110???1???0-0????????????11001000?0000010?0????????????????????????????????????????????????????????????????????????????????011010011????????????????????0?00??????1??11011010111????1??????????-?--00???000?0???01010??????-
	Dorsetichthys_bechei_         110?0100100?11001010000001012110?11-1----00011000102101110100000--?1010011020010000120???1????21??0?10???11?01100002101??-11???1??1000002020011101001??00????101000000{0 1}11111121000-1111?10-?11?????11111-0????0110?0?1?011????????????1??0-0?0??????10??21010000012???1110?111100000000000000-100--000100000000
	Ebenaqua_ritchei              111----1----1100101001-0010120?0001-1----001011000-2?00000000-00--0100011?000??010---0-??20?00-0?10?????????01000011000-0-0??????????????????1??????????????????????????????1?1???????????????????????????????011??001????01?????????????11??21?????????21010021111??11101????????-0--01???01000???000?0????000
	Elops_hawaiensis              1101-100102111001000000001012100111-1----00011100102100000100010--1101001102001000010010021--1200?02101011110100010210101-1100010??00???2000--1100-00--00--1211?010010011111120010-101????????????????0--000--000--00110111111101?11000?10-00010100111010101000001211111101010000000---000000-100--0101?0000000
	Entelognathus_primordialis    100??1-100010100??0?????0100?00?-0????????0010-????10000??0??-10???1001100000?001001?0?-?01?-0-0??0000???---0?????0?-0??00????0?0??0?????0????0?00-?0??00??0-00----------000?----?---?0???????????????????????0?????0??0???????--??0-????0-??0000????????11??????????????1?????????0---?-??????---??????????00?
	Erpetoichthys_calabaricus     1101-110000010000000010001002011101--------2111000-1003000000-010101101000000010000120?00203010001001010001001-1--00000-000-000010000-000011--0001-00--02001110???0?0010000012101000010101001011000011?0?-10??01101001000100010--101110010-000001????0?1?100-????1-?111?110????1?0-0000000?0000000001000???0000
	Eusthenopteron_foordi         110001-0000000000000001110-01010100100000-0010-000-1000001100110--0100000000000000011000000200-00102000-100001100010000-0010110000010???0100100011001--00--1-00----------00010000000001111010000---1100---00--010--001011010110--100010100-1010001---000010101000020000011001000-0-0--00-0000-00100010?0000000-
	Evenkia_eunoptera             110?0010000000?00?010?0001012001111--------?1??000?2002?002000011?0110000000????0???20???1?????0????????????11100100000000????????????????????????????????????????????????????????????????????????????????????011??0110??????????????????100?????????0??0101000?112???100?????????-010?0???01-?0????1010????000
	Fouldenia_ischiptera          11000101000?110010100?00010110????1-1----010111?00-1000?00000-?101?111?000000?10100121???20??0-0??1?0???????0110010000000110????????????????????????????????????????????????1?????????0???????????????????????0110100110?????????????????0-0?00?0??1000101010021111???0100????????-0--00???010?????01010????000
	Fukangichthys_longidorsalis   11000?1000??11001?1101?0010120?0101-1----011011?0????0310020?0?1010111000000???00001201001???100?100101000100110000000000010???01???0????????1??????????????????????????????1?1000000?0???????????????????????011??0011001001??????11?1????000??????????11010000012????011?????????00000?0?01-?0100010100???000
	Glyptolepis_groenlandica      110?-1-0000???-0?00??0111110001110010---0-0010-010-1100000000-10--01000?0??0?0000?10?0?1???2?0-00002000-1000011010100000001010100?010???010?1?001????????????00----------0001000000000011101000?---???????????01--00?101???0?10--10011?100-10?00?0???010?00??000?0????00010??????1-0--00-0000-00???000100??-00-
	Gogonasus_andrewsae           1100-1-0000?00-00000001110-00010100100000-0010-0?0-1000001?00?10--01000000000000011020?0000200-0?102000-000001100010000?00?011000?010???01001000?1001--00--1-00----------0001000100000?111010000---1?00---110-010-00?1?110?01????1100??1?0-???0001---000?1???1????????0??1????????-0--00-00000??100000?00???00-
	Gogosardina_coatesi           11000010000001001011011000-010100?1-0010?-1?????00-??000?0?0??0100?11110000001100001101?020?00-00101000000?001101000000000?0?0?????????????000?????????11001??0?????????????100??00000????????????????????????0110100110100??10??1?10?????-000001????1??11010021012????10?1?000??0-0--00-0?0?000?00010100?????-
	Guiyu_oneiros                 1?01-000000100000000000110-00000000001000-0010-110-1000000100100--01111000000?000110100?0?0000-0?1???0????0001001000000000100010???1???????0?00???????????????0???-?????-???1000000000????????????????????1?1101111001?????????--????????100??0???????????110011101???????????????-0--00-??0?0?0?00000100???00-
	Hiodon_alosoides              11000100100111001100000001012010011-1----1-0110000-2000000000-00---10010110001100001200?021--1200100101011100100000210001-0-000??000011?2010-01100-00--00----11-0100100111?112101100010????????????1110---00--010--00101111101101111001010-000101001110111011000011111101010111000-0---000001-100--011?0000000-
	Howqualepis_rostridens        1101-010000011001011011000-12010001-00100-10111?00-1100010000-01001111?000000010011110??110?00-00101000000000100100000000010?0????????????1000???10????10--1-?0???-?????????10110000000???????????????????????0110100110?0???10??????????10000000??1000111011011111???0100??000???-0--00-??000?0?00010110???00-
	Hulettia_americana            110001000011111010000000010?2010001-1----00011100102-03100200000--01000011000?100?0120???10??1200?0210100?1?0100100210101-0-???????0?????020?11???-01???0--121000?0000100?0?111101-1100???????????????????1??-0110?0011011110100010???1?00-0?0101????????1010010112111101011000000100001-00000100??000100000--0
	Ichthyokentema_purbeckensis   1101-100100111001101000001002010001-1----00011000102101101200010--01000011010010000120??0204-121010210001?110100100210101-1100010??0010?20?0011100-00--00--1211101?000011111121001-100????????????????????100-011010?110??1011001????????0-0001011???101?101001011?1??11100?11??00100001000000100??010100000000
	Kalops_monophyrum             1100010000?111001011010001-12110001-1----00010-00102003000200001010111000000?010000110???20?0??0????????????0100010000010110?????????????????1???????????????0?----------?????1???????????????????????????????0110100110?0???????????????100?00?1????0??01010021112???0100????????101000???000?0???01010????000
	Kansasiella_eatoni_           11??????????????????????000020100?1-1----0??????0????????0?????????????????????????????????????0???1???????0???????000??????0??1011001112?2001010100110111111100-?000010?00010101000000010-0111110?1111001?????????????0???11??????????????0????????????????????????????????????????????-00????????????????????
	Kentuckia_deani               110?001000000100101101000{0 1}002010001-?--?-?10111000-??000?000?-?10??111?00?000?1000011???02?????0??01000000??110000000000011?00?100100?1110200101011010011101000----------000101?00000?0010-011?100?11110?01????????0???011????????????????????????????????0???????????????????????-?--00-00????0?000101?0???000
	Lawrenciella_schaefferi       ????????????????????????0??????????-????????????0??????????????????????????????????????????????????????????0????????????????0??10110010020200101010011111101100----------000101?1000000010-011?11001111000?????????????0????????????????????????????????????????????????????????????????-00????????????????????
	Lepisosteus_osseus            11000100-12010100101000001002010111-1----00211100112-03100111101100100000100011000011001110301200100101100010110100210011-0-????0??????22020011101-00--00--1211-01101100-0011201100001001111101111101110??11100110100110111101020101011000-01011100100011101000001011111110010011000000100000-10000011100000000
	Leptolepis_bronni             1100010010111100100???0001012110011-1----00010-000-2-01110100000--?1000111020010000120???2???1210?021010?11?0110000210101-11???10??0010?2??0011??0-00--02011210??000000111??12101?-1111???????????????11-0??????????01001111111??101???110-0?0111???1?01110100000121101110?01110?0-0000100000-?0?--010100000---
	'''Ligulalepis'''             11??????????????1?1????00??00000?01?00000-0?????0???????00????????0????????????????????????????????????????-???????????????????00?0010??0?000?0?01?01????????00----------0000---------1101?????????????????????????????0????????????????????????????????????????????????????????????????--??????????????-????--
	Luederia_kempi                ??0?????0??????????????????????????????????????????????????????????????0????????????1??????????????????????0????????????????0??10?100???20200101010010?1211?100---?------0??111???0000??10?????????????????????????????0????????????????????????????????????????????????????????????????-00????????????????????
	Luganoia_lepidosteoides       11001100001011-010000??0011--11-000?1----1-1111-00-100200120?0?101010010?00000100001201?02?????0??0?101???110110000200001?????????????????????????????????????????0?????????1??0??????????????????????????????011010?1?01101?????????????0-0?21??001?101110100000121??1111????????-00001???00-?010001010????0-0
	Macrepistius_arenatus         110??10001??10????0??0?0010120????1-00000-10111?0112-03?012001?0--?100?01?0????0000120???1???1?0??????????0?010010021010???????10??0010120?0011?00-00????????101001000001111121?12-1001???????????????????10??01???0?11?????????????-????0-0??1???????????????????????11100???????000101100???1????010100???--0
	Macrosemimimus_lennieri       110001000???1110110?0000010120?0???-1----00010-?0112-?1-012111?0--0100?01100???0000120???10????0?10?1000????010010021010????????0??00???202001???0-00???0--1?11-01011100-11?12110?00?10???????????????10??????011??0011????????0?????????0-0?01?1????1???1010000012???????????????000101?????-?????01010????000
	Macrosemius_rostratus         11000100012110?01?000?0001012100?11-1----00011010112--0-00001-?0--01010011000?10000120??01040120?1001000000101001002-1101-0-???????0????2??0011??0-00????????11-0100110?-1?01211???00?????????????????1???????011??0?1101111?10??1????1?00-01?101000??011?0100?10121111111?0?00?0000--0100010?1111001010-???000
	Meemannia_eos                 11??????????0???????????0{0 1}002000??1-1----01?0???????????00????????0????????????000011???0?0?00-0?1?????????????????????????????0???0??????1???000???1?0??????0?----------000??????????????????00---0?010001111????????????????0???????????????????????????????????????????????????????0????????0???0???????????
	Melanecta_anneae              1101-01000?011001011010001002?10001-?----0??????00-1?0?0?0000-0?0?1111?00000??100001?0?????????0????????????01100000000??????0??????????????????????????????????????????????1?????????????????????????????????00???0?11??????????????????0-0?00??????1??11011011112???1100????????-0??00???000?0???010101?????0
	Trawdenia_planti              110?001?00??11001011010?010120100?1-1----001?1??00-1001?{0 1}0000-?10101110000000?10000120???1???110????????????01100000000101???0??0??????1??????????????????????0??????????????????????????0-011110001?1?0??????011010011???011????????????1000000?101000111010011111????000????????-00000???000??????1010????000
	Miguashaia_bureaui            110001-0000010-10100?00011000010000001000-?010-001?1?0000110-010--00?-------0001000120?0020?113-100??????0?001000010000-000-????????????????????????????????????????????????1?0?0?????0???????????????????10??01---0010??????1?00????????0-0010?1????0??0101?000001???0001?????0?1?0--00????0-00???000?0???-??-
	Mimipiscis_bartrami           11000010000001001011010000-01010011-00001-10111000-1000010000-01001111100000011000011010020?00-00?0100000000011010000000001000?10?00?00?0010000001001??12001000----------000100100000-0?110???????????1???????0110100110??00110??111010001000000??1?0101110100211120?0?100?0000?-1-0--00-000000000001010000-00-
	Mimipiscis_toombsi            11000010000011001011011000-02010011-00001-10111000-1000010000-01001111100000011000011010020100-0010100000000011010000000001000?10?00000000100000010010012001{0 1}00----------000100100000-0111000100---1001000101001101001101000110??11101000100000001110101?10100211110?0110010000?-1-0--00-000000000001010000-?0-
	Moythomasia_durgaringa        1101-01000?011001011010000-12010011-00010-110??100-1110011000-01001111100000001000011010010100-0010100000000011010000001001000?10?00011010100000010010011001000----------00010110000000??10???????????1?0?1?1001101101101000110??1110100010000000111?1?1110100211110??11?0?0100?-1-0--00-000000000001010000-00-
	Moythomasia_lineata           11000011000111001011010000-02010??1-00000-1101?100-1000110000-010?1111?000001010000110???1???0-0?1??????????01001000?0010010??????????????????????????????????????????????????????????????????????????????1010011010?11??????????????????100?00??11??101010?0021012???11?01???????-0--?0???000?0???01010????00-
	Moythomasia_nitida            11000110000111001011010000-02010001-00000-11111100-1102010000-01001111?000001010000110???1?????0????????????1110000000010110??????????????????????????????????????????????????????????????????????????????1?10011011?11??????????????????100?00?????01??11010021111???1100???????0-000?0???000?0???01010????00-
	Obaichthys_decoratus          110?01000?2010?0??0????0010020?0??1-1----001110?0??2--3-0?1111?0--?100001100???00???2010110??120010?10?1?0010100100210101-0-???????0???????001???0-00???0--121?-?11?1100-???1201100??1????????????????????????01101001101?1101020??1??1??0-010011?0?000111010000011?????1?001?0?10?0000100?00-10???01010?000000
	Onychodus_jandemarrai         11000000000000000001000111000010?0011---000010-010-110000-100111010111100000000000101000?00100-?0100-00?00--01100010000-000-??100?010???0??000??01?11??????1??0?????????????10000000000????????????11?????00--01---001000001010--10??00??101010010---0?0??01?0000020??001100?????1-0--00-0?0?-?0-00010?00??-00-
	Osorioichthys_marginis        1100001000?011001011001001001110?01-00000-?1010000-1001010000-010?1111?00?001010000110???1??????????????????0110000000010010?0??0???0??0????????????????????????????????????????????????110???????????????????011010011??????????????????0-0?00?0?110001110??02???????????????????-000?0???????0???01010????000
	Osteolepis_macrolepidotus     110011-000-000000?-0001110-00010100100000-0010-010-1000001000-?0--010000000000000001?00???0200-0??0??0????0001100010000100101???0?01??????????0?1????????????00----------000?????????0????????????????????????010101?101???????--???????00-1?10??????0??01010100002???00010????0?1-0--00?0?00-?0???0???00???00-
	Ozarcus_mapesae               0?-?-??-----??-??--??????????-???????????--???????-???-???----?????-???-----????-?1??-????????---?0?000??---?----------?-??-??0000001???0001--0?00-01??0---0-00----------0000---------1????????????????????????????????00000000--1010000??-?-?---????????????????????????????????------0--0-?---------------???
	Peltopleurus_lissocephalus    110?0011000111001011000001012010001-1----00010-000-10020002001010?01100100000?100001201???0????0??0?????????0110000100001?????????????????????????????????????????0???????????????????????????????????????????01???0?11?????????????-????0-002??1????1??11010000112???1010????????-00000???000?0?0001010????000
	Platysomus_superbus           111--10?----0100101101-001012010001-1----001111000-1000000000-00--01110?00000010?0012????2?????0????????????01100001000001?????????????????0?1??????????????????????????????12100?0001????????????????????????011010?11??????????????????0-0?00??????0??0?010000012???110?????????-0--?0???010?0???010100???000
	Polyodon_spathula             111---------00?10---01-000-011?0111-??????01010000-2----10000-10--00-------0-1-0000120?00-1--0-01100011-100100-0002---0-1-0-0-0000000-?20021-00100-00--10--1211-100101?0-00?12111200111111000011100???10100---011-00000000?100-??101000000-0000010010001111100000120011100?01000-1-0---000000?0-0------00000000
	Polypterus_bichir             11000110000010000000010001002011101--------2111000-10030000---010101101000000010000120100203010001001010001001001000000-000-00001??00-000011--0001-00??02001110???0?00?000001210100001?101001011000???????101001101001000100010--101110010-000001000000101010000012011101100100100-0000000000-00000011?00000000
	Porolepis_sp.                 1100100000-00000??-??0111110001110010---0-0010-010-1000000000-10--010000000000?000102001??0200-00?00000-?000011000100000001010100??10???01000000?10?????0--1-00----------0001000010000?1110?000?---???????110-01110101011000010--?00???100-1010??0???0???0010?00001???100?????????-0--?0-?000-?0???0???00???00-
	Propterus_elongatus           11000100???1101?11000?00?1002110101-1----00????1?112--0-002010?0--01010011000?10000120?????????00?0?10?0??0?01001002-1001-0-???????????????0?1????????????????????0?????????1?????????????????????????????????011????11???11?100?1???????0-00?0110??0?011?0100???02?1111100?1??100?0--0100?000111??01010000000-
	Psarolepis_romeri             1101-1-000?0??-0???0001?10-120100?????????0??0-??0-????????00?0?--??11100000000001101001000000-??1???0???0000??0??00?00?????00100??11???0?10000????01????????00----------0001000000000????????????????????1111011111????????????????????????0?0??0?????1??1???????????????????????-0??00-0?????0?00????????????
	Pteronisculus_stensioi        110000100001110?1011010001012010001-0010101111000102002100000-01010111100000101000011010010200-0110200000000110000000000011000010?10011020200101011010112101100------?--?000101010000001110111111001111010????011010011011011100111101100100000011?101??11010011112000010000000?011000000000000000001010000-11-
	Raynerius_splendens           11????????????????1?1???00-01?10001-000??-?111?00????1001??????1101111?0000?001000011010010?00-0?101000000???100000001000011??0100000???10100000010010011001000----------000100?0000000????0010????10010?01???011011???010001????111010001000?000??????????????????????????????????0--00-00????0?000?01?0?????-
	Saurichthys_madagascarensis   111--000----1?-0101-1--00?-02100??1-1----0?0011?00-?0?1?001000011?01110000000?100001201?01?????0?100?000??0001200000000-1-0-00010??00??12021-0?100-00??1?????0-----------??0121?00????00111011????????????????010-00011??????????????????0-0020011?100011101002111101111110???0??1-00000???0000?000011?0000-000
	Saurichthys_Greenland         11???00?---????0101-1--001002110??1-1----01??????0-200100?100001-0?1100000000??00???001101???0-0?1001000?0000?????00?0?-01??000100100?0120?1-00100-00??10--020-----------?01121?001-0100111011?????1101001?????????????010011????1?1??????????0???????????????????????????????????-000?0000?????00?0????000-000
	Saurichthys_ornatus           111--000----???0101-1--0010021?0001-1----010011000-20010001000011001100000000?100???101?01???0-0?1000000100001200000000-????000100100?012021-00100-00--10--020-----------?0?121?001-01?01110?1????????????????010-000110?00??????101?1???0-0020?1??100011101002111?011????0??????1-000?00000000?00?01?-0000-000
	Scanilepis_dubia              110??010????11??1?0????00100001???1-1----01111??0??1?0310-20?0?10?0111?00?00???00???20?????????0????????????11100?00000000????????????????????????????????????????????????????????????????????????????????1010011010110??????????????????0-0??????????????0100???12????0???????????000?0????10??????00????????-
	Semionotus_elegans            1100010001211-1-1101000001012000?11-1----00011100112-01-002111?0--0100000101001000012010?10?0120010?10?000010100100210101-0-??????????????????1??????????????1??01?1110?-??11?11000001????????????????????101001????011011?101101111011000-010111000??01110100201121111111?0100100000101-0?00000000010100000000
	Styloichthys_changae          1???-????0????-??0???????0-0?0100?0???????0???????????????????????010000000??0010???200???0??0-0?0???0???0?0??????0000??????1?100??11???0100000??0-1????0--1-00----------0001000000000????????????????????1111?????????1???????????????????00????1?????0??0????????????????????????0??00-00????0?000???????????
	Styracopterus_fulcratus       111--101----110010100?-0010120????1-1----010111?00-1000?00000-?101?111?000000?101001211??20??0-0??1?????????0120010000010110????????????????????????????????????????????????1?????????0???????????????????????0110100110?1???????????????0-0?00?0????0??21010021112???0100????????-0--00???000?????01010????000
	Tanaocrossus_kalliokoskii     ????????????????????????????????????1??????1???????????????????????11??????0?????????????????????????????????1100????00?????????????????????????????????????????????????????????????????????????????????????????1??0110????????????????????????????????????10?????????1?11?????????????????000?????01010???????
	Tegeolepis_clarki             11000000?0?01100101?0??0010010?0001-00000-1101??011?000010000-0????111?00?000?10010110??11?????0?????????????11??0000?000110?0??????????????????????????????????????????????1?100?000?????????????????????????00101???0?00?????????????????0?????????0??110??001011????1?????????010--00?0?00-?0???0101?????00?
	Tetragonolepis_semicincta     110001000010110010010000010020?0011-1----00011000102103-01100000--01000011000?10000120???104-??0010?10101?1?0100100210101-11?0?10??00?????20011??0-00????????0??????????????1?1??2-11?0???????????????????????011??0011011?1011101???????0-000101?110101110100201121111111??100000100101???111111??010100000001
	Venusichthys_comptus          110001000001110010110000010121?0??1-1----00011?00102-03000000000--01010000000010000120?????????0??0?????????0100001210001-1???????????????????????????????????????????????????????????????????????????????????011?10011??????????????????0-0?01?1????1??11010000012???1100?????????00001???00-?0???0201?0???000
	Watsonulus_eugnathoides       1100000011111-1-1000000001012000011-1----00010100112100010100100--01010011110010000120??010201201102101010?10110100110110-110001??0001112020011??1101--?0--1210100100010011112111000000???????????????10??????01???0?1?0111001100101???0?100020?11???1??21010000012?-?11100??00?0010--0000?0?-10000010100???001
	Wendyichthys_dicksoni         111---------110010110?-001012010001-00110-1111100111002111000-110?0111?000000010000120??0????0-0???2????????0100000000010110?0??????0???2????0???1??1???????????????????????101???????????????????????????????01???1011??????????????????{0 1}-0?00?1????1??01010010112???1100????????1000?0???00000???010100???00-
	Woodichthys_bearsdeni         1100001000?011001011010001012010001-00100-1111?000-1000100000-010?1111?000000010000110???1???0-0??0000000???011000000000010??0?????0?????0?0?10??1001??11101100----------00010100?00000???????????????????????0110110110?????????????????0-0?00???1101??11010011112???1100????????-0--00-??00??0???01010????00-
	Gibbodon_cenensis             1100010000?1???0----?00001-1200???1?-----1------00-0--1-000---?0--0100?11???00?1000120???2????????1??????01?01-1--0200001-?-??????????????????????????????????????????????????????????????????????????????????011??0011??????????????????0-???1?1????????10??121112???1111????????-000?1???111??11?1011?????---
	Yelangichthys_macrocephalus   111--100----10-0101-1--001002100001-1----01?1??000-??0??00?????????11?0000?00?000???0010020??0-0??1?10?0?010????????0???????00010??00?0010?1-00?00-00??10--?20----------???0121?10??01???11?????????????????????????????????????????????????020???????????????????????????????????????001?0????000?0????0???00-
	Eomesodon_liassicus           11000100001101-0----000001-02001001------1------00-0--0-010---?0--0?????1???0?10000120???2?????0111?1?01?01101-1--1200001-0-????????????????????????????????????????????????12????????????????????????????????011???0000?0???11?0????????0-0?01?1??00001110??02?1121??1011?01000-0-0--?1?1?0110111111101100?---
	Brembodus_ridens              110001000011??-0----?00001-1200???1------1------00-0--1-000---?0--0?????1??????1000120???2????????1??????01101-1--0200001-?-???????????????0????????????????????????????????121????0??????????????????????????011??00110?1???????????????0-0??1?1??000011101002?1121??1111?010?0-0-000?111?0110111111111111?---
	Arduafrons_prominoris         11000100001100-0----?00101-020????1------1-?????00-?--0-000---?0--?100001??0???0000120????????????1??????0??01-1---200001-0-??????????????????????????????????????????????????????????????????????????????????011??0010????????????????????0??1??????????????021111????011?01???-0-1--?1???0110?11?11101???1---
	Mesturus_sp                   11??????????????????????01-0200???1-?????1??????????????00??????????????????????????????????????????????????????????????????0??-0??0010?20?0011?10-01--0?????101000000000111121?1201111????????????????????????????????0????????????????????????????????????????????????????????????????11??????????????1???---
	Mesturus_verrucosus           11000100101100-0----000100-0200???1-?????1??????10----0-000---?0--?100011100???0000120???2???100??1??????01?01-1---200001-0-??????????????????????????????????????????????????????????????????????????????????011??00100?0?10??????????????0?21????000011101?02??1?1??1011?01????0----?1???0110?1111110??10?---
	Neoproscinetes_penalvai       11000100?02110-0????00000?-1210???1-?????0?-----0??2????--0-?-?0--?110011?000?0100012010121--100111-1001?01101-1--02010?1-0-?-0-0010000?20?0011?10-01??0---1211-?1011??0?1111210?20110?01-????????????????????01???001?01011011?0100?????0-0?21?1??00001110100201111-11010001000-0??---11100110?1111?1??1111??-
	Brachydegma_caelatum          11?????00???????????????01012000??1-1----0100???0??2003000?0000101?111000000??1000012010?1????-011001010?0??11100001000101110??10??00???10?1-0??????1???2001210??0????00????1210?200000???????????????????1010011010?1?0110011110100011000-0100011???101?10???????????????0????????000?0000???0?0000101?00??000
	NHMD_73588                    110001000?00110?101?01000??020?0001-001?100??1?0010200210?000-01010111?000001?100?0110100102012011020?00100?11001000000?????000???10?????021--01011?10?12101100------0--000010100001000?????11?????1001010????011010?110110111001??1011??10000001101111?110??0????????????1?????0?100000000??00?000010?0000-11-
	Pteronisculus_cicatrosus      11??????????110?10110100010120100?1-0011?00111100102?03?01000-01010111?000001?100?0110??010200-011000?00?0??01000000000001??00??0??0??????200?0?011?1??12????0?------?--?000101010000?????????????????????????0110?0011???????0??1?1?????10000??????0?1?110100111120000100?????????000000000000000001010000-11-
	Pteronisculus_changae         111---------010?1011010001012010001-0011?00111100102002101000-0101011100000010100001101??10??0-0?1000000????01000000000001??00?????0???????00?0????0?????????0?------?--????1?1010000?????????????????????????0110100110?1??110??1?1?????1000000????0???110100111120000100??000?011000?00?00000000001010000-11-
	Pteronisculus_magnus          11??????????110?1011010001002010001-0010100111000102002100000-0101011110000010100001101?010200-0110000001000110000000000011000?10?10011020200101011010112101100------?--?000101010000001110111111001011010????01101001101101110011?10110010000001101011?1101?011112000010000000?011000000000000000001010000-11-
	Pteronisculus_nielseni        111---------010?1011010001012010001-0010?00111100102002100000-01010111?0000010100001101??10??0-0?10?0000????010000000000011000?????0???????00?0????0?????????0?------?--????1?1010000?????????????????????????0110100110?1??110??1?1?????1000000????0???110100111120000100000000011000?00?00000000001010000-11-
	Pteronisculus_gunnari         110001000?00110?10110100010{0 1}2{0 1}10001-0010100111000102002100000-01010111100000101000011010010200-0110200001000110000000000011000010?10011020200101011010112101100------?--?000101010000001110111111001001010????01101001101101110011?10110010000001101011?110100111120000100000000011000000000000000001010000-11-

;

END;
BEGIN ASSUMPTIONS;
	OPTIONS DEFTYPE = UNORD POLYTCOUNT = MINSTEPS;
	TYPESET * UNTITLED   =  unord:  1- 89 91 93- 152 154- 173 175- 248 250- 258 260- 303, ord:  90 92 153 174 249 259;
TYPESET default   =  unord:  1- 89 91 93- 152 154- 173 175- 248 250- 254 256- 258 260- 303, ord:  90 92 153 174 249 255 259;

END;

BEGIN NOTES;

	TEXT  TAXON = 2 TEXT = Histo?;
	TEXT  TAXON = 4 TEXT = Only_dermal_skull_coded_;
	TEXT  TAXON = 47 TEXT = 'Dpt/it/stBraincaseBrainLabyrinthHistologyFins';
	TEXT  TAXON = 51 TEXT = Code_from_scan;
	TEXT  TAXON = 64 TEXT = Histology;
	TEXT  TAXON = 89 TEXT = 'Dpt/it/stAortaLabyrinthHistology';

	TEXT   CHARACTER = 1 TEXT = 'Forey, 1980; Gardiner, 1984; Zhu & Schultze, 2001; Zhu et al., 2001; Zhu & Yu, 2002; Zhu et al., 2006; Friedman, 2007; Brazeau, 2009; Zhu et al., 2009; Friedman & Brazeau; 2010; Davis et al., 2012; Zhu et al., 2013; Brazeau & Friedman, 2014; Giles et al., 2015b,c; Giles et al., 2017.';

	TEXT   CHARACTER = 2 TEXT = 'Brazeau, 2009; Zhu et al., 2013; Giles et al., 2015b; Giles et al., 2017. ^n';

	TEXT   CHARACTER = 3 TEXT = 'Hurley et al. 2007; Xu et al. 2014; Giles et al., 2017. Typically, the premaxilla is a short, paired or median bone that contributes to the orbital margin anterior to the maxilla. However, considerable variation is present, and we have attempted to consistently code this variation as laid out here and in the following character descriptions. A premaxilla may be completely absent (e.g. Acipenser, Cyranorhis) or dorsally expanded into a midline bone (possibly fused with the rostral; e.g. Bobasatrania, Styracopterus). These two latter states are coded as ?1? here. ';
	TEXT   TAXON = 10 CHARACTER = 3 TEXT = 'chaged from 0 to 1, as we consider this element to be compound (premaxilla fused with rostral) as in Saurichthys (Nielsen 1949; Romano 2007; Romano and Brinkmann 2009). All relevant characters have been recoded based on this interpretation.';
	SU  T = 10 C = 3 N = color I = 5;

	TEXT   CHARACTER = 4 TEXT = 'Cloutier & Ahlberg, 1996; Taverne, 1997; Schultze & Cumbaa, 2001; Zhu & Schultze, 2001; Zhu & Yu, 2002; Cloutier & Arratia, 2004; Friedman & Blom, 2006; Zhu et al., 2006; Friedman, 2007; Long et al., 2008; Swartz, 2009; Choo, 2011; Giles et al., 2017. Coded as inapplicable in taxa lacking any ossification in the position typically occupied by the premaxilla (e.g. Acipenser, Cyranorhis) and where the premaxilla appears fused with the rostral (e.g. Bobasatrania, Styracopterus).';
	TEXT   TAXON = 10 CHARACTER = 4 TEXT = 'changed from 0 to -';
	SU  T = 10 C = 4 N = color I = 5;
	TEXT   TAXON = 24 CHARACTER = 4 TEXT = fused;
	TEXT   TAXON = 26 CHARACTER = 4 TEXT = fused;

	TEXT   CHARACTER = 5 TEXT = 'Xu et al., 2011; Xu et al., 2015; Xu & Zhao, 2016; Giles et al., 2017. Coded as inapplicable in taxa lacking any ossification in the position typically occupied by the premaxilla (e.g. Acipenser, Cyranorhis) and where the premaxilla appears fused with the rostral (e.g. Bobasatrania, Styracopterus) and where the premaxilla appears fused with the rostral (e.g. Bobasatrania, Styracopterus).';
	TEXT   TAXON = 10 CHARACTER = 5 TEXT = 'changed from 1 to -';
	SU  T = 10 C = 5 N = color I = 5;

	TEXT   CHARACTER = 6 TEXT = 'Friedman, 2007; Giles et al., 2015b; Giles et al., 2017. Coded as inapplicable in taxa lacking any ossification in the position typically occupied by the premaxilla (e.g. Acipenser, Cyranorhis) and in taxa where the premaxillae do not contact at the midline.';

	TEXT   CHARACTER = 7 TEXT = 'Cloutier & Ahlberg, 1996; Schultze & Cumbaa, 2001; Zhu & Schultze, 2001; Zhu et al., 2001; Zhu & Yu, 2002; Cloutier & Arratia, 2004; Zhu et al., 2006; Friedman, 2007; Long et al., 2008; Swartz, 2009; Zhu et al., 2009; Xu & Gao, 2011; Zhu et al., 2013; Xu et al., 2014; Giles et al., 2017. Coded as inapplicable in taxa lacking any ossification in the position typically occupied by the premaxilla (e.g. Acipenser, Cyranorhis), where the premaxilla appears fused with the rostral (e.g. Bobosatrania, Styracopterus), and where the premaxilla is restricted anterior to the orbit.';
	TEXT   TAXON = 10 CHARACTER = 7 TEXT = 'changed from 0 to -';
	SU  T = 10 C = 7 N = color I = 5;
	TEXT   TAXON = 110 CHARACTER = 7 TEXT = 'changed from - to 0';
	SU  T = 110 C = 7 N = color I = 5;

	TEXT   CHARACTER = 8 TEXT = 'Cloutier & Arratia 2004, Xu et al. 2014; Giles et al., 2017. Coded as inapplicable in taxa lacking any ossification in the position typically occupied by the premaxilla (e.g. Acipenser, Cyranorhis).';

	TEXT   CHARACTER = 9 TEXT = 'Arratia 1999; Cavin & Suteethorn 2006; Hurley et al. 2007; Giles et al., 2017. Coded as inapplicable in taxa lacking any ossification in the position typically occupied by the premaxilla (e.g. Acipenser, Cyranorhis) and where the premaxilla appears fused with the rostral (e.g. Bobasatrania, Styracopterus)';
	TEXT   TAXON = 10 CHARACTER = 9 TEXT = 'changed from 0 to -';
	SU  T = 10 C = 9 N = color I = 5;

	TEXT   CHARACTER = 10 TEXT = 'Grande 2010; Xu et al., 2015; Xu & Shen, 2015; Xu & Zhao, 2016; Giles et al., 2017. Coded as inapplicable in taxa lacking any ossification in the position typically occupied by the premaxilla (e.g. Acipenser, Cyranorhis) and where the premaxilla appears fused with the rostral (e.g. Bobasatrania, Styracopterus).';
	TEXT   TAXON = 10 CHARACTER = 10 TEXT = 'changed from 0 to -';
	SU  T = 10 C = 10 N = color I = 5;

	TEXT   CHARACTER = 11 TEXT = 'Gardiner & Schaeffer 1989; Olsen & McCune 1991; Gardiner et al. 1996; Gardiner et al. 2005; Cavin & Suteethorn 2006; Hurley et al. 2007; Grande 2010; López-Arbarello 2012; Xu & Wu 2012; Xu et al. 2014; Bermúdez-Rochas & Poyato-Ariza 2015; Poyato-Ariza 2015; Thies & Waschkewitz, 2015; Xu & Shen, 2015; Gibson 2016; Xu & Zhao, 2016; Giles et al., 2017; López-Arbarello & Sferco 2018. Coded as inapplicable in taxa lacking any ossification in the position typically occupied by the premaxilla (e.g. Acipenser, Cyranorhis) and where the premaxilla appears fused with the rostral (e.g. Bobasatrania, Styracopterus).	';
	TEXT   TAXON = 10 CHARACTER = 11 TEXT = 'changed from 1 to -';
	SU  T = 10 C = 11 N = color I = 5;

	TEXT   CHARACTER = 12 TEXT = 'Giles et al., 2017. Coded as inapplicable in taxa lacking any ossification in the position typically occupied by the premaxilla (e.g. Acipenser, Cyranorhis) and where the premaxilla appears fused with the rostral (e.g. Bobasatrania, Styracopterus).';
	TEXT   TAXON = 10 CHARACTER = 12 TEXT = 'changed from 0 to -';
	SU  T = 10 C = 12 N = color I = 5;

	TEXT   CHARACTER = 13 TEXT = 'Cloutier & Ahlberg, 1996; Taverne, 1997; Lund, 2000; Schultze & Cumbaa, 2001; Zhu & Schultze, 2001; Lund & Poplin, 2002; Cloutier & Arratia, 2004; Friedman & Blom, 2006; Long et al., 2008; Swartz, 2009; Choo, 2011; Xu et al., 2014; Giles et al., 2015b; Giles et al., 2017.';
	TEXT   TAXON = 52 CHARACTER = 13 TEXT = how_to_code_these?_all_fused.;

	TEXT   CHARACTER = 14 TEXT = 'Gardiner & Schaeffer, 1989; Taverne, 1997; Friedman & Blom, 2006; Long et al., 2008; Swartz, 2009; Choo, 2011; Giles et al., 2015b; Poyato-Ariza 2015; Giles et al., 2017.';
	TEXT   TAXON = 31 CHARACTER = 14 TEXT = 'no rostral (paired or median)';
	TEXT   TAXON = 95 CHARACTER = 14 TEXT = 'changed from 1 to ?. Due to acuteness and incompleteness and/or fusion of the snout, its precise constituents remain unknown';
	SU  T = 95 C = 14 N = color I = 5;
	TEXT   TAXON = 106 CHARACTER = 14 TEXT = bone_absent_so_inapplicable_;

	TEXT   CHARACTER = 15 TEXT = 'Gardiner et al. 1996; Hurley et al. 2007; Giles et al., 2017.';
	TEXT   TAXON = 31 CHARACTER = 15 TEXT = 'no rostral (paired or median)';

	TEXT   CHARACTER = 16 TEXT = 'Friedman, 2007; Giles et al., 2015b; Giles et al., 2017.) ';

	TEXT   CHARACTER = 17 TEXT = 'Taverne, 1997; Schultze & Cumbaa, 2001; Friedman & Blom, 2006; Long et al., 2008; Swartz, 2009; Choo, 2011; Giles et al., 2015b; Giles et al., 2017.';
	TEXT   TAXON = 111 CHARACTER = 17 TEXT = 'unlear whether large bone is a nasal, antorbital or infraorbital';

	TEXT   CHARACTER = 18 TEXT = 'Giles et al., 2017.';

	TEXT   CHARACTER = 19 TEXT = 'Xu & Wu 2012; Xu et al. 2014; Xu & Zhao, 2016; Giles et al., 2017; López-Arbarello & Sferco 2018.';
	TEXT   TAXON = 27 CHARACTER = 19 TEXT = '0	';

	TEXT   CHARACTER = 20 TEXT = 'Lund et al., 1995; Ahlberg & Johanson, 1998; Ahlberg et al., 2000; Lund, 2000; Poplin & Lund, 2000; Schultze & Cumbaa, 2001; Lund & Poplin, 2002; Cloutier & Arratia, 2004; Zhu & Ahlberg, 2004; Daeschler et al., 2006; Long et al., 2006; Zhu et al., 2006; Zhu et al., 2009; Choo, 2011; Giles et al., 2015b; Giles et al., 2017; López-Arbarello & Sferco 2018.';
	TEXT   TAXON = 27 CHARACTER = 20 TEXT = notches_ventreal_margin_of_nasal_;
	TEXT   TAXON = 32 CHARACTER = 20 TEXT = '''nasal aperture is single as in dapedium & lepidotes (gardiner 1963)''';
	TEXT   TAXON = 34 CHARACTER = 20 TEXT = ventral;
	TEXT   TAXON = 95 CHARACTER = 20 TEXT = 'changed from 1 to - following the addition of character indicating the presence of both nostrils within single ossification';
	SU  T = 95 C = 20 N = color I = 5;

	TEXT   CHARACTER = 21 TEXT = 'In saurichthyids both nares are accommodated in the so-called nasalo-antorbital bone (Argyriou et al. 2018)';

	TEXT   CHARACTER = 22 TEXT = 'Friedman & Blom, 2006; Long et al., 2008; Choo, 2011; Giles et al., 2015b; Giles et al., 2017. This character refers to the opening of the posterior nostril with reference to the surrounding dermal bones.';
	TEXT   TAXON = 95 CHARACTER = 22 TEXT = 'changed from 0 to - following the addition of a character indicating the presence of both nostrils within single ossification';
	SU  T = 95 C = 22 N = color I = 5;

	TEXT   CHARACTER = 23 TEXT = 'Friedman & Blom, 2006; Long et al., 2008; Choo, 2011; Giles et al., 2015b; Giles et al., 2017.';

	TEXT   CHARACTER = 24 TEXT = 'Lund et al., 1995; Cloutier & Ahlberg, 1996; Lund, 2000; Schultze & Cumbaa, 2001; Zhu & Schultze, 2001; Zhu et al., 2001; Lund & Poplin, 2002; Zhu & Yu, 2002; Cloutier & Arratia, 2004; Zhu et al., 2006; Friedman, 2007; Swartz, 2009; Zhu et al., 2009; Zhu et al., 2013; Giles et al., 2015b; Giles et al., 2017. ';

	TEXT   CHARACTER = 25 TEXT = 'Cloutier & Ahlberg, 1996; Ahlberg & Johanson, 1998; Zhu & Ahlberg, 2004; Zhu & Schultze, 2001; Zhu et al., 2001; Zhu & Yu, 2002; Daeschler et al., 2006; Long et al., 2006; Zhu et al., 2006; Friedman, 2007; Brazeau, 2009; Zhu et al., 2009; Choo, 2011; Davis et al., 2012; Zhu et al., 2013; Giles et al., 2015b; Giles et al., 2017.';

	TEXT   CHARACTER = 26 TEXT = 'Cloutier & Ahlberg, 1996; Taverne, 1997; Schultze & Cumbaa, 2001; Zhu & Schultze, 2001; Zhu & Yu, 2002; Friedman & Blom, 2006; Friedman, 2007; Long et al., 2008; Brazeau, 2009; Swartz, 2009; Davis et al., 2012; Zhu et al., 2013; Xu et al., 2014; Giles et al., 2015b,c; Giles et al., 2017. A pineal foramen is variably present in Cheirolepis canadensis (Pearson & Westoll, 1979; Arratia & Cloutier, 1996), C. trailli (Pearson & Westoll, 1979), Kentuckia deani (Rayner, 1951) and Meemannia (Zhu et al., 2010), and these taxa are coded ''0/1'' to reflect this polymorphism.';

	TEXT   CHARACTER = 27 TEXT = 'Friedman, 2007; Zhu et al., 2009; Giles et al., 2015b; Giles et al., 2017. Can only be coded in taxa that lack a pineal foramen. ';
	TEXT   TAXON = 117 CHARACTER = 27 TEXT = 'changed from - to 0 to be consistent with other taxa';
	SU  T = 117 C = 27 N = color I = 5;

	TEXT   CHARACTER = 28 TEXT = 'Dietze, 2000; Schultze & Cumbaa, 2001; Cloutier & Arratia, 2004; Friedman & Blom, 2006; Long et al., 2008; Swartz, 2009; Choo, 2011; Xu et al., 2014; Giles et al., 2015b; Giles et al., 2017.';
	TEXT   TAXON = 7 CHARACTER = 28 TEXT = two_pairs..._but_together_they_are_rectangular_;
	TEXT   TAXON = 10 CHARACTER = 28 TEXT = '''l'' shaped...';
	TEXT   TAXON = 19 CHARACTER = 28 TEXT = coding_revised_from_1_to_0_following_Hilton_and_Forey_2009;
	SU  T = 19 C = 28 N = color I = 5;
	TEXT   TAXON = 123 CHARACTER = 28 TEXT = 'variability in parietal shape in Nielsen original description (p. 133)';

	TEXT   CHARACTER = 29 TEXT = 'Lund et al., 1995; Taverne, 1997; Dietze, 2000; Lund, 2000; Poplin & Lund, 2000; Schultze & Cumbaa, 2001; Lund & Poplin, 2002; Cloutier & Arratia, 2004; Friedman & Blom, 2006; Zhu et al., 2006; Long et al., 2008; Swartz, 2009; Choo, 2011; López-Arbarello 2012; Xu et al., 2014; Bermúdez-Rochas & Poyato-Ariza 2015; Thies & Waschkewitz, 2015; Gibson 2016; Giles et al., 2015b; Giles et al., 2017; López-Arbarello & Sferco 2018.';
	TEXT   TAXON = 35 CHARACTER = 29 TEXT = unclear;

	TEXT   CHARACTER = 30 TEXT = 'Arratia 1999; López-Arbarello 2012; Bermúdez-Rochas & Poyato-Ariza 2015; Thies & Waschkewitz, 2015; Gibson 2016; Giles et al., 2017; López-Arbarello & Sferco 2018.';
	TEXT   TAXON = 123 CHARACTER = 30 TEXT = 'variability in parietal shape in Nielsen original description (p. 133)';

	TEXT   CHARACTER = 31 TEXT = 'Giles et al., 2015b,c; Giles et al., 2017. Although not figured, an anterior pit line is described for Miguashaia (Cloutier 1996).';

	TEXT   CHARACTER = 32 TEXT = 'Giles et al., 2015b,c; Giles et al., 2017.';

	TEXT   CHARACTER = 33 TEXT = 'López-Arbarello 2012; Bermúdez-Rochas & Poyato-Ariza 2015; Poyato-Ariza 2015; Giles et al., 2017; López-Arbarello & Sferco 2018. The supraorbital canal may terminate in the frontal/parietal, or it may become confluent with the infraorbital canal. The exact position of this junction is highly variable, and typically occurs in the region of the frontal, dermosphenotic, dermopterotic.';
	TEXT   TAXON = 34 CHARACTER = 33 TEXT = 'p.431, not figured';
	TEXT   TAXON = 74 CHARACTER = 33 TEXT = changed_from_?_to_0;
	SU  T = 74 C = 33 N = color I = 5;

	TEXT   CHARACTER = 34 TEXT = 'Giles et al., 2017. In some taxa (e.g. Dapedium), the infraorbital canal continues anteriorly above the orbit a short way.';

	TEXT   CHARACTER = 35 TEXT = 'Lund et al., 1995; Cloutier & Ahlberg, 1996; Schultze & Cumbaa, 2001; Zhu & Schultze, 2001; Cloutier & Arratia, 2004; Long et al., 2008; Swartz, 2009; Giles et al., 2015b; Giles et al., 2017.';

	TEXT   CHARACTER = 36 TEXT = 'Giles et al., 2015b,c; Giles et al., 2017.';

	TEXT   CHARACTER = 37 TEXT = 'Gardiner & Schaeffer 1989; Cloutier & Arratia, 2004; Hurley et al. 2007; Choo, 2011; Giles et al., 2015b; Xu & Zhao, 2016; Giles et al., 2017. This character is reformulated from Choo''s character ''Dermopterotic: present/absent''. Rather than designating bones as an intertemporal (or supratemporal or dermosphenotic) a priori, we consider the number of bones carrying the otic portion of the lateral line canal between the dermosphenotic and the posterior edge of the skull roof. Where two bones are present, these are treated as the intertemporal and supratemporal; where only one is present, this is treated as the dermopterotic. Anamestic bones between the dermosphenotic and frontal are not included in this count.) ';
	TEXT   TAXON = 73 CHARACTER = 37 TEXT = 'Coates 1998 mentions "a flanking temporal series of which only the supratemporal is preserved". Revised from 1 to ?';
	SU  T = 73 C = 37 N = color I = 5;

	TEXT   CHARACTER = 38 TEXT = 'Taverne, 1997; Friedman & Blom, 2006; Choo, 2011; Giles et al., 2015b; Giles et al., 2017. Coded as inapplicable in taxa with a dermopterotic. Coded as inapplicable in taxa with a dermopterotic.';

	TEXT   CHARACTER = 39 TEXT = 'Friedman & Blom, 2006; Choo, 2011; Giles et al., 2015b; Giles et al., 2017. Coded as inapplicable in taxa with a dermopterotic.';

	TEXT   CHARACTER = 40 TEXT = 'Xu & Gao, 2011; Xu et al., 2014; Giles et al., 2015b; Giles et al., 2017. Coded as inapplicable in taxa with a dermopterotic.';

	TEXT   CHARACTER = 41 TEXT = 'Choo, 2011; Giles et al., 2015b; Giles et al., 2017. Coded as inapplicable in taxa with a dermopterotic.';

	TEXT   CHARACTER = 42 TEXT = 'Xu & Gao 2011; Xu et al. 2014; Giles et al., 2017. Coded as inapplicable in taxa with a separate intertemporal and supertemporal, and in taxa lacking these bones entirely.';
	TEXT   TAXON = 32 CHARACTER = 42 TEXT = 'tendency to co-ossification of skull roof bones in larger specimens ';
	TEXT   TAXON = 34 CHARACTER = 42 TEXT = '''fused in most specimens'' ';

	TEXT   CHARACTER = 43 TEXT = 'Lu et al. 2016; Giles et al., 2017.';
	TEXT   TAXON = 50 CHARACTER = 43 TEXT = 'changed from ? to 1 based on majority of specimens depicted in Sallan and Coates 2013, Fig. 6,7';
	SU  T = 50 C = 43 N = color I = 5;

	TEXT   CHARACTER = 44 TEXT = 'Gardiner & Schaeffer, 1989; Lund et al., 1995; Cloutier & Ahlberg, 1996; Grande & Bemis 1998; Coates, 1998; Lund, 2000; Poplin & Lund, 2000; Schultze & Cumbaa, 2001; Zhu & Schultze, 2001; Lund & Poplin, 2002; Cloutier & Arratia, 2004; Friedman & Blom, 2006; Long et al., 2008; Swartz, 2009; Choo, 2011; López-Arbarello 2012; Zhu et al., 2013; Bermúdez-Rochas & Poyato-Ariza 2015; Poyato-Ariza 2015; Giles et al., 2015b; Thies & Waschkewitz, 2015; Xu et al., 2015; Gibson 2016; Xu & Zhao, 2016; Giles et al., 2017; López-Arbarello & Sferco 2018. ';
	TEXT   TAXON = 95 CHARACTER = 44 TEXT = 'polymorphic - 0/1';
	TEXT   TAXON = 113 CHARACTER = 44 TEXT = 'canals not shown so unclear if the large ''nuchal'' posterior to the parietal is a large scale or an extrascapular';

	TEXT   CHARACTER = 45 TEXT = 'Giles et al., 2015b,c; Giles et al., 2017. The skull roof of Moythomasia as shown in Fig. 103 (Gardiner, 1984) is a restoration (see also Choo 2015: fig. 13). The only skull roof directly figured by Gardiner does not preserve the extrascapulars in situ (fig. 83). However, in specimens viewed by us, as well as in published photos of articulated material (e.g. Choo 2015: fig 8), the lateral extrascapular of M. durgaringa is clearly excluded from the lateral margin of the skull roof.';

	TEXT   CHARACTER = 46 TEXT = 'Dietze, 2000; Cloutier & Arratia, 2004; Long et al., 2008; Swartz, 2009; Choo, 2011; Xu & Gao, 2011; Zhu et al., 2013; Xu et al., 2014; Giles et al., 2015b; Giles et al., 2017.';

	TEXT   CHARACTER = 47 TEXT = 'Giles et al., 2015b,c; Giles et al., 2017. Inapplicable for taxa that possess a median extrascapular, as it is logically impossible for the lateral extrascapulae to meet in the midline.';
	TEXT   TAXON = 10 CHARACTER = 47 TEXT = recoded_from_0_to_1_based_on_Nielsen_1949;
	SU  T = 10 C = 47 N = color I = 5;

	TEXT   CHARACTER = 48 TEXT = 'Choo, 2011; Giles et al., 2015b; Giles et al., 2017.';

	TEXT   CHARACTER = 49 TEXT = 'Cloutier & Ahlberg, 1996; Ahlberg & Johanson, 1998; Zhu & Schultze, 2001; Zhu et al., 2001; Zhu & Yu, 2002; Zhu & Ahlberg, 2004; Daeschler et al., 2006; Long et al., 2006; Zhu et al., 2006; Friedman, 2007; Zhu et al., 2009; Giles et al., 2015b; Giles et al., 2017.';

	TEXT   CHARACTER = 50 TEXT = 'Gardiner & Schaeffer 1989; Lund et al. 2000; Cloutier & Arratia, 2004; Hurley et al. 2007; Choo, 2011; Xu & Gao, 2011; Xu et al., 2014; Giles et al., 2015b; Giles et al., 2017; López-Arbarello & Sferco 2018.';
	TEXT   TAXON = 29 CHARACTER = 50 TEXT = no_canal_though?_;

	TEXT   CHARACTER = 51 TEXT = 'Grande 2010; Xu & Wu 2012; Xu et al. 2014; Poyato-Ariza 2015; Xu & Shen, 2015; Xu & Zhao, 2016; Giles et al., 2017; López-Arbarello & Sferco 2018.';

	TEXT   CHARACTER = 52 TEXT = 'Cloutier & Arratia, 2004; Gardiner et al., 2005; Choo, 2011; Xu & Gao, 2011; Xu et al., 2014; Giles et al., 2015b; Giles et al., 2017; López-Arbarello & Sferco 2018.';
	TEXT   TAXON = 19 CHARACTER = 52 TEXT = 'coding revised from 1 to 0, according to Hilton and Forey 2009: "there is no trace of a postorbital in ?C. acipenseroides, as found in extant acipenserids"';
	SU  T = 19 C = 52 N = color I = 5;
	TEXT   TAXON = 27 CHARACTER = 52 TEXT = fragmented;
	TEXT   TAXON = 28 CHARACTER = 52 TEXT = fragmented;
	TEXT   TAXON = 29 CHARACTER = 52 TEXT = some_lost_so_fragmentation_unclear__;
	TEXT   TAXON = 32 CHARACTER = 52 TEXT = fragmented;
	TEXT   TAXON = 68 CHARACTER = 52 TEXT = 'coding revised from 0 to 1, Bürgin 1992 describes an anterior tube-like infraorbital element (lachrymal)';
	SU  T = 68 C = 52 N = color I = 5;
	TEXT   TAXON = 74 CHARACTER = 52 TEXT = ?_to_1;
	SU  T = 74 C = 52 N = color I = 5;
	TEXT   TAXON = 109 CHARACTER = 52 TEXT = 'secon bone appear to be anamestic - so suborbital';

	TEXT   CHARACTER = 53 TEXT = 'Taverne, 1997; Friedman & Blom, 2006; Long et al., 2008; Swartz, 2009; Choo, 2011; Giles et al., 2015b; Giles et al., 2017.';
	TEXT   TAXON = 68 CHARACTER = 53 TEXT = 'changed from - to 0. tube-like sensu Bürgin 1992';
	SU  T = 68 C = 53 N = color I = 5;

	TEXT   CHARACTER = 54 TEXT = 'Cloutier & Arratia, 2004; Friedman & Blom, 2006; Long et al., 2008; Swartz, 2009; Choo, 2011; Giles et al., 2015b; Giles et al., 2017; Xu et al., 2014. ';
	TEXT   TAXON = 74 CHARACTER = 54 TEXT = ?_to_0;
	SU  T = 74 C = 54 N = color I = 5;

	TEXT   CHARACTER = 55 TEXT = 'Gardiner & Schaeffer 1989; Olsen & McCune 1991; Taverne, 1997; Schultze & Cumbaa, 2001; Friedman & Blom, 2006; Long et al., 2008; Choo, 2011; López-Arbarello 2012; Xu & Gao 2011; Bermúdez-Rochas & Poyato-Ariza 2015; Poyato-Ariza 2015; Giles et al., 2015b; Thies & Waschkewitz, 2015; Xu et al., 2015; Gibson 2016; Xu & Zhao, 2016; Giles et al., 2017; López-Arbarello & Sferco 2018.';
	TEXT   TAXON = 35 CHARACTER = 55 TEXT = fragmented;
	TEXT   TAXON = 44 CHARACTER = 55 TEXT = coded_separately._;

	TEXT   CHARACTER = 56 TEXT = 'Giles et al., 2015b,c; Giles et al., 2017. Multiple branches radiate from the infraorbital canal in the jugal of many Carboniferous and younger actinopterygians.';

	TEXT   CHARACTER = 57 TEXT = 'Gardiner & Schaeffer, 1989; Coates, 1998; Schultze & Cumbaa, 2001; Cloutier & Arratia, 2004; Friedman & Blom, 2006; Zhu et al., 2006; Long et al., 2008; Zhu et al., 2009; Choo, 2011; Giles et al., 2015b; Xu et al., 2015; Giles et al., 2017. The dermosphenotic of Moythomasia illustrated by (Gardiner 1984, fig 69) lacks a posterior limb, but this is from a small individual and most likely reflects ontogenetic variability, with a posterior limb being developed in larger individuals (B. Choo, pers. comm.; Choo 2015: fig. 8). The posterior limb of the dermosphenotic is variably developed in Mesopoma (Coates, 1999), so this taxon is scored ''0/1'' to reflect this polymorphism.';
	TEXT   TAXON = 7 CHARACTER = 57 TEXT = no_distinct_dsphen;

	TEXT   CHARACTER = 58 TEXT = 'Friedman & Blom, 2006; Choo, 2011; Giles et al., 2015b; Giles et al., 2017.';

	TEXT   CHARACTER = 59 TEXT = 'Gardiner & Schaeffer 1989; Grande & Bemis 1998; Hurley et al. 2007; Xu & Gao 2011; Xu et al. 2014; Poyato-Ariza 2015; Xu et al., 2015; Xu & Zhao, 2016; Giles et al., 2017; López-Arbarello & Sferco 2018.';
	TEXT   TAXON = 117 CHARACTER = 59 TEXT = 'at least two are visible on the right side of the type specimen, but the complete series cannot be reconstructed';
	SU  T = 117 C = 59 N = color I = 5;

	TEXT   CHARACTER = 60 TEXT = 'Olsen & McCune 1991; Cavin & Suteethorn 2006; López-Arbarello 2012; Bermúdez-Rochas & Poyato-Ariza 2015; Thies & Waschkewitz, 2015; Gibson 2016; Giles et al., 2017; López-Arbarello & Sferco 2018.';

	TEXT   CHARACTER = 61 TEXT = 'Grande 2010; Xu & Wu 2012; Xu et al. 2014, 2015; Bermúdez-Rochas & Poyato-Ariza 2015; Poyato-Ariza 2015; Thies & Waschkewitz, 2015; Gibson 2016; Xu & Zhao, 2016; Giles et al., 2017.  The first lachrymal is regarded here as the anteriormost canal-bearing bone that contributes to orbital margin.';

	TEXT   CHARACTER = 62 TEXT = 'Wiley 1976; Lopez-Arbarello 2010.Wiley 1976; López-Arbarello 2012; Bermúdez-Rochas & Poyato-Ariza 2015; Poyato-Ariza 2015; Thies & Waschkewitz, 2015; Gibson 2016; Giles et al., 2017; López-Arbarello & Sferco 2018.';

	TEXT   CHARACTER = 63 TEXT = 'Patterson, 1982; Lauder & Liem, 1983; Gardiner, 1984; Cloutier & Arratia, 2004; Brazeau, 2009; Friedman & Brazeau, 2010; Choo, 2011; Davis et al., 2012; Zhu et al., 2013; Giles et al., 2015b,c; Giles et al., 2017.';

	TEXT   CHARACTER = 64 TEXT = 'Patterson, 1982; Gardiner & Schaeffer, 1989; Lund et al., 1995; Cloutier & Ahlberg, 1996; Coates, 1998; Dietze, 2000; Lund, 2000; Schultze & Cumbaa, 2001; Zhu & Schultze, 2001; Zhu et al., 2001; Lund & Poplin, 2002; Zhu & Yu, 2002; Cloutier & Arratia, 2004; Gardiner et al., 2005; Friedman & Blom, 2006; Zhu et al., 2006; Friedman, 2007; Long et al., 2008; Swartz, 2009; Zhu et al., 2009; Choo, 2011; Xu & Gao, 2011; Xu et al., 2014; 2015; Giles et al., 2015b; Xu & Zhao, 2016; Giles et al., 2017. This region of the cheek is missing in Coccocephalichthys (Poplin, 1974; Poplin & Véran, 1996), and was presumably removed by Watson (1925) when he first described the specimen. It is unclear from the surviving cast whether a dermohyal and/or accessory operculum were present, and as such this taxon is coded as ???.';
	TEXT   TAXON = 10 CHARACTER = 64 TEXT = 'changed from 0 to 1. Traces of the dermohyal can be seen in Pl. 18, 20 in Nielsen 1949. The position of this dermal flange agrees with the observations of Romano 2007 and Romano and Brinkmann 2009 on the dermohyal of Birgeria stensioei';
	SU  T = 10 C = 64 N = color I = 5;

	TEXT   CHARACTER = 65 TEXT = 'Giles et al., 2015b,c; Giles et al., 2017. The dermohyal is not preserved in Melanecta (Coates, 1998), but it is clear from the surrounding bones that it would not have projected above the dorsal surface of the operculum.';
	TEXT   TAXON = 10 CHARACTER = 65 TEXT = 'changed from - to ?, as this element is not well-preserved in the material figured by Nielsen (1949)';
	SU  T = 10 C = 65 N = color I = 5;
	TEXT   TAXON = 27 CHARACTER = 65 TEXT = Or_posterior_suborbital...?_;
	TEXT   TAXON = 74 CHARACTER = 65 TEXT = ?_to_0;
	SU  T = 74 C = 65 N = color I = 5;

	TEXT   CHARACTER = 66 TEXT = 'Gardiner et al., 2005; Coates, 1999; Xu & Gao, 2011; Xu et al., 2014; Giles et al., 2015b; Giles et al., 2017.';
	TEXT   TAXON = 10 CHARACTER = 66 TEXT = 'changed from - to 1, as this element appears to not be fused in the material figured by Nielsen 1949';
	SU  T = 10 C = 66 N = color I = 5;
	TEXT   TAXON = 95 CHARACTER = 66 TEXT = 'changed from 1 to ?, not known in this taxon';
	SU  T = 95 C = 66 N = color I = 5;

	TEXT   CHARACTER = 67 TEXT = 'Friedman, 2007; Zhu et al., 2009; Giles et al., 2015b; Giles et al., 2017.';

	TEXT   CHARACTER = 68 TEXT = 'Zhu & Yu, 2002; Friedman, 2007; Xu et al., 2014, 2015; Giles et al., 2015b; Xu & Zhao, 2016; Giles et al., 2017; López-Arbarello & Sferco 2018.';
	TEXT   TAXON = 88 CHARACTER = 68 TEXT = see_Grande_et_al._2002._The_bone_historically_attributed_as_the_maxilla_is_better_considered_a_dermopalatine_as_it_ensheaths_the_palatoquadrate;

	TEXT   CHARACTER = 69 TEXT = 'Lund et al., 1995; Lund, 2000; Poplin & Lund, 2000; Schultze & Cumbaa, 2001; Zhu & Schultze, 2001; Zhu et al., 2001; Zhu & Yu, 2002; Lund & Poplin, 2002; Cloutier & Arratia, 2004; Zhu et al., 2006; Friedman, 2007; Zhu et al., 2009; Zhu et al., 2013; Giles et al., 2015b,c; Giles et al., 2017; López-Arbarello & Sferco 2018. ';
	TEXT   TAXON = 86 CHARACTER = 69 TEXT = 'recoded from 0 to 1, sensu figs 136, 137 in Bürgin 1992';
	SU  T = 86 C = 69 N = color I = 5;
	TEXT   TAXON = 88 CHARACTER = 69 TEXT = 'recoded from absent to non-applicable';
	SU  T = 88 C = 69 N = color I = 5;

	TEXT   CHARACTER = 70 TEXT = 'Friedman, 2007; Zhu et al., 2009; Zhu et al., 2013; Giles et al., 2015b,c; Giles et al., 2017; López-Arbarello & Sferco 2018.';
	TEXT   TAXON = 10 CHARACTER = 70 TEXT = 'changed from 0 to 1, as a quadratojugal is not present in the material figured by Nielsen 1949';
	SU  T = 10 C = 70 N = color I = 5;

	TEXT   CHARACTER = 71 TEXT = 'Grande & Bemis 1998; Friedman, 2007; Zhu et al., 2009; Zhu et al., 2013; Giles et al., 2015b; Giles et al., 2017.';
	SU  T = 74 C = 71 N = color I = 5;

	TEXT   CHARACTER = 72 TEXT = 'Poyato-Ariza & Wenz, 2002; Cloutier & Arratia, 2004; López-Arbarello 2012; Xu et al. 2014, 2015; Bermúdez-Rochas & Poyato-Ariza 2015; Poyato-Ariza 2015; Thies & Waschkewitz, 2015; Gibson 2016; Xu & Zhao, 2016; Giles et al., 2017; Cawley & Kriwet 2018; López-Arbarello & Sferco 2018.';
	TEXT   TAXON = 5 CHARACTER = 72 TEXT = 'Changed from 1 to 0 following Sallan and Coates 2013: ''Amphicentrum, another superficially edentulous fish, has a field of apparent tooth cusps on the mesial maxilla, well dorsal to the sharpened edge of the gape''';
	SU  T = 5 C = 72 N = color I = 5;
	SU  T = 74 C = 72 N = color I = 5;

	TEXT   CHARACTER = 73 TEXT = 'Patterson 1973; Gardiner & Schaeffer 1989; Olsen & McCune 1991; Gardiner et al. 1996; Gardiner et al. 2005; Coates 1999; Hurley et al. 2007; Xu & Gao 2011; Poyato-Ariza 2015; Xu et al. 2014, 2015; Xu & Zhao, 2016; Giles et al., 2017.';

	TEXT   CHARACTER = 74 TEXT = 'Grande 2010; Xu & Wu 2012; Xu et al. 2014; Poyato-Ariza 2015; Giles et al., 2017.';

	TEXT   CHARACTER = 75 TEXT = 'Grande & Bemis 1998; Xu & Wu 2012; Xu et al. 2014, 2015. Arratia 2013; Xu & Zhao, 2016; Giles et al., 2017; López-Arbarello & Sferco 2018.';

	TEXT   CHARACTER = 76 TEXT = 'Grande & Bemis 1998; Gardiner & Schaeffer 1989; Olsen & McCune 1991; Gardiner et al. 1996; Gardiner et al. 2005; Coates 1999; Hurley et al. 2007; Xu & Gao 2011; Xu et al. 2014, 2015; Bermúdez-Rochas & Poyato-Ariza 2015; Poyato-Ariza 2015; Xu & Shen, 2015; Thies & Waschkewitz, 2015; Gibson 2016; Xu & Zhao, 2016; Giles et al., 2017; López-Arbarello & Sferco 2018.';
	TEXT   TAXON = 31 CHARACTER = 76 TEXT = 'maybe ? 	';
	TEXT   TAXON = 88 CHARACTER = 76 TEXT = 'recoded from - to 0';
	SU  T = 88 C = 76 N = color I = 5;

	TEXT   CHARACTER = 77 TEXT = 'Friedman & Blom, 2006; Long et al., 2008; Swartz, 2009; Choo, 2011; Giles et al., 2015b; Giles et al., 2017.';
	TEXT   TAXON = 31 CHARACTER = 77 TEXT = '0	';
	TEXT   TAXON = 74 CHARACTER = 77 TEXT = ?_to_0;
	SU  T = 74 C = 77 N = color I = 5;

	TEXT   CHARACTER = 78 TEXT = 'Giles et al., 2015b,c; Giles et al., 2017.';

	TEXT   CHARACTER = 79 TEXT = 'Patterson, 1982; Cloutier & Ahlberg, 1996; Coates, 1998; Schultze & Cumbaa, 2001; Zhu & Schultze, 2001; Zhu et al., 2001; Zhu & Yu, 2002; Cloutier & Arratia, 2004; Zhu et al., 2006; Friedman, 2007; Zhu et al., 2009; Choo, 2011; Zhu et al., 2013; Giles et al., 2015b; Giles et al., 2017.';

	TEXT   CHARACTER = 80 TEXT = 'Ahlberg & Johanson, 1998; Zhu et al., 2001; Zhu & Yu, 2002; Zhu & Ahlberg, 2004; Friedman, 2007; Zhu et al., 2009; Giles et al., 2015b; Giles et al., 2017.';
	TEXT   TAXON = 112 CHARACTER = 80 TEXT = roughly_half;

	TEXT   CHARACTER = 81 TEXT = 'Cloutier & Arratia 2004, Xu et al. 2014; Giles et al., 2017; López-Arbarello & Sferco 2018.';
	TEXT   TAXON = 88 CHARACTER = 81 TEXT = present_only_early_in_ontogeny;

	TEXT   CHARACTER = 82 TEXT = 'Friedman & Blom, 2006; Long et al., 2008; Swartz, 2009; Choo, 2011; Giles et al., 2015b; Giles et al., 2017.';

	TEXT   CHARACTER = 83 TEXT = 'Friedman & Blom, 2006; Long et al., 2008; Swartz, 2009; Choo, 2011; Giles et al., 2017.';
	TEXT   TAXON = 29 CHARACTER = 83 TEXT = Single_enlarged_tooth_at_anterior_end_of_dentary;

	TEXT   CHARACTER = 84 TEXT = 'Choo, 2011; Giles et al., 2015b; Giles et al., 2017.';

	TEXT   CHARACTER = 85 TEXT = 'Friedman, 2007; Giles et al., 2015b; Poyato-Ariza 2015; Giles et al., 2017; López-Arbarello & Sferco 2018. Coates (1998) states that the maxilla of Melanecta bears large teeth interspersed with smaller teeth, but it is unclear how these teeth are arranged. As such, this taxon is coded ''?''. ';
	TEXT   TAXON = 26 CHARACTER = 85 TEXT = monocuspid;
	TEXT   TAXON = 27 CHARACTER = 85 TEXT = monocuspid;
	TEXT   TAXON = 28 CHARACTER = 85 TEXT = bicuspid;
	TEXT   TAXON = 74 CHARACTER = 85 TEXT = ?_to_2;
	SU  T = 74 C = 85 N = color I = 5;

	TEXT   CHARACTER = 86 TEXT = 'Giles et al. 2017. In Styracopterus, Fouldenia and Amphicentrum, a lateral lamina of bone obscures the maxillary dentition (Sallan & Coates 2013).';

	TEXT   CHARACTER = 87 TEXT = 'Patterson, 1982; Gardiner, 1984; Maisey, 1986; Gardiner & Schaeffer, 1989; Cloutier & Ahlberg, 1996; Taverne, 1997; Coates, 1999; Poplin & Lund, 2000; Schultze & Cumbaa, 2001; Zhu & Schultze, 2001; Zhu et al., 2001; Zhu & Yu, 2002; Cloutier & Arratia, 2004; Gardiner et al., 2005; Friedman & Blom, 2006; Zhu et al., 2006; Friedman, 2007; Long et al., 2008; Zhu et al., 2009; Friedman & Brazeau, 2010; Choo, 2011; Xu & Gao, 2011; Zhu et al., 2013; Xu et al., 2014; Giles et al., 2015b,c; Giles et al., 2017. ';
	SU  T = 117 C = 87 N = color I = 16;

	TEXT   CHARACTER = 88 TEXT = 'Zhu & Yu 2002; Friedman 2007; López-Arbarello 2012; Bermúdez-Rochas & Poyato-Ariza 2015; Poyato-Ariza 2015; Thies & Waschkewitz, 2015; Gibson 2016; Giles et al., 2017; López-Arbarello & Sferco 2018.';
	SU  T = 117 C = 88 N = color I = 15;

	TEXT   CHARACTER = 89 TEXT = 'Friedman & Blom, 2006; Long et al., 2008; Swartz, 2009; Grande 2010; Choo, 2011; Xu et al., 2014; Giles et al., 2015b; Poyato-Ariza 2015; Giles et al., 2017.';
	TEXT   TAXON = 4 CHARACTER = 89 TEXT = nb_three_ossifications;
	TEXT   TAXON = 64 CHARACTER = 89 TEXT = nb_two_ossifications;

	TEXT   CHARACTER = 90 TEXT = 'Friedman & Blom, 2006; Friedman, 2007; Long et al., 2008; Choo, 2011; Xu & Gao, 2011; Xu et al., 2014; Giles et al., 2015b; Poyato-Ariza 2015; Giles et al., 2017; López-Arbarello & Sferco 2018.';
	TEXT   TAXON = 74 CHARACTER = 90 TEXT = ?_to_1;
	SU  T = 74 C = 90 N = color I = 5;

	TEXT   CHARACTER = 91 TEXT = 'Schultze and Cumbaa, 2001; Zhu and Schultze, 2001; Zhu et al., 2001; Zhu and Yu, 2002; Zhu et al., 2006; Friedman, 2007; Zhu et al., 2009; Giles et al., 2015b,c; Poyato-Ariza 2015; Giles et al., 2017; López-Arbarello & Sferco 2018. ';
	TEXT   TAXON = 117 CHARACTER = 91 TEXT = changed_from_0_to_?;
	SU  T = 117 C = 91 N = color I = 5;

	TEXT   CHARACTER = 92 TEXT = 'Ahlberg & Clack, 1998; Daeschler et al., 2006; Long et al., 2006; Friedman, 2007; Zhu et al., 2009; Zhu et al., 2013; Giles et al., 2015b; Giles et al., 2017. A single specimen of Pteronisculus stensioi has at least five or six coronoids anterior to the prearticular region. However, these appear to correspond to the three coronoids present in most specimens, so the taxon is coded here as ''2''. Two coronoids are reported in Boreosomus (Nielsen, 1942).';
	TEXT   TAXON = 5 CHARACTER = 92 TEXT = fused;
	TEXT   TAXON = 106 CHARACTER = 92 TEXT = Hmm._The_one_figured_in_G&B_only_has_two._;

	TEXT   CHARACTER = 93 TEXT = 'Cloutier & Ahlberg, 1996; Ahlberg & Johanson, 1998; Zhu & Ahlberg, 2004; Daeschler et al., 2006; Long et al., 2006; Giles et al., 2015b; Giles et al., 2017.';

	TEXT   CHARACTER = 94 TEXT = 'Patterson 1973; Gardiner & Schaeffer 1989; Olsen & McCune 1991; Poyato-Ariza & Wenz, 2002; Zhu & Yu, 2002; Friedman, 2007; Friedman, 2007; Xu & Gao 2011; Xu et al. 2014, 2015; Giles et al., 2015b; Poyato-Ariza 2015; Xu & Zhao, 2016; Giles et al., 2017; Cawley & Kriwet 2018; Cawley & Kriwet 2018; López-Arbarello & Sferco 2018.';
	TEXT   TAXON = 74 CHARACTER = 94 TEXT = ?_to_1;
	SU  T = 74 C = 94 N = color I = 5;
	SU  T = 117 C = 94 N = color I = 16;

	TEXT   CHARACTER = 95 TEXT = 'Modified from Patterson 1973; Olsen & McCune 1991; Gardiner et al. 2005; Poyato-Ariza 2015; Giles et al., 2017; López-Arbarello & Sferco 2018.';
	TEXT   TAXON = 24 CHARACTER = 95 TEXT = Splenail_rather_than_dentary;
	TEXT   TAXON = 26 CHARACTER = 95 TEXT = Splenail_rather_than_dentary;
	TEXT   TAXON = 29 CHARACTER = 95 TEXT = Splenail_rather_than_dentary;
	TEXT   TAXON = 74 CHARACTER = 95 TEXT = ?_to_1;
	SU  T = 74 C = 95 N = color I = 5;

	TEXT   CHARACTER = 96 TEXT = 'Arratia 2013; Giles et al., 2017; López-Arbarello & Sferco 2018. A distinct notch in the posterior margin of the dentary is seen in taxa such as Leptolepis.';

	TEXT   CHARACTER = 97 TEXT = 'Grande & Bemis 1998; Olsen & McCune 1991; Grande 2010; Xu & Wu 2012; Xu et al. 2014, 2015; López-Arbarello 2012; Bermúdez-Rochas & Poyato-Ariza 2015; Poyato-Ariza 2015; Thies & Waschkewitz, 2015; Gibson 2016; Xu & Zhao, 2016; Giles et al., 2017; López-Arbarello & Sferco 2018.';
	TEXT   TAXON = 10 CHARACTER = 97 TEXT = 'Changed from 0 to ?, as Nielsen 1949 provides an interpretative drawing showing the presence of a symplectic articulating with the retroarticular process of the lower jaw';
	SU  T = 10 C = 97 N = color I = 5;
	TEXT   TAXON = 88 CHARACTER = 97 TEXT = 'condition similar to that in Acipenser, but the symplectic/interhyal remains cartilaginous';

	TEXT   CHARACTER = 98 TEXT = 'Friedman, 2007; Giles et al., 2015b; Giles et al., 2017.';

	TEXT   CHARACTER = 99 TEXT = 'Giles et al., 2017.';

	TEXT   CHARACTER = 100 TEXT = 'Revised from Friedman, 2007; Brazeau, 2009; Zhu et al., 2009; Friedman & Brazeau, 2010; Davis et al., 2012; Zhu et al., 2013; Giles et al., 2015b,c; Giles et al., 2017; López-Arbarello & Sferco 2018. This character is expanded from previous formulations, which only considered whether a basipterygoid fenestra was absent or present. Where a basipterygoid process is absent, the dorsal margin of the palate may be flat, or the metapterygoid may bear a distinct notch.';
	TEXT   TAXON = 7 CHARACTER = 100 TEXT = mpt_proc;
	TEXT   TAXON = 10 CHARACTER = 100 TEXT = 'changed from 2 to 0, as there is no obvious process in the palate of Birgeria (Nielsen 1949).';
	SU  T = 10 C = 100 N = color I = 5;

	TEXT   CHARACTER = 101 TEXT = 'Giles et al., 2017.)';
	TEXT   TAXON = 95 CHARACTER = 101 TEXT = 'changed from 0 to ?, as this feature is not well-preserved in this taxon';
	SU  T = 95 C = 101 N = color I = 5;
	TEXT   TAXON = 117 CHARACTER = 101 TEXT = changed_from_0_to_?;
	SU  T = 117 C = 101 N = color I = 15;

	TEXT   CHARACTER = 102 TEXT = 'Giles et al., 2017. This character captures whether the palatoquadrates contact at the midline. ';

	TEXT   CHARACTER = 103 TEXT = 'Giles et al., 2017.';

	TEXT   CHARACTER = 104 TEXT = 'Giles et al., 2017.';

	TEXT   CHARACTER = 105 TEXT = 'Friedman, 2007; Giles et al., 2015b; Giles et al., 2017.';

	TEXT   CHARACTER = 106 TEXT = 'Poyato-Ariza 2015; Giles et al., 2017; López-Arbarello & Sferco 2018.';

	TEXT   CHARACTER = 107 TEXT = 'Patterson 1973; Olsen & McCune 1991; López-Arbarello 2012; Arratia 2013; Xu & Wu, 2012; Bermúdez-Rochas & Poyato-Ariza 2015; Poyato-Ariza 2015; Thies & Waschkewitz, 2015; Gibson 2016; Xu & Zhao, 2016; Giles et al., 2017; López-Arbarello & Sferco 2018.';
	TEXT   TAXON = 56 CHARACTER = 107 TEXT = vomers_fuse_ontogenetically;
	TEXT   TAXON = 106 CHARACTER = 107 TEXT = sutured_to_parasphenoid._Check_in_Grande_&_Bemis._;

	TEXT   CHARACTER = 108 TEXT = 'Patterson 1973; Olsen & McCune 1991; Hurley et al. 2007; Giles et al., 2017.';

	TEXT   CHARACTER = 109 TEXT = 'Schultze & Cumbaa, 2001; Cloutier & Arratia, 2004; Friedman & Blom, 2006; Long et al., 2008; Swartz, 2009; Giles et al., 2015b; Giles et al., 2017. This region of the cheek was removed in Coccocephalichthys (Poplin, 1974; Poplin & Véran, 1996), presumably by Watson (1925) when he first described the specimen. It is unclear from the surviving cast whether a dermohyal and/or accessory operculum were present, and as such this taxon is coded as ???.';
	TEXT   TAXON = 110 CHARACTER = 109 TEXT = changed_from_0_to_?;

	SU  T = 1 C = 110 N = color I = 5;

	TEXT   CHARACTER = 111 TEXT = 'Modified from Lund et al., 1995; Lund, 2000; Lund & Poplin, 2002; Cloutier & Arratia, 2004; Long et al., 2008; Swartz, 2009; Choo, 2011; López-Arbarello 2012; Bermúdez-Rochas & Poyato-Ariza 2015; Xu et al., 2015; Giles et al., 2015b; Thies & Waschkewitz, 2015; Gibson 2016; Xu & Zhao, 2016; Giles et al., 2017; López-Arbarello & Sferco 2018.';
	TEXT   TAXON = 1 CHARACTER = 111 TEXT = 'changed from ? to -';
	SU  T = 1 C = 111 N = color I = 5;
	TEXT   TAXON = 2 CHARACTER = 111 TEXT = changed_from_2_to_0;
	SU  T = 2 C = 111 N = color I = 5;
	TEXT   TAXON = 10 CHARACTER = 111 TEXT = 'changed from 1 to ?, the structure identified by Nielsen (1949) as the subopercle corresponds to the subopercle and branchiostegal rays (Romano and Brinkmann 2009)';
	SU  T = 10 C = 111 N = color I = 5;
	TEXT   TAXON = 85 CHARACTER = 111 TEXT = 'changed from ? to -';
	SU  T = 85 C = 111 N = color I = 5;
	TEXT   TAXON = 95 CHARACTER = 111 TEXT = 'changed from - to 2, following Argyriou et al 2018';
	SU  T = 95 C = 111 N = color I = 5;

	TEXT   CHARACTER = 112 TEXT = 'Poyato-Ariza & Wenz, 2002; Xu et al. 2014; Giles et al., 2017; Cawley & Kriwet 2018.';
	TEXT   TAXON = 1 CHARACTER = 112 TEXT = 'changed from ? to -';
	SU  T = 1 C = 112 N = color I = 5;
	TEXT   TAXON = 10 CHARACTER = 112 TEXT = multiple_suboperc;
	TEXT   TAXON = 95 CHARACTER = 112 TEXT = 'changed from 1 to 0, following Argyriou et al 2018';
	SU  T = 95 C = 112 N = color I = 5;
	TEXT   TAXON = 106 CHARACTER = 112 TEXT = 'changed from 1 to 0, following Olsen 1984';
	SU  T = 106 C = 112 N = color I = 5;
	TEXT   TAXON = 110 CHARACTER = 112 TEXT = 'although the expanded gill covering bone likely corresponds to a subopercle as in other saurichthyiforms, we prefer to recode it as uncertain, given the lack of an associated hyomandibula (see Wu et al. 2013)';
	SU  T = 110 C = 112 N = color I = 5;

	TEXT   CHARACTER = 113 TEXT = 'Long et al., 2008; Choo, 2011; López-Arbarello 2012; Bermúdez-Rochas & Poyato-Ariza 2015; Giles et al., 2015b; Thies & Waschkewitz, 2015; Poyato-Ariza 2015; Gibson 2016; Giles et al., 2017; López-Arbarello & Sferco 2018. An anterodorsal process is present some non-neopterygian actinopterygians and ''semionotiforms''. Although these processes do not appear to be homologous they are coded within the character.';
	TEXT   TAXON = 1 CHARACTER = 113 TEXT = 'changed from ? to -';
	SU  T = 1 C = 113 N = color I = 5;
	TEXT   TAXON = 85 CHARACTER = 113 TEXT = 'changed from ? to -';
	SU  T = 85 C = 113 N = color I = 5;
	TEXT   TAXON = 95 CHARACTER = 113 TEXT = 'changed from - to 0, following Argyriou et al 2018';
	SU  T = 95 C = 113 N = color I = 5;
	TEXT   TAXON = 110 CHARACTER = 113 TEXT = 'although the expanded gill covering bone likely corresponds to a subopercle as in other saurichthyiforms, we prefer to recode it as uncertain, given the lack of an associated hyomandibula (see Wu et al. 2013)';
	SU  T = 110 C = 113 N = color I = 5;
	TEXT   TAXON = 118 CHARACTER = 113 TEXT = process_absent_from_original_description_but_present_in_fossil;
	TEXT   TAXON = 123 CHARACTER = 113 TEXT = process_absent_from_original_description_but_present_in_fossil;

	TEXT   CHARACTER = 114 TEXT = 'Giles et al., 2017.';
	TEXT   TAXON = 1 CHARACTER = 114 TEXT = 'changed from ? to -';
	SU  T = 1 C = 114 N = color I = 5;
	TEXT   TAXON = 95 CHARACTER = 114 TEXT = 'changed from - to 0, following Argyriou et al 2018';
	SU  T = 95 C = 114 N = color I = 5;
	TEXT   TAXON = 110 CHARACTER = 114 TEXT = 'although the expanded gill covering bone likely corresponds to a subopercle as in other saurichthyiforms, we prefer to recode it as uncertain, given the lack of an associated hyomandibula (see Wu et al. 2013)';
	SU  T = 110 C = 114 N = color I = 5;

	TEXT   CHARACTER = 115 TEXT = 'Friedman, 2007; Zhu et al., 2009; Zhu et al., 2013; Giles et al., 2015b; Xu & Zhao, 2016; Giles et al., 2017.';
	TEXT   TAXON = 2 CHARACTER = 115 TEXT = lots_of_small_ossicles...;
	TEXT   TAXON = 26 CHARACTER = 115 TEXT = loads!_suborbs_carry_it..._;
	TEXT   TAXON = 111 CHARACTER = 115 TEXT = 'Apparently two. But dorsal pop might be infraorbital? Can''t tell. 	';

	TEXT   CHARACTER = 116 TEXT = 'Modified from Patterson 1973; Olsen & McCune 1991; Grande & Bemis 1998; Gardiner et al., 2005; Swartz 2009; López-Arbarello 2012; Bermúdez-Rochas & Poyato-Ariza 2015; Thies & Waschkewitz, 2015; Poyato-Ariza 2015; Gibson 2016; Giles et al., 2017; López-Arbarello & Sferco 2018. This character is reformulated from its original (compound) formulation, which considered both maxilla and preoperculum shape. Primitively in actinopterygians the preoperculum is wholly or partially developed dorsal to the maxilla as an anterodorsal-posteroventrally oriented bone, either with (e.g. Mimipiscis, Moythomasia) or without (e.g. Cheirolepis) a dorsoventrally oriented limb. The preoperculum may also be near-vertical, with no distinct anterodorsal or anteroventral extensions (e.g. Boreosomus, Peltopleurus), or developed as an anteroventral-posterodorsally-directed bone largely ventral to the maxilla (e.g. Discoserra, Propterus).';
	TEXT   TAXON = 112 CHARACTER = 116 TEXT = hmm._or_state_2;

	TEXT   CHARACTER = 117 TEXT = 'Modified from López-Arbarello 2012; Bermúdez-Rochas & Poyato-Ariza 2015; Poyato-Ariza 2015; Thies & Waschkewitz, 2015; Gibson 2016; Giles et al., 2017; López-Arbarello & Sferco 2018.';

	TEXT   CHARACTER = 118 TEXT = 'López-Arbarello 2012; Bermúdez-Rochas & Poyato-Ariza 2015; Giles et al., 2017; Thies & Waschkewitz, 2015; Gibson 2016; López-Arbarello & Sferco 2018.';

	TEXT   CHARACTER = 119 TEXT = 'Patterson 1973; Grande & Bemis 1998; Gardiner & Schaeffer 1989; Olsen & McCune 1991; Poyato-Ariza & Wenz, 2002; Xu & Gao 2011; Xu et al. 2014, 2015. Gardiner & Schaeffer 1989; Olsen & McCune 1991; Gardiner et al. 1996; Gardiner et al. 2005; Cavin & Suteethorn 2006; Hurley et al. 2007; López-Arbarello 2012; Bermúdez-Rochas & Poyato-Ariza 2015; Poyato-Ariza 2015; Thies & Waschkewitz, 2015; Gibson 2016; Xu & Zhao, 2016; Giles et al., 2017; Cawley & Kriwet 2018; López-Arbarello & Sferco 2018. ';
	TEXT   TAXON = 96 CHARACTER = 119 TEXT = changed_from_0_to_?;
	SU  T = 96 C = 119 N = color I = 5;

	TEXT   CHARACTER = 120 TEXT = 'Lund et al., 1995; Cloutier & Arratia, 2004; Choo, 2011; Giles et al., 2015b; Giles et al., 2017.';

	TEXT   CHARACTER = 121 TEXT = 'Olsen & McCune 1991; Xu et al., 2014; Giles et al., 2017. ';

	TEXT   CHARACTER = 122 TEXT = 'Gardiner & Schaeffer, 1989; Cloutier & Ahlberg, 1996; Taverne, 1997; Lund & Poplin, 1997; Coates, 1999; Schultze & Cumbaa, 2001; Zhu & Schultze, 2001; Cloutier & Arratia, 2004; Friedman & Blom, 2006; Long et al., 2008; Swartz, 2009; Brazeau, 2009; Xu & Gao, 2011; Davis et al., 2012; Zhu et al., 2013; Xu et al., 2014; Giles et al., 2015b,c; Giles et al., 2017. The condition in Boreosomus (Nielsen, 1942) is unique: instead of lateral gulars flanking a median gular, there appears to be a second median gular. This may well represent a fusion of the two, longer lateral gulars, is coded as such.';
	TEXT   TAXON = 74 CHARACTER = 122 TEXT = ?_to_1;
	SU  T = 74 C = 122 N = color I = 5;

	TEXT   CHARACTER = 123 TEXT = 'Olsen & McCune 1991; Lund et al., 1995; Cloutier & Ahlberg, 1996; Coates, 1999; Lund, 2000; Schultze & Cumbaa, 2001; Zhu & Schultze, 2001; Zhu et al., 2001: Lund & Poplin, 2002; Zhu & Yu, 2002; Cloutier & Arratia, 2004; Zhu et al., 2006; Friedman, 2007; Zhu et al., 2009; Xu & Gao, 2011; López-Arbarello 2012; Zhu et al., 2013; Bermúdez-Rochas & Poyato-Ariza 2015; Poyato-Ariza 2015; Xu et al., 2014, 2015; Giles et al., 2015b,c; Thies & Waschkewitz, 2015; Xu & Zhao, 2015; Gibson 2016; Giles et al., 2017; López-Arbarello & Sferco 2018. Pearson & Westoll (1979: p. 365) state that a median gular is not known in Cheirolepis canadensis. Although a median gular is reconstructed by Cloutier & Arratia (1996: fig. 7), this bone is not present in any specimen photos and is not mentioned in the text. As such, this taxon is coded as ???.';
	TEXT   TAXON = 117 CHARACTER = 123 TEXT = changed_from_0_to_1;
	SU  T = 117 C = 123 N = color I = 5;

	TEXT   CHARACTER = 124 TEXT = 'Giles et al., 2015b,c; Giles et al., 2017. The condition in Boreosomus (Nielsen, 1942) is unique: instead of lateral gulars flanking a median gular, there appears to be a second median gular. This may well represent a fusion of the two, longer lateral gulars, and is coded as such.';

	TEXT   CHARACTER = 125 TEXT = 'Schultze & Cumbaa, 2001; Zhu & Schultze, 2001; Zhu et al., 2001; Zhu & Yu, 2002; Zhu et al., 2006; Friedman, 2007; Zhu et al., 2009; Zhu et al., 2013; Giles et al., 2015b; Giles et al., 2017. ';

	TEXT   CHARACTER = 126 TEXT = 'Zhu & Yu, 2002; Friedman, 2007; Giles et al., 2015b; Giles et al., 2017. Character can only be coded in taxa which possess all of these bones.';

	TEXT   CHARACTER = 127 TEXT = 'Ahlberg & Johanson, 1998; Zhu & Yu, 2002; Zhu & Ahlberg, 2004; Daeschler et al., 2006; Long et al., 2006; Friedman, 2007; Zhu et al., 2009; Zhu et al., 2013; Giles et al., 2015b,c; Giles et al., 2017.';

	TEXT   CHARACTER = 128 TEXT = 'Friedman, 2007; Zhu et al., 2009; Brazeau, 2009; Friedman & Brazeau, 2010; Davis et al., 2012; Zhu et al., 2013; Giles et al., 2015b,c; Giles et al., 2017. ';
	TEXT   TAXON = 2 CHARACTER = 128 TEXT = 'changed from 1 to 0, following our investigation of the neurocranium of Acipenser (Argyriou et al 2018)';
	SU  T = 2 C = 128 N = color I = 5;
	SU  T = 117 C = 128 N = color I = 5;

	TEXT   CHARACTER = 129 TEXT = 'Giles et al., 2017. Primitively in actinopts, the optic nerve exits the cranial cavity into the orbit through paired foramina approximately halfway up the orbital wall. In many Carboniferous and younger taxa, much of the orbital wall is unossified (the optic fenestra). The optic nerves may exit through openings just posterior to (e.g. Pteronisculus) or confluent with (e.g. Pholidophorus) the optic fenestra. In polypterids and Fukangichthys, the optic nerve exits ventrally through paired foramina that abut the parasphenoid.';

	TEXT   CHARACTER = 130 TEXT = 'Giles et al., 2015b,c; Giles et al., 2017. Carboniferous and younger actinopts such as Lawrenciella (Hamel & Poplin, 2008) have a median crista anterior to the anterior dorsal fontanelle upon which the skull roof sits.';

	TEXT   CHARACTER = 131 TEXT = 'Giles et al., 2015b,c; Giles et al., 2017. The anterior dorsal fontanelle of many Carboniferous and younger actinopts is greatly expanded, in contrast to the smaller fontanelle of Devonian taxa such as Mimipiscis (Gardiner, 1984).';
	TEXT   TAXON = 95 CHARACTER = 131 TEXT = 'changed from 0 to ?, as the braincase is not well-known in S, madagascariensis';
	SU  T = 95 C = 131 N = color I = 5;

	TEXT   CHARACTER = 132 TEXT = 'Cloutier & Ahlberg, 1996; Ahlberg & Johanson, 1998; Zhu & Ahlberg, 2004; Zhu et al., 2001; Zhu & Yu, 2002; Daeschler et al., 2006; Long et al., 2006; Friedman, 2007; Brazeau, 2009; Zhu et al., 2009; Friedman & Brazeau, 2010; Davis et al., 2013; Zhu et al., 2013; Giles et al., 2015b,c; Giles et al., 2017.';
	SU  T = 117 C = 132 N = color I = 16;

	TEXT   CHARACTER = 133 TEXT = 'Zhu & Schultze, 2001; Zhu et al., 2001; Zhu & Yu, 2002; Zhu et al., 2006; Friedman, 2007; Zhu et al., 2009; Zhu et al., 2013; Giles et al., 2015b,c; Giles et al., 2017. This character is coded as absent for taxa that possess a large interorbital fenestra (e.g. Pteronisculus, Coccocephalichthys, Kentuckia deani), as, if present, the eyestalk area would be visible posterior to the opening for the optic nerve.';

	TEXT   CHARACTER = 134 TEXT = 'Coates, 1999; Giles et al., 2015b; Giles et al., 2017.';

	TEXT   CHARACTER = 135 TEXT = 'Coates, 1999; Giles et al., 2015b; Giles et al., 2017.';

	TEXT   CHARACTER = 136 TEXT = 'Patterson 1973; Olsen & McCune 1991; Coates, 1999; Gardiner et al. 1996; Hurley et al. 2007; Xu & Gao, 2011; Xu et al., 2014, 2015; Giles et al., 2015b; Poyato-Ariza 2015; Xu & Zhao, 2016; Giles et al., 2017.';
	TEXT   TAXON = 95 CHARACTER = 136 TEXT = 'changed from ? to 1. Fen 1 of Beltan 1968, Fig 12 is homologous to the anterodorsal myodome';
	SU  T = 95 C = 136 N = color I = 5;

	TEXT   CHARACTER = 137 TEXT = 'Modified from Patterson 1973; Olsen & McCune 1991; Coates, 1999; Xu et al., 2014. Wiley 1976; Gardiner, 1984; Gardiner & Schaeffer, 1989; Gardiner et al. 1996; Hurley et al. 2007; López-Arbarello 2012; Xu & Gao, 2011; Bermúdez-Rochas & Poyato-Ariza 2015; Xu et al. 2014, 2015; Giles et al., 2015b; Thies & Waschkewitz, 2015; Gibson 2016; Xu & Zhao, 2016; Giles et al., 2017; López-Arbarello & Sferco 2018.';
	SU  T = 117 C = 137 N = color I = 15;

	TEXT   CHARACTER = 138 TEXT = 'Ahlberg & Johanson, 1998; Zhu et al., 2001; Zhu & Yu, 2002; Zhu & Ahlberg, 2004; Friedman, 2007; Zhu et al., 2009; Zhu et al., 2013; Giles et al., 2015b,c; Giles et al., 2017. ';

	TEXT   CHARACTER = 139 TEXT = 'Modified from Patterson, 1982; Gardiner, 1984; Gardiner & Schaeffer, 1989; Taverne, 1997; Coates, 1999; Gardiner et al., 2005; Xu & Gao, 2011; Xu et al., 2014; Giles et al., 2015b; Giles et al., 2017; López-Arbarello & Sferco 2018. Taxa that lack a groove on the lateral commissure are coded as inapplicable for this character.) ';

	TEXT   CHARACTER = 140 TEXT = 'Gardiner et al., 2005; Xu & Gao, 2011; Xu et al., 2014, 2015; Giles et al., 2015b; Xu & Zhao, 2016; Giles et al., 2017; López-Arbarello & Sferco 2018.';
	TEXT   TAXON = 2 CHARACTER = 140 TEXT = 'changed from 0 to 1, as the basitrabecular process in the posteroventral ethmoidal region of acipenserids is not homologous to the basipterygoid process in the sphenoid region of other actinopterygians.';
	SU  T = 2 C = 140 N = color I = 5;
	TEXT   TAXON = 19 CHARACTER = 140 TEXT = 'changed from 0 to ?, braincase is unknown';
	SU  T = 19 C = 140 N = color I = 5;
	SU  T = 117 C = 140 N = color I = 15;
	TEXT   TAXON = 118 CHARACTER = 140 TEXT = 'basipterygoid processes absent in fossil, present in original description by Nielsen (1942)';
	TEXT   TAXON = 123 CHARACTER = 140 TEXT = 'basipterygoid processes absent in fossil, present in original description by Nielsen (1942)';

	TEXT   CHARACTER = 141 TEXT = 'Ahlberg & Johanson, 1998; Zhu & Schultze, 2001; Zhu et al., 2001; Zhu & Yu, 2002; Zhu & Ahlberg, 2004; Zhu et al., 2006; Friedman, 2007; Zhu et al., 2009; Davis et al., 2012; Zhu et al., 2013; Giles et al., 2015b,c; Giles et al., 2017.';
	TEXT   TAXON = 2 CHARACTER = 141 TEXT = 'changed from 0 to 1, as the basitrabecular process in the posteroventral ethmoidal region of acipenserids is not homologous to the basipterygoid process in the sphenoid region of other actinopterygians.';
	TEXT   TAXON = 19 CHARACTER = 141 TEXT = 'changed from 0 to ?, braincase is unknown';
	SU  T = 19 C = 141 N = color I = 5;
	TEXT   TAXON = 85 CHARACTER = 141 TEXT = '? to -';
	SU  T = 85 C = 141 N = color I = 5;
	SU  T = 117 C = 141 N = color I = 15;
	TEXT   TAXON = 118 CHARACTER = 141 TEXT = 'basipterygoid processes absent in fossil, present in original description by Nielsen (1942)';
	TEXT   TAXON = 123 CHARACTER = 141 TEXT = 'basipterygoid processes absent in fossil, present in original description by Nielsen (1942)';

	TEXT   CHARACTER = 142 TEXT = 'Gardiner, 1984; Gardiner & Schaeffer, 1989; Taverne, 1997; Coates, 1999; Giles et al., 2015b; Giles et al., 2017.';
	TEXT   TAXON = 85 CHARACTER = 142 TEXT = '? to -';
	SU  T = 85 C = 142 N = color I = 5;
	TEXT   TAXON = 88 CHARACTER = 142 TEXT = 'changed from - to 0';
	SU  T = 88 C = 142 N = color I = 5;
	TEXT   TAXON = 96 CHARACTER = 142 TEXT = 'changed from - to 0';
	SU  T = 96 C = 142 N = color I = 5;
	TEXT   TAXON = 97 CHARACTER = 142 TEXT = 'changed from - to 0';
	SU  T = 97 C = 142 N = color I = 5;
	TEXT   TAXON = 110 CHARACTER = 142 TEXT = 'changed from - to 0';
	SU  T = 110 C = 142 N = color I = 5;
	TEXT   TAXON = 118 CHARACTER = 142 TEXT = 'basipterygoid processes absent in fossil, present in original description by Nielsen (1942)';
	TEXT   TAXON = 123 CHARACTER = 142 TEXT = 'basipterygoid processes absent in fossil, present in original description by Nielsen (1942)';

	TEXT   CHARACTER = 143 TEXT = 'Gardiner et al. 2005; Gardiner et al. 1996; Hurley et al. 2007; Xu & Gao 2011; Xu et al. 2015; Giles et al., 2017; López-Arbarello & Sferco 2018.	';

	TEXT   CHARACTER = 144 TEXT = 'Gardiner, 1984; Gardiner & Schaeffer, 1989; Taverne, 1997; Coates, 1999; Xu & Gao, 2011; Xu et al., 2014; Giles et al., 2015b; Giles et al., 2017.';
	TEXT   TAXON = 117 CHARACTER = 144 TEXT = changed_from_0_to_?;
	SU  T = 117 C = 144 N = color I = 5;

	TEXT   CHARACTER = 145 TEXT = 'Zhu & Yu, 2002; Friedman, 2007.';

	TEXT   CHARACTER = 146 TEXT = 'Friedman, 2007; Brazeau, 2009; Zhu et al., 2009; Friedman & Brazeau, 2010; Davis et al., 2012; Zhu et al., 2013; Brazeau & Friedman, 2014; Giles et al., 2015b,c; Giles et al., 2017.';

	TEXT   CHARACTER = 147 TEXT = 'Coates, 1999; Giles et al., 2015b; Giles et al., 2017. We follow Coates (1999) in coding Howqualepis as ''0'' on the basis of Long 1988 fig. 16 and AMF65495 (pers. obs. S.G.), rather than the braincase reconstruction (Long, 1988: fig. 18.';
	TEXT   TAXON = 88 CHARACTER = 147 TEXT = 'changed from ? to -';
	SU  T = 88 C = 147 N = color I = 5;

	TEXT   CHARACTER = 148 TEXT = 'Friedman, 2007; Zhu et al., 2009; Giles et al., 2015b; Giles et al., 2017.';

	TEXT   CHARACTER = 149 TEXT = 'Friedman, 2007; Brazeau, 2009; Davis et al., 2012; Zhu et al., 2013; Giles et al., 2015b,c; Giles et al., 2017.';
	TEXT   TAXON = 106 CHARACTER = 149 TEXT = check_this_coding..._;

	TEXT   CHARACTER = 150 TEXT = 'Giles et al., 2015b; Giles et al., 2017. Variable in Boreosomus: the posterior dorsal fontanelle is closed in the specimen figured in Nielsen (1942: plate 25F), but developed in the specimen figured in plate 28. This taxon is coded ?0/1? to reflect this polymorphism.';

	TEXT   CHARACTER = 151 TEXT = 'Giles et al., 2015b; Giles et al., 2017. This projection is somewhat reduced in Pteronisculus (Nielsen, 1942), but is coded ''1'' here. Variable in Boreosomus: the posterior dorsal fontanelle is closed in the specimen figured in Nielsen 1942 plate 25F, but developed in the specimen figured in plate 28. This taxon is coded ?0/1? to reflect this polymorphism. Cheirolepis trailli is coded ''0'' (2015a).';

	TEXT   TAXON = 25 CHARACTER = 152 TEXT = 'A small process pierced by anastomosing occipital nerves is present (Latimer and Giles 2018), which could correspond to a reduced craniospinal process';
	TEXT   TAXON = 53 CHARACTER = 152 TEXT = 'changed from 1 to 0, miscoded in Argyriou et al. 2018';
	SU  T = 53 C = 152 N = color I = 5;
	SU  T = 117 C = 152 N = color I = 5;

	TEXT   CHARACTER = 153 TEXT = 'Coates & Sequeira, 1998; Coates & Sequeira, 2001a, b; Coates, 1999; Friedman, 2007; Zhu et al., 2009; Friedman & Brazeau, 2010; Zhu et al., 2013; Giles et al., 2015b,c; Giles et al., 2017; López-Arbarello & Sferco 2018. This character is coded as inapplicable in taxa that lack a canal for the dorsal aorta. Cheirolepis trailli is coded ''0'' (Giles et al., 2015a). The aortic canal of Moythomasia is not figured by Gardiner (1984), but a clear posterior notch in the aortic canal can be seen in Long & Trinajstic (2010:fig 5b). The neurocranium of Gogosardina is not yet described, but this character can be coded on the basis of Choo et al. (2009: fig. 9).';
	TEXT   TAXON = 28 CHARACTER = 153 TEXT = 'v hard to interpret: Thies 1999 fig 2. So ? ';
	TEXT   TAXON = 43 CHARACTER = 153 TEXT = 'Rayner says canal, Patetrson says no canal...';
	TEXT   TAXON = 67 CHARACTER = 153 TEXT = Draw_in_notebook!;
	TEXT   TAXON = 89 CHARACTER = 153 TEXT = changed_from_?_to_2;
	SU  T = 89 C = 153 N = color I = 5;

	TEXT   CHARACTER = 154 TEXT = 'Giles et al., 2015b; Giles et al., 2017. In Mimipiscis bartrami and M. toombsi, a groove for one of the efferent branchial arteries branches off from the lateral dorsal aorta immediately before the articular area for the first infrapharyngobranchial. However, it is uncertain which, so both taxa coded as ''?'' for these characters. The neurocranium of Gogosardina is not yet described, but this character can be coded on the basis of Choo et al. (2009: fig. 9).';
	TEXT   TAXON = 65 CHARACTER = 154 TEXT = 'Recoded from 1 to 0. Based on Rayner 1937, the anterior opening of the aortic canal leads to a keel on the parasphenoid, which indicates that it is the lateral dorsal aortae that exited the aortic canal at this point. The overal basicranial circulation pattern matches that of other neopterygians and teleosts figured in Patterson 1975.';
	SU  T = 65 C = 154 N = color I = 5;
	TEXT   TAXON = 89 CHARACTER = 154 TEXT = changed_from_?_to_0;
	SU  T = 89 C = 154 N = color I = 5;

	TEXT   CHARACTER = 155 TEXT = 'Giles et al., 2015b; Giles et al., 2017. In Mimipiscis bartrami and M. toombsi, a groove for one of the efferent branchial arteries branches off from the lateral dorsal aorta immediately before the articular area for the first infrapharyngobranchial. However, it is uncertain which, so both taxa coded as ''?'' for these characters. The neurocranium of Gogosardina is not yet described, but this character can be coded on the basis of Choo et al. (2009: fig. 9).';
	TEXT   TAXON = 89 CHARACTER = 155 TEXT = changed_from_?_to_0;
	SU  T = 89 C = 155 N = color I = 5;

	TEXT   CHARACTER = 156 TEXT = 'Coates & Sequeira, 1998; Coates & Sequeira, 2001a, b; Coates, 1999; Friedman, 2007; Zhu et al., 2009; Friedman & Brazeau, 2010; Zhu et al., 2013; Giles et al., 2015b,c; Giles et al., 2017.';
	TEXT   TAXON = 89 CHARACTER = 156 TEXT = changed_from_?_to_1;
	SU  T = 89 C = 156 N = color I = 5;

	TEXT   CHARACTER = 157 TEXT = 'Coates, 1999; Giles et al., 2015b; Giles et al., 2017. In Mimipiscis toombsi, the bifurcation point of the dorsal aorta can be open (Giles & Friedman, 2014: fig. 2) or closed (Gardiner 1984: fig. 15). This taxon is coded ''0/1'' to reflect this polymorphism. The aortic canal of Moythomasia is not figured by Gardiner (1984), but the bifucation into the lateral dorsal aortae can be seen in Long & Trinajstic (2010:fig 5b). We modified this character (sensu Argyriou et al. 2018) to capture the variation in actinopterygians taxa that lack an aortic canal (e.g., Saurichthys), or taxa that have an aortic canal, but whose dorsal aortae or common carotids bifurcate and extend ventral to the parasphenoid (e.g., Brachydegma). This character is coded as inapplicable in taxa that host both dorsal aorta and lateral dorsal aortae in endoskeletal grooves. In Mimipiscis toombsi, the bifurcation point of the dorsal aorta can be open (Giles & Friedman, 2014: fig. 2) or closed (Gardiner 1984: fig. 15). This taxon is coded ''0/1'' to reflect this polymorphism. The aortic canal of Moythomasia is not figured by Gardiner (1984), but the bifucation into the lateral dorsal aortae can be seen in Long & Trinajstic (2010:fig 5b). In Acipenser there are variably present foramina for the efferent branchial arteries (Marinelli and Strenger 1973; Hilton et al. 2011), which are here accounted for similarly to the lateral dorsal aortae or common carotids, and are coded as polymorphic 1/2.';
	TEXT   TAXON = 2 CHARACTER = 157 TEXT = 'changed from - to 1/2 following Marinelli and Strenger 1973 and Argyriou et al. 2018';
	SU  T = 2 C = 157 N = color I = 5;
	TEXT   TAXON = 4 CHARACTER = 157 TEXT = 'changed from 0 to 2 following the addition of the new character state (Argyriou et al. 2018)';
	SU  T = 4 C = 157 N = color I = 5;
	SU  T = 6 C = 157 N = color I = 5;
	TEXT   TAXON = 10 CHARACTER = 157 TEXT = changed_from_?_to_2_given_the_absence_of_an_aortic_canal_and_the_presence_of_an_elongate_parasphenoidean_stalk;
	SU  T = 10 C = 157 N = color I = 5;
	TEXT   TAXON = 45 CHARACTER = 157 TEXT = 0_to_2;
	SU  T = 45 C = 157 N = color I = 5;
	SU  T = 58 C = 157 N = color I = 5;
	SU  T = 59 C = 157 N = color I = 5;
	TEXT   TAXON = 64 CHARACTER = 157 TEXT = changed_from_0_to_2_following_the_addition_of_the_new_character_state;
	SU  T = 64 C = 157 N = color I = 5;
	SU  T = 65 C = 157 N = color I = 5;
	TEXT   TAXON = 81 CHARACTER = 157 TEXT = changed_from_0_to_2_following_the_addition_of_the_new_character_state;
	SU  T = 81 C = 157 N = color I = 5;
	TEXT   TAXON = 89 CHARACTER = 157 TEXT = changed_from_?_to_1;
	SU  T = 89 C = 157 N = color I = 5;
	SU  T = 106 C = 157 N = color I = 5;
	SU  T = 117 C = 157 N = color I = 5;

	TEXT   CHARACTER = 158 TEXT = 'Giles et al., 2017.';
	TEXT   TAXON = 24 CHARACTER = 158 TEXT = intercalar_only_differentiated;
	TEXT   TAXON = 27 CHARACTER = 158 TEXT = intercalar_only_differentiated;
	TEXT   TAXON = 118 CHARACTER = 158 TEXT = 'juvenile fossil specimen has braincase ossification in different "areas" rather than separate bones (occiput-orbitotemporal-ethmoidal-basisphenoid)';
	TEXT   TAXON = 123 CHARACTER = 158 TEXT = 'juvenile fossil specimen has braincase ossification in different "areas" rather than separate bones (occiput-orbitotemporal-ethmoidal-basisphenoid)';

	TEXT   CHARACTER = 159 TEXT = 'Wiley 1976; López-Arbarello 2012; Bermúdez-Rochas & Poyato-Ariza 2015; Thies & Waschkewitz, 2015; Gibson 2016; Giles et al., 2017; López-Arbarello & Sferco 2018.';
	SU  T = 117 C = 159 N = color I = 15;

	TEXT   CHARACTER = 160 TEXT = 'Gardiner et al. 1996; Hurley et al. 2007; Giles et al., 2017. This character can only be coded when separate braincase ossifications can be identified.';

	TEXT   CHARACTER = 161 TEXT = 'Hurley et al. 2007; Giles et al., 2017. This character can only be coded when separate braincase ossifications can be identified.';

	TEXT   CHARACTER = 162 TEXT = 'Olsen & McCune 1991; Gardiner et al. 1996; Cavin & Suteethorn 2006; Hurley et al. 2007; López-Arbarello 2012; Bermúdez-Rochas & Poyato-Ariza 2015; Thies & Waschkewitz, 2015; Gibson 2016; Giles et al. 2017; López-Arbarello & Sferco 2018. This character can only be coded when separate braincase ossifications can be identified.';

	TEXT   CHARACTER = 163 TEXT = 'Grande 2010; López-Arbarello 2012; Xu & Wu 2012; Xu et al. 2014, 2015; Arratia 2013; Bermúdez-Rochas & Poyato-Ariza 2015; Poyato-Ariza 2015; Thies & Waschkewitz, 2015; Gibson 2016; Xu & Zhao, 2016; Giles et al. 2017; López-Arbarello & Sferco 2018. This character can only be coded when separate braincase ossifications can be identified.';

	TEXT   CHARACTER = 164 TEXT = 'Gardiner et al. 1996; Grande & Bemis, 1998; Hurley et al. 2007; Xu et al. 2014; Poyato-Ariza 2015; Xu & Shen, 2015; Xu & Zhao, 2016; Giles et al. 2017; López-Arbarello & Sferco 2018. This character can only be coded when separate braincase ossifications can be identified.';

	TEXT   CHARACTER = 165 TEXT = 'Wiley 1976; Olsen & McCune 1991; Grande & Bemis 1998; Cavin & Suteethorn 2006; Hurley et al. 2007; Grande 2010; López-Arbarello 2012; Bermúdez-Rochas & Poyato-Ariza 2015; Poyato-Ariza 2015; Thies & Waschkewitz, 2015; Xu et al., 2014, 2015; Gibson 2016; Xu & Zhao, 2016; Giles et al. 2017; López-Arbarello & Sferco 2018. This character can only be coded when separate braincase ossifications can be identified.';

	TEXT   CHARACTER = 166 TEXT = 'Olsen & McCune 1991; Olsen 1994; Gardiner et al. 1996; López-Arbarello 2012; Xu et al. 2014; Poyato-Ariza 2015; Xu & Shen, 2015; Thies & Waschkewitz, 2015; Gibson 2016; Xu & Zhao, 2016; Giles et al. 2017; López-Arbarello & Sferco 2018. This character can only be coded when separate braincase ossifications can be identified. ';
	TEXT   TAXON = 118 CHARACTER = 166 TEXT = 'intercalar present in fossil NHMD-75388-all other codings changed to "?" to reflect this';

	TEXT   CHARACTER = 167 TEXT = 'This character (Argyriou et al. 2018) applies only to taxa with differentiated braincase ossifications. Acipenseriforms are coded as ? due to the putative homologization of the occipital perichondral ossification with a compound exoccipital-epioccipital (see Hilton et al. 2011). The occipital region is co-mineralized in polypterids (e.g., Allis 1922).';

	TEXT   CHARACTER = 168 TEXT = 'Grande 2010; Poyato-Ariza 2015; Xu et al. 2014, 2015; Xu & Zhao, 2016; Giles et al. 2017; López-Arbarello & Sferco 2018. This character can only be coded when separate braincase ossifications can be identified. ';
	SU  T = 117 C = 168 N = color I = 5;

	TEXT   CHARACTER = 169 TEXT = 'Patterson 1973; Olsen & McCune 1991; Gardiner et al. 1996; Hurley et al. 2007; Giles et al. 2017; López-Arbarello & Sferco 2018. This character can only be coded when separate braincase ossifications can be identified. ';
	TEXT   TAXON = 118 CHARACTER = 169 TEXT = 'intercalar present in fossil NHMD-75388-all other codings changed to "?" to reflect this';

	TEXT   CHARACTER = 170 TEXT = 'Patterson 1973; Gardiner, 1984; Gardiner & Schaeffer, 1989; Olsen & McCune 1991; Coates, 1999; Hurley et al. 2007; López-Arbarello 2012; Xu & Gao, 2011; Xu et al. 2014, 2015; Bermúdez-Rochas & Poyato-Ariza 2015; Giles et al., 2015b; Poyato-Ariza 2015; Thies & Waschkewitz, 2015; Gibson 2016; Xu & Zhao, 2016; Giles et al. 2017.';

	TEXT   CHARACTER = 171 TEXT = 'Gardiner, 1984; Gardiner & Schaeffer, 1989; Gardiner et al. 1996; Hurley et al. 2007; Xu & Gao, 2011; Xu et al. 2014, 2015; Xu & Zhao, 2016.';

	TEXT   CHARACTER = 172 TEXT = 'Gardiner, 1984; Gardiner & Schaeffer, 1989; Coates, 1999; Gardiner et al. 1996; Hurley et al. 2007; Xu & Gao, 2011; Xu et al. 2014, 2015; Xu & Zhao, 2016; Giles et al. 2017.';
	TEXT   TAXON = 41 CHARACTER = 172 TEXT = Unclear_where_hmd_articulates_so_not_sure._Also_why_foramen?;

	TEXT   CHARACTER = 173 TEXT = 'Gardiner, 1984; Brazeau, 2009; Davis et al., 2012; Zhu et al., 2013; Giles et al., 2015b,c; Giles et al. 2017.';

	TEXT   CHARACTER = 174 TEXT = 'Coates, 1999; Zhu & Yu, 2002; Gardiner et al., 2005; Friedman, 2007, Xu & Gao, 2011; Xu et al., 2014, 2015; Giles et al., 2015b; Xu & Zhao, 2016; Giles et al. 2017; López-Arbarello & Sferco 2018.';
	TEXT   TAXON = 99 CHARACTER = 174 TEXT = changed_from_1_to_?_as_the_posterior_boundary_of_the_endochondral_braincase_is_not_visible_in_illustrated_specimens;
	SU  T = 99 C = 174 N = color I = 5;
	TEXT   TAXON = 117 CHARACTER = 174 TEXT = changed_from_1_to_2;
	SU  T = 117 C = 174 N = color I = 5;

	TEXT   CHARACTER = 175 TEXT = 'Patterson, 1982; Coates, 1999; Dietze, 2000; Schultze & Cumbaa, 2001; Zhu & Schultze, 2001; Cloutier & Arratia, 2004; Gardiner et al., 2005; Friedman & Blom, 2006; Zhu et al., 2006; Zhu et al., 2009; Choo, 2011; Xu & Gao, 2011; Zhu et al., 2013; Xu et al., 2014; Giles et al., 2015b,c; Giles et al. 2017; López-Arbarello & Sferco 2018.';

	TEXT   CHARACTER = 176 TEXT = 'Friedman & Blom, 2006; Friedman, 2007; Zhu et al., 2009; Choo, 2011; Zhu et al., 2013; Giles et al., 2015b,c; Giles et al. 2017.';

	TEXT   CHARACTER = 177 TEXT = 'Giles et al., 2015b,c; Giles et al. 2017. The buccohypophyseal canal typically enters the dorsal surface of the parasphenoid, but whether it exits via the ventral surface is more variable, and this distribution is captured by this character.';
	TEXT   TAXON = 5 CHARACTER = 177 TEXT = 'nb median myodome means this is not trye bhc, but mmaybe an anterior extension';
	TEXT   TAXON = 117 CHARACTER = 177 TEXT = scan_quality_does_not_allow_us_to_securely_assess_this_feature;
	SU  T = 117 C = 177 N = color I = 5;
	TEXT   TAXON = 118 CHARACTER = 177 TEXT = prasphenoid_pierced_by_canal_in_fossil_NHMD_73588;

	TEXT   CHARACTER = 178 TEXT = 'Arratia 2013; Giles et al. 2017; López-Arbarello & Sferco 2018.';

	SU  T = 4 C = 179 N = color I = 5;
	SU  T = 114 C = 179 N = color I = 5;

	TEXT   CHARACTER = 180 TEXT = 'Gardiner et al. 1996; Hurley et al. 2007; Xu & Wu, 2012; Xu et al., 2015; Xu & Zhao, 2016; Giles et al. 2017.';
	TEXT   TAXON = 118 CHARACTER = 180 TEXT = 'paired foramina on orsal surface, near the base of the ascending processes. Canals from the foramina then join the buccohypophyseal canal';

	TEXT   CHARACTER = 181 TEXT = 'Gardiner et al. 1996; Hurley et al. 2007; Xu & Wu, 2012; Xu et al., 2015; Xu & Zhao, 2016; Giles et al. 2017.) ';

	TEXT   CHARACTER = 182 TEXT = 'Modified from Gardiner et al. 2005; Giles et al. 2017.';

	TEXT   CHARACTER = 183 TEXT = 'Xu and Gao 2011; Xu et al. 2014; Giles et al. 2017.';
	TEXT   TAXON = 10 CHARACTER = 183 TEXT = 'changed from 1 to ?, as this feature has not been investigated';
	SU  T = 10 C = 183 N = color I = 5;
	TEXT   TAXON = 25 CHARACTER = 183 TEXT = '0	';

	TEXT   CHARACTER = 184 TEXT = 'Coates, 1999; Giles & Friedman, 2014; Giles et al., 2015b; Giles et al. 2017.';

	TEXT   CHARACTER = 185 TEXT = 'Brazeau, 2009; Friedman & Brazeau 2010; Davis et al., 2012; Zhu et al., 2013; Brazeau & Friedman, 2014; Giles & Friedman, 2014; Giles et al., 2015b,c; Giles et al. 2017. The olfactory tracts of Osorioichthys are elongate (pers. obs. unpubl. scan data S.G.).';

	TEXT   CHARACTER = 186 TEXT = 'Coates, 1999; Giles & Friedman, 2014; Giles et al., 2015b; Giles et al. 2017. The olfactory nerves are carried in separate tracts in Osorioichthys (pers. obs. unpubl. scan data S.G.).';
	TEXT   TAXON = 6 CHARACTER = 186 TEXT = 'changed from 0 to 1, based on PIMUZ A/I 4171a';
	SU  T = 6 C = 186 N = color I = 5;
	TEXT   TAXON = 67 CHARACTER = 186 TEXT = changed_from_?_to_0._Described_as_median_by_Schaeffer_and_Dalquest_1978;
	SU  T = 67 C = 186 N = color I = 5;
	TEXT   TAXON = 74 CHARACTER = 186 TEXT = revised_from_?_to_0;
	SU  T = 74 C = 186 N = color I = 5;

	TEXT   CHARACTER = 188 TEXT = 'Coates, 1999; Xu & Gao, 2011; Xu et al., 2014; Giles et al., 2015b; Giles et al. 2017.';
	TEXT   TAXON = 13 CHARACTER = 188 TEXT = changed_from_1_to_?_as_it_is_still_unknown_in_Boreosomus;
	SU  T = 13 C = 188 N = color I = 5;

	TEXT   CHARACTER = 189 TEXT = 'Giles & Friedman, 2014; Giles et al., 2015b; Giles et al. 2017.';

	TEXT   CHARACTER = 190 TEXT = 'Coates 1999; Hurley et al. 2007; Xu et al. 2014; Giles et al. 2017. ';

	TEXT   CHARACTER = 191 TEXT = 'Coates, 1999; Giles et al., 2015b; Giles et al. 2017.';
	TEXT   TAXON = 95 CHARACTER = 191 TEXT = changed_from_1_to_?._The_endocast_is_very_poorly_known;
	SU  T = 95 C = 191 N = color I = 5;

	TEXT   CHARACTER = 192 TEXT = 'Giles et al., 2015b,c; Giles et al. 2017. The region posterior to the cerebellar auricles in Lawrenciella was considered to be the area octavolateralis by Hamel & Poplin (2005). However, we interpret it as the corpus cerebellum, and this taxon is coded ''1''.';

	TEXT   CHARACTER = 193 TEXT = 'Coates 1999; Hurley et al. 2007; Xu & Gao 2011; Xu et al. 2014; Giles et al. 2017. ';

	TEXT   CHARACTER = 194 TEXT = 'Coates 1999; Hurley et al. 2007; Xu & Gao 2011; Xu et al. 2014; Giles et al. 2017. ';

	TEXT   CHARACTER = 195 TEXT = 'Coates 1999; Hurley et al. 2007; Xu & Gao 2011; Xu et al. 2014; Giles et al. 2017. ';

	TEXT   CHARACTER = 196 TEXT = 'Davis et al., 2012; Zhu et al., 2013; Giles & Friedman, 2014; Giles et al., 2015b,c; Giles et al. 2017.';
	TEXT   TAXON = 2 CHARACTER = 196 TEXT = 'changed from ? to 1 based on observations on endocast (Argyriou et al. 2018)';
	SU  T = 2 C = 196 N = color I = 5;

	TEXT   CHARACTER = 197 TEXT = 'Giles et al., 2015b,c; Giles et al. 2017. In certain stem actinopterygians, such as Mimipiscis (Giles and Friedman, 2014), a short length of canal lies between the posterior ampulla and the remainder of the labyrinth.';
	TEXT   TAXON = 2 CHARACTER = 197 TEXT = 'changed from ? to 0 based on observations on endocast (Argyriou et al. 2018';
	SU  T = 2 C = 197 N = color I = 5;

	TEXT   CHARACTER = 198 TEXT = 'Giles & Friedman, 2014; Giles et al., 2015b; Giles et al. 2017.';
	TEXT   TAXON = 2 CHARACTER = 198 TEXT = 'changed from ? to 1 based on observations on endocast (Argyriou et al. 2018)';
	SU  T = 2 C = 198 N = color I = 5;
	TEXT   TAXON = 96 CHARACTER = 198 TEXT = 'changed from ? to 0, since crural superimposition is clearly seen in Argyriou et al 2018';
	SU  T = 96 C = 198 N = color I = 5;

	TEXT   CHARACTER = 199 TEXT = 'Gardiner, 1984; Gardiner & Schaeffer, 1989; Coates, 1999; Cloutier & Arratia, 2004; Gardiner et al., 2005; Zhu et al., 2006; Zhu et al., 2009; Zhu et al., 2013; Giles & Friedman, 2014; Xu et al., 2014; Giles et al., 2015b,c; Giles et al. 2017. The presence of a lateral cranial canal in Ligulalepis and Psarolepis is uncertain, but its presence in Meemannia is confirmed following Lu et al. (2016). Erpetoichthys is conservatively coded as ''?''. ';
	TEXT   TAXON = 2 CHARACTER = 199 TEXT = 'changed from ? to 0 based on observations on endocast (Argyriou et al. 2018';
	SU  T = 2 C = 199 N = color I = 5;

	TEXT   CHARACTER = 200 TEXT = Giles_et_al._2017.;
	TEXT   TAXON = 2 CHARACTER = 200 TEXT = 'changed from ? to 1 based on observations on endocast (Argyriou et al. 2018)';
	SU  T = 2 C = 200 N = color I = 5;
	TEXT   TAXON = 118 CHARACTER = 200 TEXT = 'juvenile specimen has two openings: posterior one as usual and anterior one below anterior semicircular canal. Juvenile feature only, so coded "0"';

	TEXT   TAXON = 24 CHARACTER = 201 TEXT = 'changed from - to 0';
	SU  T = 24 C = 201 N = color I = 5;
	TEXT   TAXON = 62 CHARACTER = 201 TEXT = changed_from_0_to_?;
	SU  T = 62 C = 201 N = color I = 5;
	TEXT   TAXON = 94 CHARACTER = 201 TEXT = changed_from_1_to_?;
	SU  T = 94 C = 201 N = color I = 5;

	TEXT   TAXON = 4 CHARACTER = 202 TEXT = 'changed from - to 0';
	TEXT   TAXON = 7 CHARACTER = 202 TEXT = 'changed from - to 0';
	SU  T = 7 C = 202 N = color I = 5;
	TEXT   TAXON = 24 CHARACTER = 202 TEXT = changed_from_0_to_?;
	SU  T = 24 C = 202 N = color I = 5;
	TEXT   TAXON = 47 CHARACTER = 202 TEXT = 'changed from ? to -';
	SU  T = 47 C = 202 N = color I = 5;

	TEXT   CHARACTER = 203 TEXT = 'Giles et al., 2015b,c, 2017:Characters 147-150 form part of an atomisation of the compound characters ''ganoine'' (typically defined as a single or multilayer enamel covering) and ''cosmine'' (typically defined as a single layer of enamel with a well-defined pore canal network) (e.g. Cloutier & Ahlberg, 1996; Ahlberg & Johanson, 1998; Zhu & Ahlberg, 2004; Schultze & Cumbaa, 2001; Zhu & Schultze, 2001; Zhu et al., 2001; Zhu & Yu, 2002; Daeschler et al., 2006; Long et al., 2006; Zhu et al., 2006; Zhu et al., 2009; Davis et al., 2012; Zhu et al., 2013). A similar approach to atomization was taken by Friedman (2007), Brazeau & Friedman (2010) and Giles et al. (2015b). As detailed histological investigations have not been carried out for the majority of early actinopterygians (rather, they have simply been described as being covered in/bearing ridges of ganoine), many of these characters cannot be coded for a number of taxa.';

	TEXT   CHARACTER = 204 TEXT = See_notes_above_for_c._147.;

	TEXT   CHARACTER = 205 TEXT = See_notes_above_for_c._147.;

	TEXT   CHARACTER = 206 TEXT = See_notes_above_for_c._147.;

	TEXT   CHARACTER = 207 TEXT = 'Giles et al. 2017.)';
	TEXT   TAXON = 2 CHARACTER = 207 TEXT = 'changed from - to 0 to account for the scutes';
	SU  T = 2 C = 207 N = color I = 5;
	TEXT   TAXON = 10 CHARACTER = 207 TEXT = 'changed from 1 to 0, see Nielsen 1949: Plate 18, fig. 2';
	SU  T = 10 C = 207 N = color I = 5;
	TEXT   TAXON = 95 CHARACTER = 207 TEXT = reduced;

	TEXT   CHARACTER = 208 TEXT = 'Cloutier & Arratia, 2004; Friedman & Blom, 2006; Long et al., 2008; Swartz, 2009; Zhu et al., 2009; Choo, 2011; Giles et al., 2015b; Giles et al. 2017.';
	TEXT   TAXON = 2 CHARACTER = 208 TEXT = 'changed from - to 1';
	SU  T = 2 C = 208 N = color I = 5;

	TEXT   CHARACTER = 209 TEXT = 'Maisey, 1986; Gardiner & Schaeffer, 1989; Cloutier & Ahlberg, 1996; Coates, 1999; Dietze, 2000; Poplin & Lund, 2000; Schultze & Cumbaa, 2001; Cloutier & Arratia, 2004; Friedman & Blom, 2006; Friedman, 2007; Long et al., 2008; Brazeau, 2009; Swartz, 2009; Zhu et al., 2009; Friedman & Brazeau, 2010; López-Arbarello 2012; Xu & Gao, 2011; Choo, 2011; Davis et al., 2012; Zhu et al., 2013; Xu et al., 2014; Giles et al., 2015b,c; Xu & Zhao, 2016; Giles et al. 2017; López-Arbarello & Sferco 2018. This character is coded only for taxa that possess rhombic scales.';
	TEXT   TAXON = 2 CHARACTER = 209 TEXT = 'changed from - to 0 to account for scutes and caudal scales';
	SU  T = 2 C = 209 N = color I = 5;
	TEXT   TAXON = 10 CHARACTER = 209 TEXT = 'changed from ? to 0, as the lateral line scales lack this feature';
	SU  T = 10 C = 209 N = color I = 5;

	TEXT   CHARACTER = 210 TEXT = 'Patterson, 1982; Cloutier & Ahlberg, 1996; Dietze, 2000; Schultze & Cumbaa, 2001; Zhu & Schultze, 2001; Zhu et al., 2001: Zhu & Yu, 2002; Cloutier & Arratia, 2004; Friedman & Blom, 2006; Zhu et al., 2006; Friedman, 2007; Zhu et al., 2009; Giles et al. 2017. Although peg-and-socket articulation of is present between the scales of Limnomis (Daeschler, 2000), the nature of the peg is not described. As such, this taxon is conservatively coded ''?''.';

	TEXT   CHARACTER = 211 TEXT = 'Patterson, 1982; Gardiner, 1984; Gardiner & Schaeffer, 1989; Schultze & Cumbaa, 2001; Zhu & Schultze, 2001; Zhu et al., 2001; Zhu & Yu, 2002; Cloutier & Arratia, 2004; Friedman & Blom, 2006; Zhu et al., 2006; Friedman, 2007; Long et al., 2008; Swartz, 2009; Zhu et al., 2009; Choo, 2011; Zhu et al., 2013; Giles et al., 2015b,c; Giles et al. 2017.';
	TEXT   TAXON = 2 CHARACTER = 211 TEXT = 'changed from - to 0';
	SU  T = 2 C = 211 N = color I = 5;
	TEXT   TAXON = 10 CHARACTER = 211 TEXT = changed_from_?_to_0;
	SU  T = 10 C = 211 N = color I = 5;

	TEXT   CHARACTER = 212 TEXT = 'Friedman & Blom 2006; Long et al., 2008; Swartz, 2009; Choo, 2011; Xu et al., 2014; Giles et al., 2015b; Giles et al. 2017. Scales from the posterior half of the flank in Wendichthys bear pores on the enamel surface, whereas those from the anterior part of the flank lack these pores (Lund & Poplin, 1997: fig. 6). This taxon is scored ''1''.';
	TEXT   TAXON = 2 CHARACTER = 212 TEXT = 'changed from - to 0';
	SU  T = 2 C = 212 N = color I = 5;

	TEXT   CHARACTER = 213 TEXT = Giles_et_al._2017.;
	TEXT   TAXON = 74 CHARACTER = 213 TEXT = check;

	TEXT   CHARACTER = 214 TEXT = 'Friedman, 2007; Brazeau, 2009; Zhu et al., 2009; Friedman & Brazeau, 2010; Davis et al., 2012; Zhu et al., 2013; Brazeau & Friedman, 2014; Giles et al., 2015b,c; Giles et al. 2017. ';

	TEXT   CHARACTER = 215 TEXT = 'Patterson, 1982; Gardiner & Schaeffer, 1989; Grande & Bemis 1998; Coates, 1999; Dietze, 2000; Schultze & Cumbaa, 2001; Poyato-Ariza & Wenz, 2002; Cloutier & Arratia, 2004; Friedman & Blom, 2006; Friedman, 2007; Long et al., 2008; Swartz, 2009; Zhu et al., 2009; Choo, 2011; Xu & Gao, 2011; López-Arbarello 2012; Zhu et al., 2013; Xu et al., 2014, 2015; Bermúdez-Rochas & Poyato-Ariza 2015; Poyato-Ariza 2015; Giles et al., 2015b; Thies & Waschkewitz, 2015; Gibson 2016; Xu & Zhao, 2016; Giles et al. 2017; Cawley & Kriwet 2018; López-Arbarello & Sferco 2018.';
	TEXT   TAXON = 2 CHARACTER = 215 TEXT = changed_from_0_to_1_following_Hilton_et_al._2011;
	SU  T = 2 C = 215 N = color I = 5;

	TEXT   CHARACTER = 216 TEXT = 'Cloutier & Ahlberg, 1996; Zhu & Schultze, 2001; Schultze & Cumbaa, 2001; Zhu et al., 2001; Zhu & Yu, 2002; Zhu et al., 2006; Friedman, 2007; Zhu et al., 2009; Friedman & Brazeau, 2010; Zhu et al., 2013; Giles et al., 2015b,c; Giles et al. 2017; López-Arbarello & Sferco 2018.';

	TEXT   CHARACTER = 217 TEXT = 'Friedman, 2007; Zhu et al., 2009; Friedman & Brazeau, 2010; Xu & Gao, 2011; Zhu et al., 2013; Brazeau & Friedman, 2014; Xu et al., 2014, 2015; Giles et al., 2015b; Xu & Zhao, 2016; Giles et al. 2017. Although Long (1988: p.24) mentions the presence of a depression for the hyomandibular nerve in Howqualepis, it is unclear whether this perforated the hyomandibula. This taxon is conservatively coded as ''?''.';

	TEXT   CHARACTER = 218 TEXT = 'Gardiner & Schaeffer, 1989 Poyato-Ariza & Wenz, 2002;; Giles et al., 2015b; Giles et al. 2017; Cawley & Kriwet 2018; López-Arbarello & Sferco 2018.';

	TEXT   CHARACTER = 219 TEXT = 'Gardiner et al., 2005; Xu & Gao, 2011; Giles et al., 2015b; Giles et al. 2017; Xu et al., 2014.';
	TEXT   TAXON = 2 CHARACTER = 219 TEXT = changed_from_1_to_0_following_our_reinterpretation_of_accessory_hyoidean_elements_in_chondrosteans;
	SU  T = 2 C = 219 N = color I = 5;
	TEXT   TAXON = 10 CHARACTER = 219 TEXT = changed_from_0_to_?_as_the_ventral_part_of_the_hyoid_arch_is_not_completely_preserved_in_the_material_figured_by_Nielsen_1949;
	SU  T = 10 C = 219 N = color I = 5;

	TEXT   CHARACTER = 220 TEXT = 'Revised from Coates, 1999; Giles et al., 2015b; Giles et al. 2017; López-Arbarello & Sferco 2018.The character captures whether the ceratohyal (or the anterior ossification if an anterior and posterior ceratohyal are present) is medially constricted (hourglass-shaped) or plate-like in lateral view.';

	TEXT   CHARACTER = 221 TEXT = 'Coates, 1999; Giles et al., 2015b; Giles et al. 2017. The groove for the afferent hyoidean artery in the ceratohyal of Gogosardina is visible in Choo 2009 (fig 6).';
	TEXT   TAXON = 51 CHARACTER = 221 TEXT = 'Changed from 0 to 1, see Giles et al. 2017 extended figure 5';
	SU  T = 51 C = 221 N = color I = 5;
	TEXT   TAXON = 117 CHARACTER = 221 TEXT = changed_from_0_to_1;
	SU  T = 117 C = 221 N = color I = 5;

	TEXT   CHARACTER = 222 TEXT = 'Davis et al., 2012; Zhu et al., 2013; Giles et al., 2015b,c; Giles et al. 2017.';
	TEXT   TAXON = 88 CHARACTER = 222 TEXT = 'the two accessory hyoidean elements remain cartilaginous (Grande and Bemis 1991)';

	TEXT   CHARACTER = 223 TEXT = 'The homology of some of the constituents of the hyoid arch?the so-called intermediate or accessory hyoid elements?in modern ray-fins remains controversial. The history attached to naming these elements is very complex (see Patterson 1973, Patterson 1982, Véran 1988, Gardiner et al. 1996, etc.), and we have tried here to apply a simple, consistent approach and code for the number of ossified accessory hyoidean elements alone. Polypterids exhibit a single ossified element connecting the hyomandibula with the ceratohyal, which has been identified as the primitively present interhyal (e.g., Allis 1922, Jollie 1984, Patterson 1982, Veran 1988). A second independently ossified element (a ?symplectic?) does not develop. Neopterygians (e.g. Amia, Lepisosteus, Hiodon, ?Dorsetichthys, ?Macrosemionotus, ?Watsonulus etc.) exhibit two intermediate hyoidean elements, a symplectic and an interhyal. The interhyal connects the hyomandibula with the ceratohyal. This may be very reduced (e.g. ?Watsonulus, Elops), or entirely cartilaginous (e.g. Amia, Lepisosteus) in many actinopts. The ossification that contacts the hyomandibula (and typically the quadrate and the articular), but does not articulate with the ceratohyal, is termed the symplectic. This element may brace the quadrate, and in ?Watsonulus, ?Caturus, Amia and possibly pycnodonts additionally forms an articulation with the lower jaw. Although the presence of a true symplectic in stem actinopterygians (?paleoniscoids?) has been rejected by many authors(e.g, Patterson 1982, Gardiner and Schaeffer 1989, Gardiner 1996), Véran (1988) identified a second, symplectic-like, intermediate ossification in the hyoid arch in a number of ?palaeoniscoids?. Our reexamination of part of her material (?Boreosomus reuteskioldi) confirmed her observations of two ossified accessory hyoidean elements. These include a symplectic-like element associated with the anteroventral portion of the hyomandibula and in close association with the palatoquadrate and lower jaw; and a small interhyal articulating with the posterodorsal tip of the ceratohyal. ?Brachydegma, which is not a crown neopterygian, also exhibits both a symplectic and an interhyal. ';
	TEXT   TAXON = 2 CHARACTER = 223 TEXT = 'The condition in modern Chondrostei is extremely complicated. Two intermediate hyoidean elements are present, but unlike in other extant ray-fins these are serially arranged. Dorsally, a hypertrophied element connects the hyomandibula with the quadrate and lower jaw, and might partially ossify in very large sturgeons (Hilton et al 2011). This element has been variably homologized with the symplectic or the interhyal. A second, smaller, element suspends the ventral portion of the hyoid arch from the former hypertrophied element, but does not ossify. This element has been identified as either an interhyal, or a posterior ceratohyal. Given the fact that only one of the two elements ossify, we code Acipenser as 0.';
	TEXT   TAXON = 4 CHARACTER = 223 TEXT = the_interhyal_remains_cartilaginous;
	TEXT   TAXON = 58 CHARACTER = 223 TEXT = 'changed from 1 to 0, following the description in Schaeffer and Patterson 1984';
	SU  T = 58 C = 223 N = color I = 5;

	TEXT   CHARACTER = 224 TEXT = 'Giles et al. 2017; López-Arbarello & Sferco 2018.)';
	TEXT   TAXON = 2 CHARACTER = 224 TEXT = 'changed from - to ?, given the uncertainty regarding the hypertrophic element variably identified as symplectic and interhyal';
	SU  T = 2 C = 224 N = color I = 5;
	TEXT   TAXON = 7 CHARACTER = 224 TEXT = 'changed from - to ?. Nielsen 1949 figures an accessory element, which he calls symplectic, but the identity of which needs to be clarified';
	SU  T = 7 C = 224 N = color I = 5;
	TEXT   TAXON = 9 CHARACTER = 224 TEXT = 'changed from - to ?, given the uncertainty regarding the identity of preserved accessory hyoidean oddifications';
	SU  T = 9 C = 224 N = color I = 5;
	TEXT   TAXON = 10 CHARACTER = 224 TEXT = 'changed from - to ?, due to the uncertain identity of the "symplectic" element described by Nielsen 1949:Fig 72.';
	SU  T = 10 C = 224 N = color I = 5;
	TEXT   TAXON = 12 CHARACTER = 224 TEXT = 'changed from - to ?, given the uncertainty regarding the identity of preserved accessory hyoidean oddifications';
	SU  T = 12 C = 224 N = color I = 5;
	TEXT   TAXON = 13 CHARACTER = 224 TEXT = 'changed from - to ?, given the uncertainty regarding the identity of preserved accessory hyoidean oddifications';
	SU  T = 13 C = 224 N = color I = 5;
	TEXT   TAXON = 18 CHARACTER = 224 TEXT = 'changed from - to ?, given the uncertainty regarding the identity of preserved accessory hyoidean oddifications';
	SU  T = 18 C = 224 N = color I = 5;
	TEXT   TAXON = 41 CHARACTER = 224 TEXT = 'changed from - to ?, given the uncertainty regarding the identity of preserved accessory hyoidean oddifications';
	TEXT   TAXON = 42 CHARACTER = 224 TEXT = 'changed from - to ?, as these elements are now preserved in this taxon';
	TEXT   TAXON = 51 CHARACTER = 224 TEXT = 'changed from - to ?, since accessory hyoidean elements are not preserved in the specimens described in Giles et al. 2017';
	SU  T = 51 C = 224 N = color I = 5;
	SU  T = 54 C = 224 N = color I = 5;
	SU  T = 57 C = 224 N = color I = 5;
	TEXT   TAXON = 62 CHARACTER = 224 TEXT = 'chenged from - to ?, since the accessory hyoid elements are unknown in this taxon';
	SU  T = 62 C = 224 N = color I = 5;
	TEXT   TAXON = 72 CHARACTER = 224 TEXT = 'changed from - to ?, given the uncertainty regarding the identity of preserved accessory hyoidean oddifications';
	SU  T = 72 C = 224 N = color I = 5;
	TEXT   TAXON = 76 CHARACTER = 224 TEXT = 'changed from - to ?, given the uncertainty regarding the identity of preserved accessory hyoidean oddifications';
	SU  T = 76 C = 224 N = color I = 5;
	TEXT   TAXON = 77 CHARACTER = 224 TEXT = 'changed from - to ?, given the uncertainty regarding the identity of preserved accessory hyoidean oddifications';
	SU  T = 77 C = 224 N = color I = 5;
	TEXT   TAXON = 78 CHARACTER = 224 TEXT = 'changed from - to ?, given the uncertainty regarding the identity of preserved accessory hyoidean oddifications';
	SU  T = 78 C = 224 N = color I = 5;
	TEXT   TAXON = 93 CHARACTER = 224 TEXT = 'changed from - to ?, given the fact that the preserved accessory hyoidean element is likely a symplectic, rather than an interhyal';
	SU  T = 93 C = 224 N = color I = 5;
	TEXT   TAXON = 94 CHARACTER = 224 TEXT = 'changed from - to ?';
	SU  T = 94 C = 224 N = color I = 5;

	TEXT   CHARACTER = 226 TEXT = 'Friedman & Brazeau, 2010; Brazeau & Friedman, 2014; Giles et al., 2015b,c; Giles et al. 2017.';
	TEXT   TAXON = 56 CHARACTER = 226 TEXT = 'two pairs: dors & vent';
	TEXT   TAXON = 58 CHARACTER = 226 TEXT = single_hypohyal;

	TEXT   CHARACTER = 227 TEXT = 'Davis et al., 2012; Zhu et al., 2013; Giles et al., 2015b,c; Giles et al. 2017; López-Arbarello & Sferco 2018.';

	TEXT   CHARACTER = 228 TEXT = 'Giles et al., 2015b,c; Giles et al. 2017.';

	TEXT   CHARACTER = 229 TEXT = Giles_et_al._2017.;
	TEXT   TAXON = 10 CHARACTER = 229 TEXT = 'Poorly preserved: only know first three arches';
	SU  T = 117 C = 229 N = color I = 16;

	TEXT   CHARACTER = 230 TEXT = 'Grande, 2010; Xu & Wu, 2012; Xu et al., 2014, 2015; Xu & Zhao, 2016; Giles et al. 2017; López-Arbarello & Sferco 2018. ';
	TEXT   TAXON = 2 CHARACTER = 230 TEXT = 'changed from 0 to 0/1 as 4th hypobranchials are variably present sensu Hilton et al. 2011';
	SU  T = 2 C = 230 N = color I = 5;

	TEXT   CHARACTER = 231 TEXT = 'Coates, 1999; Xu & Gao, 2011; Xu et al., 2014, 2015; Poyato-Ariza 2015; Xu & Zhao, 2016; Giles et al. 2017; López-Arbarello & Sferco 2018. An uncinate process is a dorsally-directed extension on the epibranchial that articulates with the pharyngobranchial skeleton. ';

	TEXT   CHARACTER = 232 TEXT = 'Friedman, 2007; Friedman & Brazeau, 2010; Giles et al., 2015b,c; Poyato-Ariza 2015; Giles et al. 2017; López-Arbarello & Sferco 2018.';

	TEXT   CHARACTER = 233 TEXT = 'Arratia 2013; Giles et al. 2017.';

	TEXT   CHARACTER = 234 TEXT = 'Patterson, 1982; Gardiner, 1984; Gardiner & Schaeffer, 1989; Taverne, 1997; Lund, 2000; Schultze & Cumbaa, 2001; Zhu & Schultze, 2001; Zhu et al., 2001; Lund & Poplin, 2002; Zhu & Yu, 2002; Cloutier & Arratia, 2004; Gardiner et al., 2005; Friedman & Blom, 2006; Zhu et al., 2006; Friedman, 2007; Long et al., 2008; Swartz, 2009; Zhu et al., 2009; Choo, 2011; Xu & Gao, 2011; Zhu et al., 2013; Xu et al., 2014; Giles et al., 2015b; Giles et al. 2017; López-Arbarello & Sferco 2018. An elongate bone termed the ''anocleithrum'' is variably present in Wendichthys (Lund & Poplin, 1997) in the position occupied by the presupracleithrum in other taxa. We regard this as a positional homologue, and code the taxon ''0/1'' to reflect this polymorphism. Coded as ''?'' in C. trailli following arguments in Friedman & Blom (2006).';
	TEXT   TAXON = 74 CHARACTER = 234 TEXT = ?_to_1;
	SU  T = 74 C = 234 N = color I = 5;

	TEXT   CHARACTER = 235 TEXT = 'Xu et al. 2014; Giles et al. 2017.';
	TEXT   TAXON = 74 CHARACTER = 235 TEXT = ?_to_0;
	SU  T = 74 C = 235 N = color I = 5;

	TEXT   CHARACTER = 236 TEXT = 'Cloutier & Ahlberg, 1996; Schultze & Cumbaa, 2001; Zhu & Schultze, 2001; Zhu et al., 2001; Zhu & Yu, 2002; Cloutier & Arratia, 2004; Zhu et al., 2006; Friedman, 2007; Zhu et al., 2009; Giles et al., 2015b,c; Giles et al. 2017.';
	SU  T = 117 C = 236 N = color I = 5;

	TEXT   CHARACTER = 237 TEXT = 'Cavin & Suteethorn 2006; Poyato-Ariza 2015; Giles et al. 2017; López-Arbarello & Sferco 2018.';
	TEXT   TAXON = 74 CHARACTER = 237 TEXT = ?_to_0;
	SU  T = 74 C = 237 N = color I = 5;
	SU  T = 117 C = 237 N = color I = 5;

	TEXT   CHARACTER = 238 TEXT = 'Gardiner & Schaeffer, 1989; Lund et al., 1995; Cloutier & Ahlberg, 1996; Dietze, 2000; Poplin & Lund, 2000; Schultze & Cumbaa, 2001; Zhu & Schultze, 2001; Zhu et al., 2001; Zhu & Yu, 2002; Cloutier & Arratia, 2004; Zhu et al., 2006; Friedman, 2007; Zhu et al., 2009; Zhu et al., 2013; Giles et al., 2015b; Giles et al. 2017; López-Arbarello & Sferco 2018.';
	TEXT   TAXON = 35 CHARACTER = 238 TEXT = two;

	TEXT   CHARACTER = 239 TEXT = 'Patterson 1973; Olsen & McCune 1991; Coates 1999; Xu & Gao 2011; Xu et al. 2014; Poyato-Ariza 2015; Xu & Zhao, 2016; Giles et al. 2017; López-Arbarello & Sferco 2018.';
	TEXT   TAXON = 86 CHARACTER = 239 TEXT = changed_from_1_to_?._not_preserved_in_the_specimens_described_by_Bürgin_1992.;
	SU  T = 86 C = 239 N = color I = 5;

	TEXT   CHARACTER = 240 TEXT = 'Arratia 2013; Giles et al. 2017; López-Arbarello & Sferco 2018. The serrated organ (or appendage) is a small, elongate element, typically ornamented with serrated ridges, present near the anterior margin of the cleithrum. ';
	TEXT   TAXON = 74 CHARACTER = 240 TEXT = ?_to_0;
	SU  T = 74 C = 240 N = color I = 5;
	SU  T = 117 C = 240 N = color I = 16;

	TEXT   CHARACTER = 241 TEXT = 'Cloutier & Arratia, 2004; Giles et al. 2017; Xu et al., 2014. ';

	TEXT   CHARACTER = 242 TEXT = 'Olsen & McCune 1991; Zhu & Schultze, 2001; Zhu et al., 2001; Zhu & Yu, 2002; Zhu et al., 2006; Friedman, 2007; Zhu et al., 2009; Xu & Gao, 2011; Zhu et al., 2013; Xu et al., 2014; Giles et al., 2015b; Giles et al. 2017.';
	TEXT   TAXON = 74 CHARACTER = 242 TEXT = ?_to_1;
	SU  T = 74 C = 242 N = color I = 5;
	TEXT   TAXON = 97 CHARACTER = 242 TEXT = changed_from_0_to_?_to_account_for_the_poor_preservation_of_the_specimens_figured_in_Stensio_1925;
	SU  T = 97 C = 242 N = color I = 5;

	TEXT   CHARACTER = 243 TEXT = 'Patterson, 1982; Gardiner, 1984; Gardiner & Schaeffer, 1989; Rosen, 1989; Taverne, 1997; Coates, 1999; Zhu & Schultze, 2001; Zhu et al., 2001; Zhu & Yu, 2002; Zhu et al., 2006; Brazeau, 2009; Zhu et al., 2009; Friedman & Brazeau, 2010; Xu & Gao, 2011; Davis et al., 2012; Zhu et al., 2013; Xu et al., 2014; Giles et al., 2015b,c.';
	TEXT   TAXON = 74 CHARACTER = 243 TEXT = ?_to_0;
	SU  T = 74 C = 243 N = color I = 5;

	TEXT   CHARACTER = 244 TEXT = 'Patterson, 1982; Gardiner, 1984; Gardiner & Schaeffer, 1989; Taverne, 1997; Coates, 1999; Schultze & Cumbaa, 2001; Zhu & Schultze, 2001; Friedman & Blom, 2006; Long et al., 2008; Swartz, 2009; Choo, 2011; Xu & Gao, 2011; Giles et al., 2015b; Giles et al. 2017. ';
	TEXT   TAXON = 10 CHARACTER = 244 TEXT = 'changed from - to ?, as it is not clear in Nielsen 1949';
	SU  T = 10 C = 244 N = color I = 5;
	TEXT   TAXON = 74 CHARACTER = 244 TEXT = ?_to_1;
	SU  T = 74 C = 244 N = color I = 5;

	TEXT   CHARACTER = 245 TEXT = 'Giles et al. 2017; López-Arbarello & Sferco 2018.';
	TEXT   TAXON = 10 CHARACTER = 245 TEXT = 'changed from - to ?, as it is not clear in Nielsen 1949';
	SU  T = 10 C = 245 N = color I = 5;
	TEXT   TAXON = 74 CHARACTER = 245 TEXT = ?_to_0;
	SU  T = 74 C = 245 N = color I = 5;
	TEXT   TAXON = 118 CHARACTER = 245 TEXT = 'not described in original description, but present in NHMD 75388';
	TEXT   TAXON = 123 CHARACTER = 245 TEXT = 'not described in original description, but present in NHMD 75388';

	TEXT   CHARACTER = 246 TEXT = 'Taverne, 1997; Coates, 1999; Friedman & Blom, 2006; Long et al., 2008; Swartz, 2009; Xu & Gao, 2011; Xu et al., 2014; Giles et al., 2015b; Giles et al. 2017.';
	TEXT   TAXON = 74 CHARACTER = 246 TEXT = ?_to_0;
	SU  T = 74 C = 246 N = color I = 5;

	TEXT   CHARACTER = 247 TEXT = 'Zhu & Yu, 2002; Friedman, 2007; Giles et al., 2015b; Giles et al. 2017. Two series of pectoral fin radials are described (but not figured) for Cheirolepis candensis (Arratia & Cloutier, 2004). Although we consider this arrangement to be unlikely, for now this taxon is coded ''1''.  ';
	TEXT   TAXON = 74 CHARACTER = 247 TEXT = ?_to_0;
	SU  T = 74 C = 247 N = color I = 5;

	TEXT   CHARACTER = 248 TEXT = 'Zhu & Schultze, 2001; Zhu et al., 2001; Zhu & Yu, 2002; Zhu et al., 2006; Friedman, 2007; Zhu et al., 2009; Friedman & Brazeau, 2010; Zhu et al., 2013; Giles et al., 2015b,c; Giles et al. 2017. ';
	TEXT   TAXON = 74 CHARACTER = 248 TEXT = ?_to_1;
	SU  T = 74 C = 248 N = color I = 5;

	TEXT   CHARACTER = 249 TEXT = 'Coates, 1999; Friedman & Blom, 2006; Long et al., 2008; Choo, 2011; Xu & Gao, 2011; Xu et al., 2014; Giles et al., 2015b; Giles et al. 2017.';
	TEXT   TAXON = 10 CHARACTER = 249 TEXT = 'changed from 3 to 1 based on Nielsen 1949: Pl. 19, 20';
	SU  T = 10 C = 249 N = color I = 5;

	TEXT   CHARACTER = 250 TEXT = 'Giles et al., 2015b; Giles et al. 2017.';
	TEXT   TAXON = 97 CHARACTER = 250 TEXT = changed_from_?_to_1;
	SU  T = 97 C = 250 N = color I = 5;

	TEXT   CHARACTER = 251 TEXT = 'Zhu et al., 2001; Zhu & Yu, 2002; Friedman, 2007; Zhu et al., 2009; Brazeau, 2009; Davis et al., 2012; Zhu et al., 2013; Giles et al., 2015b,c; Giles et al. 2017.';

	TEXT   CHARACTER = 252 TEXT = 'Friedman & Blom, 2006; Friedman, 2007; Brazeau, 2009; Choo, 2011; Davis et al., 2012; Zhu et al., 2013; Brazeau & Friedman, 2014; Giles et al., 2015b,c; Giles et al. 2017. ';

	TEXT   CHARACTER = 253 TEXT = 'Gardiner & Schaeffer, 1989; Coates, 1998; Coates, 1999; Lund, 2000; Schultze & Cumbaa, 2001; Cloutier & Arratia, 2004; Friedman & Blom, 2006; Zhu et al., 2006; Long et al., 2008; Swartz, 2009; Zhu et al., 2009; Choo, 2011; Xu et al., 2014; Giles et al., 2015b; Giles et al. 2017.';

	TEXT   CHARACTER = 254 TEXT = 'Friedman, 2007; Zhu & Yu, 2002; López-Arbarello 2012; Bermúdez-Rochas & Poyato-Ariza 2015; Giles et al., 2015b; Thies & Waschkewitz, 2015; Gibson 2016; Giles et al. 2017; López-Arbarello & Sferco 2018.';

	TEXT   CHARACTER = 255 TEXT = 'Olsen & McCune 1991; Lund, 2000; Poplin & Lund, 2000; Poyato-Ariza & Wenz, 2002; Cloutier & Arratia, 2004; Friedman & Blom, 2006; Long et al., 2008; Swartz, 2009; Choo, 2011; Giles et al., 2015b; Thies & Waschkewitz, 2015; Gibson 2016; Giles et al. 2017; Cawley & Kriwet 2018; López-Arbarello & Sferco 2018.';

	TEXT   CHARACTER = 256 TEXT = 'Patterson, 1982; Taverne, 1997; Friedman & Blom, 2006; Long et al., 2008; Choo, 2011; Giles et al., 2015b; Giles et al. 2017; López-Arbarello & Sferco 2018.';

	TEXT   CHARACTER = 257 TEXT = 'Cloutier & Arratia, 2004; Friedman & Blom, 2006; Long et al., 2008; Swartz, 2009; Choo, 2011; Giles et al., 2015b; Giles et al. 2017.';

	TEXT   CHARACTER = 258 TEXT = 'Gardiner & Schaeffer, 1989; Schultze & Cumbaa, 2001; Zhu & Schultze, 2001; Zhu et al., 2001; Zhu & Yu, 2002; Cloutier & Arratia, 2004; Friedman & Blom, 2006; Zhu et al., 2006; Friedman, 2007; Long et al., 2008; Brazeau, 2009; Swartz, 2009; Zhu et al., 2009; Choo, 2011; Davis et al., 2012; Zhu et al., 2013; Giles et al., 2015b,c; Giles et al. 2017.';

	TEXT   CHARACTER = 259 TEXT = 'Poplin & Lund, 2000; Cloutier & Arratia, 2004; Friedman & Blom, 2006; Long et al., 2008; Swartz, 2009; Choo, 2011; López-Arbarello 2012; Bermúdez-Rochas & Poyato-Ariza 2015; Giles et al., 2015b; Thies & Waschkewitz, 2015; Gibson 2016; Giles et al. 2017; López-Arbarello & Sferco 2018.';

	TEXT   CHARACTER = 260 TEXT = 'Gardiner et al., 2005; Xu & Gao, 2011; Xu et al., 2014, 2015; Xu & Zhao, 2016; Giles et al. 2017; López-Arbarello & Sferco 2018.';

	TEXT   CHARACTER = 261 TEXT = Giles_et_al._2017.;

	TEXT   CHARACTER = 262 TEXT = Giles_et_al._2017.;

	TEXT   CHARACTER = 263 TEXT = 'Patterson, 1982; Cloutier & Ahlberg, 1996; Coates, 1999; Schultze & Cumbaa, 2001; Zhu & Schultze, 2001; Friedman & Blom, 2006; Long et al., 2008; Swartz, 2009; Choo, 2011; Giles et al., 2015b; Giles et al. 2017.';

	TEXT   CHARACTER = 264 TEXT = 'Patterson, 1982; Taverne, 1997; Gardiner & Schaeffer, 1989; Gardiner et al., 2005; Friedman & Blom, 2006; Long et al., 2008; Choo, 2011; López-Arbarello 2012; Bermúdez-Rochas & Poyato-Ariza 2015; Giles et al., 2015b; Giles et al. 2017; López-Arbarello & Sferco 2018. ';

	TEXT   CHARACTER = 265 TEXT = 'Modified from Gardiner et al., 2005; Giles et al. 2017. A long chordal lobe is considered to be present when the notochord reaches the posterior margin of the caudal fin.';
	TEXT   TAXON = 86 CHARACTER = 265 TEXT = 'revised from 0 to 1, sensu Bürgin 1990, 1992';
	SU  T = 86 C = 265 N = color I = 5;

	TEXT   CHARACTER = 266 TEXT = 'Grande & Bemis 1998; Xu & Gao 2011; Xu et al. 2014; Xu & Zhao, 2016; Giles et al. 2017; López-Arbarello & Sferco 2018.';

	TEXT   CHARACTER = 267 TEXT = 'Arratia 2013. Most Paleozoic and early Mesozoic non-neopterygian actinopterygians and many teleosts bear strong posterolaterally expanding epineural processes on lateral surfaces of their neural arches.';
	TEXT   TAXON = 79 CHARACTER = 267 TEXT = 'Gardiner (1984) mentions the similarity of arcual elements between Mimipiscis and Moythomasia, but he does not figure the latter.';
	TEXT   TAXON = 118 CHARACTER = 267 TEXT = 'not described in original description, but present in NHMD 75388';
	TEXT   TAXON = 123 CHARACTER = 267 TEXT = 'not described in original description, but present in NHMD 75388';

	TEXT   CHARACTER = 268 TEXT = 'Grande & Bemis 1998; Arratia 2013; Poyato-Ariza 2015; Giles et al. 2017; López-Arbarello & Sferco 2018.';

	TEXT   CHARACTER = 269 TEXT = 'Coates 1999; Hurley et al. 2007; Xu et al. 2014; Giles et al. 2017; López-Arbarello & Sferco 2018.';

	TEXT   CHARACTER = 270 TEXT = 'Pinna, 1996; Hurley et al., 2007; Xu & Wu, 2012, Xu et al., 2014; Xu & Zhao, 2016; Giles et al. 2017; López-Arbarello & Sferco 2018.';
	TEXT   TAXON = 11 CHARACTER = 270 TEXT = changed_from_?_to_0;
	SU  T = 11 C = 270 N = color I = 5;

	TEXT   CHARACTER = 271 TEXT = 'Pinna, 1996, Xu & Wu, 2012, Xu et al., 2014; Xu & Zhao, 2016; Giles et al. 2017.';
	TEXT   TAXON = 11 CHARACTER = 271 TEXT = changed_from_?_to_0;
	SU  T = 11 C = 271 N = color I = 5;

	TEXT   CHARACTER = 272 TEXT = 'Giles et al. 2017; López-Arbarello & Sferco 2018.';
	TEXT   TAXON = 11 CHARACTER = 272 TEXT = changed_from_?_to_0;
	SU  T = 11 C = 272 N = color I = 5;

	TEXT   CHARACTER = 273 TEXT = 'Wiley, 1976; López-Arbarello 2012; Bermúdez-Rochas & Poyato-Ariza 2015; Thies & Waschkewitz, 2015; Poyato-Ariza 2015; Gibson 2016; Giles et al. 2017; López-Arbarello & Sferco 2018. Non applicable in taxa without ossified centra';
	TEXT   TAXON = 10 CHARACTER = 273 TEXT = 'changed from 0 to -';
	SU  T = 10 C = 273 N = color I = 5;
	TEXT   TAXON = 11 CHARACTER = 273 TEXT = 'changed from ? to -';
	SU  T = 11 C = 273 N = color I = 5;
	TEXT   TAXON = 13 CHARACTER = 273 TEXT = 'changed from 0 to - as the notochord is unconstricted';
	SU  T = 13 C = 273 N = color I = 5;
	TEXT   TAXON = 15 CHARACTER = 273 TEXT = 'changed from 0 to - as the notochord is unconstricted';
	SU  T = 15 C = 273 N = color I = 5;
	TEXT   TAXON = 48 CHARACTER = 273 TEXT = 'changed from 0 to - as the notochord is unconstricted';
	SU  T = 48 C = 273 N = color I = 5;
	TEXT   TAXON = 65 CHARACTER = 273 TEXT = changed_from_0_to_?_as_vertebral_morphology_is_now_well_known_in_this_taxon;
	SU  T = 65 C = 273 N = color I = 5;
	TEXT   TAXON = 76 CHARACTER = 273 TEXT = 'changed from 0 to - as the notochord is unconstricted';
	SU  T = 76 C = 273 N = color I = 5;
	TEXT   TAXON = 77 CHARACTER = 273 TEXT = 'changed from 0 to - as the notochord is unconstricted';
	SU  T = 77 C = 273 N = color I = 5;
	TEXT   TAXON = 78 CHARACTER = 273 TEXT = 'changed from 0 to - as the notochord is unconstricted';
	SU  T = 78 C = 273 N = color I = 5;
	TEXT   TAXON = 82 CHARACTER = 273 TEXT = changed_from_0_to_?_as_vertebral_morphology_is_now_well_known_in_this_taxon;
	TEXT   TAXON = 88 CHARACTER = 273 TEXT = 'changed from 0 to - as the notochord is unconstricted';
	SU  T = 88 C = 273 N = color I = 5;
	TEXT   TAXON = 93 CHARACTER = 273 TEXT = 'to change from 0 to - as the notochord is unconstricted';
	SU  T = 93 C = 273 N = color I = 8;
	TEXT   TAXON = 111 CHARACTER = 273 TEXT = 'changed from 0 to - notochord is unconstricted';
	SU  T = 111 C = 273 N = color I = 5;
	TEXT   TAXON = 112 CHARACTER = 273 TEXT = 'changed from 0 to - notochord is unconstricted';
	SU  T = 112 C = 273 N = color I = 5;
	TEXT   TAXON = 113 CHARACTER = 273 TEXT = 'changed from 0 to - notochord is unconstricted^n';
	SU  T = 113 C = 273 N = color I = 5;
	TEXT   TAXON = 115 CHARACTER = 273 TEXT = changed_from_0_to_?;
	SU  T = 115 C = 273 N = color I = 5;

	TEXT   CHARACTER = 274 TEXT = Giles_et_al._2017.;

	TEXT   CHARACTER = 275 TEXT = 'López-Arbarello 2012; Thies & Waschkewitz, 2015; Gibson 2016; López-Arbarello & Sferco 2018.)';

	TEXT   CHARACTER = 276 TEXT = 'The infraorbitals normally form as a single row partially surrounding the orbit. In some taxa, they may be fragmented into multiple irregular ossifications.';

	TEXT   CHARACTER = 277 TEXT = 'Grande & Bemis 1998; López-Arbarello 2011; López-Arbarello & Sferco 2018. The suborbitals normally form as a single row partially surrounding the orbit, posterior and/or ventral to the infraorbitals. In some taxa, they may be fragmented into multiple irregular ossifications.';

	TEXT   CHARACTER = 278 TEXT = 'López-Arbarello 2011; López-Arbarello & Sferco 2018';

	TEXT   CHARACTER = 279 TEXT = 'Coronoids are normally restricted to the medial (lingual) surface of the jaw. In some dapediids, they may extend laterally to contribute to the labial tooth row.';
	TEXT   TAXON = 5 CHARACTER = 279 TEXT = check;

	TEXT   CHARACTER = 280 TEXT = 'López-Arbarello 2012; Bermúdez-Rochas & Poyato-Ariza 2015; Poyato-Ariza 2015; Thies & Waschkewitz, 2015; Gibson 2016; López-Arbarello & Sferco 2018.';
	TEXT   TAXON = 43 CHARACTER = 280 TEXT = hmm;
	TEXT   TAXON = 112 CHARACTER = 280 TEXT = anterior;

	TEXT   CHARACTER = 281 TEXT = 'The parasphenoid is normally restricted to covering the ventral face of the braincase. In some taxa, extensions may extend dorsally and laterally to cloak the basiocciput.';

	TEXT   CHARACTER = 282 TEXT = 'The parasphenoid extends dorsally between the orbits, contributing to the interorbital septum, in some taxa.';

	TEXT   CHARACTER = 283 TEXT = 'A deep ligament pit, dorsal to the foramen magnum, is found on the posterior face of the braincase in some taxa.';

	TEXT   CHARACTER = 284 TEXT = Gibson_2016.;
	TEXT   TAXON = 27 CHARACTER = 284 TEXT = present;

	TEXT   TAXON = 24 CHARACTER = 286 TEXT = check;

	TEXT   CHARACTER = 287 TEXT = 'Poyato-Ariza & Wenz, 2002; Cawley & Kriwet 2018.)';

	TEXT   CHARACTER = 288 TEXT = 'Poyato-Ariza & Wenz, 2002; Thies & Waschkewitz, 2015; Gibson 2016; Cawley & Kriwet 2018; López-Arbarello & Sferco 2018. The supratemporal commissure is normally carried by the extrascapulars, but is sometimes borne in the parietal.';
	TEXT   TAXON = 5 CHARACTER = 288 TEXT = '0	';
	TEXT   TAXON = 27 CHARACTER = 288 TEXT = long_mpl;
	TEXT   TAXON = 29 CHARACTER = 288 TEXT = long_mpl;
	TEXT   TAXON = 36 CHARACTER = 288 TEXT = 'travels onto lateral extrascapular then onto parietal (no median exscp)';
	TEXT   TAXON = 104 CHARACTER = 288 TEXT = 'travels onto lateral extrascapular then onto parietal (no median exscp)';

	TEXT   CHARACTER = 289 TEXT = 'Poyato-Ariza & Wenz, 2002; Cawley & Kriwet 2018; López-Arbarello & Sferco 2018.';
	TEXT   TAXON = 4 CHARACTER = 289 TEXT = conical;
	TEXT   TAXON = 15 CHARACTER = 289 TEXT = conical;
	SU  T = 117 C = 289 N = color I = 16;

	TEXT   CHARACTER = 290 TEXT = 'Poyato-Ariza & Wenz, 2002; Cawley & Kriwet 2018.';
	TEXT   TAXON = 5 CHARACTER = 290 TEXT = 'borne on ?fused coronoids+prearticulars';
	SU  T = 117 C = 290 N = color I = 16;

	TEXT   CHARACTER = 291 TEXT = 'Nursall 2010.) ';
	SU  T = 117 C = 291 N = color I = 16;

	TEXT   CHARACTER = 292 TEXT = 'Poyato-Ariza & Wenz, 2002; Cawley & Kriwet 2018. A median bone, typically referred to as a dermal supraoccipital, is present between the parietals in some taxa.';

	TEXT   CHARACTER = 293 TEXT = 'Poyato-Ariza & Wenz, 2002; Cawley & Kriwet 2018.';

	TEXT   CHARACTER = 294 TEXT = 'Poyato-Ariza & Wenz, 2002; Cawley & Kriwet 2018.';
	TEXT   TAXON = 95 CHARACTER = 294 TEXT = 'changed from 0 to 1 following the reinterpretation of the saurichthyid opercle (Argyriou et al. 2018)';
	SU  T = 95 C = 294 N = color I = 5;

	TEXT   CHARACTER = 295 TEXT = 'Poyato-Ariza & Wenz, 2002; Cawley & Kriwet 2018.';

	TEXT   CHARACTER = 297 TEXT = Nursall_2010.;

	TEXT   CHARACTER = 298 TEXT = 'Poyato-Ariza & Wenz, 2002; Cawley & Kriwet 2018.';
	SU  T = 117 C = 298 N = color I = 16;

	TEXT   CHARACTER = 299 TEXT = 'Poyato-Ariza & Wenz, 2002; Cawley & Kriwet 2018.';

	TEXT   CHARACTER = 300 TEXT = Nursall_2010.;

	TEXT   CHARACTER = 179 STATE = 0 TEXT = 'From Argyriou et al. 2018. In saurichthyids and likely Yelangichthys the common carotids bifurcate after piercing the parasphenoid, at the level of the ascending processes. In some sturgeons the parasphenoid can be pierced by the descending 1st and 2nd efferent branchial arteries, but a circulus cephalicus is absent (Findeis 1993). Inapplicable In taxa like most stem teleosts where there are two foramina piercing the parasphenoid, but the arrangement of their corresponding grooves on the dorsal part of the parasphenoid suggests that they carried the internal carotid (posterior foramen) and pseudobranchial (anterior foramen) arteries (Patterson 1975, fig. 142). This pattern is reconstructed for many fossil neopterygians (Patterson 1975).';
	TEXT   CHARACTER = 187 STATE = 0 TEXT = 'Modified from Wu et al. 2013, ch.13. Applicable only in taxa that do not exhibit a single olfactory tract.';
	TEXT   CHARACTER = 201 STATE = 1 TEXT = From_Argyriou_et_al._2018;
	TEXT   CHARACTER = 202 STATE = 0 TEXT = From_Argyriou_et_al._2018;
	TEXT   CHARACTER = 225 STATE = 1 TEXT = Arratia_2013.;
	TEXT   CHARACTER = 272 STATE = 0 TEXT = single;

	AN T = 37 C = 3  A = 0 DC = 2017.1.15.15.23.58 DM = 2017.1.15.15.23.58 ID = 0159a2ba59981 I = _ TF = (CM ) ;

END;

BEGIN MESQUITECHARMODELS;
	ProbModelSet * UNTITLED   =  'Mk1 (est.)':  1- 300;
END;

Begin MESQUITE;
		MESQUITESCRIPTVERSION 2;
		TITLE AUTO;
		tell ProjectCoordinator;
		timeSaved 1702898646189;
		getEmployee #mesquite.minimal.ManageTaxa.ManageTaxa;
		tell It;
			setID 0 8638558614273783985;
			tell It;
				setDefaultOrder  63 75 70 91 93 107 80 69 77 73 10 82 0 13 85 1 2 3 71 61 4 5 68 101 26 96 28 35 29 30 31 32 33 34 36 37 6 135 48 85 74 7 95 75 96 59 61 26 71 110 69 46 45 8 47 83 10 92 106 65 11 13 15 79 103 16 17 78 87 107 90 49 18 19 50 20 21 22 23 24 102 51 39 52 58 86 133 88 60 53 103 54 40 41 133 96 97 72 90 55 107 99 42 106 103 80 43 44 109 110 111 112 113 114 118 115 116 117 118 119 120 121 122;
				attachments ;
			endTell;
		endTell;
		getEmployee #mesquite.charMatrices.ManageCharacters.ManageCharacters;
		tell It;
			setID 0 2279276808437032474;
			tell It;
				setDefaultOrder  0 1 243 2 267 3 4 225 233 266 211 251 5 6 265 7 8 280 210 9 20 10 11 12 36 13 14 15 16 235 17 18 342 341 20 21 28 23 24 27 29 184 284 30 31 32 33 34 35 37 207 38 39 40 41 42 25 26 185 261 262 236 43 44 45 46 48 56 57 59 60 224 187 214 215 186 62 63 64 65 225 66 67 68 69 345 70 240 71 75 72 73 74 76 259 244 218 77 344 80 343 277 279 273 79 275 239 230 49 111 50 218 51 345 61 346 282 236 192 52 231 53 54 55 81 82 83 84 255 85 86 87 88 89 90 91 92 93 94 95 96 97 187 98 99 100 101 102 103 104 105 153 106 107 108 109 111 256 236 224 225 231 200 208 209 210 168 210 247 175 176 177 112 113 114 115 116 240 180 264 265 247 178 117 118 119 188 120 121 193 122 123 194 195 196 124 125 126 127 261 202 203 139 140 141 142 253 143 144 145 146 147 250 148 149 131 132 133 128 129 130 136 225 265 226 137 135 138 244 209 186 134 237 153 189 150 266 151 190 240 210 158 154 155 259 159 157 160 162 161 163 164 165 169 170 171 172 173 174 191 331 332 166 168 273 192 268 239 213 214 215 273 234 271 321 280 281 284 278 320 287 313 288 277 314 282 285 294 289 290 301 292 304 295 296 291 301 297 298 302 300 301 302;
				attachments ;
			endTell;
			mqVersion 361;
			checksumv 0 3 1388748793 null  getNumChars 303 numChars 303 getNumTaxa 123 numTaxa 123   short true   bits 2305843009213693983   states 31   sumSquaresStatesOnly 53321.0 sumSquares -8.762203435012037E19 longCompressibleToShort false usingShortMatrix true   NumFiles 1 NumMatrices 1;
			mqVersion;
		endTell;
		getWindow;
		tell It;
			suppress;
			setResourcesState false false 100;
			setPopoutState 300;
			setExplanationSize 0;
			setAnnotationSize 0;
			setFontIncAnnot 0;
			setFontIncExp 0;
			setSize 1440 811;
			setLocation 0 25;
			setFont SanSerif;
			setFontSize 10;
			getToolPalette;
			tell It;
			endTell;
			desuppress;
		endTell;
		getEmployee  #mesquite.minimal.ManageTaxa.ManageTaxa;
		tell It;
			showTaxa #8638558614273783985 #mesquite.lists.TaxonList.TaxonList;
			tell It;
				setTaxa #8638558614273783985;
				getWindow;
				tell It;
					useTargetValue off;
					setTargetValue ;
					newAssistant  #mesquite.lists.TaxonListCurrPartition.TaxonListCurrPartition;
					setExplanationSize 30;
					setAnnotationSize 20;
					setFontIncAnnot 0;
					setFontIncExp 0;
					setSize 1340 739;
					setLocation 0 25;
					setFont SanSerif;
					setFontSize 10;
					getToolPalette;
					tell It;
						setTool mesquite.lists.TaxonList.TaxonListWindow.arrow;
					endTell;
				endTell;
				showWindow;
				getEmployee #mesquite.lists.ColorTaxon.ColorTaxon;
				tell It;
					setColor Red;
					removeColor off;
				endTell;
				getEmployee #mesquite.lists.TaxonListAnnotPanel.TaxonListAnnotPanel;
				tell It;
					togglePanel off;
				endTell;
			endTell;
		endTell;
		getEmployee  #mesquite.charMatrices.BasicDataWindowCoord.BasicDataWindowCoord;
		tell It;
			showDataWindow #2279276808437032474 #mesquite.charMatrices.BasicDataWindowMaker.BasicDataWindowMaker;
			tell It;
				getWindow;
				tell It;
					getTable;
					tell It;
						rowNamesWidth 132;
					endTell;
					setExplanationSize 50;
					setAnnotationSize 80;
					setFontIncAnnot 0;
					setFontIncExp 0;
					setSize 1340 659;
					setLocation 0 25;
					setFont SanSerif;
					setFontSize 10;
					getToolPalette;
					tell It;
						setTool mesquite.charMatrices.BasicDataWindowMaker.BasicDataWindow.arrow;
					endTell;
					setActive;
					setTool mesquite.charMatrices.BasicDataWindowMaker.BasicDataWindow.arrow;
					colorCells  #mesquite.charMatrices.ColorCells.ColorCells;
				tell It;
					setColor Red;
					removeColor off;
				endTell;
					colorRowNames  #mesquite.charMatrices.TaxonGroupColor.TaxonGroupColor;
					colorColumnNames  #mesquite.charMatrices.CharGroupColor.CharGroupColor;
					colorText  #mesquite.charMatrices.NoColor.NoColor;
					setBackground White;
					toggleShowNames off;
					toggleShowTaxonNames on;
					toggleTight off;
					toggleThinRows off;
					toggleShowChanges on;
					toggleSeparateLines off;
					toggleShowStates on;
					toggleReduceCellBorders off;
					toggleAutoWCharNames on;
					toggleAutoTaxonNames off;
					toggleShowDefaultCharNames off;
					toggleConstrainCW on;
					toggleBirdsEye off;
					toggleShowPaleGrid off;
					toggleShowPaleCellColors off;
					toggleShowPaleExcluded off;
					togglePaleInapplicable on;
					togglePaleMissing off;
					toggleShowBoldCellText off;
					toggleAllowAutosize on;
					toggleColorsPanel on;
					toggleDiagonal on;
					setDiagonalHeight 80;
					toggleLinkedScrolling on;
					toggleScrollLinkedTables off;
				endTell;
				showWindow;
				getWindow;
				tell It;
					forceAutosize;
				endTell;
				getEmployee #mesquite.charMatrices.AlterData.AlterData;
				tell It;
					toggleBySubmenus off;
				endTell;
				getEmployee #mesquite.charMatrices.ColorByState.ColorByState;
				tell It;
					setStateLimit 9;
					toggleUniformMaximum on;
				endTell;
				getEmployee #mesquite.categ.StateNamesEditor.StateNamesEditor;
				tell It;
					makeWindow;
					tell It;
						getTable;
						tell It;
							rowNamesWidth 232;
						endTell;
						setExplanationSize 50;
						setAnnotationSize 40;
						setFontIncAnnot 0;
						setFontIncExp 0;
						setSize 1340 699;
						setLocation 0 25;
						setFont SanSerif;
						setFontSize 10;
						getToolPalette;
						tell It;
							setTool mesquite.categ.StateNamesEditor.StateNamesWindow.arrow;
						endTell;
						rowsAreCharacters on;
						toggleConstrainChar on;
						toggleConstrainCharNum 3;
						togglePanel off;
						toggleSummaryPanel off;
					endTell;
					showWindow;
				endTell;
				getEmployee #mesquite.categ.StateNamesStrip.StateNamesStrip;
				tell It;
					showStrip off;
				endTell;
				getEmployee #mesquite.charMatrices.AnnotPanel.AnnotPanel;
				tell It;
					togglePanel off;
				endTell;
				getEmployee #mesquite.charMatrices.CharReferenceStrip.CharReferenceStrip;
				tell It;
					showStrip off;
				endTell;
				getEmployee #mesquite.charMatrices.QuickKeySelector.QuickKeySelector;
				tell It;
					autotabOff;
				endTell;
				getEmployee #mesquite.charMatrices.SelSummaryStrip.SelSummaryStrip;
				tell It;
					showStrip off;
				endTell;
				getEmployee #mesquite.categ.SmallStateNamesEditor.SmallStateNamesEditor;
				tell It;
					panelOpen true;
				endTell;
			endTell;
		endTell;
		getEmployee  #mesquite.charMatrices.ManageCharacters.ManageCharacters;
		tell It;
			showCharacters #2279276808437032474 #mesquite.lists.CharacterList.CharacterList;
			tell It;
				setData 0;
				getWindow;
				tell It;
					useTargetValue off;
					setTargetValue ;
					newAssistant  #mesquite.lists.DefaultCharOrder.DefaultCharOrder;
					newAssistant  #mesquite.lists.CharListInclusion.CharListInclusion;
					newAssistant  #mesquite.lists.CharListPartition.CharListPartition;
					newAssistant  #mesquite.parsimony.CharListParsModels.CharListParsModels;
					setExplanationSize 30;
					setAnnotationSize 20;
					setFontIncAnnot 0;
					setFontIncExp 0;
					setSize 1340 739;
					setLocation 0 25;
					setFont SanSerif;
					setFontSize 10;
					getToolPalette;
					tell It;
						setTool mesquite.lists.CharacterList.CharacterListWindow.arrow;
					endTell;
				endTell;
				showWindow;
				getEmployee #mesquite.lists.CharListAnnotPanel.CharListAnnotPanel;
				tell It;
					togglePanel off;
				endTell;
			endTell;
		endTell;
		endTell;
end;
