## Supplementary material phylogenetic analyses for "A Portrait of a Young Fish: Redescription of *Pteronisculus gunnari* (Nielsen, 1942) from a juvenile specimen from the Early Triassic of East Greenland, with implications for ontogenetic development in early actinopterygians": changes_to_phylogenetic_matrix.pdf

**1) General notes.** This matrix is based on that of Argyriou et al. (2022), with 3 additional characters and 6 additional taxa. These taxa were added to the analysis in order to understand the relationships among the *Pteronisculus* genus. These comprise NHMD–73588, *Pteronisculus gunnari*, *Pteronisculus magnus* (Nielsen, 1942), *Pteronisculus changae* (Ren & Xu, 2021), *Pteronisculus nielseni* (Xu et al., 2014) and *Pteronisculus cicatrosus* (Lehman, 1952). The characters added were sourced from the character matrix of Ren & Xu (2021). The present matrix comprises 303 characters and 123 taxa.

**2) Character codes updated from Argyriou et al. (2022).**

*Pteronisculus stensioi*:

char. 37: 1 to 0;

char. 38: - to 0;

char. 39: 0 to 1;

char. 40: 1 to 0;

char. 41: 0 to 1;

char. 66: 0 to 1;

char. 68: 0 to 1;

char. 111: 1 to 0;

char. 127: 1 to 0;

char. 166: 0 to ?;

char. 224: ? to 0;

char. 225: ? to 1;

char. 231: 0 to 1;

char. 249: 0 to 1;

char. 281: 1 to 0.

**3) New characters List**

**301: Lacrimal contributing to oral margin:** 0, absent; 1, present.

**302 Teeth on lacrimal:** 0, absent; 1, present.

**303 Dermosphenotic position:** 0, extending well below dermopterotic; 1, located at same horizontal level than dermopterotic.
