## Supplementary material for "A Portrait of a Young Fish: Redescription of *Pteronisculus gunnari* (Nielsen, 1942) from a juvenile specimen from the Early Triassic of East Greenland, with implications for ontogenetic development in early actinopterygians": Figure S1

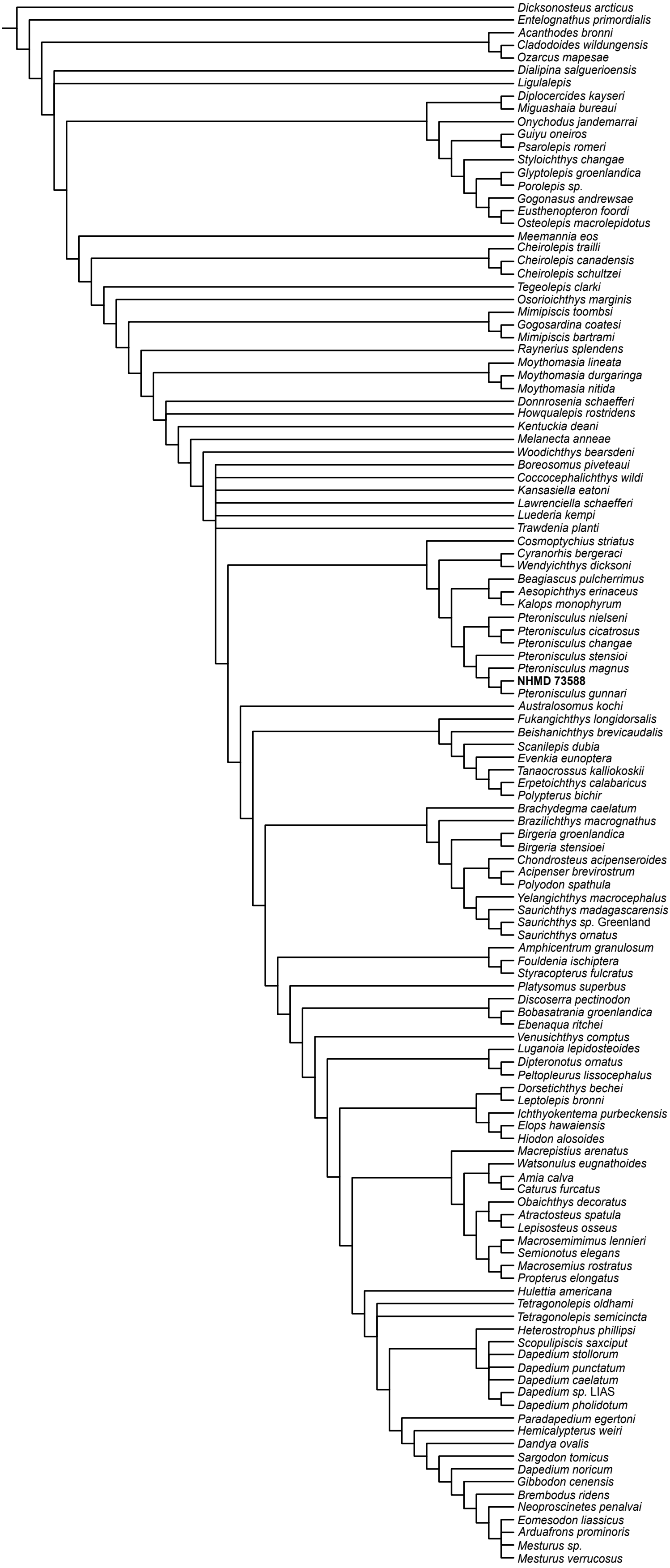


**Figure S1**: strict consensus tree of 135 most parsimonious trees, implied weights analysis (K=100) with outgroup topology contrained.
